# Wing musculature reconstruction in extinct flightless auks (*Pinguinus* and *Mancalla*) reveals incomplete convergence with penguins (Spheniscidae) due to differing ancestral states

**DOI:** 10.1101/2020.07.22.215707

**Authors:** Junya Watanabe, Daniel J. Field, Hiroshige Matsuoka

## Abstract

Despite longstanding interest in convergent evolution, factors that result in deviations from fully convergent phenotypes remain poorly understood. In birds, the evolution of flightless wing-propelled diving has emerged as a classic example of convergence, having arisen in disparate lineages including penguins (Sphenisciformes) and auks (Pan-Alcidae, Charadriiformes). Nevertheless, little is known about the functional anatomy of the wings of flightless auks because all such taxa are extinct, and their morphology is almost exclusively represented by skeletal remains. Here, in order to re-evaluate the extent of evolutionary convergence among flightless wing-propelled divers, wing muscles and ligaments were reconstructed in two extinct flightless auks, representing independent transitions to flightlessness: *Pinguinus impennis* (a crown-group alcid), and *Mancalla* (a stem-group alcid). Extensive anatomical data were gathered from dissections of 12 species of extant charadriiforms and 4 aequornithine waterbirds including a penguin. It was found that the wings of both flightless auk taxa were characterized by an increased mechanical advantage of wing elevator/retractor muscles, and decreased mobility of distal wing joints, both of which are likely advantageous for wing-propelled diving and parallel similar functional specializations in penguins. However, the conformations of individual muscles and ligaments underlying these specializations differ markedly between penguins and flightless auks, instead resembling those in each respective group’s close relatives. Thus, the wings of these flightless wing-propelled divers can be described as convergent as overall functional units, but are incompletely convergent at lower levels of anatomical organization—a result of retaining differing conditions from each group’s respective volant ancestors. Detailed investigations such as this one may indicate that, even in the face of similar functional demands, courses of phenotypic evolution are dictated to an important degree by ancestral starting points.

## Introduction

Convergent evolution, in which distantly-related lineages acquire similar traits, has been regarded as evidence for the predictability of organismal evolution under natural selection (e.g., Conway Morris 2003, 2010; Melville et al. 2006; Mahler et al. 2013). Convergence may arise as a result of a tight relationship between phenotype and functional performance, and/or evolutionary constraints or biases inherent to certain organismal designs that result in a limitation of possible phenotypic solutions (Wake 1991; Losos 2011; Wake et al. 2011). These factors often operate simultaneously, and may lead to idiosyncratic outcomes because of differences in ancestral conditions and/or evolvability between lineages (historical contingency; e.g., Gould 2002; Agrawal 2017; Blount et al. 2018). Recent studies have demonstrated that idiosyncrasies among lineages occupying similar niches, termed “incomplete” convergence (sensu Herrel et al. 2004), might be more prevalent than previously recognized (Losos 2010; Moen et al. 2016; Hulsey et al. 2019). As such, close examination into the nature of apparently convergent phenotypes and their ancestral conditions is required to fully comprehend the various evolutionary processes underlying convergence.

The evolution of avian wing-propelled diving provides a classic example of convergent evolution. Wing-propelled diving describes a mode of underwater locomotion whereby birds propel themselves by flapping their forelimbs (aquatic flight; Townsend 1909; Storer 1960). This locomotor mode has arisen independently in multiple avian lineages: penguins (Sphenisciformes; throughout the paper, “Spheniscidae” is reserved for crown-group penguins while Sphenisciformes applies to the total group); auks (Pan-Alcidae [=Mancallinae + crown-group Alcidae], Charadriiformes); diving petrels (*Pelecanoides*, Procellariidae, Procellariiformes); dippers (Cinclidae, Passeriformes); and extinct plotopterids (Plotopteridae, Suliformes). Additionally, some petrels and shearwaters (Procellariidae), gannets (Sulidae), and certain waterfowl (Anatidae) are known to use their wings, sometimes along with their feet, in underwater movement (e.g., Townsend 1909; Kuroda 1954; Storer 1960; Ashmole 1971).

Along with the independent acquisition of wing-propelled diving, penguins, great auks (*Pinguinus*, Alcidae), mancalline auks (Mancallinae, Pan-Alcidae), and plotopterids lost the capacity for aerial flight. Whereas the diversity of volant wing-propelled divers encapsulates a continuous spectrum between casual and dedicated divers, flightless wing-propelled divers are best regarded as occupying a distinct adaptive zone unto themselves (Simpson 1946; Livezey 1989), characterized by distinct morphological and ecological specializations.

Extant penguins are the product of a long, independent evolutionary history as flightless wing-propelled divers. With the technical exception of the extant Galápagos penguin (*Spheniscus mendiculus*), all extant and fossil penguin species are known from the Southern Hemisphere, and their oldest known fossil record dates to the Paleocene (>60 Ma; e.g., Slack et al. 2006; Ksepka and Ando, 2011; Blockland et al. 2019). Great auks are a lineage of crown-group Alcidae known from the North Atlantic. The only Recent representative of the lineage, *Pinguinus impennis*, became extinct in the 19th century as a consequence of human exploitation (Lucas 1890; Fuller 1999). The only known prehistoric member of the lineage is *Pinguinus alfrednewtoni*, known from isolated fossil bones from the Pliocene (∼4.4 Ma) of North Carolina (Olson 1977; Olson and Rasmussen 2001). Mancalline auks are an extinct lineage of flightless auks representing the sister group to crown-group Alcidae (Smith 2011). Two genera are currently recognized: *Miomancalla* is known from the Miocene–Pliocene of California (∼10–4.9 Ma), and *Mancalla* is known from the Pliocene (perhaps extending into the Miocene) – Pleistocene of the Pacific coasts of North America and Japan (∼5.0–0.12 Ma; e.g., Lucas 1901; Miller and Howard 1949; Chandler 1990; Smith 2011; Smith and Clarke 2015; Watanabe et al. 2020a, 2020b). Plotopteridae is an extinct lineage of Suliformes (although there is some dispute about their exact phylogenetic position; Smith 2010; Mayr et al. 2015, 2020a). Eight genera have been described from the upper Eocene – middle Miocene of the Pacific coast of North America and Japan (∼35–17 Ma; e.g., Howard 1969; Olson and Hasegawa 1985, 1996; Sakurai et al. 2008; Mayr and Goedart 2016, 2018).

Because sea water is ∼800 times denser than air (Pennycuick 1987; Vogel 1994), aquatic flight imposes different functional demands on the wing than does aerial flight. In water, the downward force of gravity is largely offset by the buoyancy of water. At the same time, resistance and drag against movement is much greater in water than in air. Several morphological attributes have been ascribed to wing-propelled diving: relatively small wings characterized by shortened bones and flight feathers which induce less drag and provide increased rigidity (Pennycuick 1987; Livezey 1988, 1989; Louw 1992); dorsoventrally flattened wing bones with thick cortices apparently providing hydrodynamic efficiency and resistance to bending stress (Stettenheim 1959; Habib and Ruff 2008; Habib 2010; Smith and Clarke 2014); well-developed wing elevator muscles presumably involved in active upstroke of the wings in a dense, viscous medium (Stettenheim 1959; Schreiweis 1982; Bannasch 1986b, 1994; Kovacs and Meyers, 2000); and reduced mobility of the elbow, wrist, and digital joints providing increased rigidity to the wings (specifically noted in penguins, but absent in volant auks; Bannasch 1986a, 1994; Raikow et al. 1988; Louw 1992). The specialized, rigid wings of penguins are often referred to as flippers, analogous to the modified limbs of other secondarily aquatic tetrapods (e.g., Thewissen and Taylor 2007; Kelley and Pyenson 2015; DeBlois and Motani 2019).

During aquatic flight in extant penguins and volant auks, upstroke of the wings produces substantial forward thrust (propulsive force) (e.g., Clark and Bemis 1979; Johansson and Wetterholm Aldrin 2002; Watanuki et al. 2006; Lapsansky and Tobalske 2019). This contrasts with aerial flight in birds, in which generation of forward thrust is typically restricted to the downstroke phase (Rayner 1988). Active thrust generated by the upstroke during aquatic flight is presumably facilitated by well-developed wing elevator muscles, enabling energetically efficient swimming at relatively steady speeds (Lovvorn 2001).

Storer (1960) once posited parallels in the evolutionary history of Alcidae in the Northern Hemisphere and Sphenisciformes + Procellariiformes in the Southern Hemisphere, hypothesizing a similar three-stage transition toward wing-propelled diving in both lineages; that is, 1) wings used for aerial flight only (exemplified by extant non-diving taxa), 2) wings used for both aquatic and aerial flight (exemplified by extant volant alcids and diving petrels), and 3) wings used for aquatic flight only (exemplified by extant penguins, and the extinct great and mancalline auks). Storer (1960) additionally hypothesized that morphological features of the wings of birds in the second “stage” reflect a compromise between the differing demands of aquatic and aerial flight, which should favor smaller and larger wing areas for reduced drag and reduced wing loading, respectively. However, although some sort of evolutionary trade-off probably exists regarding wing area/loading in volant alcids (Thaxter et al. 2010), empirical measurements of diving parameters during extensive flight feather molt indicate that a smaller wing area on its own does not improve diving performance in volant alcids (Bridge 2004). In addition, neither the joint mobility (Raikow et al. 1988) nor muscle histochemistry (Kovacs and Meyers 2000) of volant alcids clearly exemplify a compromise or intermediate condition between non-diving birds and penguins. Indeed, even though volant alcids and diving petrels were regarded as representatives of the same “stage” in the evolution of wing-propelled diving in Storer’s scheme, they exhibit striking osteological differences (Kuroda 1967; Harrison 1977). Therefore, the simplistic typology of Storer (1960) does not adequately encapsulate the evolutionary history of wing-propelled diving in Pan-Alcidae and Sphenisciformes, demanding close examination of the extent of convergence in these groups.

One obstacle to studying the evolution of wing-propelled diving in birds is that many key taxa—stem penguins, plotopterids, and mancalline auks—are extinct, and known only from fossilized bones. Only in very exceptional circumstances have remnants of feathers and skin been recovered from fossil penguins (Clarke et al. 2010; Acosta Hospitaleche et al. 2020). Hence, data available for investigations into the convergence of flightless wing-propelled divers are mostly restricted to skeletal elements. This holds largely true even for great auks, which became extinct in the 19th Century before much was learned about their anatomy (Lucas 1890; Fuller 1999). Despite much work on various morphological aspects of these extinct wing-propelled divers, including morphometrics (Livezey 1988, 1989), limb bone histology (Smith and Clarke 2014; Ksepka et al. 2015), feeding morphology (Haidr and Acosta Hospitaleche 2012, 2014; Degrange et al. 2018; Chávez-Hoffmeister 2020), and neuroanatomy (Ksepka et al. 2012b; Smith and Clarke 2012; Kawabe et al. 2014; Tambussi et al. 2015; Proffitt et al. 2016), surprisingly little is known about the musculoskeletal anatomy of the wings in extinct wing-propelled diving birds, perhaps with the exception of specific aspects of the musculature in stem penguins (Acosta Hospitaleche and Di Carlo 2012; Haidr and Acosta Hospitaleche 2019).

This study reconstructs the wing musculature of the extinct flightless auks *Pinguinus* and *Mancalla*, and undertakes thorough comparisons with extant charadriiforms and aequornithine waterbirds in order to explore the evolution of wing-propelled diving from a detailed anatomical perspective. These reconstructions draw on osteological correlates observable in fossil and subfossil bones, evaluated through dissection of extant relatives and application of the Extant Phylogenetic Bracket (Witmer 1995). In short, the Extant Phylogenetic Bracket is a framework enabling justified inferences about soft parts in extinct organisms known only from fossilized hard parts (e.g., bones in vertebrates), based on hypothesized homological correspondence between soft and hard parts (so-called osteological correlates), inferred under a phylogenetic hypothesis (see also Bryant and Russell 1992; Witmer 1997). Although limitations of this methodology exist (e.g., Bryant and Seymour 1990; Hutchinson 2001a, 2001b), it provides a means of testing hypotheses regarding soft parts in extinct organisms that are not directly observable.

Although a good number of mounted skins of recently extinct *Pinguinus impennis* have been preserved (Fuller 1999), the irreplaceable nature of these skins prohibits attempts at direct observation of remnant musculature via dissection or contrast-enhanced computed tomography (Lautenschlager et al. 2014; Gignac et al. 2016). Hence, reconstructing soft parts through well-justified osteological correlates provides the only feasible means of investigating the gross topological features of wing musculature in this species (although, as noted later, a small number of skeletal specimens are preserved with partial remnants of associated soft parts). Fortunately, an adequate number of well-preserved bones are known for this species, as well as for the extinct taxon *Mancalla*, to enable reliable identification of osteological correlates in these groups. A comprehensive reconstruction of the wing musculature of these taxa was developed, based on original anatomical data obtained from extant representatives of Charadriiformes, and close interrogation of osteological correlates and application of the Extant Phylogenetic Bracket. This reconstructed musculature for extinct flightless auks was subsequently compared with anatomical data on the wing musculature of extant penguins and their relatives, in order to identify similarities and differences in the wings of these evolutionarily independent examples of flightless wing-propelled divers. Importantly, observations from penguins were not consulted during reconstruction of the musculature of extinct auks, in order to avoid logical circularity in the identification of convergent features in these groups.

## Materials and Methods

### Taxon sampling

In order to reconstruct wing musculature in extinct pan-alcids, anatomical information was gathered from dissection of extant alcids and other charadriiform birds. For extant alcids, 17 individuals representing 7 species were examined (Table 1). Although taxonomic sampling was limited by availability of specimens, the sample covers a substantial portion of genus-level diversity of extant Alcidae (6 out of 9 extant genera). The sample also covers a large part of the body size spectrum of extant alcids, ranging from *Synthliboramphus antiquus* to *Uria lomvia* (∼200 g to ∼950 g, respectively; Gaston and Jones 1998), with only some members of *Aethia*, *Ptychoramphus*, and *Synthliboramphus* falling clearly outside the lower end of this range, and *Alle alle* and *Uria aalge* overlapping with the lower and upper margins of the range, respectively. Other groups of Charadriiformes were also sampled to cover major extant subclades, and to place Mancallinae within an extant phylogenetic bracket: 11 individuals of 5 species were examined, representing Charadriidae, Scolopacidae, Laridae, and Stercorariidae (Table 1). The phylogenetic framework generally follows the family-level relationships inferred by Prum et al. (2015), and for detailed relationships within Charadriiformes, the phylogenetic relationships of Smith and Clarke (2015) were followed. The relevant aspects of the topologies of these two phylogenies are consistent (Fig. 1).

**Figure 1.**
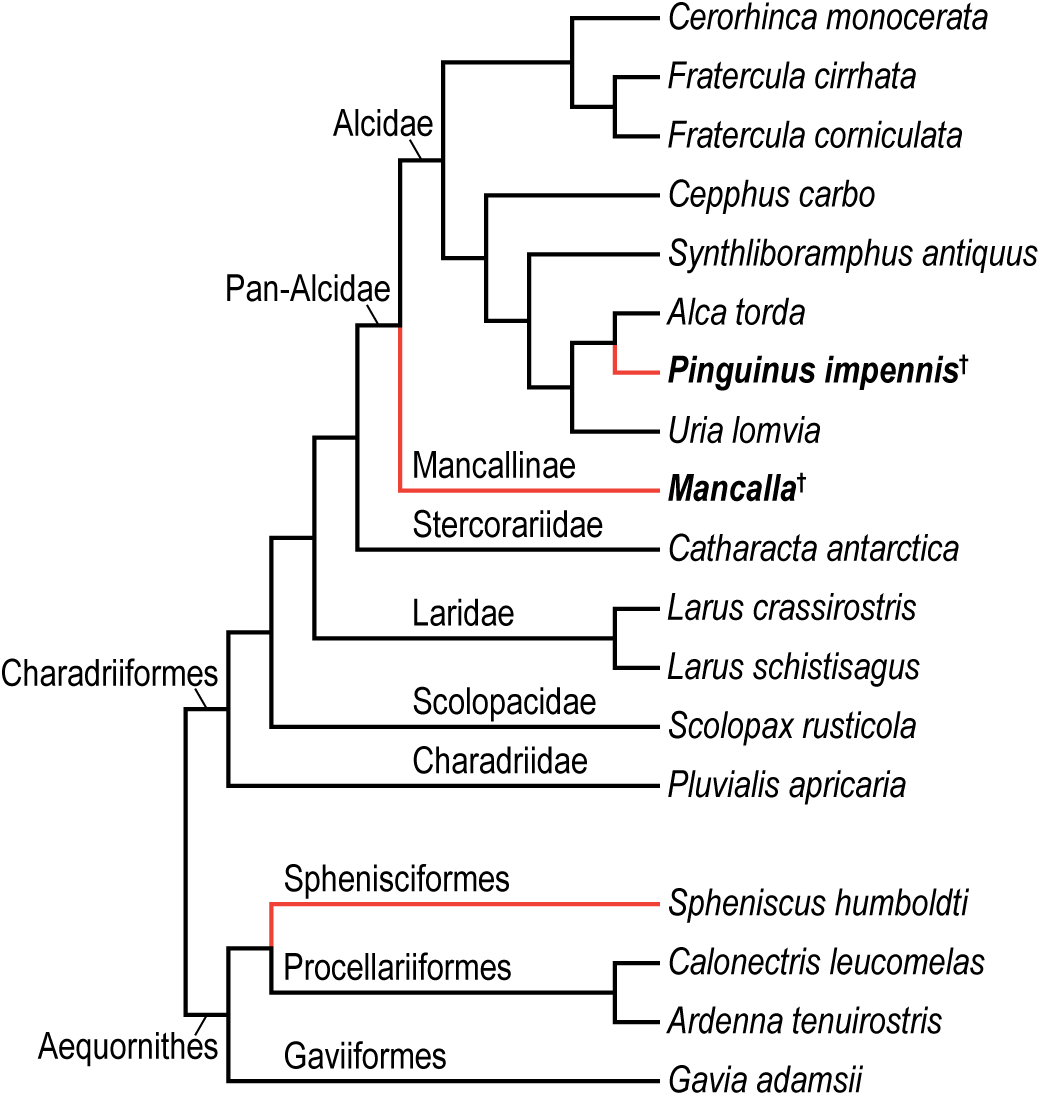
Working phylogeny. Simplified from the family-level relationships of Prum et al. (2015), supplemented by Smith and Clarke (2015) for the relationships within Charadriiformes. The two focal taxa of flightless auks are shown with boldface. Daggers denote extinct taxa, and orange branches denote flightless wing-propelled diving lineages (flightless auks and penguins).

**Table 1.**
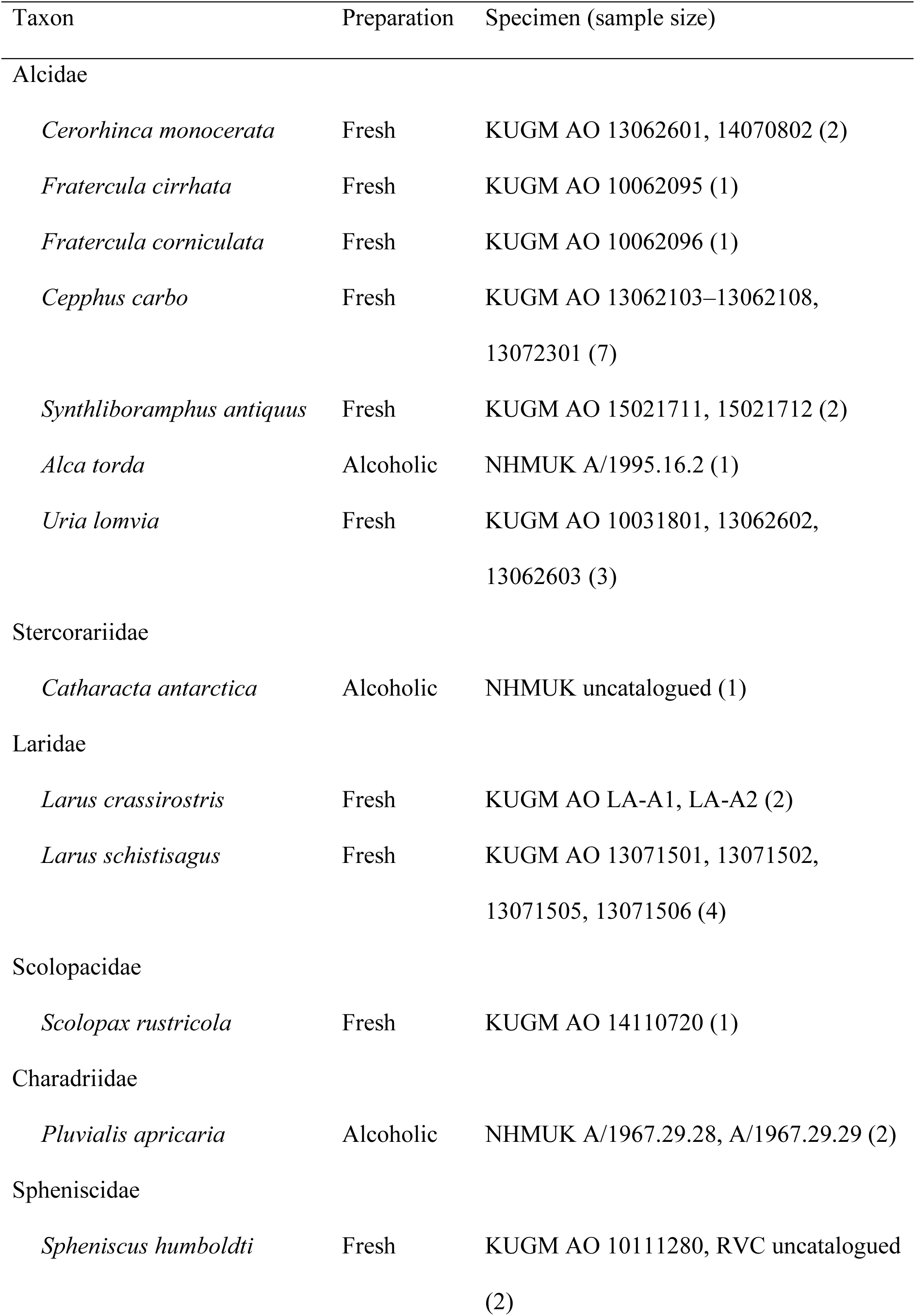
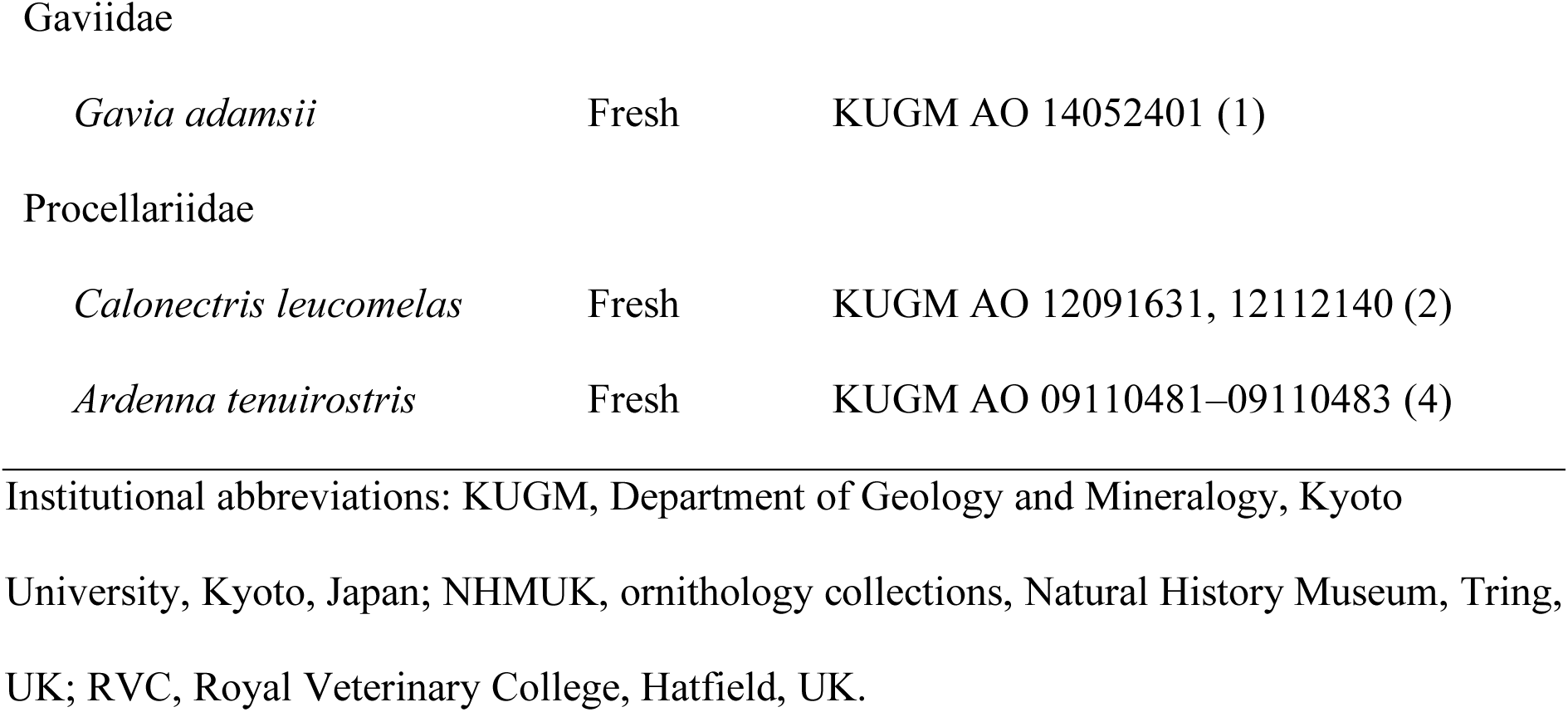
List of the extant taxa and specimens examined.

Reconstructed musculature of the extinct auks was subsequently compared with that of extant Sphenisciformes and their close relatives (Procellariiformes and Gaviiformes). For this purpose, 2 individuals of *Spheniscus humboldti* (Sphenisciformes) and 7 individuals representing 3 species of Procellariiformes and Gaviiformes were examined (Table 1; Fig. 1). Because one of the *Spheniscus humboldti* specimens examined was a chick (with natal down retained on the entire body), results for this species should be viewed cautiously. However, the observations did not differ markedly between chick and adult individuals, nor from what has been previously described (Schreiweis 1982; Bannasch 1986b); thus, the new observations appear to be valid for the purposes of this investigation.

### Dissection of modern specimens

Dissections were made on unfixed carcasses and spirit specimens. Unfixed specimens were obtained through salvaging dead wild individuals, rescued individuals that subsequently died, victims of by-catch by research vessels, or individuals that died in captivity (in the case of *Spheniscus humboldti*). In the case of *Larus crassirostris*, specimens previously collected in the wild for another project (Watanabe 2018a, 2018b, 2018c) were examined. No animals were killed or captured for this project, and all specimens were legally obtained and transferred under local regulations. The unfixed carcasses were stored frozen, and thawed overnight at room temperature prior to dissection. The skin and viscera were removed after taking external measurements, and detailed dissection was conducted on one randomly selected wing per individual, assuming bilateral symmetry. Flight feathers and coverts were removed either before or during dissection of the wing; hence, some muscles and ligaments (e.g., m. expansor secondariorum) connected to the feathers were not observed for all individuals. Nevertheless, such muscles and ligaments generally do not exhibit osteological correlates and therefore could not be included in the present reconstructions of extinct taxa even if they were present. Most of the unfixed specimens examined were subsequently prepared as skeletal specimens, and are stored in the Department of Geology and Mineralogy, Kyoto University, Kyoto, Japan, or the Royal Veterinary College, Hatfield, UK.

The fixed, spirit (alcoholic) specimens examined in this study are stored in the anatomical collections of the Natural History Museum, Tring, UK. For these specimens, only one side of the wings and thorax were skinned and dissected. Subsequent to dissection, muscles removed from these bodies were individually labelled and stored together with the main specimens.

Muscles and ligaments on the dissected wings were removed one after another, while recording the detailed positions of their attachment sites. Descriptions in the literature (Stettenheim 1959; Hudson et al. 1969; Schreiweis 1982; McKitrick 1991; Bannasch 1994; Kovacs and Meyers 2000) were consulted for identification of muscles, but the descriptions presented here are based entirely on original observations. Osteological correlates were identified either on the specimens dissected or on additional skeletal specimens, and the positions and extent of attachment sites were recorded as precisely as possible during dissection.

### (Sub)fossil specimens and reconstruction of soft parts

Most known skeletal specimens of *Pinguinus impennis* are subfossil bones that were collected after the species had been driven to extinction. Therefore, associated skeletons of this species are vanishingly rare in museum collections (Livezey 1988). As a result, any attempt to reconstruct the musculature of this species will inevitably rely on isolated bones from multiple individuals. Nevertheless, given little intraspecific variation in the relative positions of osteological correlates within the extant charadriiforms examined, the composite nature of the *Pinguinus* specimens is unlikely to affect the qualitative inferences drawn in this study. Subfossil bones of *Pinguinus impennis* from the collections of the Museum of Zoology, University of Cambridge, Cambridge, UK (UMZC 187.d and 187.G) were the primary source of osteological data for the musculature reconstruction for this species.

Thousands of fossil specimens of Mancallinae are available in museum collections, from which several species of *Mancalla* have been described (e.g., Chandler 1990; Smith 2011). Nevertheless, no single specimen represents a sufficient component of the pectoral girdle and wing skeleton to enable reconstruction of the complete musculature of the wing. This situation necessitated that the reconstructions be based on observations of multiple specimens. Several associated, partial skeletons were available in the extensive collections of fossil birds at the Natural History Museum of Los Angeles County, Los Angeles, and the San Diego Museum of Natural History, San Diego (both California, USA). These constituted the primary basis of the reconstructions for *Mancalla*. In addition, other well-preserved specimens were also examined to complement observations on these associated skeletons (Table 2). Many *Mancalla* specimens could not be identified to species level due to a lack of diagnostic features, and they may represent multiple species. Nevertheless, only negligible qualitative variation was observed in the relative positions of osteological correlates. Thus, these specimens were collectively treated as representatives of a single taxon (*Mancalla*) for the purpose of wing musculature reconstruction. Although this is admittedly a coarse assumption that may overlook potential interspecific variation, this approach was necessary in the light of a lack of complete skeletons. Some of the specimens had originally been identified as belonging to species that were considered invalid by Smith (2011), but no attempt was made to re-identify them to species level, apart from confirming their assignment to *Mancalla*.

**Table 2.** Fossil specimens of *Mancalla* primarily consulted in this study. For associated skeletons, only elements of the pectoral girdle and wing skeleton are listed.

| Specimen | Identification | Locality | Element |
| --- | --- | --- | --- |
| LACM 15373 | *Mancalla cedrosensis*<br>(holotype) | A | R coracoid, R scapula, R humerus, ulnae, radii, carpometacarpi |
| LACM 15410 | *Mancalla cedrosensis* | A | Incomplete sternum, furcula, coracoids, scapulae, L humerus, L carpometacarpus |
| LACM 56259 | *Mancalla* sp. | C | Incomplete L coracoid, R scapula, R humerus, L carpometacarpus |
| SDSNH 21295 | *Mancalla* sp. | SD | Incomplete sternum, furcula, L coracoid |
| SDSNH 25237 | *Mancalla lucasi*<br>(holotype) | N | Scapulae, humeri |
| SDSNH 77966 | *Mancalla* sp. | SD | Incomplete R humerus, R ulna, R carpometacarpus |
| SDSNH 26242 | *Mancalla* “*emlongi*” | SD | Incomplete sternum |
| SDSNH 27292 | *Mancalla* sp. | SD | Incomplete sternum |
| SDSNH 21021 | *Mancalla* sp. | SD | L coracoid |
| SDSNH 22844 | *Mancalla* sp. | SD | L scapula |
| SDSNH 24983 | *Mancalla* “*diegensis*” | SD | L humerus |

Table 2 (continued)
| Specimen | Identification | Locality | Element |
| --- | --- | --- | --- |
| SDSNH 32760 | *Mancalla* sp. | SD | L humerus |
| SDSNH 21454 | *Mancalla* “*milleri*” | SD | L ulna |
| SDSNH 24991 | *Mancalla* sp. | SD | R ulna |
| SDSNH 71922 | *Mancalla* sp. | SD | L ulna |
| SDSNH 77268 | *Mancalla* sp. | SD | L ulna |
| SDSNH 71927 | *Mancalla* sp. | SD | R radius |
| SDSNH 126338 | *Mancalla* sp. | SD | R radius |
| SDSNH 23758 | *Mancalla* “*milleri*” | SD | L carpometacarpus |
| SDSNH 40969 | *Mancalla* sp. | SD | L carpometacarpus |
| SDSNH 59051 | *Mancalla* sp. | SD | L carpometacarpus |

L and R denote left and right sides, respectively. Abbreviations for localities: A, Almejas Formation, Cedros Island, Mexico; C, Capistrano Formation, California, USA; N, Niguel Formation, California, USA; SD, San Diego Formation, California, USA. Institutional abbreviations: LACM, vertebrate paleontology collections, Natural History Museum of Los Angeles County, Los Angeles, California, USA; SDSNH, vertebrate paleontology collections, San Diego Museum of Natural History, San Diego, California, USA.

There was no complete sternum for Mancallinae available for the present study, nor have any been reported in the literature. As a result, the sternal morphology of *Mancalla* needed to be reconstructed from multiple specimens. This reconstruction was accomplished by photographic collage of three well-preserved partial sterna; photographs of two specimens (LACM 2180 and SDSNH 77399), taken in lateral and ventral views, were overlaid onto photographs of another specimen (SDSNH 26242), with the former ones rescaled such that the outlines of their preserved portions matched those of the latter as closely as possible, while retaining their original aspect ratios. This procedure may have introduced some inaccuracies in scaling into the reconstruction of this portion of the skeleton, as the specimens involved differed distinctly in size.

Osteological correlates on the bones of the extinct species were identified by comparison with those in extant species, based on their shape, nature (e.g., tubercles, scars, lines), and positions relative to other landmarks. In most cases, the presence of muscles and ligaments could be inferred by “Level I” inferences of the Extant Phylogenetic Bracket framework (Witmer 1995, 1997). That is, the presence of a muscle/ligament in an extinct species was inferred based on the presence of the corresponding osteological correlate in that species and the conserved relationship between the soft parts and the osteological correlate in at least two extant species that phylogenetically “bracket” the extinct species. In some cases, however, only weaker inferences could be made, for which specific notes are given below.

For *Pinguinus impennis*, a dried partial skeleton with remnants of the elbow and forearm musculature (NHMUK 1972.1.156) became available after the reconstruction based on osteological correlates was complete. The reconstructed musculature was subsequently compared with this desiccated specimen in order to verify the validity of the reconstruction based only on osteological correlates.

### Anatomical terminology

Anatomical terminology largely follows Nomina Anatomica Avium 2nd Edition (Baumel et al. 1993), especially those chapters regarding musculoskeletal anatomy (Baumel and Witmer 1993; Baumel and Raikow 1993; Vanden Berge and Zweers 1993). Terminological notes are given in the text as required, especially when nomenclatural inconsistency was noted, or appropriate names were not available in this publication. A list of the muscles and ligaments examined is given in Table 3. The following abbreviations are used throughout the text: artc., articulatio; lig., ligamentum/ligamenti (singular); ligg., ligamenti (plural); m., musculus/musculi (singular); mm., musculi (plural).

**Table 3.** List of ligaments and muscles examined, with abbreviations for figures.

| Name | Abbreviation | Note |
| --- | --- | --- |
| ***Ligaments of the shoulder*** |  |  |
| Lig. acrocoracohumerale | L. acr-hum. |  |
| Lig. coracohumerale dorsale | L. cor-hum. dor. |  |
| Lig. scapulohumerale dorsale | L. scap-hum. dor. |  |
| Retinaculum originis m.<br>scapulotricipitis | R. or. scaptri. | Tentative term |
| Plica synovialis transversa | P. syn. tr. |  |
| ***Ligaments of the elbow*** |  |  |
| Lig. collaterale ventrale | L. col. ven. |  |
| Lig. collaterale dorsale | L. col. dor. | See text for discussion |
| Lig. dorsale cubiti | L. dor. cub. | Tentative term; see text<br>for discussion |
| Lig. craniale cubiti | L. cran. cub. |  |
| Lig. radioulnare transversum | L. rad-uln. tr. | See text for discussion |
| Meniscus radioulnaris | Men. rad-uln. |  |
| Lig. radioulnare ventrale | L. rad-uln. ven. | Tentative term; see text<br>for discussion |
| Trochlea humeroulnaris | T. hum-uln. |  |
| Lig. tricipitale | L. tri. |  |
| ***Ligaments of the wrist*** |  |  |
| Aponeurosis ventralis | A. ven. |  |
| Lig. radioulnare interosseum | L. rad-uln. int. |  |
| Lig. ulno-ulnocarpale proximale | L. uln-uc. prox. |  |
| Lig. ulno-ulnocarpale distale | L. uln-uc. dist. |  |
| Lig. ulno-radiocarpale ventrale | L. uln-rc. ven. |  |
| Lig. ulno-radiocarpale interosseum | L. uln-rc. int. |  |
| Lig. ulno-radiocarpale dorsale | L. uln-rc. dor. | Tentative term |
| Lig. ulno-metacarpale ventrale | L. uln-met. ven. |  |
| Lig. radio-radiocarpale craniale | L. rad-rc. cran. |  |
| Lig. radio-radiocarpale ventrale | L. rad-rc. ven. |  |
| Lig. radio-radiocarpale dorsale | L. rad-rc. dor. |  |
| Meniscus intercarpalis | Men. intercar. |  |
| Lig. radiocarpo-metacarpale craniale | L. rc-met. cran. |  |
| Lig. radiocarpo-metacarpale dorsale | L. rc-met. dor. |  |
| Lig. radiocarpo-metacarpale ventrale | L. rc-met. ven. |  |
| Lig. ulnocarpo-metacarpale ventrale | L. uc-met. ven. |  |
| Lig. ulnocarpo-metacarpale dorsale | L. uc-met. dor. |  |
| Lig. ulnocarpo-metacarpale caudale | L. uc-met. caud. | Tentative term |
| Lig. ulno-metacarpale externum | L. uln-met. ext. | Tentative term after<br>Stettenheim (1959) |
| ***Ligaments of the manus*** |  |  |
| Lig. obliquum alulae | L. obl. al. |  |
| Lig. collaterale caudale [of artc.<br>metacarpophalangealis alulae] | L. col. caud. al. |  |
| Lig. collaterale dorsale [of artc.<br>metacarpophalangealis alulae] | L. col. dor. al. | Tentative term |
| Lig. collaterale ventrale [of artc.<br>metacarpophalangealis digiti<br>majoris] | L. col. ven. maj. |  |
| Lig. collaterale caudale [of artc.<br>metacarpophalangealis digiti<br>majoris] | L. col. caud. maj. |  |
| Lig. obliquum intra-articulare | L. obl. int. |  |
| Lig. collaterale ventrale [of artc.<br>metacarpophalangealis digiti<br>minoris] | L. col. ven. min. |  |
| Lig. collaterale dorsale [of artc.<br>metacarpophalangealis digiti<br>minoris] | L. col. dor. min. |  |
| Lig. collaterale caudale [of artc.<br>metacarpophalangealis digiti<br>minoris] | L. col. caud. min. | Tentative term |
| Lig. interosseum | L. inteross. |  |
| *Accessory ligaments* |  |  |
| Lig. propatagiale | L. prop. |  |
| Lig. limitans cubiti | L. lim. cub. |  |
| Lig. humerocarpale | L. hum-car. |  |
| Membrana interossea antebrachii | Mem. int. |  |
| Retinaculum m. scapulotricipitis | R. scaptri. |  |
| Retinaculum m. extensoris metacarpi<br>ulnaris | R. m. e. uln. |  |
| ***Wing muscles*** |  |  |
| M. rhomboideus superficialis | M. rhom. sup. |  |
| M. rhomboideus profundus | M. rhom. prof. |  |
| M. serratus superficialis pars cranialis | M. ser. cran. |  |
| M. serratus superficialis pars caudalis | M. ser. caud. |  |
| M. serratus superficialis pars<br>metapatagialis | M. ser. metap. |  |
| M. serratus profundus | M. ser. prof. |  |
| M. scapulohumeralis cranialis | M. scap-hum. cran. |  |
| M. scapulohumeralis caudalis | M. scap-hum. caud. |  |
| Mm. subcoracoscapulares | Mm. subcorscap. |  |
| M. subscapularis caput laterale | M. subscap. lat. |  |
| M. subscapularis caput mediale | M. subscap. med. |  |
| M. subcoracoideus | M. subcor. |  |
| M. coracobrachialis cranialis | M. cor-br. cran. |  |
| M. coracobrachialis caudalis | M. cor-br. caud. |  |
| M. pectoralis pars sternobrachialis | M. pect. ster-br. |  |
| M. pectoralis pars costobrachialis | M. pect. cost-br. |  |
| M. pectoralis pars profundus | M. pect. prof. | Tentative term after<br>Kuroda (1960, 1961) |
| M. supracoracoideus | M. supracor. |  |
| M. latissimus dorsi pars cranialis | M. lat. dor. cran. |  |
| M. latissimus dorsi pars caudalis | M. lat. dor. caud. |  |
| M. latissimus dorsi pars metapatagialis | M. lat. dor. metap. |  |
| M. deltoideus pars propatagialis | M. delt. prop. |  |
| M. deltoideus pars major | M. delt. maj. |  |
| M. deltoideus pars minor | M. delt. min. |  |
| M. deltoideus pars minor caput dorsale | M. delt. min. dor. |  |
| M. deltoideus pars minor caput ventrale | M. delt. min. ven. |  |
| M. scapulotriceps | M. scaptri. |  |
| M. humerotriceps | M. humtri. |  |
| M. biceps brachii | M. bic. |  |
| M. biceps brachii pars propatagialis | M. bic. prop. |  |
| M. brachialis | M. brach. |  |
| M. pronator superficialis | M. pron. sup. |  |
| M. pronator profundus | M. pron. prof. |  |
| M. flexor carpi ulnaris | M. f. car. uln. |  |
| M. flexor digitorum superficialis | M. f. d. sup. |  |
| M. flexor digitorum profundus | M. f. d. prof. |  |
| M. extensor carpi radialis | M. e. car. rad. |  |
| M. extensor carpi ulnaris | M. e. car. uln. |  |
| M. extensor digitorum communis | M. e. d. com. |  |
| M. extensor longus alulae | M. e. lon. al. |  |
| M. extensor longus digiti majoris | M. e. lon. d. maj. |  |
| M. extensor longus digiti majoris pars<br>proximalis | M. e. lon. d. maj.<br>prox. |  |
| M. extensor longus digiti majoris pars<br>distalis | M. e. lon. d. maj.<br>dist. |  |
| M. supinator | M. supin. |  |
| M. ectepicondylo-ulnaris | M. ect-uln. |  |
| M. ulnometacarpalis dorsalis | M. uln-met. dor. |  |
| M. ulnometacarpalis ventralis | M. uln-met. ven. |  |
| M. interosseus dorsalis | M. int. dor. |  |
| M. interosseus ventralis | M. int. ven. |  |
| M. extensor brevis alulae | M. e. br. al. |  |
| M. abductor alulae | M. abd. al. |  |
| M. flexor alulae | M. f. al. |  |
| M. adductor alulae | M. add. al. |  |
| M. abductor digiti majoris | M. abd. dig. maj. |  |
| M. flexor digiti minoris | M. f. dig. min. |  |

## Results

### Musculature in extant birds

Among the charadriiform birds examined, most wing ligaments and muscles were observed in generally consistent positions. In many cases, the attachments of ligaments and tendons (indirect attachments of muscles) corresponded to distinct tubercles or scars, which could be easily delineated. In contrast, the margins of fleshy (direct) attachments could not be clearly discerned unless delineated by intermuscular lines or other osteological landmarks, as pointed out previously (Bryant and Seymour 1990). Descriptions of major wing ligaments and muscles are given below, as well as illustrations of the overall musculature in a representative taxon (*Alca*; Figs. 2, 3), and osteological correlates in selected taxa (*Catharacta*, *Alca*, *Spheniscus*, Figs. 4–23; *Pluvialis*, *Scolopax*, *Larus schistisagus*, *Cerorhinca*, *Cepphus*, *Synthliboramphus*, *Uria*, *Gavia*, *Ardenna*, Supplementary Material Figs. S1–S32). Results for *Larus crassirostris*, *Fratercula*, and *Calonectris* were mostly similar to those of *Larus schistisagus*, *Cerorhinca*, and *Ardenna*, respectively.

**Figure 2.**
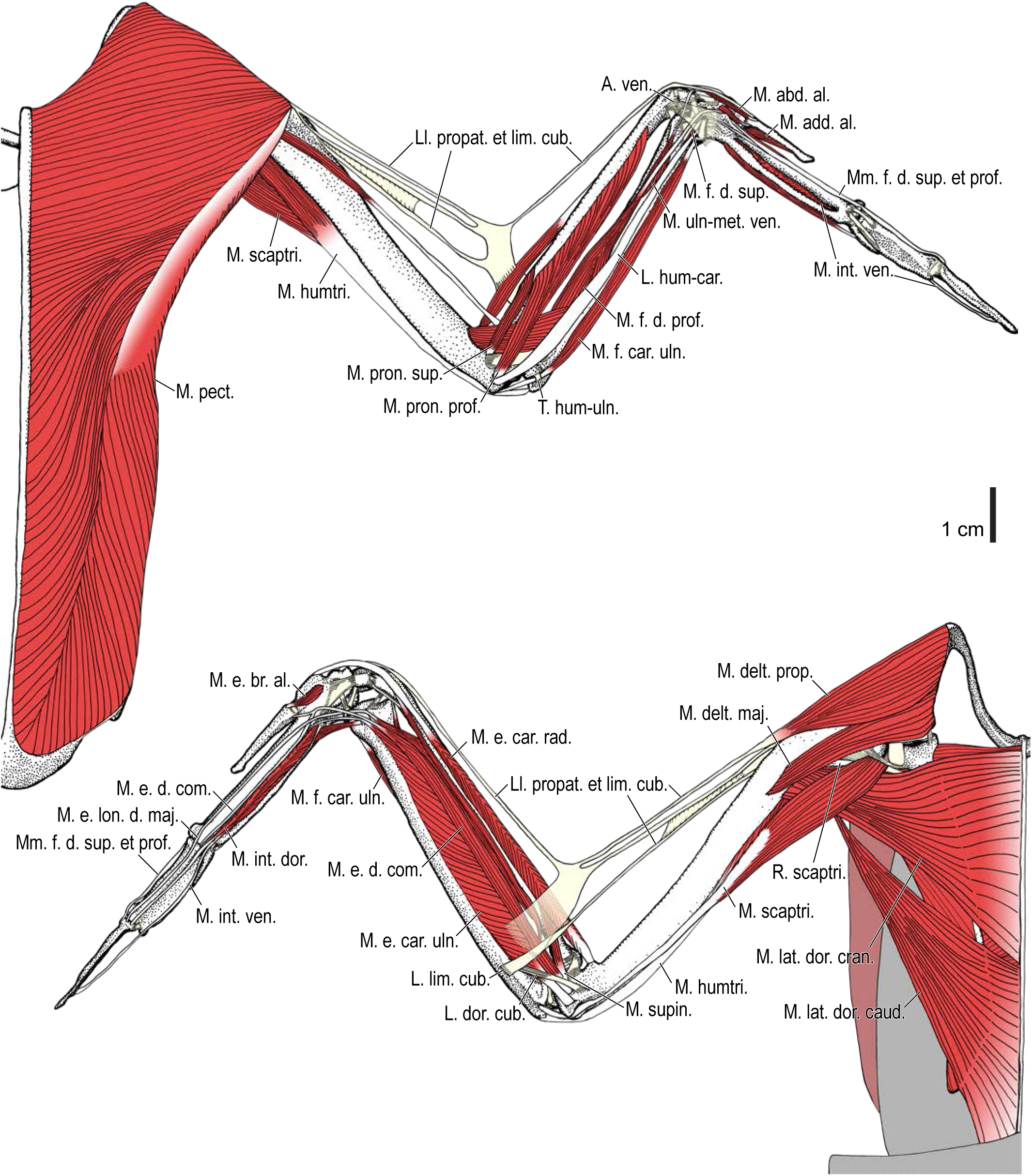
Wing musculature in extant *Alca torda*; ventral (top) and dorsal (bottom) views, superficial layer. This illustration is partly schematic, and is not an accurate representation of muscle volume, pennation, or other architectural properties. See Table 3 for abbreviations.

**Figure 3.**
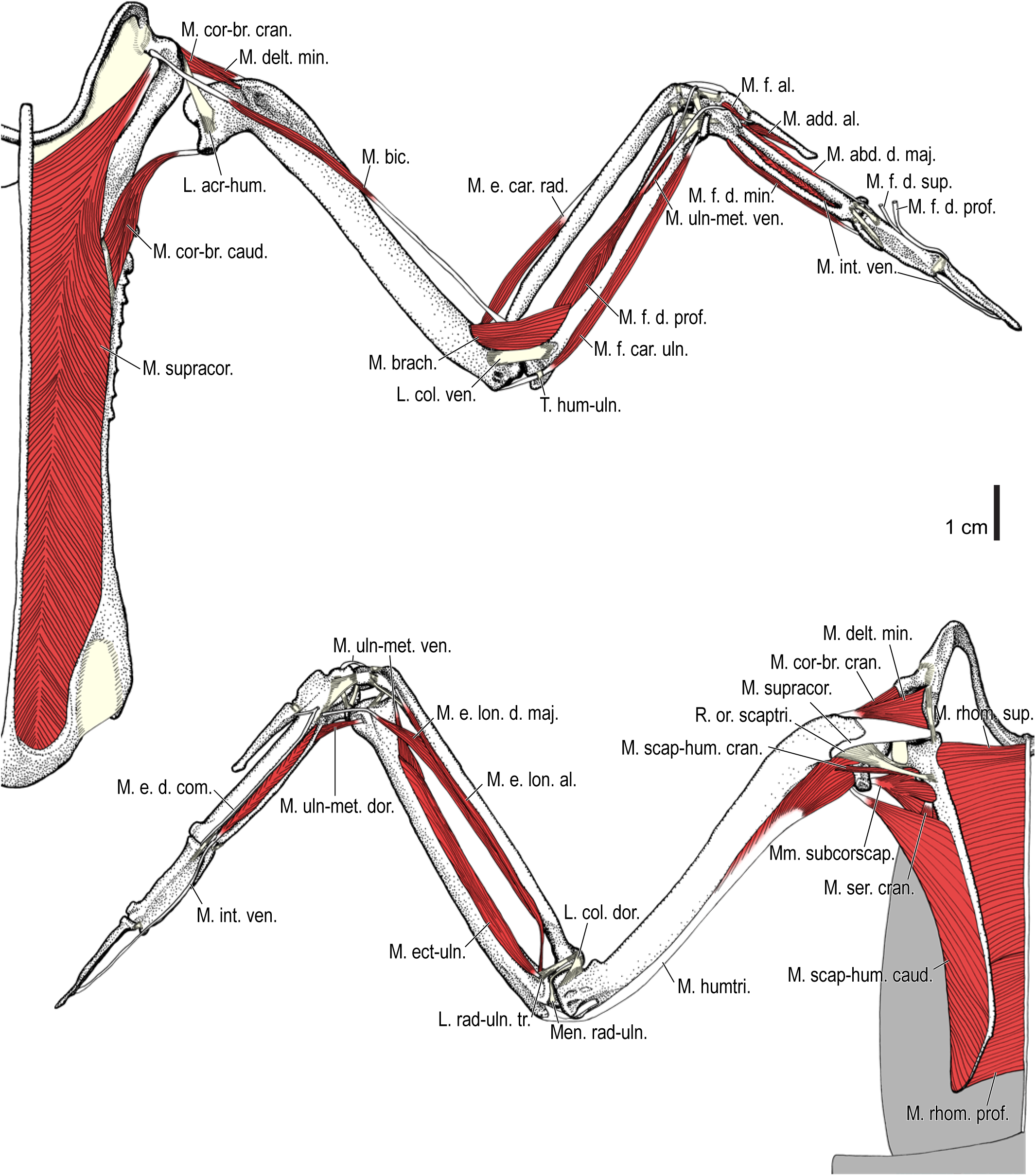
Wing musculature in extant *Alca torda*; ventral (top) and dorsal (bottom) views, deep layer. See Table 3 for abbreviations and Figure 2 for further information.

**Figure 4.**
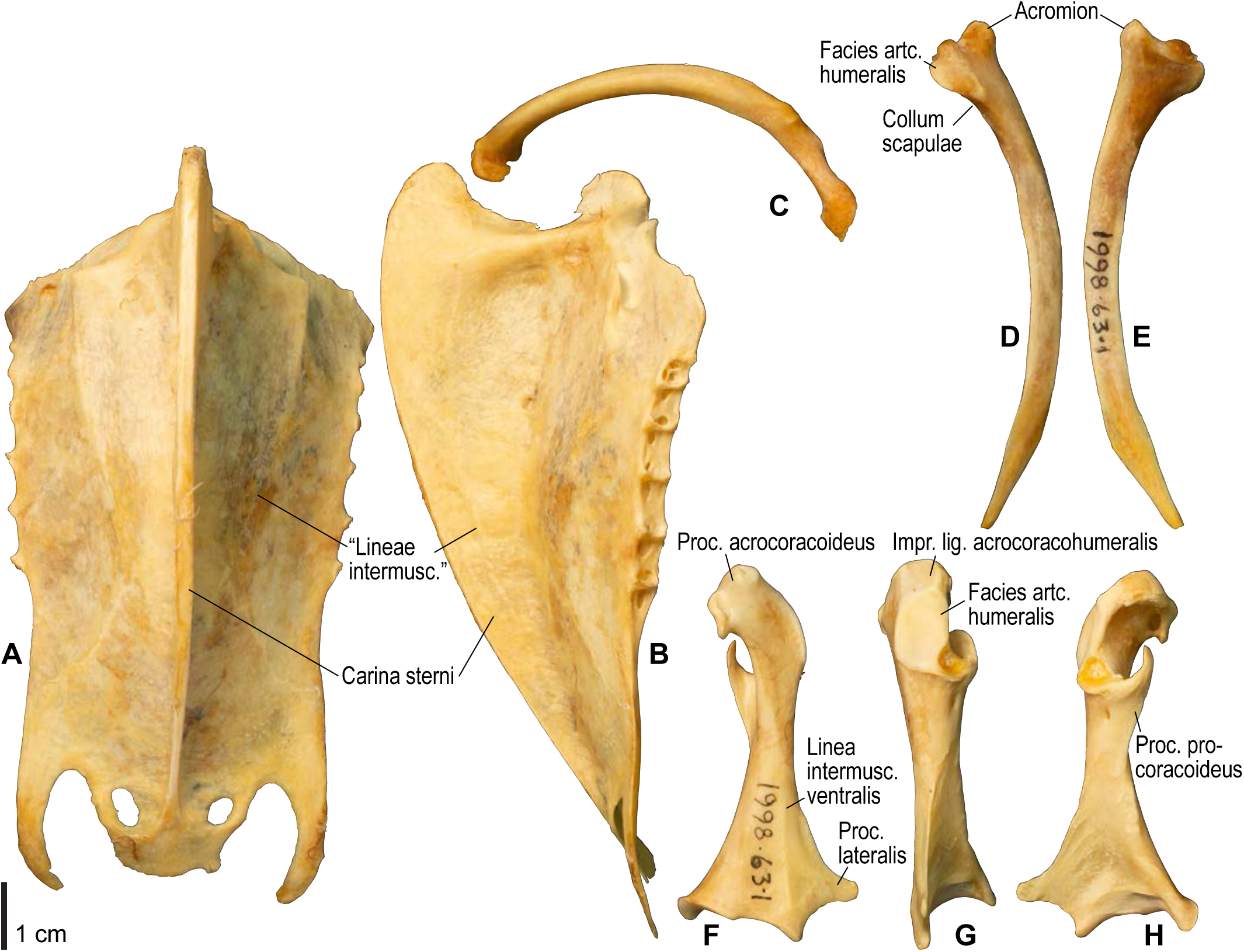
Osteology of *Catharacta antarctica*, pectoral girdle elements. Drawn on NHMUK 1998.63.1. Sternum in ventral (**A**) and left lateral (**B**) views; furcula in left lateral view (**C**); left scapula in lateral (**D**) and medial (**E**) views; left coracoid in ventral (**F**), lateral (**G**), and dorsal (**H**) views. Major osteological landmarks mentioned in text are designated. Abbreviations: artc., articularis; impr., impressio; intermusc., intermuscularis/intermusculares; lig., ligamenti; m., musculi; proc., processus; tuberc., tuberculum.

**Figure 5.**
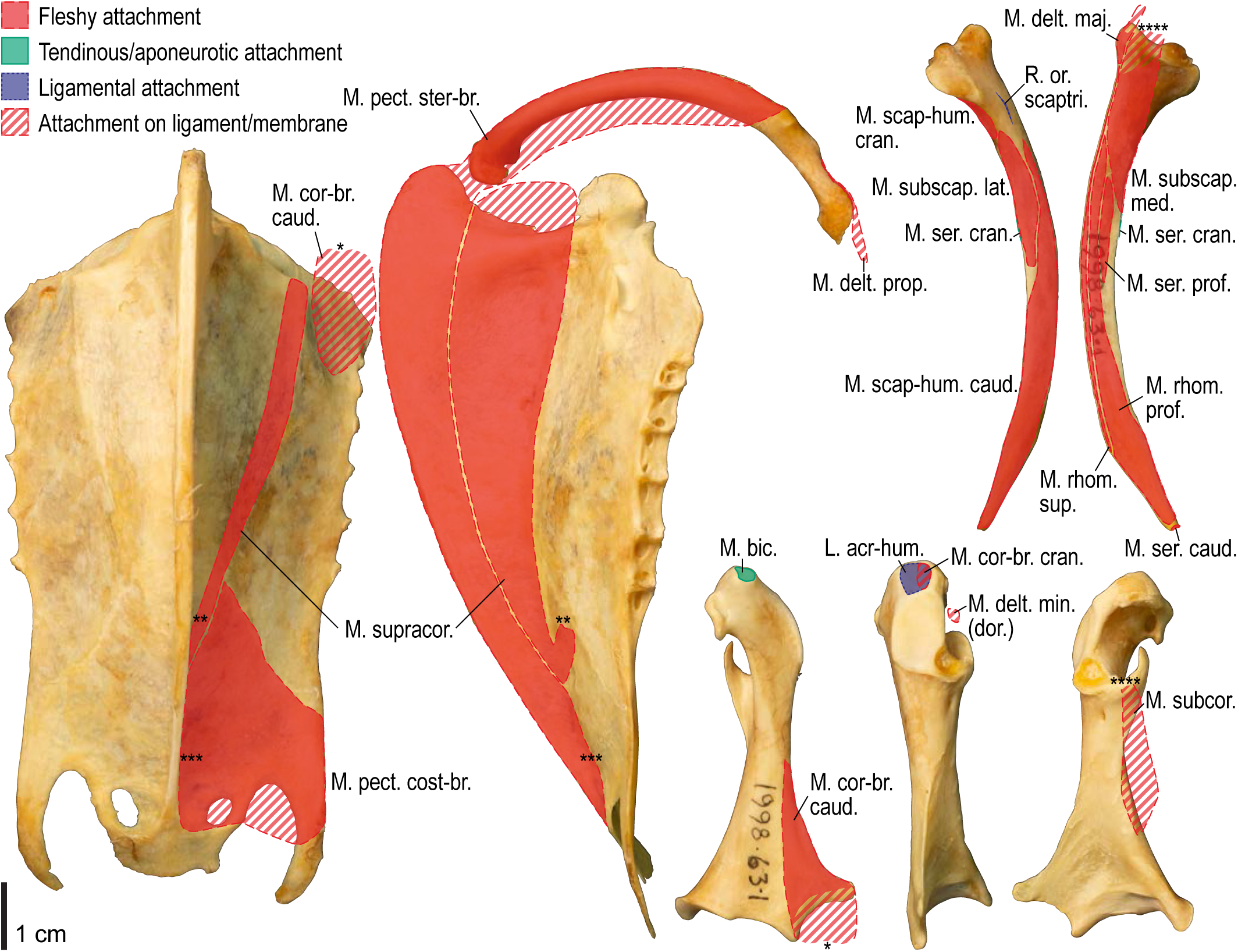
Osteological correlates of major wing muscles and ligaments in *Catharacta antarctica*, pectoral girdle elements. Drawn on NHMUK 1998.63.1. Note that only reliably identified attachment sites are shown, and the gaps between some adjacent attachment sites are exaggerated for distinction. Asterisks denote continuous attachment sites across panels. Red fill with broken outline, fleshy (direct) attachment of muscles; green fill with solid outline, tendinous/aponeurotic (indirect) attachment of muscles; blue fill with dotted outline, attachment of ligaments; stroked fill, attachment on ligaments/membranes.

**Figure 6.**
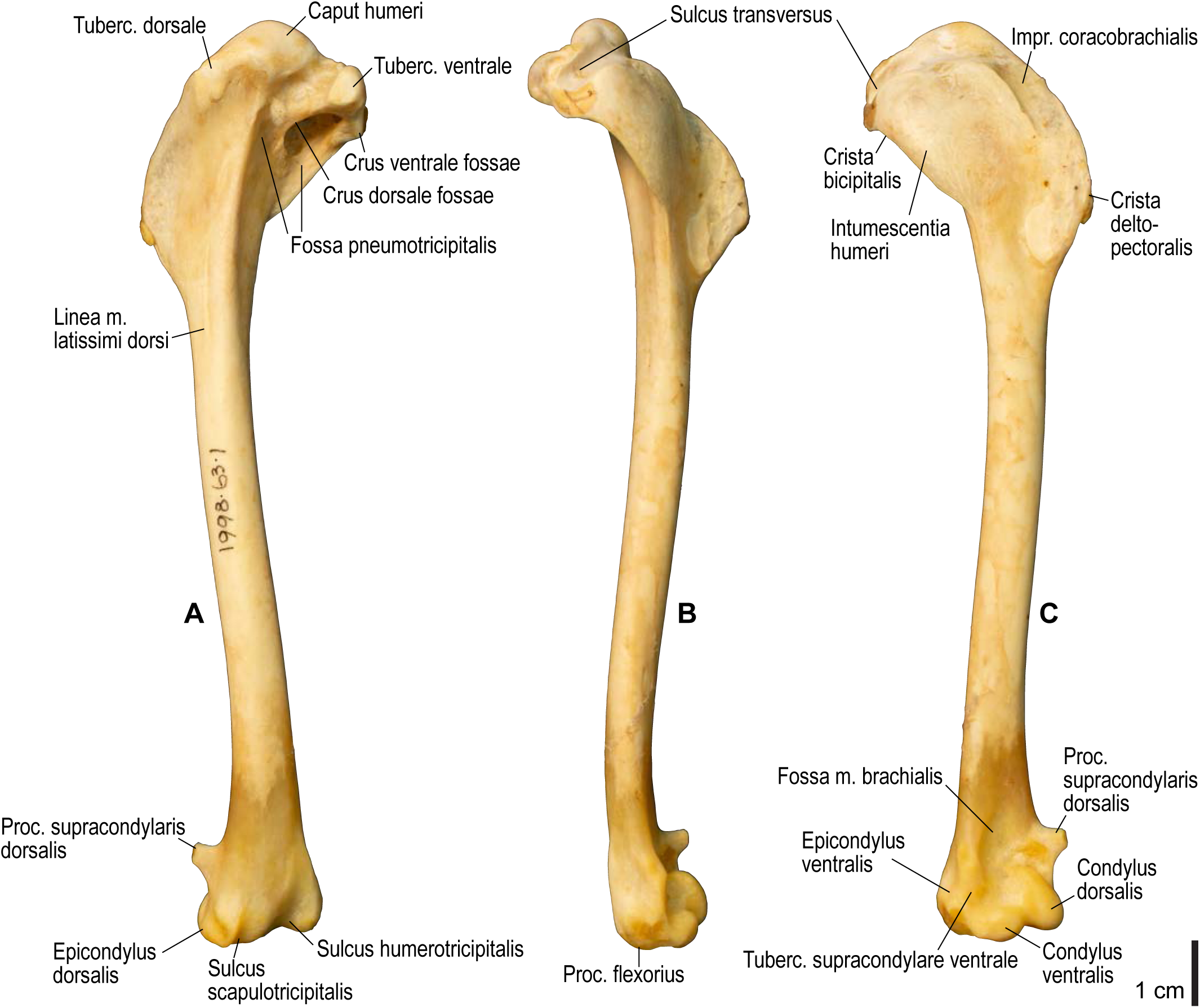
Osteology of *Catharacta antarctica*, humerus. Drawn on NHMUK 1998.63.1. Left humerus in caudal (**A**), ventral (**B**), and cranial (**C**) views. See Figure 4 for abbreviations.

**Figure 7.**
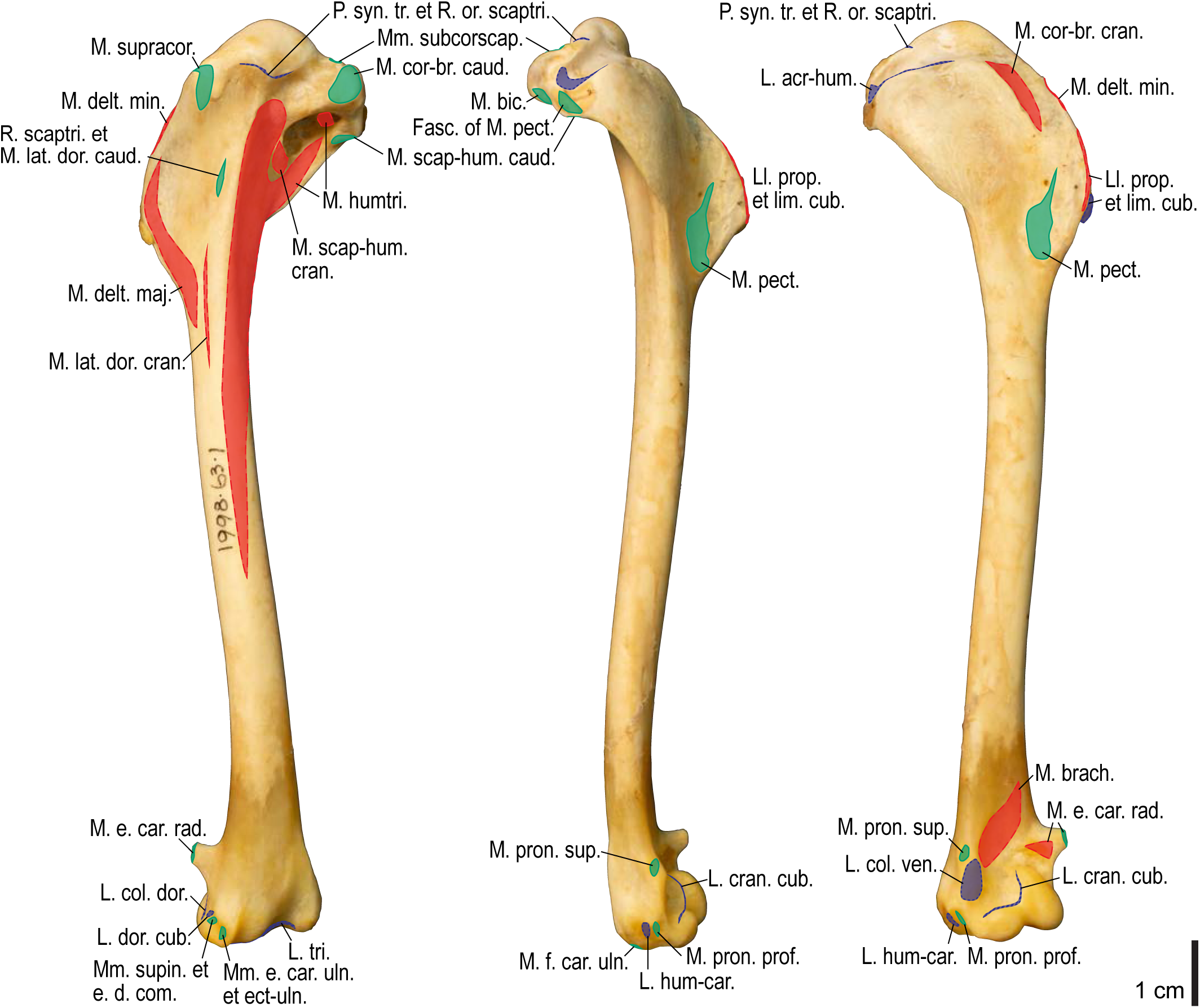
Osteological correlates of major wing muscles and ligaments in *Catharacta antarctica*, humerus. Drawn on NHMUK 1998.63.1. See Figure 5 for legends.

**Figure 8.**
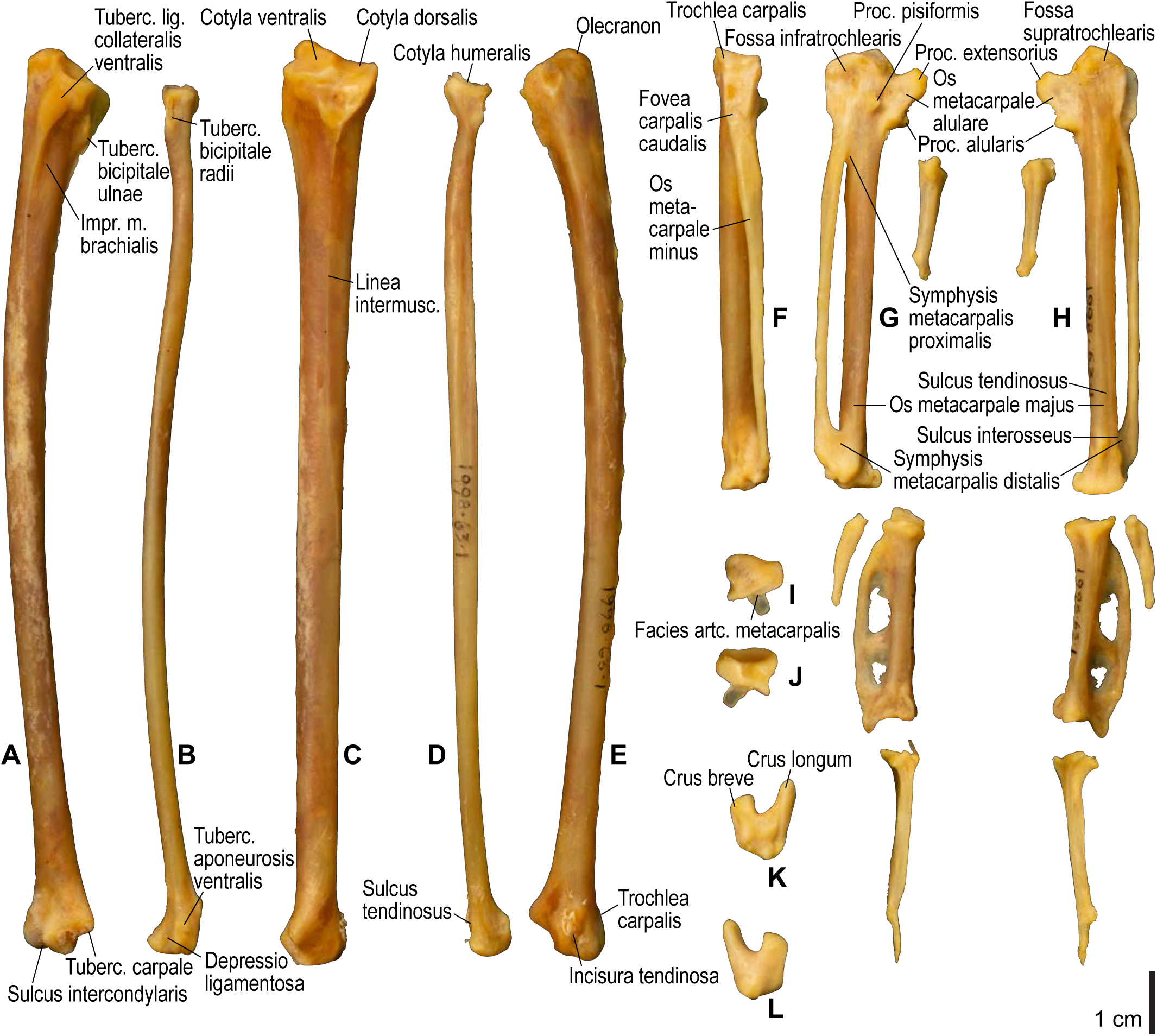
Osteology of *Catharacta antarctica*, distal wing elements. Drawn on NHMUK 1998.63.1. Left ulna in ventral (**A**), cranial (**C**), and dorsal (**E**) views; left radius in ventral (**B**) and dorsocaudal (**D**) views; left carpometacarpus and phalanges in caudal (**F**, phalanges not shown), ventral (**G**), and dorsal views; left radiale in cranial (**I**) and caudal (**J**) views; left ulnare in proximal (**K**) and distal (**L**) views. See Figure 4 for abbreviations.

**Figure 9.**
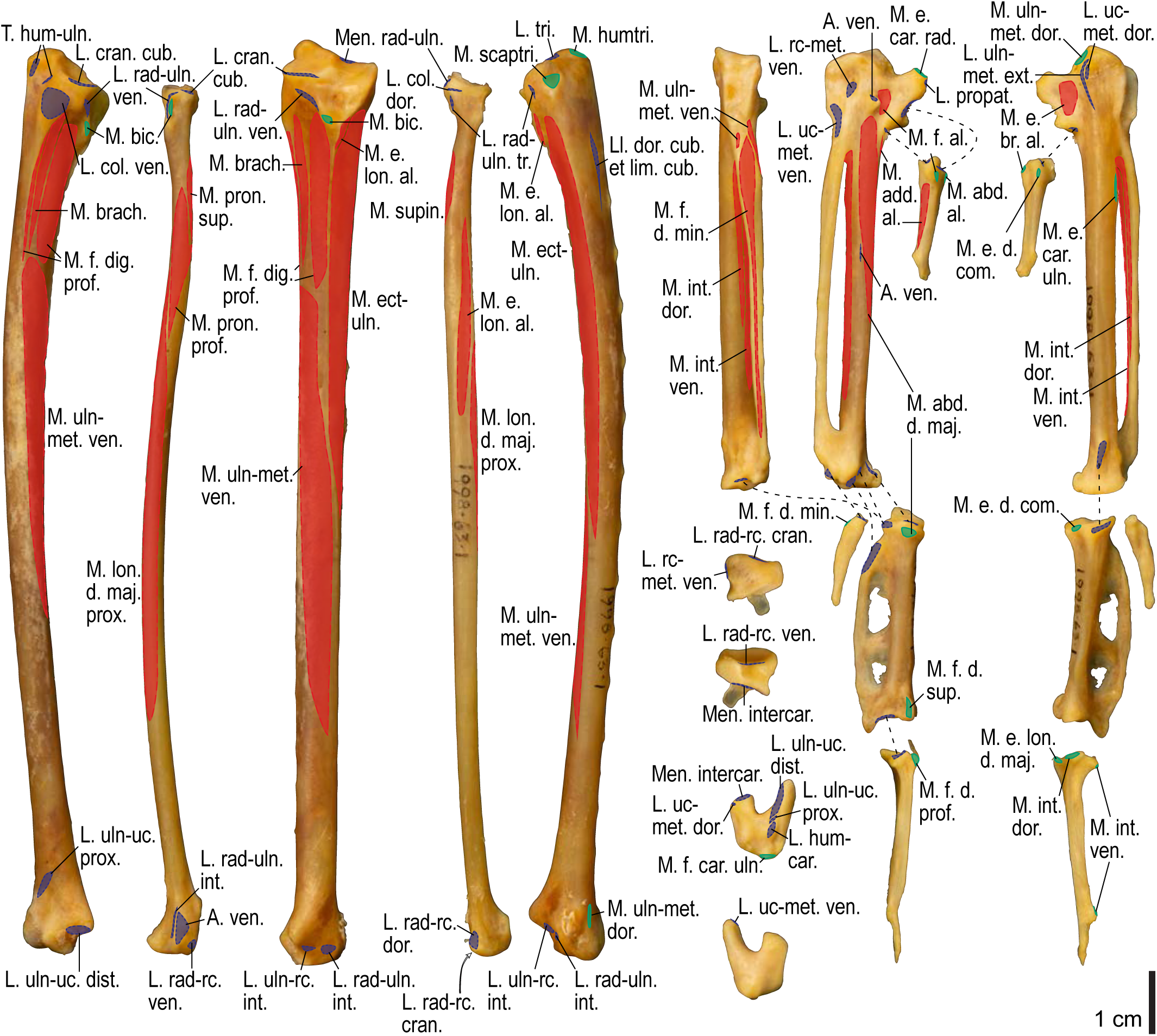
Osteological correlates of major wing muscles and ligaments in *Catharacta antarctica*, distal wing elements. Drawn on NHMUK 1998.63.1. Due to space restrictions, the labels for some distal wing ligaments are not shown; broken lines show correspondence of attachment sites for these ligaments. See Figure 5 for legends.

**Figure 10.**
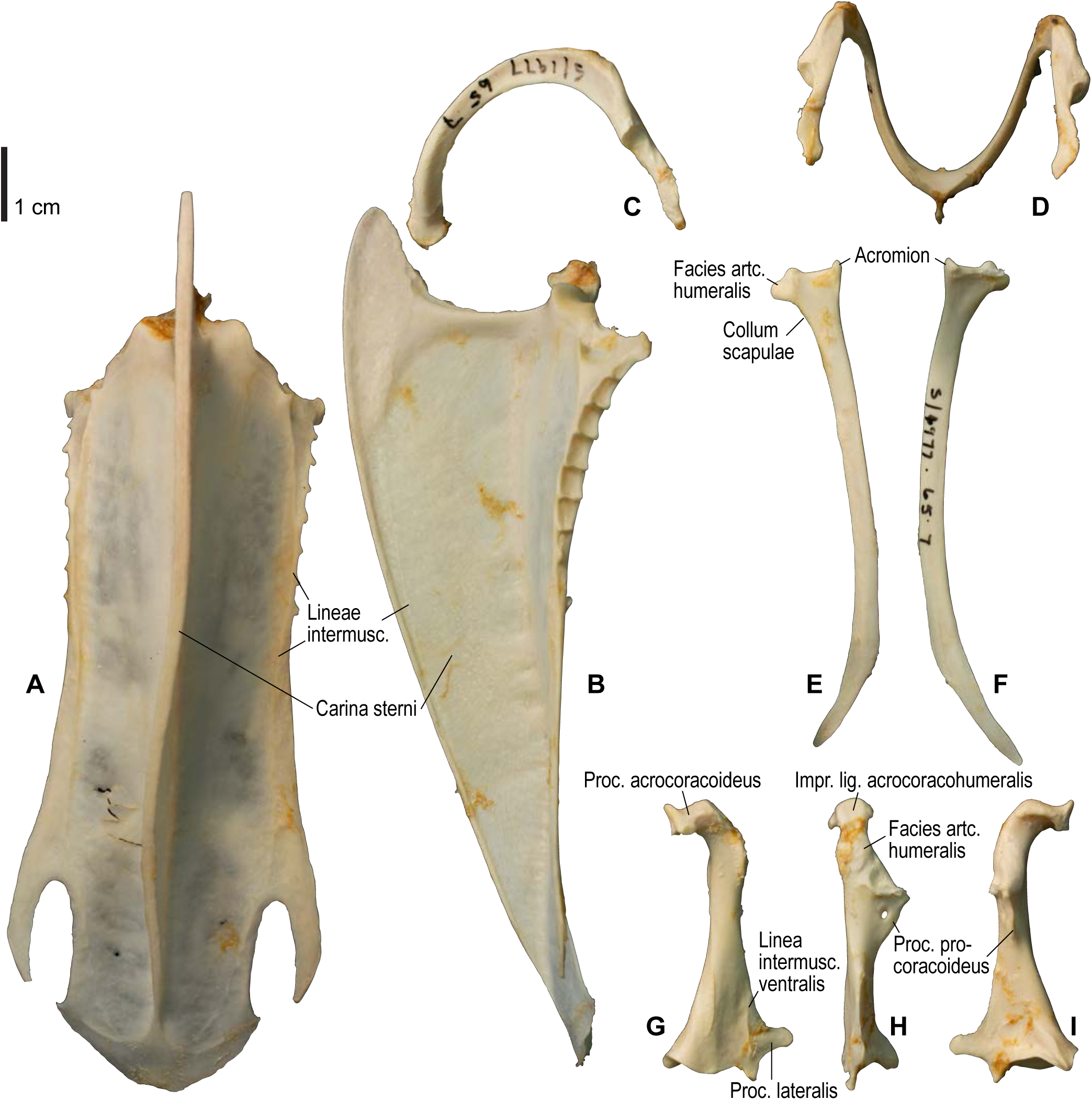
Osteology of *Alca torda*, pectoral girdle elements. Drawn on NHMUK S/1977.65.7. Sternum in ventral (**A**) and left lateral (**B**) views; furcula in left lateral (**C**) and dorsal (**D**) views; left scapula in lateral (**E**) and medial (**F**) views; left coracoid in ventral (**G**), lateral (**H**), and dorsal (**I**) views. See Figure 4 for abbreviations.

**Figure 11.**
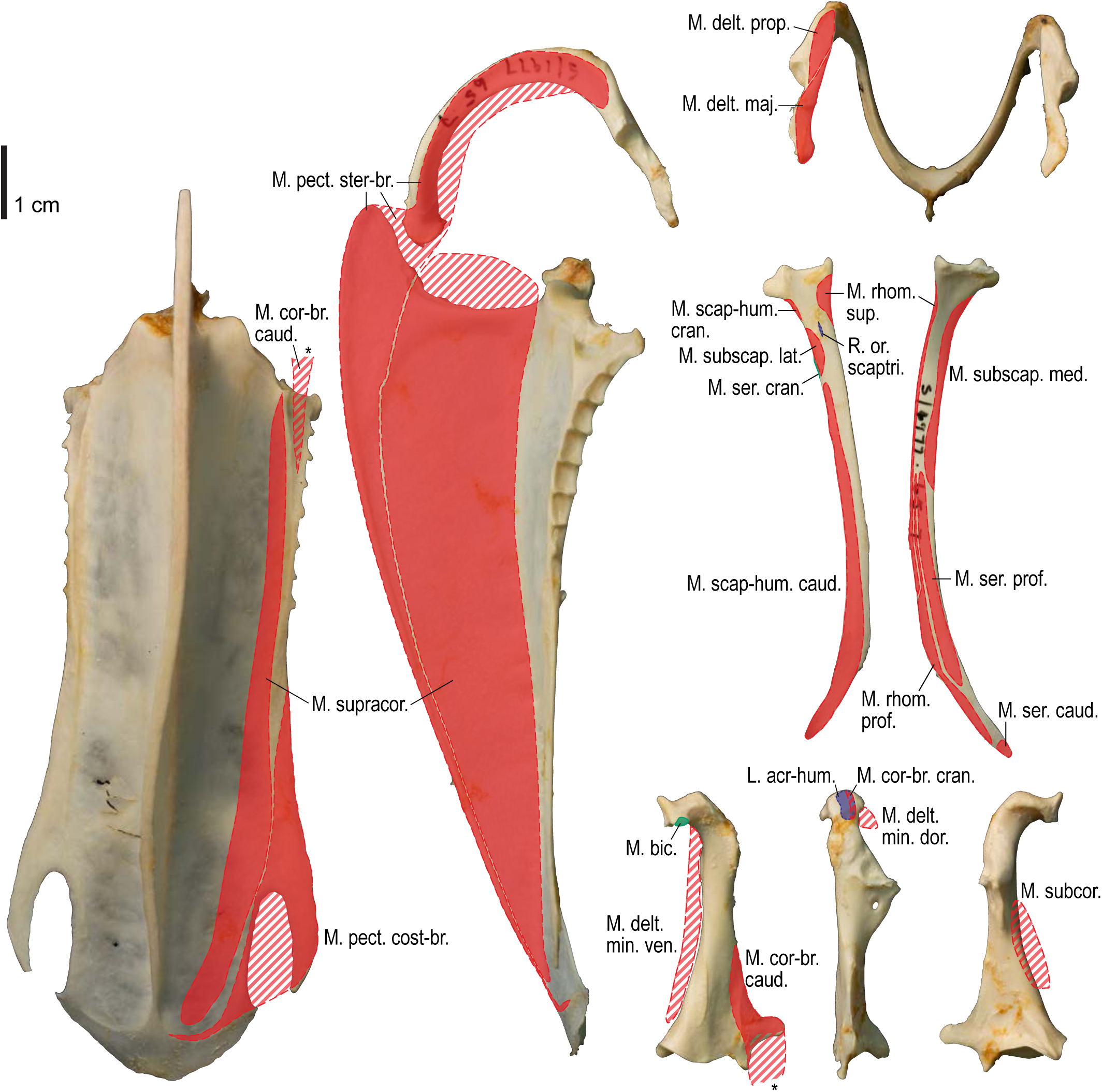
Osteological correlates of major wing muscles and ligaments in *Alca torda*, pectoral girdle elements. Drawn on NHMUK S/1977.65.7. See Figure 5 for legends.

**Figure 12.**
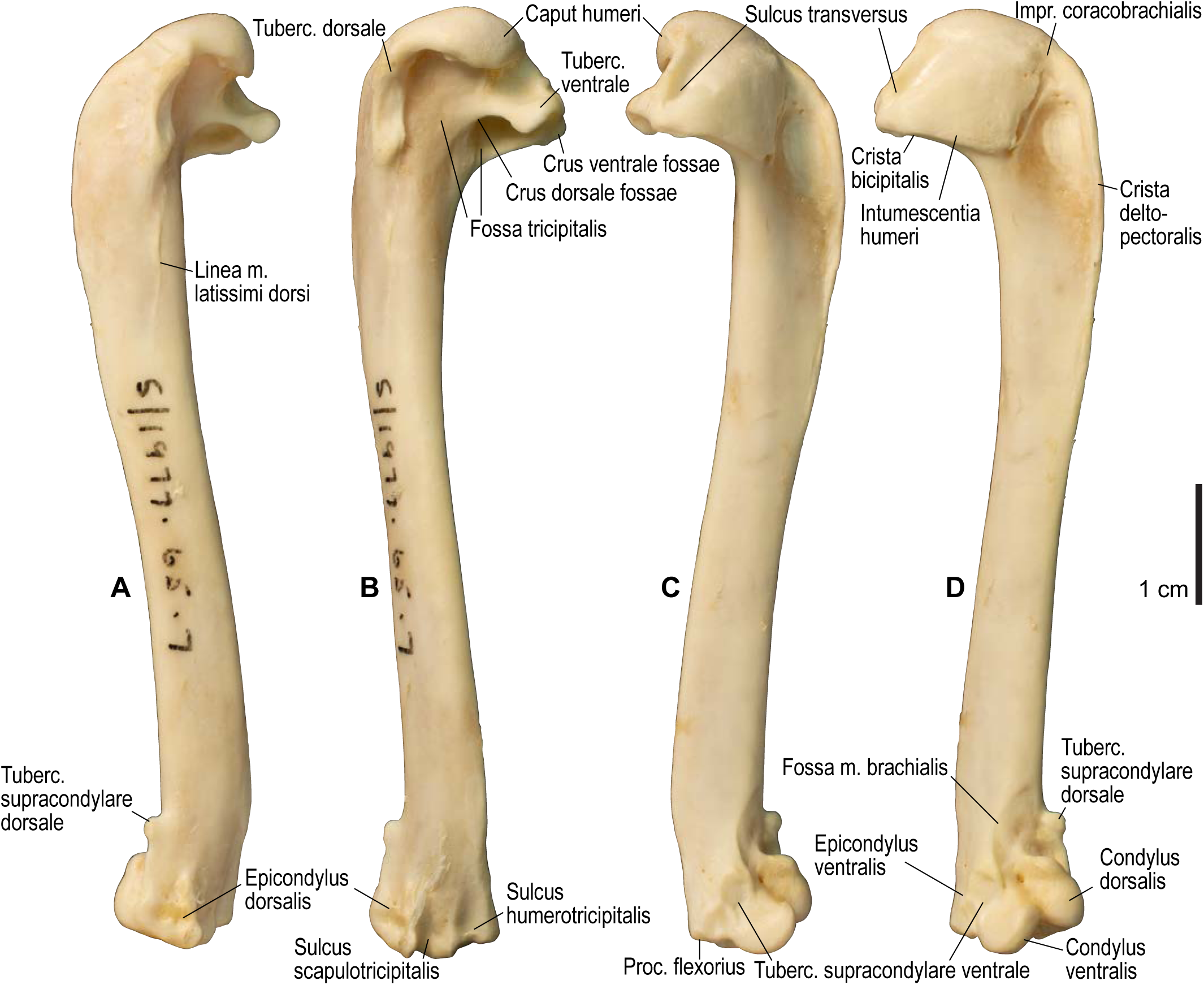
Osteology of *Alca torda*, humerus. Drawn on NHMUK S/1977.65.7. Left humerus in dorsal (**A**), caudal (**B**), ventral (**C**), and cranial (**D**) views. See Figure 4 for abbreviations.

**Figure 13.**
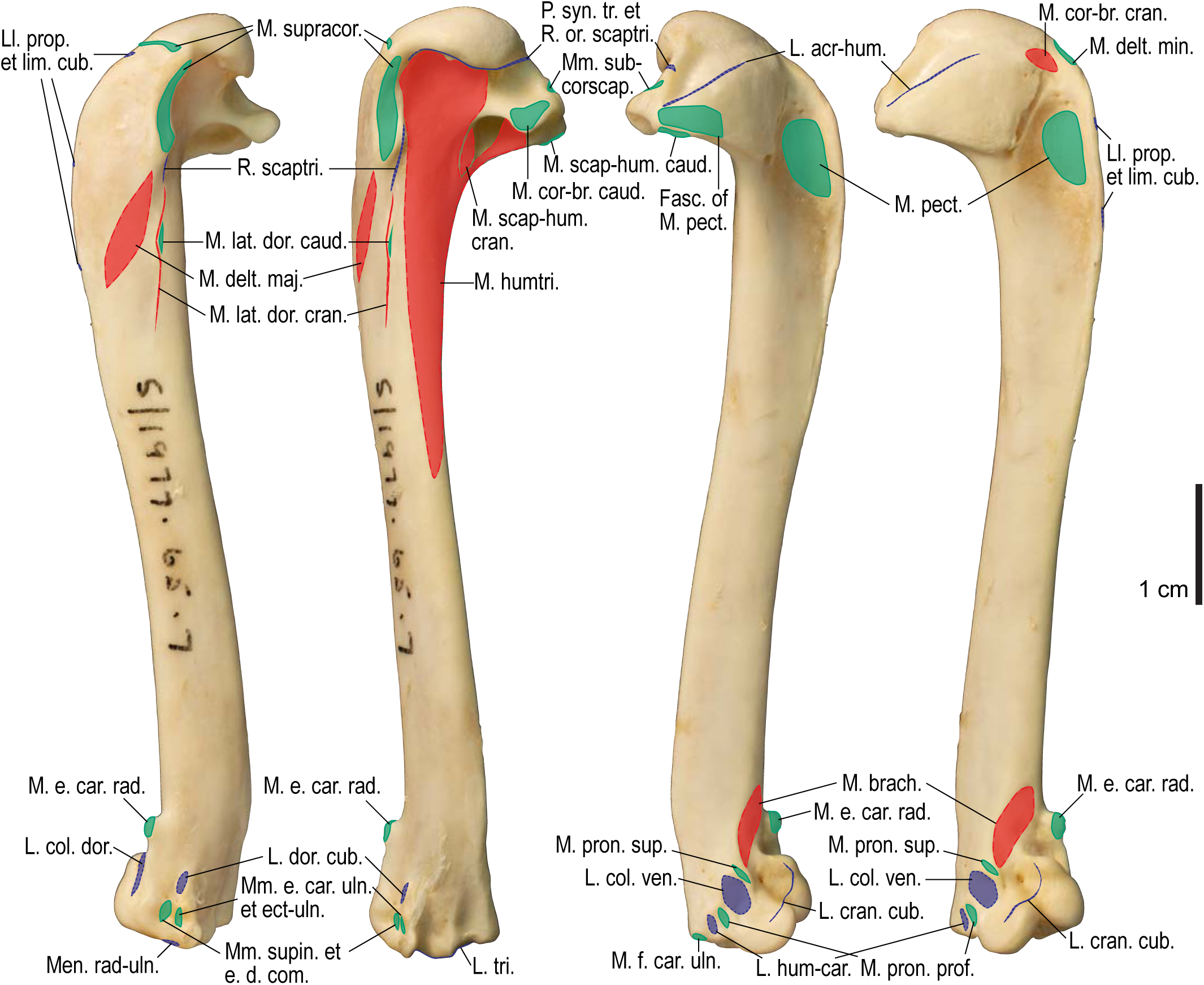
Osteological correlates of major wing muscles and ligaments in *Alca torda*, humerus. Drawn on NHMUK S/1977.65.7. See Figure 5 for legends.

**Figure 14.**
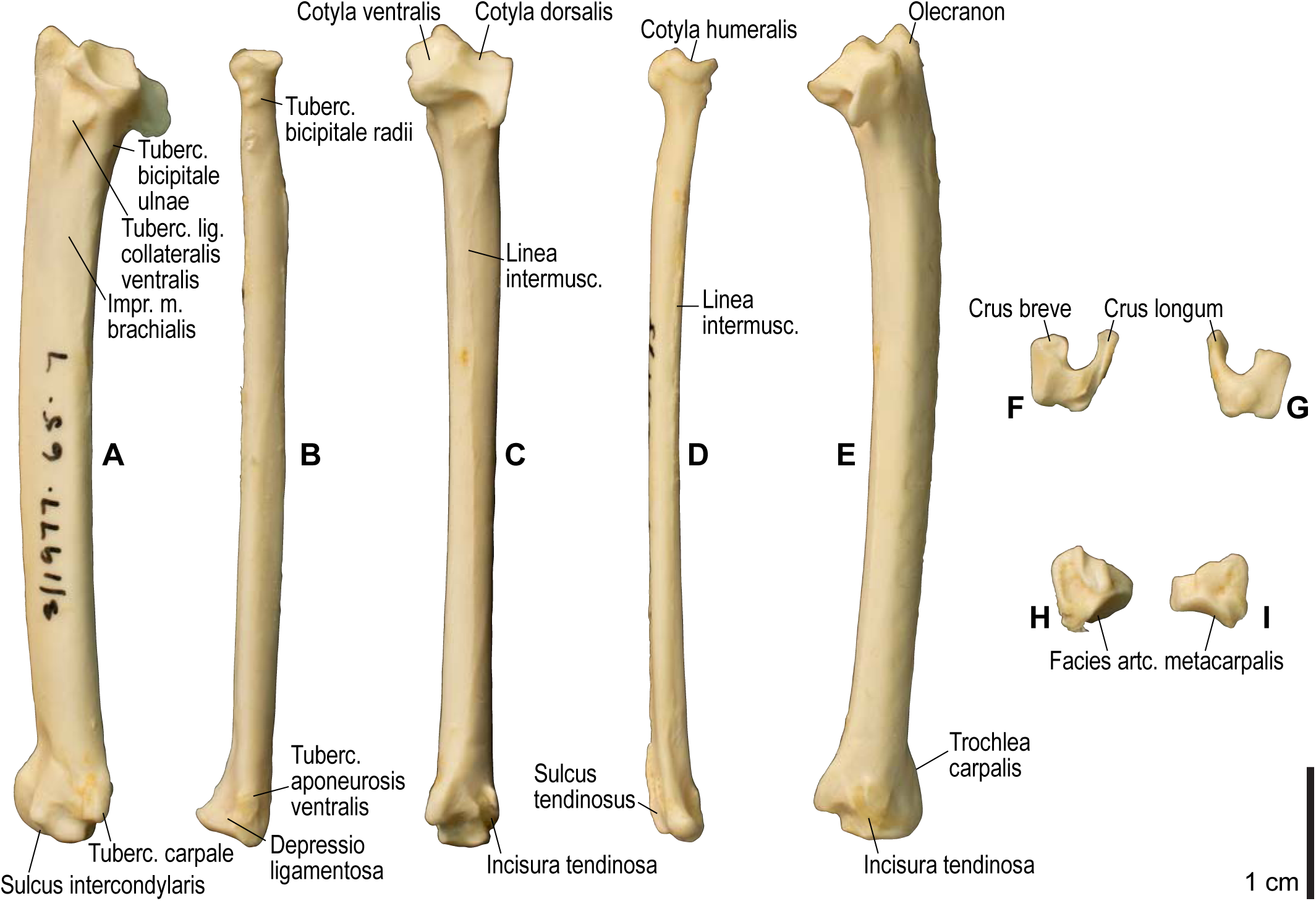
Osteology of *Alca torda*, forearm and free carpal elements. Drawn on NHMUK S/1977.65.7. Left ulna in ventral (**A**), cranial (**C**), and dorsal (**E**) views; left radius in ventral (**B**) and dorsocaudal (**D**) views; left ulnare in proximal (**F**) and distal (**G**) views; left radiale in cranial (**H**) and caudal (**I**) views. See Figure 4 for abbreviations.

**Figure 15.**
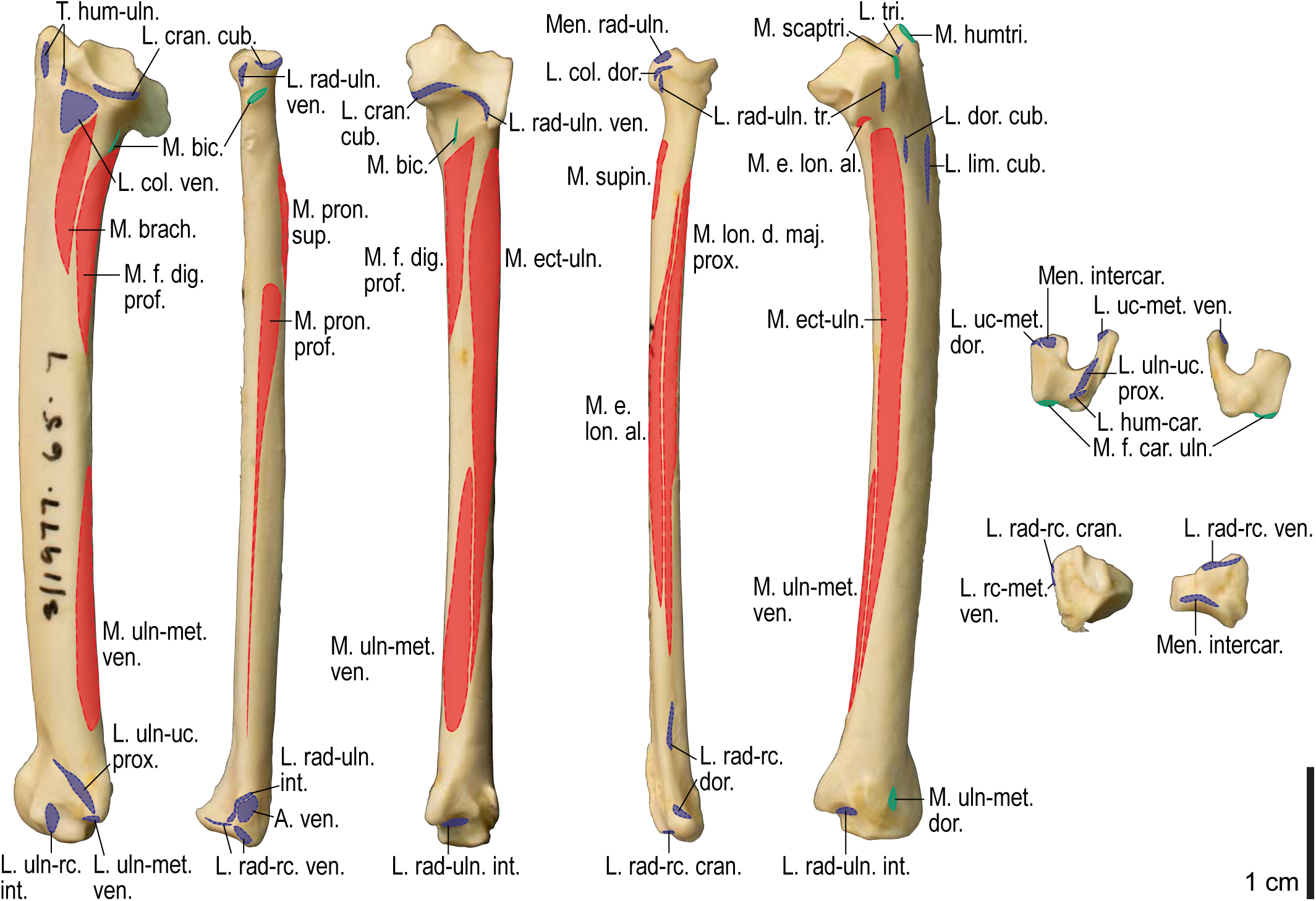
Osteological correlates of major wing muscles and ligaments in *Alca torda*, forearm and free carpal elements. Drawn on NHMUK S/1977.65.7. See Figure 5 for legends.

**Figure 16.**
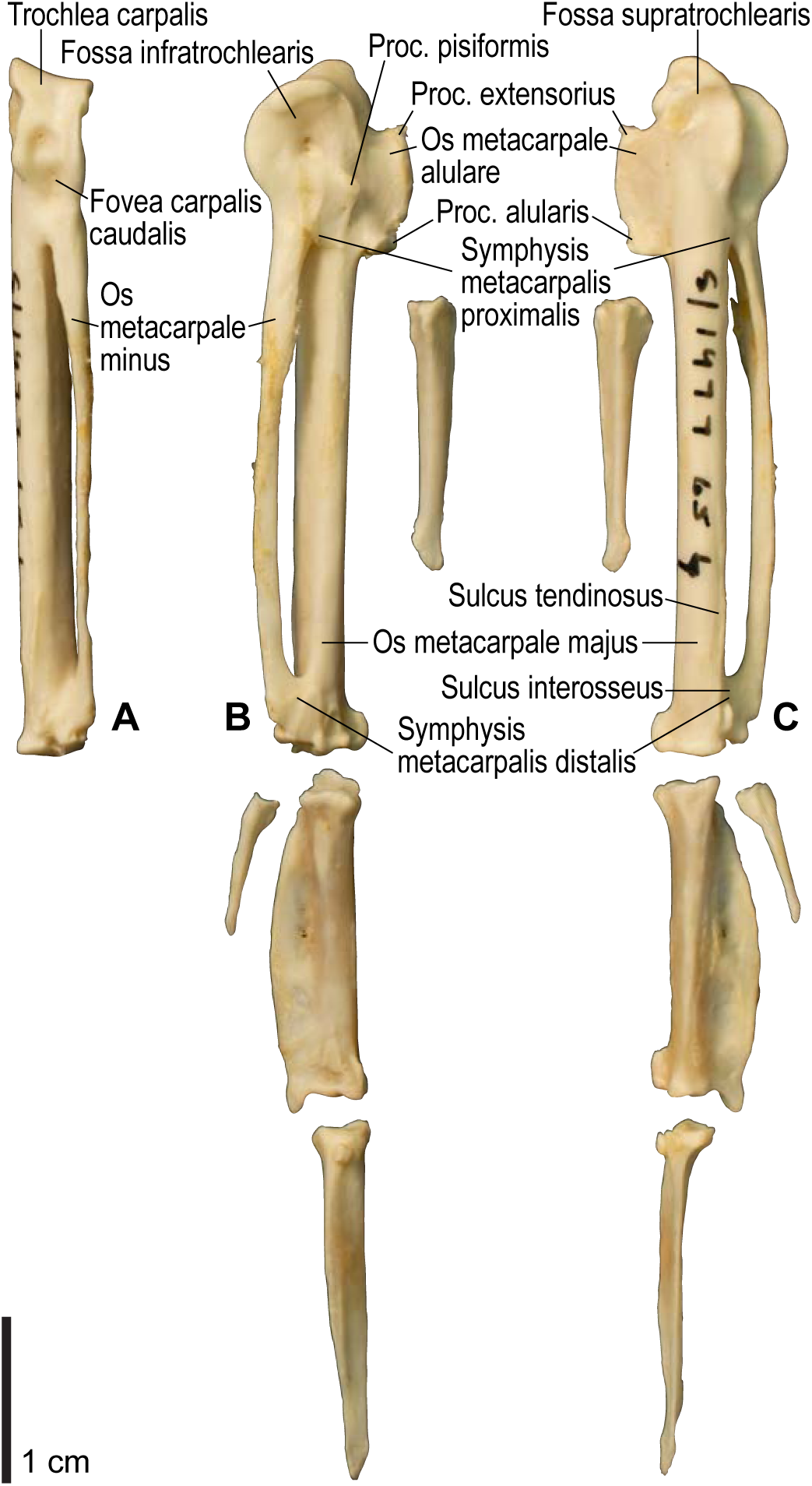
Osteology of *Alca torda*, manual elements. Drawn on NHMUK S/1977.65.7. Left carpometacarpus and phalanges in caudal (**A**, phalanges not shown), ventral (**B**), and dorsal (**C**) views. See Figure 4 for abbreviations.

**Figure 17.**
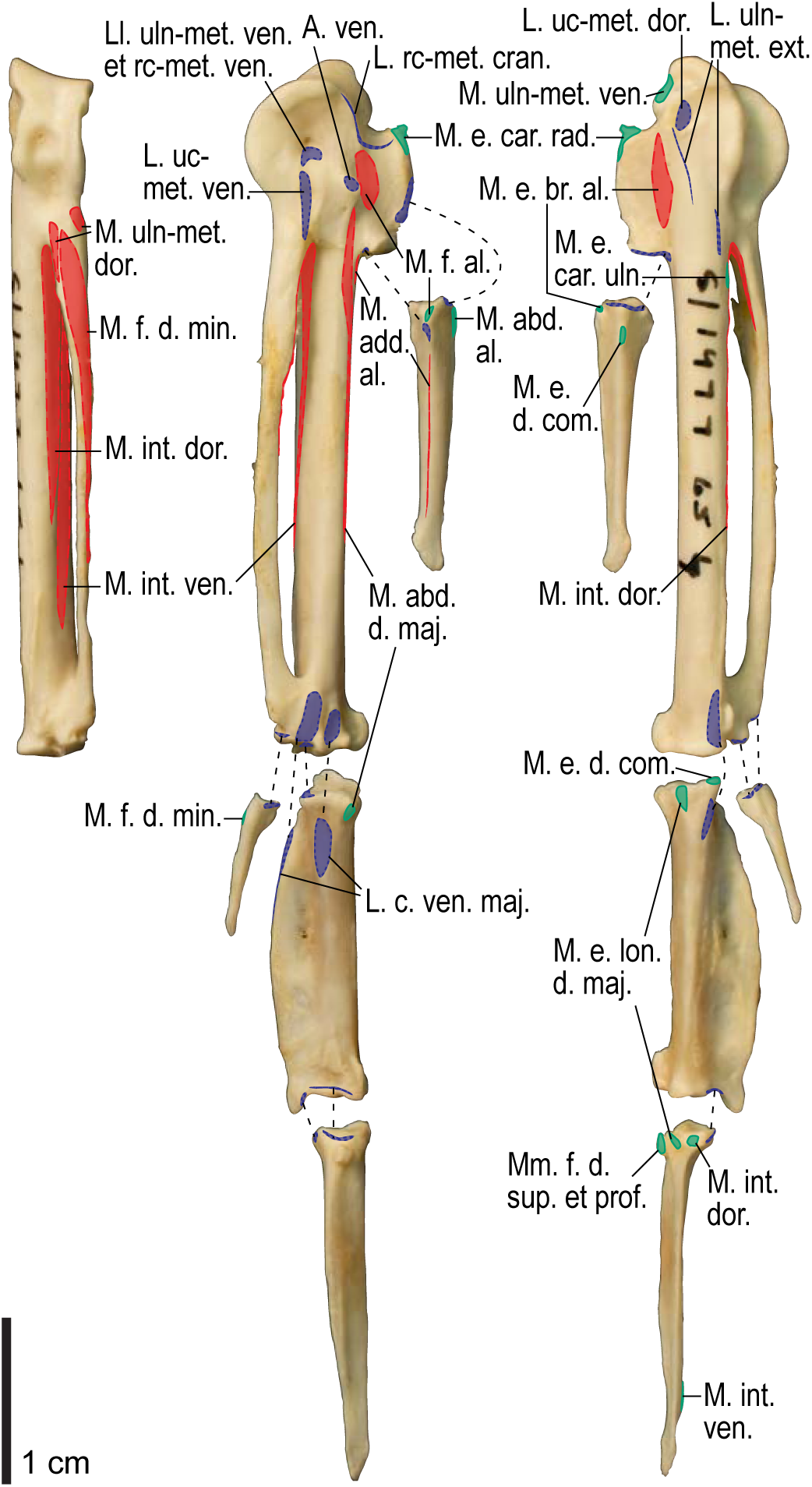
Osteological correlates of major wing muscles and ligaments in *Alca torda*, manual elements. Drawn on NHMUK S/1977.65.7. See Figures 5 and 9 for legends.

**Figure 18.**
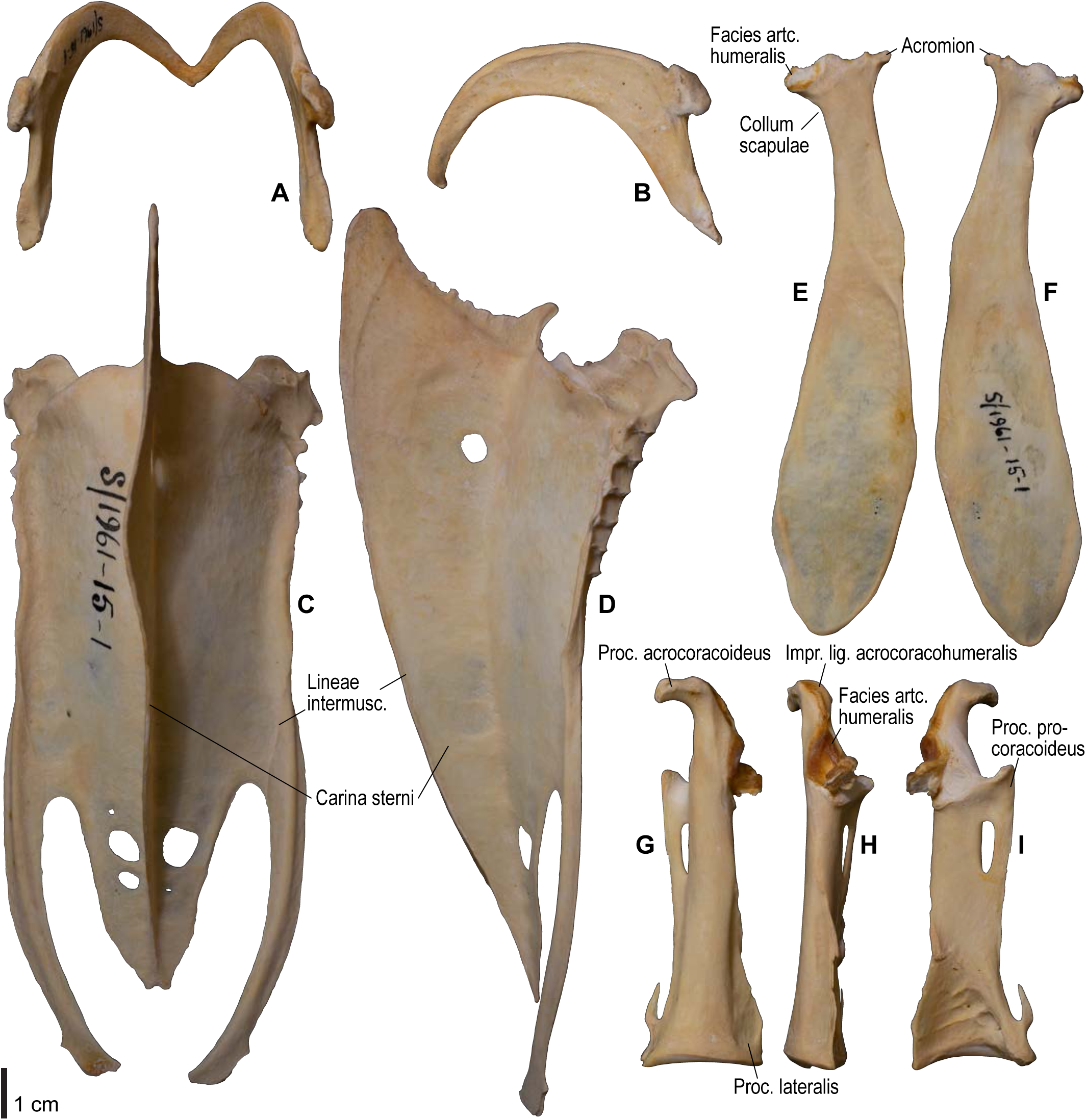
Osteology of *Spheniscus humboldti*, pectoral girdle elements. Drawn on NHMUK S/1961.15.1. Furcula in dorsal (**A**) and left lateral (**B**) views; sternum in ventral (**C**) and left lateral (**D**) views; left scapula in lateral (**E**) and medial (**F**) views; left coracoid in ventral (**G**), lateral (**H**), and dorsal views. See Figure 4 for abbreviations.

**Figure 19.**
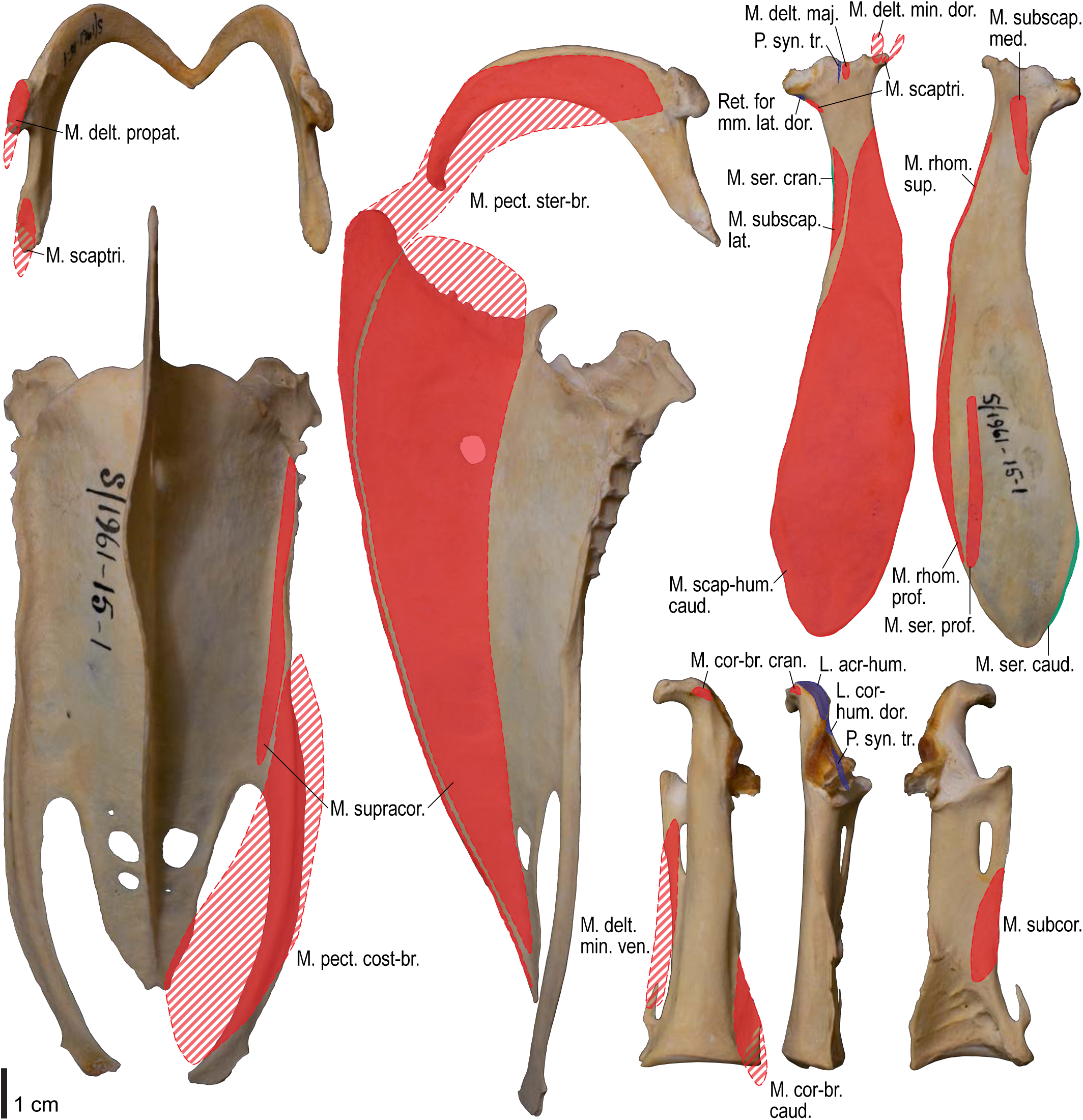
Osteological correlates of major wing muscles and ligaments in *Spheniscus humboldti*, pectoral girdle elements. Drawn on NHMUK S/1961.15.1. See Figure 5 for legends.

**Figure 20.**
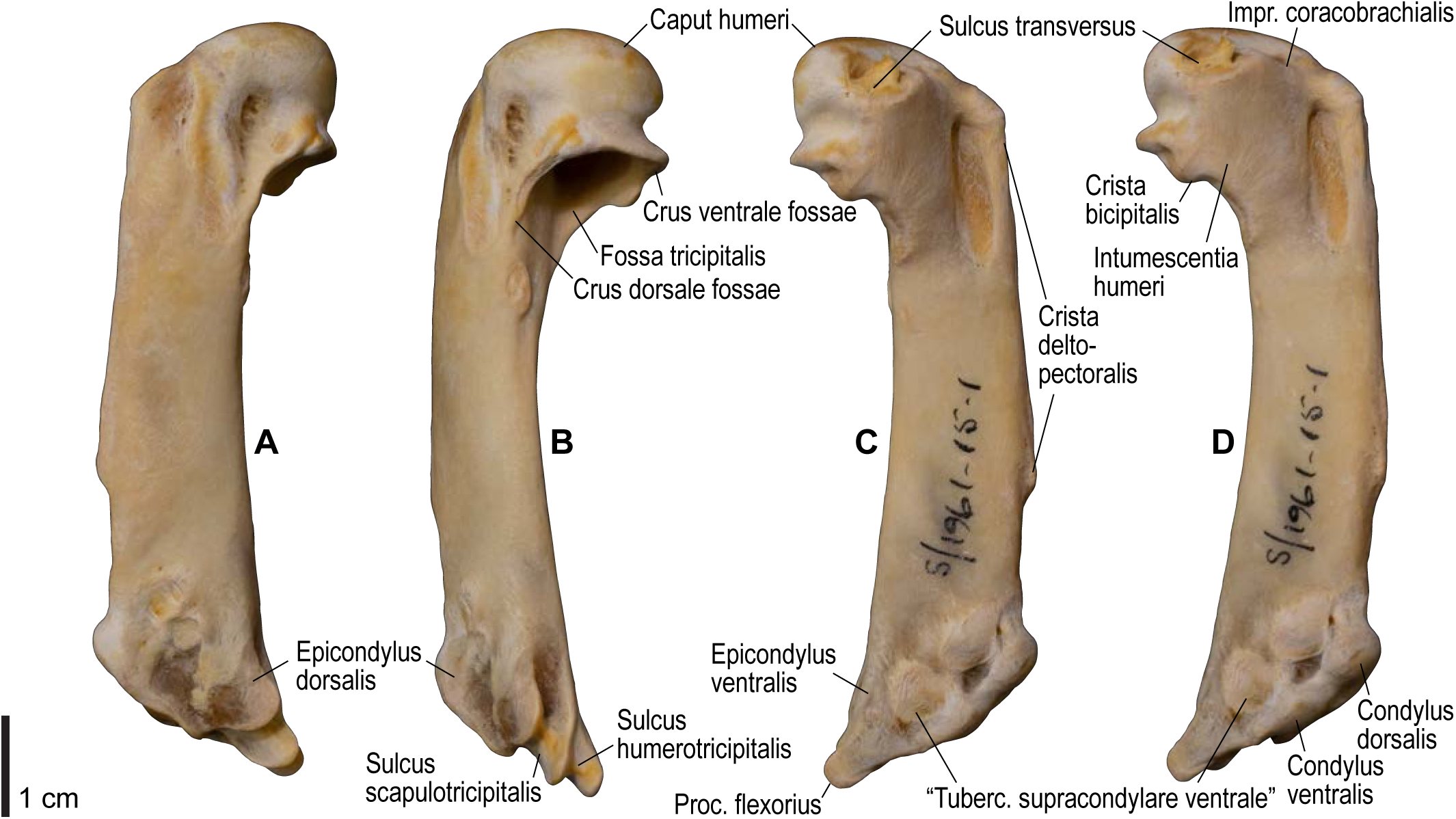
Osteology of *Spheniscus humboldti*, humerus. Drawn on NHMUK S/1977.65.7. Left humerus in dorsal (**A**), caudal (**B**), ventral (**C**), and cranial (**D**) views. See Figure 4 for abbreviations.

**Figure 21.**
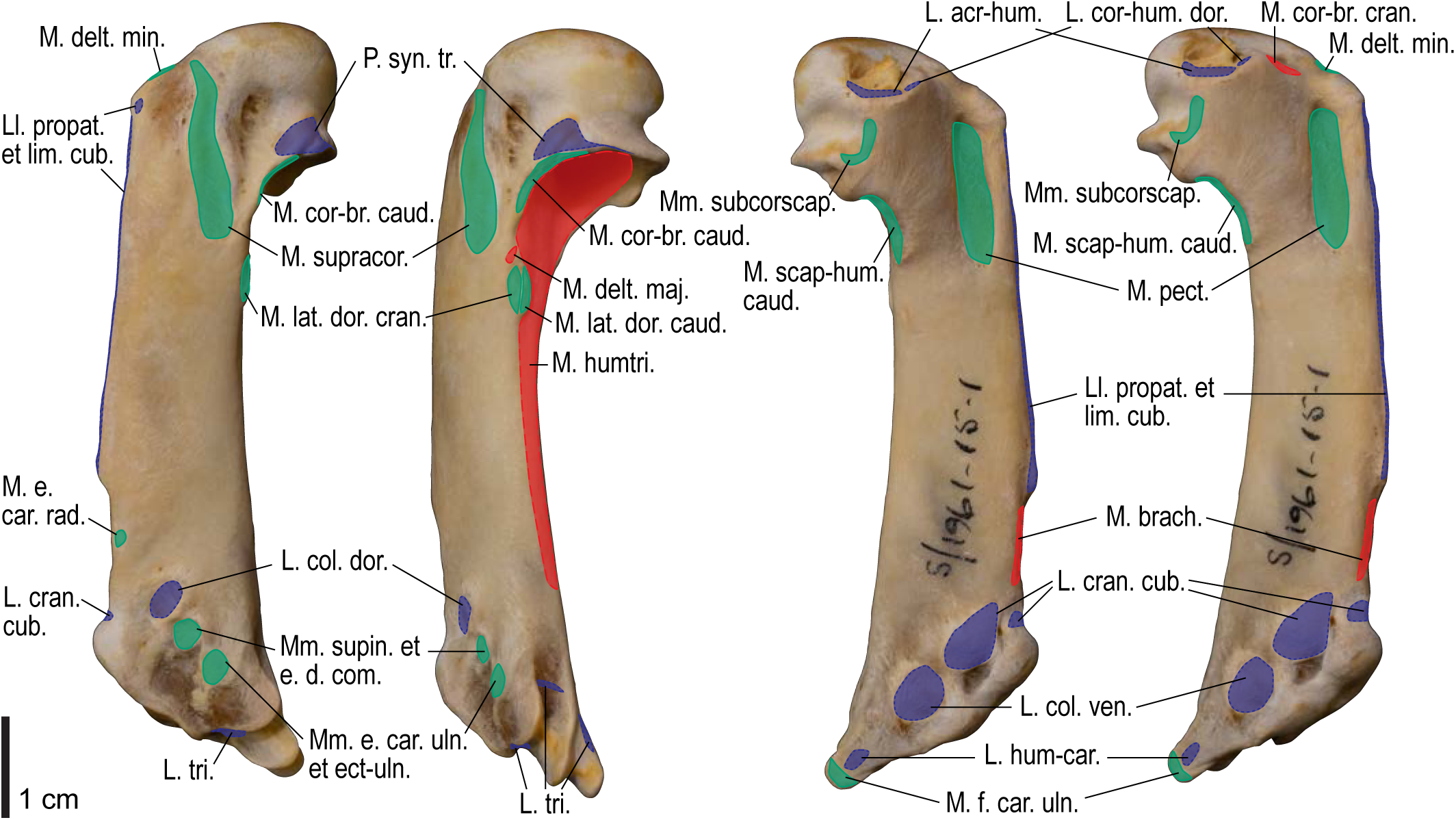
Osteological correlates of major wing muscles and ligaments in *Spheniscus humboldti*, humerus. Drawn on NHMUK S/1961.15.1. See Figure 5 for legends.

**Figure 22.**
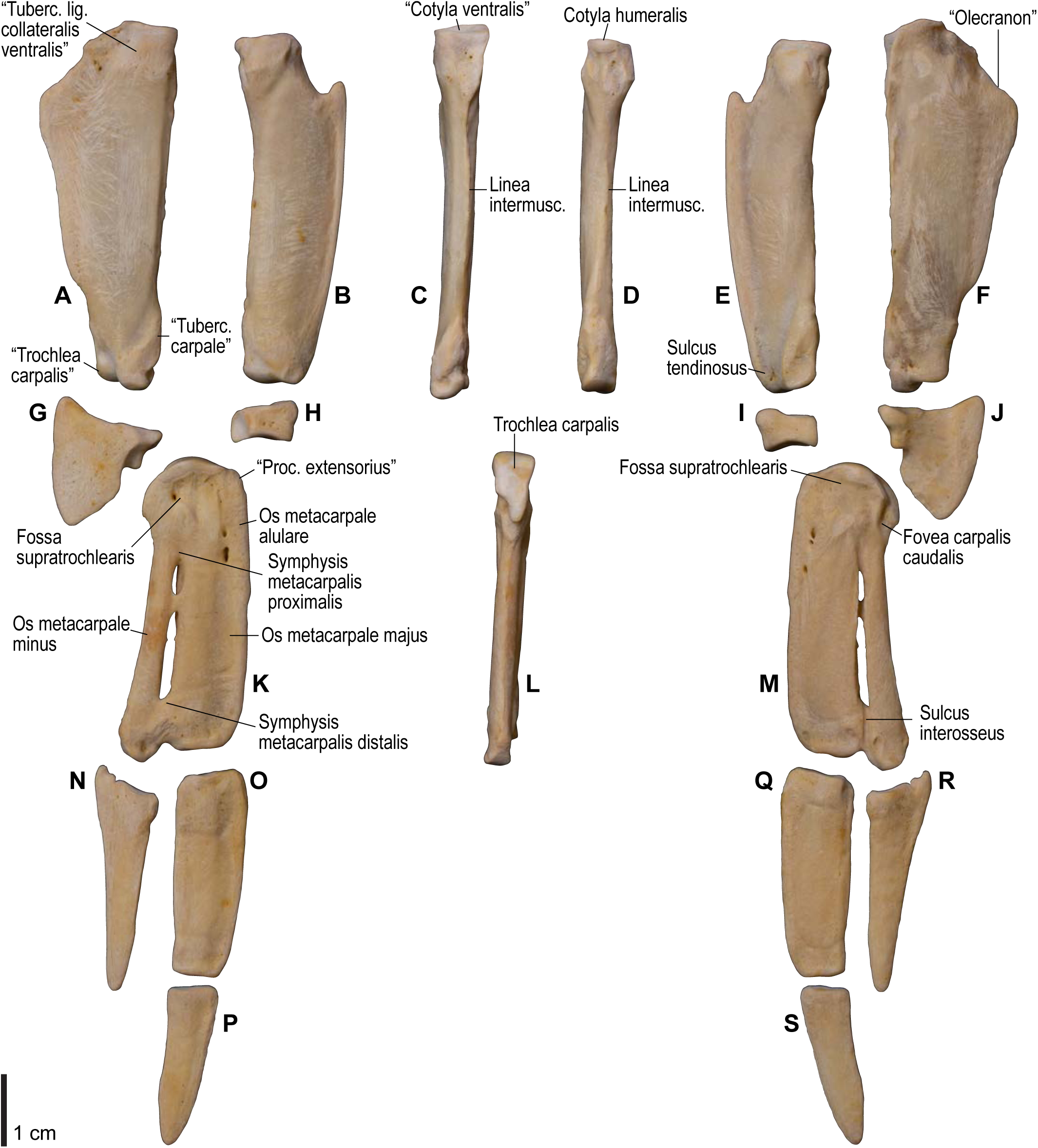
Osteology of *Spheniscus humboldti*, distal wing elements. Drawn on NHMUK S/1961.15.1. Left ulna (**A**, **C**, **F**), radius (**B**, **D**, **E**), ulnare (**G**, **J**), radiale (**H**, **I**), carpometacarpus (**K**–**M**), and phalanges (**N**–**S**) in ventral (**A**, **B**, **G**, **H**, **K**, **N**–**P**), cranial (**C**), caudal (**D**, **L**), and dorsal (**E**, **F**, **I**, **J**, **M**, **Q**–**S**) views. See Figure 4 for abbreviations.

**Figure 23.**
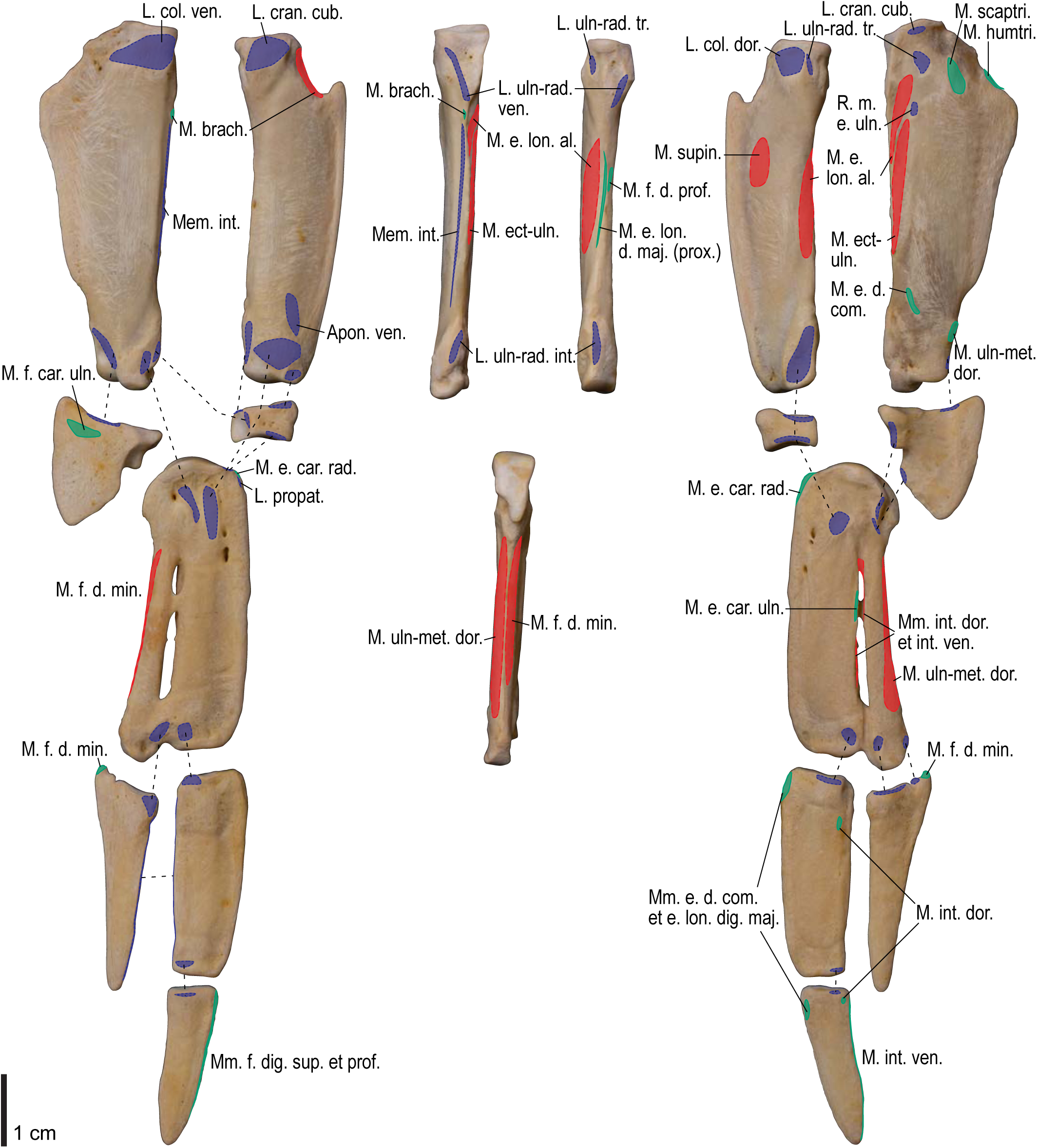
Osteological correlates of major wing muscles and ligaments in *Spheniscus humboldti*, distal wing elements. Drawn on NHMUK S/1961.15.1. See Figures 5 and 9 for legends.

#### Ligaments of the shoulder

The lig. acrocoracohumerale is a prominent ligament connecting the proximal end of the humerus to the processus acrocoracoideus of the coracoid (Fig. 3). Its origin on the coracoid is marked by a broad scar (impressio lig. acrocoracohumeralis) on the dorsolateral margin of the processus acrocoracoideus, typically between the facies articularis humeralis and the omal end of the coracoid (Figs. 4, 5, 10, 11). Its humeral insertion lies on the ventral margin of the sulcus transversus on the cranial aspect of the proximal humerus (Figs. 6, 7, 12, 13). In *Spheniscus*, the caudodorsal part of this ligament is somewhat differentiated, and could be termed the lig. coracohumerale dorsale; its origin extends onto the dorsal margin of the glenoid cavity, and its insertion is on the craniodistal margin of the sulcus transversus, adjacent to the typical insertion of the lig. acrocoracohumerale (Figs. 19, 21).

In most taxa examined, a thick, distinct ligament or retinaculum bridges between the lateral margin of the collum scapulae and the caudodistal margin of the caput humeri, providing an origin for the m. scapulotriceps (Fig. 3). This ligament is apparently not formally named in Baumel and Raikow (1993). Here, this ligament is tentatively referred to as the retinaculum originis m. scapulotricipitis. The scapular attachment of the retinaculum originis m. scapulotricipitis is marked by a tubercle on the lateroventral aspect of the collum scapulae (Figs. 4, 5, 10, 11). The retinaculum is closely associated with the caudal part of the joint capsule (plica synovialis transversa; below). The humeral end of this retinaculum is attached to the caudodistal and ventral margins of the caput humeri (Figs. 6, 7, 12, 13). This retinaculum is absent in *Spheniscus*, where the m. scapulotriceps arises directly from the scapula (see below).

The caudodorsal side of the shoulder joint capsule is sometimes developed as a distinct ligament that spans between the caudal margin of the glenoid cavity and the caudodistal margin of the caput humeri. This ligament is tentatively named the plica synovialis transversa. In most taxa (except *Spheniscus*), this ligament cannot be clearly differentiated from the retinaculum originis m. scapulotricipitis except at their proximal ends (the caudal margin of the glenoid cavity). In *Spheniscus*, where that retinaculum is absent, this ligament is distinctly developed, originating from a large area on the dorsal margin of the glenoid, and inserting on the caudal aspect of the caput humeri with a distinct scar (Figs. 19– 21).

#### Ligaments of the elbow

The lig. collaterale ventrale is a prominent ligament lying deep on the ventral side of the elbow joint, connecting the distal end of the humerus and the proximal end of the ulna (Fig. 3). Its humeral attachment is marked by a distinct tubercle (tuberculum supracondylare ventrale) lying proximoventral to the distal condyles of the humerus (Figs. 6, 7, 12, 13), whereas the ulnar attachment is marked by another tubercle (tuberculum lig. collateralis ventralis) on the ventral aspect of the proximal end of the ulna, just distal to the ventral margin of the cotyla ventralis (Figs. 8, 9, 14, 15).

A terminological clarification is required for the “lig. collaterale dorsale” in the avian elbow joint. Baumel and Raikow (1993: p. 163) state that this ligament is attached to the proximal end of the ulna, citing Stettenheim (1959). However, the same authors use this term to designate another ligament attached to the radius (Baumel and Raikow 1993: fig. 5.4). In fact, Stettenheim (1959: pp. 74–75) made clear that his use of the term was different from that in some previous studies, and development of this structure as a distinct ligament is apparently a unique feature of Charadriiformes (see below). Hence, Stettenheim’s (1959) structure attached to the ulna is here referred to as the lig. dorsale cubiti and the ligament attached to the radius as the lig. collaterale dorsale.

The lig. collaterale dorsale (as defined above) is a thin ligament on the dorsal side of the elbow joint connecting the distal end of the humerus and the proximal end of the radius (Fig. 3). It originates from the ventral and dorsal margins of the shallow groove between the condylus dorsalis and epicondylus dorsalis (Figs. 6, 7). In Alcidae, the origin also extends along the blunt crest extending distally from the tuberculum supracondylare dorsale (Figs. 12, 13). It is attached on the proximal aspect of the tubercle that lies on the dorsocranial margin of the cotyla humeralis of the radius, near the radial attachment of the meniscus radioulnaris (Figs. 8, 9, 14, 15).

As defined above, the lig. dorsale cubiti refers to a ligament on the dorsal side of the elbow joint that directly connects the humerus (or at least the proximal bellies of dorsal muscles of the forearm) and ulna (Fig. 2). This ligament is closely associated with the dorsal aponeurosis of the proximal forearm (aponeurosis dorsalis antebrachii) rather than the joint capsule; when present, this ligament is superficial to the m. extensor carpi ulnaris, m. supinator, m. extensor digitorum communis, and m. ectepicondylo-ulnaris. In *Pluvialis* and *Scolopax*, the ligament appears to arise from the dorsal surface of m. extensor digitorum communis, around the transition between the proximal tendon and fleshy belly (whose proximalmost parts are common with the m. supinator; see below). The osteological correlates of this ligament are not distinct in these taxa, but at least in *Scolopax*, the pit for the attachment of the mm. supinator et extensor digitorum communis on the humerus is slightly elongated proximally (Figs. S3, S4, S7, S8). In *Larus*, *Catharacta* and Alcidae, the ligament is relatively more distinct at its humeral origin; in *Larus* and *Catharacta*, the origin is marked by a faint depression proximoventrally adjacent to the pit for the mm. supinator et extensor digitorum communis (Figs. 6, 7, S9, S10). In Alcidae, it is marked by a separate tubercle lying proximal to the pit (Figs. 12, 13, S11–S14, S17, S18, S21, S22). In all these charadriiform taxa, the ligament inserts on the dorsocaudal surface of the proximal ulna, typically with a proximodistally elongated scar, but its distinctness from the attachment of the lig. limitans cubiti (below) varies: in *Pluvialis*, *Scolopax*, *Larus*, and *Catharacta*, the insertions of these ligaments are almost confluent with each other so that they cannot be distinguished on the bone (Figs. 9, S4, S8, S10); in Alcidae, the two insertions are separate from each other, with that for the lig. dorsale cubiti lying close to the craniodorsal margin of the bone and that for the lig. limitans cubiti lying near the caudal margin (Figs. 15, S12, S14, S18, S22). This ligament is also closely associated with the tendon of the m. extensor metacarpi ulnaris (see below) in *Larus*. In contrast to the conditions in Charadriiformes, the lig. dorsale cubiti is apparently absent in *Gavia*, Procellariidae, and *Spheniscus*. In *Gavia* and Procellariidae, the single scar on the dorsocaudal surface of the proximal ulna corresponds to the attachment of the aponeurosis dorsalis antebrachii associated with the lig. limitans cubiti or the dorsal branch of the lig. propatagiale (Figs. S28, S32). No direct ligamental connection is observed between the proximal ulna and the epicondylus dorsalis of the humerus in these taxa.

The lig. craniale cubiti is a broad but thin ligament lying deep on the cranial aspect of the elbow joint, connecting the forelimb bones to the distal end of the humerus. Its humeral origin lies along the proximal margins of the condyli dorsalis et ventralis on the cranial aspect of the humerus (Figs. 7, 13). It forms a major part of the joint capsule, and inserts on the cranial aspect of the proximal ulna just distal to the margins of the cotylae dorsalis et ventralis, and on the proximal radius along the ventral margin of the cotyla humeralis (Figs. 9, 15). In *Spheniscus*, the ventral portion of this ligament is exceptionally well-developed, with the attachments marked by distinct scars on the distal humerus and proximal radius; there is also a small branch, barely distinct from the joint capsule, that connects the proximal margin of the condylus dorsalis of the humerus and the dorsodistal margin of the proximal end of the ulna (Figs. 20–23).

The lig. radioulnare transversum (lig. cubiti teres in Stettenheim 1959) is a short but distinct ligament lying deep on the dorsal side of the elbow joint, bridging the proximal ends of the ulna and radius (Fig. 3). Its ulnar attachment lies on the dorsal aspect of the proximal ulna, typically within a convexity just distal to the dorsal margin of the cotyla dorsalis. The radial attachment is on the dorsodistal aspect of the tubercle on the dorsocranial margin of the cotyla humeralis, distal to the radial attachment of the lig. collaterale dorsale (Figs. 8, 9, 14, 15).

The meniscus radioulnaris is a thick ligament bridging between the proximal ends of the ulna and radius partly within the joint capsule (Fig. 3). Dorsally, it rims the articulation between the condylus dorsalis of the humerus and the cotyla dorsalis of the ulna. Its ulnar attachment lies along the caudoproximal margin of the cotyla dorsalis, but is poorly delineated as the area is covered by articular cartilage. After running along the dorsal margins of the cotyla dorsalis of the ulna and the cotyla humeralis of the radius, the meniscus ends on the dorsocranial margin of the latter cotyla, proximal to the tubercle that hosts the ligg. collaterale dorsale et radioulnare transversum (Figs. 9, 15).

Another ligament connects the proximal ends of the radius and ulna, deep within the interosseal space of the elbow joint. This ligament was termed “lig. transversum” by Stettenheim (1959), but was not treated by Baumel and Raikow (1993). Here it is referred to as the “lig. radioulnare ventrale” to avoid confusion with the lig. radioulnare transversum (above). The ulnar attachment is either restricted to the craniodistal margin of the proximal articular cotylae or extends distally along the distal leg of the cotyla dorsalis. The radial attachment is marked by a short, rugose ridge on the caudal (interosseal) aspect of the proximal end of the radius which extends ventrodistally from the rim of the cotyla humeralis (Figs. 8, 9, 14, 15).

The trochlea humeroulnaris is a retinaculum on the ventrocaudal aspect of the proximal ulna which braces the proximal tendon of the m. flexor carpi ulnaris (Figs. 2, 3). In all taxa examined except *Spheniscus*, where the presence of this structure was not confirmed, the trochlea primarily lies in a groove on the caudoventral margin of the proximal ulna (sulcus tendinosus), and is mainly formed by a ligamentous bridge spanning over the groove. At least the middle part of this ligament consists of two layers, forming a loop through which the tendon of m. flexor carpi ulnaris passes. The ligament is attached on both margins of the groove, one on the ventral aspect of the caudal margin of the ulna just distal to the olecranon, and the other caudal to the tuberculum lig. collateralis ventralis of the ulna (Figs. 9, 15). The pars humeralis accessoria of the trochlea (Bentz and Zusi 1982) was confirmed in *Gavia*, *Larus*, and *Catharacta*, but not in the other taxa examined; when present, it connects the main trochlea with the distal humerus, attached to the epicondylus ventralis caudodistal to the attachments of the lig. collateralis ventralis and m. pronator superficialis. However, the attachment site of the pars humeralis accessoria is hardly discernible on the bone. The main part of the trochlea humeroulnaris typically contains a sesamoid on the ventral part of the superficial layer.

The lig. tricipitale is a ligament lying deep within the caudal side of the elbow joint, anchoring the distal tendons of the mm. scapulotriceps et humerotriceps to the distal end of the humerus. The humeral attachment lies along most of the caudal margin of the fossa olecrani (Figs. 7, 13). Typically, it is also attached to the proximal end of the ulna caudal to the proximal cotylae (Figs. 9, 15).

#### Ligaments of the wrist and manus

The aponeurosis ventralis of the wrist is a broad aponeurosis which lies over the wrist musculature (Fig. 2). It spans from the ventral aspect of the distal radius to some of the remiges, while a portion (the so-called retinaculum flexorium) is attached on the tip of the processus pisiformis of the carpometacarpus, the ventrocranial tip of the crus longum of the ulnare, and, in *Catharacta*, the mid-shaft of the os metacarpale majus of the carpometacarpus (Figs. 9, 17). As such, it prevents the tendons of the mm. flexores digitorum superficialis et profundus from being displaced. The radial attachment of the aponeurosis is marked by a distinct tubercle (tuberculum aponeurosis ventralis) on the ventrocaudal aspect of the distal end of the radius (Figs. 8, 9, 14, 15). This feature is not correctly designated in a published illustration (Baumel and Witmer 1993: fig. 4.13).

The lig. radioulnare interosseum is a short ligament which connects the internal sides of the distal ends of the ulna and radius. Its ulnar attachment lies on the ventral margin of the depression (depressio radialis) on the cranial aspect of the distal end of the ulna, near the base of the tuberculum carpale, just proximal to the attachments of the ligg. ulno-radiocarpale interosseum et ventrale (see below). The radial attachment lies on the depression (depressio ligamentosa) on the caudoventral side of the distal end of the radius, just caudal to the tuberculum aponeurosis ventralis and proximal to the attachment of the lig. radio-radiocarpale ventrale (see below). These attachment scars are sometimes barely differentiated from adjacent ones (Figs. 9, 15).

The lig. ulno-ulnocarpale proximale is a broad ligament which connects the distal end of the ulna and the ulnare. It arises from the caudal surface of the tuberculum carpale of the ulna, and ends on the caudal part of the proximal surface of the ulnare, just dorsocranial to the tubercle for the lig. humerocarpale (Figs. 9, 15).

The lig. ulno-ulnocarpale distale is another ligament connecting the distal end of the ulna and the ulnare. Some variation in this ligament is evident among various charadriiform taxa. In *Pluvialis*, *Scolopax*, *Larus*, and *Catharacta*, this ligament arises from the tip of the tuberculum carpale and ends on the proximocranial aspect of the crus longum of the ulnare (Fig. 8, 9). In Alcidae, the ligament is apparently absent (or at least not distinct from the lig. ulno-ulnocarpale proximale), and the attachment site for this ligament is largely replaced by that of the lig. ulno-metacarpale ventrale (see below) (Fig. 15). It is unclear whether Stettenheim (1959) referred to this ligament by his “lig. obliquus carpi ulnaris”, as he did not specify the exact location of its insertion.

The lig. ulno-metacarpale ventrale was not confirmed in *Scolopax*, *Larus*, and *Catharacta*. When present, it arises from the distal aspect of the tip of the tuberculum carpale (Figs. 14, 15), directly connecting the ulna with the proximal end of the carpometacarpus. It ends in a distinct depression (fossa infratrochlearis) on the ventral surface of the proximal carpometacarpus proximocaudal to the processus pisiformis, along with the lig. radiocarpo-metacarpale ventrale (see below). In Alcidae, these two ligaments merge into a common ligament before insertion, so that their insertion sites cannot be told apart from each other (Fig. 17). In *Gavia*, this ligament merges into the aponeurosis ventralis to share the same insertion site on the ventrocaudal side of the processus pisiformis.

Both the lig. ulno-radiocarpale interosseum and lig. ulno-radiocarpale ventrale connects the distal end of the ulna and the radiale. These are not always clearly separated from each other; when they are (in *Larus* and *Uria*), the former arises from the sulcus intercondylaris of the ulna and ends on the caudal aspect of the radiale, whereas the latter arises more ventrally, near the distal aspect of the tuberculum carpale of the ulna, and ends in a relatively ventroproximal position on the radiale (Fig. S22).

The lig. radio-radiocarpale ventrale arises from the ventral aspect of the thickened distal end of the radius, just adjacent to the tuberculum aponeurosis ventralis, and ends on the ventral aspect of the radiale (Figs. 9, 15).

The lig. radio-radiocarpale dorsale usually arises from the tubercle on the dorsal margin of the distal end of the radius; however, in Alcidae, there seems to be a separate origin for the ligament along with the usual one, arising from the dorsal margin of the radius about one-tenth along the length of the bone from the distal end (Fig. 15). Both of these parts end on the proximodorsal margin of the radiale.

The meniscus intercarpalis is a thick, stiff ligament (or cartilage) lying within the wrist joint, bridging the gap between the radiale and ulnare. The radial side encloses nearly the entire caudodistal margin of the radiale, along the caudal margin of the facies articularis metacarpalis. The ulnar end lies in a depression on the proximal aspect of the tip of the crus breve of the ulnare (Figs. 8, 9, 14, 15).

The lig. radiocarpo-metacarpale craniale is a thin ligament lying on the cranioventral aspect of the wrist joint. The presence of this ligament was confirmed in most taxa examined, except *Catharacta*, *Gavia* and *Spheniscus*. When present, the ligament originates from the ventrodistal tip of the radiale, but its attachment is not clearly discernible on the bone. The ligament spreads before ending on the ventral aspect of the proximal carpometacarpus, along the cranial part of the ventral rim of the trochlea carpalis and the proximoventral margin of the os metacarpale alulare (Figs. 16, 17).

The lig. radiocarpo-metacarpale dorsale is a rather thin ligament on the dorsal side of the wrist joint. It arises near the dorsal tip of the facies articularis metacarpalis of the radiale, and ends on the proximal end of a slight depression (fossa supratrochlearis) on the dorsal surface of the proximal carpometacarpus, along with, but slightly proximal to, the attachment of the lig. ulnocarpo-metacarpale dorsale (see below).

The lig. radiocarpo-metacarpale ventrale is a distinct ligament lying on the ventral side of the wrist joint. The ligament originates from the ventral margin of the facies articularis metacarpalis of the radiale. As mentioned above, it ends in the fossa infratrochlearis of the carpometacarpus, along with the lig. ulno-metacarpale ventrale (Figs. 8, 9, 16, 17).

The lig. ulnocarpo-metacarpale ventrale is a short ligament connecting the ulnare and carpometacarpus on the ventral side of the wrist joint. It arises from the distal aspect of the tip of the crus longum of the ulnare, deep to the anchor of the retinaculum ventrale (Figs. 9, 15). In Alcidae, its insertion is marked by a depression lying just caudal to the processus pisiformis (distocaudal of the fossa infratrochlearis; Fig. 17), whereas in *Pluvialis*, *Scolopax*, *Larus*, and *Catharacta*, the attachment is elongated and lies near, but not along, the caudal margin of the ventral rims of the trochlea carpalis and fovea carpalis caudalis (Fig. 9). The insertion in Procellariidae is similar to that in Alcidae, but the depression is much less distinct (Figs. S31, S32). In *Gavia*, the insertion is marked by a distinct oval scar, rather than a depression (Figs. S27, S28). In *Spheniscus*, this ligament is not distinctly developed.

The lig. ulnocarpo-metacarpale dorsale is a ligament on the dorsal side of the wrist joint, and is much thicker than the lig. radiocarpo-metacarpale dorsale which lies adjacent to it. It arises from the distal aspect of the tip of the crus breve of the ulnare, distal to the attachment of the meniscus intercarpalis. It ends with a distinct scar near the proximal margin of the fossa supratrochlearis (Figs. 8, 9, 16, 17).

In most taxa examined (except *Spheniscus*), a part of the dorsal side of the complex wrist joint capsule is developed as a ligament or a retinaculum. This is treated as the “lig. ulnare externum metacarpi” in Stettenheim (1959), and is apparently not treated by Baumel and Raikow (1993). Hereafter, this ligament is tentatively referred to as the lig. ulno-metacarpale externum for terminological consistency. Proximally, this ligament is attached to the ulna, near the tip of the tubercle associated with the incisura tendinosa of the ulna, just cranial to the origin of the m. ulnometacarpalis dorsalis, although the corresponding attachment sites cannot be clearly discerned on the bone. It is also closely associated with the lig. radiocarpo-metacarpale dorsale, and is partly attached to the dorsal aspects of the radiale and ulnare. This ligament passes over the tendons of extensor muscles on the dorsal side of the wrist, and ends as a thin aponeurosis on the dorsal surface of the proximal carpometacarpus. In most taxa examined (except *Gavia*), the attachment scar is quite indistinct, but extends distocaudally from the distal end of the attachment of the lig. ulnocarpo-metacarpale dorsale (Figs. 9, 17). In *Gavia*, the attachment of this ligament is further apart distally from that of the latter ligament, and is marked by a distinct scar (Figs. S27, S28).

The lig. obliquum alulae is a distinct ligament on the alula, originating from the distocranial slope of the processus extensorius of the carpometacarpus and inserting on the cranioventral margin of the proximal end of the alular phalanx.

The lig. collaterale caudale (of artc. metacarpophalangealis alulae) lies deep within the alular articulation, connecting the caudal margins of the facies articularis alularis of the carpometacarpus and the proximal end of the alular phalanx.

The lig. collaterale ventrale (of artc. metacarpophalangealis digiti majoris) consists of two distinct parts on the ventral side of the joint. Both the cranial and caudal parts arise from the ventral side of the distal end of the carpometacarpus, where the attachments are marked by two distinct tubercles in Alcidae. The cranial part ends on the proximal end of the phalanx (or slightly offset from the proximal articular surface in Alcidae), whereas the caudal part ends on the caudal margin of the ventral surface of the proximal phalanx (Figs. 9, 17).

The lig. collaterale caudale (of artc. metacarpophalangealis digiti majoris) is present on the dorsocaudal aspect of the joint between the carpometacarpus and the proximal phalanx of the major digit. Its origin is marked by a tubercle which is slightly offset from the distal end of the carpometacarpus (near the level of the proximal margin of the symphysis metacarpalis distalis) and lies cranial to the sulcus interosseus. The insertion is on the dorsal part of the craniodorsal margin of the proximal articular surface of the phalanx (Figs. 9, 17).

The lig. obliquum intra-articulare (of artc. metacarpophalangealis digiti majoris) lies deep within the joint between the carpometacarpus and the proximal phalanx of the major digit. It originates from the groove between the two articular surfaces of the distal end of the carpometacarpus, and inserts on the caudal margin of the proximal articular surface of the phalanx (Figs. 9, 17).

The ligg. collaterale ventrale et dorsale (of artc. metacarpophalangealis digiti minoris) appear to be present in most taxa examined, but they are usually not quite differentiated from the articular capsule, and their attachment sites on the bones are hardly discernible. The lig. interosseum (of artc. interphalangealis lateralis) connects nearly the entire cranial margin of the minor digit to the caudal margin of the proximal phalanx of the major digit.

#### Accessory ligaments

The propatagium is spanned by a ligamental complex which typically consists of several interconnected ligamentous bands (Fig. 2). Following Baumel and Raikow (1993), the long ligamentous band forming the cranial edge of the propatagium is referred to as the lig. propatagiale, whereas the caudal band running along the humerus and inserting on the proximal forearm is referred to as the lig. limitans cubiti. In most taxa examined (except in *Spheniscus*, where these ligaments are undifferentiated), these two ligaments largely share the same origin.

In most taxa examined, the ligg. propatagiale et limitans cubiti together arise as the m. deltoideus pars propatagialis (and partly as the m. pectoralis pars propatagialis; see below). These are proximally anchored to the tip of the crista deltopectoralis of the humerus (Figs. 6, 7, 12, 13). In *Larus* and *Catharacta*, the ligaments arise separately from the distally bifurcated belly of the m. deltoideus pars propatagialis; the lig. propatagiale is further bifurcated at its proximal end, with the caudal branch anchored to the crista deltopectoralis. Typically, the middle part of the lig. propatagiale is flared and partly bifurcated, and around the flexion of the propatagium the cranialmost part is thickened and consists of elastic fibers (the so-called pars elastica). In *Gavia*, the pars elastica is rather enlarged, and the ligament consists almost entirely of elastic fibers except near the proximal and distal ends. In Procellariidae, the distal part of the ligament is largely bifurcated, and these divisions merge with each other near the wrist joint. In all cases, the lig. propatagiale passes the cranial edge of the wrist joint along the thickened cranioventral margin of the distal radius, where the ligament hosts a sesamoid (os prominens) in Procellariidae. The ligament inserts on the proximoventral margin of the processus extensorius of the carpometacarpus and the ventral margin of the proximal end of the alular phalanx (Figs. S4, S6, S18, S22), but the attachment sites on the bones are often hardly discernible.

Typically, two short branches (hereafter, the ventral and dorsal branches) arise around the pars elastica of the lig. propatagiale, inserting on the dorsal and ventral sides of the proximal forearm. The ventral branch is a thin ligament, and ends on the ventral fascia of the proximal forearm (aponeurosis ventralis antebrachii). In *Gavia*, the ventral branch is also anchored to the ventral surface of the belly of m. extensor carpi radialis (see below). Among the taxa examined, the conformation of the dorsal branch is rather variable. It is usually broad, lying on the most superficial layer of the dorsal surface of the forearm musculature. In *Gavia* and Procellariidae, the dorsal branch seems to be merged with the lig. limitans cubiti, and together these are attached to the dorsal fascia of the forearm (aponeurosis dorsalis antebrachii); in turn, the fascia is anchored to the dorsal aspect of the proximal ulna with an elongated scar. In Procellariidae, the dorsal branch is also attached to a sigmoidal sesamoid within the cranial side of the elbow joint, which has a ligamentous connection with the tip of the processus supracondylaris dorsalis of the humerus. In Charadriiformes, the dorsal branch of the lig. propatagiale merges either with the aponeurosis dorsalis antebrachii, with the lig. limitans cubiti, or with both of these. Either the dorsal branch of the lig. propatagiale or the lig. limitans cubiti is attached on the dorsal surface of the proximal ulna, along with the lig. dorsale cubiti (see above for the relative positions between the attachment sites). From the attachment scars alone, it is generally impossible to discern which of the dorsal branch of the lig. propatagiale or the lig. limitans cubiti is attached on the ulna.

The lig. limitans cubiti runs along the cranial margin of the humerus, caudal to the lig. propatagiale (Fig. 2). In Charadriiformes, it is more or less distinct from, and lies deep (ventral) to, the dorsal branch of the lig. propatagiale. In Alcidae, the proximal part of the lig. limitans cubiti is loosely connected to the cranial margin of the humerus, where a blunt, elongated ridge is present in some taxa (e.g., *Uria*, *Synthliboramphus*; Figs. S17, S18, S21, S22). Therefore, the ridge is considered to be an osteological correlate indicating strong attachment of the ligament to the humerus. As described above, it merges with the aponeurosis dorsalis antebrachii or ends on the ulna.

In *Spheniscus*, there is virtually no distinction between the ligg. propatagiale et limitans cubiti, and this single ligament is attached along the entire cranial margin of the crista deltopectoralis of the humerus (Figs. 20, 21). It then extends along the cranioventral margin of the radius up to its insertion on the processus extensorius of the carpometacarpus (Figs. 22, 23).

The lig. humerocarpale is a long, broad ligamentous band on the superficial layer of the ventral side of the forearm, connecting the distal humerus and the ulnare (Fig. 2). It arises from the caudodistal-most of two distinct pits on the ventral surface of the epicondylus ventralis (the other being for the m. pronator profundus; see below) (Figs. 6, 7, 12, 13). It ends on a distinct tubercle on the proximocaudal aspect of the ulnare, which lies ventral to the attachment of the lig. ulno-ulnocarpale proximale (see above) (Figs. 8, 9, 14, 15). In *Cepphus*, the ligament is also attached on the base of the processus pisiformis of the carpometacarpus (Fig. S14). The m. flexor digitorum superficialis arises from the deep surface of this ligament (see below). In *Spheniscus*, the ligament arises from the ventral surface of the caudodistal extension of the epicondylus ventralis (where no distinct pit is discernible), proximocranial to the origin of the m. flexor carpi ulnaris (Fig. 21). The ligament then becomes the m. flexor digitorum superficialis (which is entirely tendinous/ligamentous; see below) without attaching to the ulnare.

The retinaculum m. extensoris metacarpi ulnaris is a short retinaculum that anchors the proximal belly of the m. extensor carpi ulnaris to the dorsal aspect of the proximal ulna (this inconsistency in terminology is as per Baumel and Raikow [1993] and Vanden Berge and Zweers [1993]). The retinaculum lies deep to the lig. limitans cubiti, and also to the lig. dorsale cubiti when the latter ligament is present. The ulnar attachment of the retinaculum is often common with these ligaments, thus is not distinctly discernible on the bone (but see Figs. 23, S4, S14, S22).

#### Wing muscles

The m. rhomboideus superficialis is a thin, sheet-like muscle connecting the scapula with the vertebral column (Fig. 3). It lies deep to the m. latissimus dorsi cranialis and superficial to the m. rhomboideus profundus. The muscle arises as a thin fleshy sheet from the processus spinosi of several consecutive vertebrae (exact positions vary, but typically from the caudalmost one or two cervical and the cranialmost few thoracic vertebrae), and ends fleshily on the cranial part of the dorsal margin of the medial side of the scapular blade (Figs. 5, 11). In *Gavia* and Procellariidae, unlike in Charadriiformes, the insertion extends cranially to the medial aspect of some acromial ligaments (e.g., the lig. acromioclaviculare) and associated membranes (Figs. S24, S30).

The m. rhomboideus profundus is another muscle connecting the scapula with the vertebral column (Fig. 3). The muscle lies slightly caudally and deep to the m. rhomboideus superficialis, by which it is largely overlain. The muscle arises from the processus spinosi of several consecutive vertebrae (typically from the cranialmost to caudalmost thoracic vertebrae) with a partly aponeurotic origin, and ends fleshily on a broad area on the caudal part of the medial surface of the scapular blade, ventral to the attachment of the m. rhomboideus superficialis and dorsal to those of the mm. serratia (Figs. 5, 11).

Three more or less distinct muscles connect the scapula with the rib cage: the m. serratus superficialis pars cranialis, m. serratus superficialis pars caudalis, and m. serratus profundus. All of these muscles arise as partly separate aponeuroses from the lateral surfaces of some vertebral ribs; the first two typically arise from the dorsal margin of the processus uncinatus, whereas the last one arises from the facies lateralis of the rib body. Typical origins are the first two true (complete) ribs for the m. serratus superficialis pars cranialis, 3rd to 6th true ribs for the pars caudalis, and the last floating (incomplete) and the first few true ribs for the m. serratus profundus. The m. serratus superficialis pars cranialis ends as a thin but distinct aponeurosis on the margo ventralis of the scapular blade, between the two heads of the m. subscapularis (see below), marked by a sharp ridge. The m. serratus superficialis pars caudalis ends fleshily on the medial surface of the scapula around its caudal tip. The m. serratus profundus ends fleshily on the ventral area of the scapular blade, just cranial to the attachment of the previous muscle (Figs. 5, 11). The attachment sites of the last two muscles are hardly delineated on the bones. Another muscle, the m. serratus superficialis pars metapatagialis, was confirmed in most taxa examined, with the exception of *Spheniscus*. It arises as aponeuroses from the facies laterales and/or processus uncinati of a few caudal vertebral ribs, and ends fleshily on the dermis deep to the humeral feather tract.

The m. scapulohumeralis cranialis is a small muscle lying deep on the caudal aspect of the shoulder joint (Fig. 3). The muscle originates fleshily from the ventrolateral aspect of the collum scapulae, just caudoventral to the caudal tip of the facies articularis humeralis and slightly cranioventral to the attachment of the retinaculum originis m. scapulotricipitis (Figs. 5, 11); this area is marked by a slight depression in some alcids, e.g., *Uria* and *Synthliboramphus*. The muscle ends fleshily on a restricted area within the fossa tricipitalis of the humerus, just distal to the crus dorsale fossae, in the incision of the head of the m. humerotriceps (see below) (Figs. 7, 13). In many charadriiform taxa (*Larus*, *Catharacta*, and Alcidae), the humeral attachment is marked by a slightly elevated relief, whose margins are sometimes indistinct. This muscle is not present in *Spheniscus* (see also Schreiweis 1982).

The m. scapulohumeralis caudalis is a bulky muscle lying on the caudal aspect of the shoulder joint (Fig. 3). It arises fleshily from most of the lateral surface of the scapula that is unoccupied by other attachment sites (the origin is especially large in *Spheniscus*; Figs. 5, 11, 19). It inserts tendinously on the thickened part of the crus ventrale fossae (Figs. 7, 13), which slightly protrudes distally in some alcids (e.g., *Cerorhinca* and *Fratercula*; Fig. S12).

The mm. subcoracoscapulares complex lies deep within the caudal aspect of the shoulder joint (Fig. 3). The complex has three fleshy heads: the m. subscapularis caput laterale, m. subscapularis caput mediale, and m. subcoracoideus. The m. subscapularis caput laterale arises from the ventral part of the lateral surface of the scapular blade between the attachment sites of the retinaculum originis m. scapulotricipitis and the m. scapulohumeralis caudalis (Figs. 5, 11). The caput mediale has a somewhat larger origin on the cranial half of the medial surface of the scapular blade and neck (Figs. 5, 11). The margins of the attachment sites of these two heads are only faintly delineated. The two heads are separated by the aponeurosis of the m. serratus superficialis pars cranialis. In most taxa observed, the m. subcoracoideus arises largely from the dorsal surface of the membrana sternocoracoclavicularis around the processus procoracoideus (especially lig. intercoracoideum). The actual attachment on the coracoid, if any, is restricted to a small area of the dorsomedial aspect of the coracoidal body around the processus procoracoideus (Figs. 5, 11). Therefore, the attachment site of the muscle is not clearly discernible on the coracoid. The bellies of these heads merge into a common tendon, which then inserts into a pit on the proximal aspect of the tuberculum ventrale of the humerus (Figs. 7, 13).

The m. coracobrachialis cranialis is a short but bulky muscle on the cranial side of the shoulder joint (Fig. 3). Its origin is fleshy, and lies predominantly on the dorsal surface of the lig. acrocoracohumerale along its origin (Figs. 5, 11). As a result, the attachment on the coracoid cannot usually be traced on the bone. An exception is *Spheniscus*, where the muscle directly arises from the ventral aspect of the processus acrocoracoideus (Fig. 19). The muscle inserts on a depression (impressio coracobrachialis) on the dorsal part of the cranial aspect of the proximal humerus, which lies just ventrodistal to the attachment of the m. deltoideus pars minor and dorsal to the sulcus transversus (Figs. 7, 13). The depression is rather small in Alcidae and *Spheniscus* compared to the other taxa examined.

The m. coracobrachialis caudalis is a large muscle on the ventrocaudal aspect of the shoulder joint (Fig. 3). The muscle arises fleshily from the sternal end of the coracoid, on its ventral surface around the processus lateralis, and also from the adjacent lig. sternocoracoideum laterale (Figs. 5, 11). The medial border of the attachment site does not necessarily correspond with the linea intermuscularis ventralis of the coracoid. The muscle ends as a thick tendon on a distinct scar lying on the caudal aspect of the tuberculum ventrale of the humerus (Figs. 7, 13). In *Spheniscus*, the humeral insertion is displaced dorsally, and lies on the crus dorsale fossae (Fig. 21).

The m. pectoralis is the largest and the most superficial breast muscle (Fig. 2). The pars sternobrachialis of this muscle arises fleshily along the ventral part of the facies lateralis carinae of the sternum, with the dorsal margin marked by a linea intermuscularis, as well as from a large part of the lateral surface of the furcula and adjacent membrana sternocoracoclavicularis (Figs. 5, 11). In *Gavia*, the sternal origin of the pars sternobrachialis extends cranially past the apex carinae by ∼1 cm, resulting in a direct contact between muscle fibers on both contralateral sides of the muscle (Fig. S24). The pars costobrachialis of this muscle arises from the caudolateral part of the facies muscularis sterni and the associated membranae incisurarum sterni; the mediocranial border of this attachment site is marked by a faint ridge, which coincides with the linea intermuscularis in Alcidae (but not in the other taxa examined; see below). Despite its name, the pars costobrachialis does not directly arise from the rib cage in the taxa examined. The partes sternobrachialis et costobrachialis together form an overall bipennate structure of the muscle; most fibers insert on the aponeurosis intramuscularis, which then becomes a thick tendon of insertion. The tendon inserts on a distinct scar (impressio m. pectoralis) on the ventral aspect of the distal part of the crista deltopectoralis (and hence not on the entire surface of the crista deltopectoralis; Figs. 7, 13). The deep side of the fascia of this muscle is also partly attached to the ventral margin of the intumescentia humeri. Some of the cranialmost fibers of the pars sternobrachialis (or the so-called pars propatagialis) arising from the furcula do not contribute to this tendon, but instead merge with the m. deltoideus pars propatagialis.

In Procellariidae, a distinct part of the m. pectoralis is developed, known as the pars profundus (Kuroda 1960, 1961; Meyers and Stakebake 2005). This is a pinkish, fan-shaped muscle lying between the m. supracoracoideus and the main part of the m. pectoralis described above. This part arises both from the facies lateralis carinae and facies muscularis sterni. The carinal origin lies on the cranial part of the carina between the attachment sites for the m. pectoralis pars sternobrachialis and the m. supracoracoideus (Figs. S29, S30). As a result of this conformation, the attachment site of the pars sternobrachialis on the facies lateralis carinae is restricted to a small area near the ventral margin of the carina. The origin on the facies muscularis is restricted cranially, lying between the linea intermuscularis and the ridge for the lig. sternocoracoideum laterale. The distal tendon of this muscle lies on the deep side of the main parts of the m. pectoralis, and is closely associated with them. Hence, their insertions are in almost exactly the same place on the crista deltopectoralis of the humerus, and their separate attachment sites may not be clearly distinguished on the bone (Figs. S31, S32).

The m. supracoracoideus is a large, pennate muscle lying deep in the breast region (Fig. 3). It arises from the dorsocranial part of the facies lateralis carinae and the craniomedial part of the facies muscularis sterni, as well as from a restricted area of the membrana sternocoracoclavicularis adjacent to them (Figs. 5, 11). No muscle fibers of this muscle were confirmed to arise from the coracoid in the taxa examined. The attachment of this muscle on the sternum is clearly bordered by distinct ridges (lineae intermusculares), which form a subtriangular area in most taxa examined. The attachment site on the sternal plate is usually restricted craniomedially so that it is apart from that of the m. pectoralis pars costobrachialis, but it is rather extended caudolaterally in Alcidae and *Spheniscus*, resulting in direct contact between both attachment sites. In Alcidae, the passage of this muscle appears to be partly marked as a flattened scar on the medial surface of the coracoidal body. The muscle turns into a thick, flattened tendon as it passes the canalis triosseus which apparently acts as a pulley for this muscle. The tendon inserts on a distinct scar on the caudal aspect of the tuberculum dorsale of the humerus (Figs. 7, 13). In addition, the tendon is flared and partly bifurcated in Charadriiformes and *Gavia*, inserting also on a shallow furrow on the dorsal margin of the caput humeri (or the proximal aspect of the tuberculum dorsale), proximally adjacent to the main insertion (see also Kovacs and Meyer 2000).

The m. latissimus dorsi complex is among the most superficial muscles of the dorsum, lying superficial to the mm. rhomboideus and the scapula (Fig. 2). In most taxa examined, the partes cranialis et caudalis have distinct origins, passages, and insertions. The pars cranialis of this muscle is a thin, sheet-like muscle, which arises aponeurotically from the processus spinosi of a few consecutive vertebrae (typically from the caudalmost cervical vertebra to the cranialmost few thoracic vertebrae). The pars caudalis is somewhat bulkier, and arises as an aponeurosis spanning from more caudally positioned thoracic vertebrae (typically the caudalmost several thoracic vertebrae) to the area deep to the thigh musculature (e.g., mm. iliotibiales); in some taxa (e.g., *Catharacta* and *Spheniscus*), the entire origin of the pars caudalis lies deep to the thigh musculature. As they enter the brachium, these two parts pass deep to the proximal belly of the m. scapulotriceps and cross each other with the pars cranialis lying superficial (dorsal) to the pars caudalis. The pars cranialis ends fleshily on the dorsocaudal aspect of the proximal humerus, along a faint ridge (linea m. latissimi dorsi) extending distally from the tuberculum for the retinaculum m. scapulotricipitis (Figs. 7, 13). The pars caudalis turns into a tendon (or aponeurosis) which becomes closely associated with the retinaculum m. scapulotricipitis. Its insertion lies just ventrodistal to that of the retinaculum and ventral to the proximal margin of that of the pars cranialis (Figs. 7, 13). However, these attachment sites on the bone are sometimes hardly distinguishable from each other. The presence of the pars metapatagialis of this muscle was only confirmed in *Gavia* and one individual of *Larus schistisagus* examined in this study, in contrast to Hudson et al. (1969) who stated that this part was present in all larids and alcids they examined. This part might be damaged during skinning and overlooked in most birds examined here. In *Gavia*, this part arises as an aponeurosis from the processus spinosi of the few caudalmost thoracic and the cranialmost synsacral vertebrae. In *Larus*, the origin of this part is largely fused with that of the pars caudalis. In both cases, the pars metapatagialis ends on the dermis deep to the humeral feather tract. This muscle is highly modified in *Spheniscus* (see also Schreiweis 1982); the partes cranialis et caudalis pass through a ligamentous loop on the caudal side of the shoulder joint; both parts turn into a partly fused tendon which then inserts on a distinct tubercle lying distal to the crus dorsale fossae (Figs. 20, 21).

The m. deltoideus pars propatagialis is a moderately bulky muscle on the cranialmost part of the shoulder (Fig. 2). It arises fleshily from the dorsolateral margin of the furcula cranial to the processus acrocoracoideus claviculae (Figs. 5, 11). Just past the shoulder joint, it typically turns into the common ligament of the ligg. propatagiale et limitans cubiti; in *Larus* and *Catharacta*, however, the belly bifurcates before shedding the common ligament, so that the two ligaments arise separately from the muscle (see above). In the taxa examined, the muscle is consistently single-headed, and the delineation of multiple heads (see Vanden Berge and Zweers 1993) was not confirmed.

The m. deltoideus pars major is a bulky muscle on the dorsal aspect of the shoulder joint (Fig. 2). This muscle lies deep to the m. deltoideus pars propatagialis and typically superficial to the m. deltoideus pars minor and the tendon of the m. supracoracoideus. In most taxa, this muscle arises fleshily from the dorsolateral margin of the omal (dorsocaudal) end of the furcula and the adjacent lig. acromioclaviculare (Figs. 5, 11). In *Pluvialis*, *Gavia*, and Procellariidae, the origin is slightly extended caudally to reach the acromion of the scapula (Figs. S2, S24, S30). In *Spheniscus*, the origin seems to lie on the cranial end of the scapula between the facies articularis humeralis and acromion (Fig. 19). The insertion is usually fleshy (aponeurotic in *Gavia*), and lies on the dorsal aspect of the proximal humerus, with its relative position varying substantially among taxa: in *Gavia*, the insertion extends proximally from the middle part of the crista deltopectoralis and distally past the distal end of the crista deltopectoralis (Fig. S26); in Procellariidae, the insertion lies in a depression along the entire length of the crista deltopectoralis (Fig. S32); the conditions in *Pluvialis*, *Larus* and *Catharacta* are similar, but the insertion does not extend proximally past the tip of the crista deltopectoralis (Figs. 7, S4, S10); in *Scolopax*, where the crista deltopectoralis is relatively small, the insertion largely lies on the shaft and extends as far distally as the midshaft region (Fig. S8); in Alcidae, the insertion is restricted to a narrow area just craniodorsal to the insertions of the retinaculum m. scapulotricipitis and the m. latissimus dorsi partes cranialis et caudalis (Figs. 13, S12, S14, S18, S22). In *Spheniscus*, the insertion is restricted to a small area just proximal to the tubercle for the insertion of the two parts of the m. latissimus dorsi (Fig. 21).

The m. deltoideus pars minor is a two-headed muscle of the shoulder joint with a complex conformation (Fig. 3). The caput dorsale arises from the lateral aspect of the lig. acromioclaviculare or the lateral surface of the furcula (in *Gavia*), deep (ventral) to the origin of the m. deltoideus pars major. The caput ventrale lies ventral to it, arising from the dorsolateral margin of the membrana sternocoracoclavicularis along the coracoid, deep to the belly of the m. supracoracoideus (Fig. 11). The latter head is undeveloped in the non-alcid charadriiform taxa examined (*Pluvialis*, *Scolopax*, *Larus*, and *Catharacta*). In any case, the origins of these heads are not associated with any osteological correlates. A single belly is formed by the two heads as the muscle passes the canalis triosseus, where the belly lies dorsal to the tendon of the m. supracoracoideus. It ends largely fleshily (but partly tendinously in *Calonectris* and *Scolopax*, and exclusively tendinously in *Spheniscus*) on a restricted area on the proximal tip of the crista deltopectoralis of the humerus, with an indistinct scar (Figs. 7, 13).

The m. scapulotriceps is a prominent, two-joint muscle on the dorsocaudal aspect of the brachium (Fig. 2). In most taxa examined (except *Spheniscus*), the muscle fibers originate on the retinaculum originis m. scapulotricipitis (see above); its proximal belly is anchored to the dorsocaudal aspect of the humerus by a ligamentous retinaculum (retinaculum m. scapulotricipitis), whose position is clearly marked by a tubercle on the margo caudalis of the humerus, distal to the tuberculum dorsale and caudal to the crista deltopectoralis (Figs. 7, 13). In *Spheniscus*, both of these retinacula are absent (see above), and this muscle arises from two separate heads on the cranial end of the scapula. One head originates from the ventral margin of the scapula, and the other originates from the acromion (Fig. 19). In any case, around the proximal half of the brachium, the muscle turns to a thick ligament, which then passes the dorsal part of the fossa olecrani (sulcus scapulotricipitalis) of the distal humerus. The tendon of this muscle ends on the proximal end of the ulna, in a depression lying just caudal to the cotyla dorsalis (Figs. 9, 15).

The m. humerotriceps is a prominent, one-joint muscle on the caudal aspect of the brachium (Fig. 3). The origin is largely fleshy, and the head occupies most of the fossa (pneumo-)tricipitalis on the caudal aspect of the proximal humerus (Figs. 7, 13). The head is proximally incised by the crus dorsale fossae and the insertion of the m. scapulohumeralis cranialis (see above) in the taxa examined except *Spheniscus*. In most taxa examined, the portion of the head dorsal to the incision is extended so proximally that the dorsoproximal margin of the attachment more or less excavates the distal margin of the caput humeri (especially pronounced in some charadriiforms including *Larus* and *Fratercula*). The ventral portion of the head lies deeply in the ventral part of the fossa tricipitalis. The origin may extend as far distally as around the midshaft of the humerus, but its distal margin is not clearly discernible on the bone. Its bellies become a thick tendon near the elbow joint, which then passes the ventral part of the fossa olecrani (sulcus humerotricipitalis). The tendon of this muscle is closely associated with that of the m. scapulotriceps, and these ligaments are anchored to the fossa olecrani by the lig. tricipitale (see above). The tendon ends on the caudoproximal aspect of the olecranon of the ulna, where the attachment is marked by a prominent scar (Figs. 9, 15).

The m. biceps brachii is a two-joint muscle lying deep in the cranioventral aspect of the brachium (Fig. 3). This muscle appears to be absent in *Spheniscus* (see also Schreiweis 1982). Although two heads (the caput coracoideum and caput humerale) are recognized in the literature, the caput humerale is apparently absent in *Gavia*, Procellariidae, and most alcids (with the exception of *Uria*). The caput coracoideum arises tendinously from a distinct tuberculum on the ventral aspect of the processus acrocoracoideus of the coracoid (Figs. 5, 11). The caput humerale, when present, arises tendinously or aponeurotically from the proximal part of the crista bicipitalis of the humerus, just proximal to the insertion of the m. scapulohumeralis caudalis (Figs. 7, S4, S8, S10). This head is closely associated with the deep fascia of the m. pectoralis (see above). These two heads immediately merge into a single belly, which then turns into a tendon around the midshaft of the humerus. This tendon lies deep to the bellies of the m. brachialis and the mm. pronatores superficialis et profundus on the ventral aspect of the elbow joint. It then bifurcates just proximal to its terminus, and inserts on both the radius and ulna (Figs. 9, 15). The radial insertion is marked by a tuberculum (tuberculum bicipitale radii) on the caudal (interosseal) aspect of the proximal end of the bone, lying ventral to the attachment of the lig. radioulnare internum. The ulnar insertion is marked by another tuberculum (tuberculum bicipitale ulnae), which lies on the cranial (interossesal) aspect of the proximal ulna, slightly offset distally from the margins of the proximal cotylae and the attachment of the lig. radioulnare internum.

Another part of the muscle, m. biceps brachii pars propatagialis, is present in most taxa examined, regardless of the presence/absence of the caput humerale. This part arises as an aponeurosis, adjacently to the caput humerale, from the cranioventral aspect of the intumescentia humeri around the crista bicipitalis. The belly inserts either on the lig. propatagiale, the lig. limitans cubiti, or the common ligament of the two.

The m. brachialis is a bulky muscle lying deep in the cranial aspect of the elbow joint (Fig. 3). It arises fleshily from a distinct depression (fossa m. brachialis) on the cranial aspect of the distal humerus (Figs. 7, 13), and inserts fleshily onto another depression (impressio m. brachialis) on the ventrocranial aspect of the proximal ulna (Figs. 9, 15). This muscle is highly modified in *Spheniscus* (see also Schreiweis 1982); it arises fleshily from the cranial margin of the shaft distal to the crista deltopectoralis (Fig. 21), and mainly ends fleshily on the craniodorsal margin of the proximal radius. However, a small branch of fibers from this muscle also passes across the ventral side of the proximal radius and ends on the cranial margin of the ulna (Fig. 23).

The m. pronator superficialis is a small, fan-shaped muscle on the cranioventral aspect of the elbow joint (Fig. 2). It arises either tendinously (*Gavia*, *Pluvialis*, and *Scolopax*) or fleshily (other taxa examined) from the ventral aspect of the distal humerus, around the tuberculum supracondylare ventrale. The exact position and distinctness of the origin is rather variable among the taxa examined: it may arise without a distinct pit either from the ventrodistal margin of the tuberculum (in *Gavia*; Fig. S26), from the proximoventral margin (in Procellariidae and *Catharacta*; Figs. 7, S32), from the proximodorsal margin (in *Larus*; Fig. S10), from the ventral margin (in *Uria*; Fig. S22), or from the proximal margin (in *Synthliboramphus* and *Pluvialis*; Figs. S4, S18). Alternatively, it may arise from a distinct pit lying either proximoventral (in *Scolopax*; Fig. S8), proximal (*Cepphus* and *Alca*; Figs. 15, S14), or ventral (in *Cerorhinca* and *Fratercula*; Fig. S12) to the tuberculum. The belly becomes rather thin near its insertion site, and it inserts either fleshily (in Procellariidae, *Pluvialis*, *Catharacta*, and Alcidae) or aponeurotically (in *Gavia*, *Scolopax*, and *Larus*) on the ventrocranial margin of the proximal radius (Figs. 9, 15).

The m. pronator profundus is an elongated, fan-shaped muscle lying caudodistal to the previous muscle (Fig. 2). Its origin is tendinous, lying in a proximocranially positioned pit on the epicondylus ventralis of the humerus (one of the two distinct pits on the epicondyle) (Figs. 7, 13). The belly lies partly deep to the m. pronator superficialis, but is superficial to the lig. collaterale ventrale and to the m. brachialis. The insertion is fleshy, and occupies a large part of the ventrocaudal aspect of the radius (on the flattened ventral surface in Alcidae), extending past the midshaft of the bone in most taxa examined except in *Larus* and *Catharacta* (Figs. 9, 15). Both mm. pronator superficialis et profundus are lacking in *Spheniscus* (see also Schreiweis 1982).

The m. flexor carpi ulnaris is a distinct, two-joint muscle on the caudoventral aspect of the antebrachium (Fig. 2). The muscle arises tendinously from the distal aspect of the distal extension of the epicondylus ventralis of the humerus (processus flexorius), where the attachment is marked by a prominent scar (Figs. 7, 13). The tendon passes the trochlea humeroulnaris (see above) and then runs along the caudoventral margin of the ulna. Along the antebrachium, the caudal margin of the belly (the so-called pars remigalis) is associated with the lig. elasticum interremigale minor which spans the bases of the secondaries. The insertion is tendinous, lying on the concavity on the caudal aspect of the ulnare (Figs. 9, 15). In *Gavia*, the distal tendon is partly ossified.

The m. flexor digitorum superficialis is a thin, multi-joint muscle whose belly lies on the ventral aspect of the antebrachium (Figs. 2, 3). As mentioned above, the belly of this muscle arises from the deep surface of the lig. humerocarpale in the mid-distal part of the antebrachium. It soon turns into a thin tendon around the wrist joint, and then passes the retinaculum on the proximoventral aspect of the crus longum of the ulnare, where the passage is marked by a sulcus and bony canal in some taxa (e.g., in *Larus* and *Catharacta*; Figs. 9, S10). After passing the caudal side of the processus pisiformis of the carpometacarpus, the tendon of this muscle runs along the ventral margin of the os metacarpale majus, being parallel and deep to that of the m. flexor digitorum profundus. As these two tendons remain in close association along the remainder of their trajectories, it is not easy to distinguish the insertions of these muscles from one another. Nevertheless, the m. flexor digitorum superficialis appears to end on the ventrocranial margins of the proximal ends of either or both of the proximal and distal phalanges of the major digit (Figs. 9, 17). In Alcidae, the insertion on the distal phalanx lies on an elongated area on the ventral margin of the bone, rather than on its proximal end. In *Spheniscus*, this muscle is continuous with the lig. humerocarpale, and is entirely tendinous (see also Schreiweis 1982); it merges with the tendon of the m. flexor digitorum profundus and ends on the cranioventral margin of the distal phalanx of the major digit (Fig. 23).

The m. flexor digitorum profundus is a slender, multi-joint muscle whose belly lies deep in the interosseal space of the antebrachium (Figs. 2, 3). The muscle arises fleshily from the cranial (interosseal) aspect of the ulna, with the extent and exact position of the origin being rather variable among taxa. In *Gavia*, the origin occupies a large part of the midshaft of the ulna between the attachments of the m. brachialis and m. ulnometacarpale ventrale (Fig. S28). In Procellariidae, it is restricted to a small area distal to the attachment of the m. brachialis, and does not reach the midpoint of the bone (Fig. S32). In *Scolopax*, it extends proximally from that position, reaching as far proximally as the attachments of the m. biceps brachii and lig. radioulnare ventrale, and also extends proximoventrally to the area ventral to the impressio m. brachialis (as a result, the area of attachment is proximally incised by the impressio m. brachialis; Fig. S8). In *Pluvialis*, *Larus*, and *Catharacta*, it occupies a similar position to that in *Scolopax*, but the origin ventral to the impressio m. brachialis is not continuous with the main origin, thereby essentially forming a small, separate head (Figs. 9, S4, S10). Also, in these taxa, the distal margin of the attachment lies more proximally due to the proximal extension of the attachment of the m. ulnometacarpale ventrale. In Alcidae, the origin does not extend as far distally as in other taxa, and the distal end of the origin either lies more proximal to the distal margin of the impressio m. brachialis (in *Uria*, *Alca*, and *Cepphus*; Figs. 15, S14, S22), or extends only slightly further distally than that impression (in *Synthliboramphus*, *Cerorhinca*, and *Fratercula*; Figs. S12, S18). In *Spheniscus*, the origin of this muscle is so closely associated with the membrana interossea antebrachii that an attachment site of the muscle separate from the membrane cannot be identified on the bone. Past the middle antebrachium, the muscle turns into a thin tendon, which passes below the aponeurosis ventralis. In the proximal manus, the tendon changes its direction on the cranial side of the processus pisiformis of the carpometacarpus, which acts as a pulley for this muscle. The tendon then goes on the cranioventral margin of the os metacarpale majus, superficial to that of the m. flexor digitorum superficialis, with which it is partly associated. The main insertion of this muscle lies on the cranioventral margin of the proximal end of the distal (second) phalanx of the major digit (Figs. 9, 17).

The m. extensor carpi radialis is a prominent, two-joint muscle on the craniodorsal aspect of the antebrachium (Fig. 2). It usually arises from the tuberculum supracondylare dorsale of the humerus (which is developed into the processus supracondylaris dorsalis in Procellariidae, *Pluvialis*, *Larus*, and *Catharacta*) with two closely associated heads: the largely tendinous caput dorsale and the fleshy caput ventrale. In most taxa lacking the processus supracondylaris dorsalis (*Gavia*, *Scolopax*, and Alcidae), these two heads are not always distinguishable, and arise adjacently from the tuberculum supracondylare dorsale (Figs. 12, 13, S7, S8, S25, S26). In *Larus* and *Catharacta*, where the tuberculum is well-developed into a processus, the caput dorsale is restricted around the tip of the dorsal side of the processus, whereas the caput ventrale arises from the middle, rather than the tip, of the ventral surface of the processus (Figs. 6, 7, S9, S10). On the other hand, in Procellariidae, although the caput ventrale arises from the ventral surface of the processus near the base, the caput dorsale does not directly arise from the processus, but from the distal margin of the sesamoid connected to the processus; the tip of the processus is occupied by bilateral ligaments for the sesamoid (Figs. S31, S32). The belly of the caput dorsale often receives anchors from the lig. limitans cubiti and branches of the lig. propatagiale. The bellies of the two heads merge in the proximal antebrachium, and then turn into a thick tendon around the middle of the antebrachium. In *Spheniscus*, the distinction between the two heads is indistinct and the muscle arises from an indistinct scar on the dorsal surface of the distal humerus (Figs. 20, 21). The tendon passes a ligamentous retinaculum formed in the sulcus tendinosus of the distal radius, where it merges with the tendon of the m. extensor longus alulae, and then ends on the proximocranial tip of the os metacarpale alulare (processus extensorius) of the carpometacarpus (Figs. 9, 17).

The m. extensor carpi ulnaris is a long, two-joint muscle on the dorsal antebrachium (Fig. 2). It arises tendinously from the caudodistal of two pits on the epicondylus dorsalis of the humerus, in a common tendon with the m. ectepicondylo-ulnaris (Figs. 7, 13). It is also anchored to the dorsocranial margin of the proximal ulna by the retinaculum m. extensoris metacarpi ulnaris (see above). The belly passes through the dorsocaudal part of the interosseal space of the antebrachium, and turns into a tendon in the distal antebrachium. The tendon passes through the incisura tendinosa of the distal ulna which acts as a pulley for this muscle. The tendon ends on a distinct tubercle on the caudodorsal aspect of the os metacarpale majus around the symphysis metacarpalis proximalis of the carpometacarpus (Figs. 9, 17).

The m. extensor digitorum communis is a long, multi-joint muscle on the dorsal aspects of the antebrachium and manus (Figs. 2, 3). It arises from the cranioproximal of two pits on the epicondylus dorsalis of the humerus, in a common tendon with the m. supinator (Figs. 7, 13). In *Pluvialis* and *Scolopax*, the proximal part of this muscle is anchored to the dorsal aspect of the proximal ulna by the lig. dorsale cubiti (see above). The belly of this muscle lies in the dorsal aspect of the interosseal space of the antebrachium, just cranial to the m. extensor carpi ulnaris. The belly of this muscle turns into a thin tendon near the wrist joint, which then passes through the incisura tendinosa of the distal ulna, along with that of the m. extensor carpi ulnaris. On the dorsal side of the proximal manus, the tendon of this muscle lies superficial to that of the m. extensor carpi ulnaris, and bifurcates near the os metacarpale alulare. The cranial branch ends on a tubercle on the dorsocaudal tip of the proximal end of the alular phalanx (Figs. 9, 17). By contrast, the caudal branch runs distocaudally along the os metacarpale majus, crosses with the tendon of the m. extensor longus digiti majoris which lies superficial to it, and lies within the sulcus tendinosus on the dorsocaudal aspect of the shaft. The tendon finally ends on the dorsal aspect of the proximal end of the proximal phalanx of the major digit, just cranial to the scar for the lig. collaterale caudale (Figs. 9, 17). In *Spheniscus*, this muscle is present in much the same conformation, except that it completely lacks the branch leading the alular phalanx; the distal tendon of this muscle is merged with that of the m. extensor longus digiti majoris, and inserts on the proximal ends of the two phalanges of the major digit (Fig. 23).

The m. extensor longus alulae is a thin muscle lying deep within the antebrachium (Fig. 3). In most taxa examined, the muscle has two heads, on the proximal ulna and on the midshaft of the radius. The ulnar head is partly tendinous (entirely fleshy in *Spheniscus*), and arises from the cranial aspect of the ulna just distal to the proximal articular surfaces (Figs. 9, 15); in Alcidae, this head lies in a small concavity formed by the hook-like distal extension of the cotyla dorsalis; otherwise, this origin is so vaguely marked that it is hardly discernible on the bone (perhaps except in *Spheniscus*). Presence of the ulnar head was not confirmed in Procellariidae and *Larus*. By contrast, the radial head was confirmed to be present in all taxa examined. This head arises fleshily from the caudodorsal aspect of the proximal radius, between the proximal end of the m. extensor longus digiti majoris and the distal end of the m. supinator (Figs. 9, 15). This origin lies dorsal to the linea intermuscularis on the caudal (interosseal) margin of the radius, but is not clearly demarcated on the bone. The two heads merge in the interosseal space, and the resultant belly crosses the dorsal side of the radius to lie within the sulcus tendinosus of the distal radius, where it turns into a thin tendon. The tendon runs alongside that of the m. extensor carpi radialis, with which it merges before inserting on the processus extensorius of the carpometacarpus.

The m. extensor longus digiti majoris is a multi-joint muscle on the dorsal aspects of the antebrachium and manus (Fig. 2). The muscle seems to have two separate heads, partes proximalis et distalis. The pars proximalis arises fleshily from the caudodorsal aspect of the radius (Figs. 9, 15), typically occupying a large area of the radial shaft caudodorsal to the attachment of the m. pronator profundus (except in *Larus*, where the origin is restricted to the midshaft region, and in *Spheniscus*, where the origin is largely aponeurotic; Figs. 23, S10). The proximal margin of the origin is marked by a convergence of two lineae intermusculares. The belly turns into a thin tendon which passes the wrist joint along the dorsal rim of the trochlea carpalis of the ulna, just cranioventral to the incisura tendinosa. The much more indistinct pars distalis arises fleshily from the dorsal aspect of the proximal manus, but its origin in most taxa examined lies on the ligaments and aponeuroses spanning between the carpal bones, thus it does not typically correspond to osteological correlates. An exception is *Gavia*, where the pars distalis arises from the dorsal aspect of the os metacarpale majus, although the origin is only indistinctly marked (Fig. S28). In *Spheniscus*, the pars distalis is absent (see also Schreiweis 1982). In the proximal manus, the tendon from the pars proximalis lies along the dorsocaudal margin of the os metacarpale majus, but distally it crosses with the distal branch of the tendon of the m. extensor digitorum communis, lying superficial to the latter. In all taxa except *Gavia* and *Spheniscus*, the tendons from both parts merge to form a common tendon. When present, this common tendon then passes the cranial aspects of the metacarpo-phalangeal and interphalangeal joints, and ends on the cranial aspect of the proximal ends of the proximal and distal phalanges of the major digit (Figs. 9, 17). In *Gavia*, where the major digit has three free phalanges, the tendon of the pars proximalis inserts on the second phalanx of the major digit, whereas that of the pars distalis inserts on the first (proximal) phalanx (Fig. S28).

The m. supinator is a fan-shaped muscle on the dorsocranial aspect of the elbow joint (Fig. 2). The muscle arises in a common tendon with the m. extensor digitorum communis from the epicondylus dorsalis of the humerus (see above). The insertion is fleshy, and lies on the cranial margin of the dorsal aspect of the radial shaft (Figs. 9, 15). The insertion expands somewhat craniodistally in *Gavia* (Fig. S28), whereas it is rather restricted in area in *Larus* and *Catharacta* (Figs. 9, S10).

The m. ectepicondylo-ulnaris is a large muscle on the dorsal aspect of the elbow joint (Fig. 3). This muscle arises in a common tendon with the m. extensor carpi ulnaris from the epicondylus dorsalis of the humerus (see above). The belly lies deep (ventral) to that of the m. extensor carpi ulnaris, and inserts fleshily on the craniodorsal aspect of the ulnar shaft (Figs. 9, 15). The insertion is extended distally in *Gavia*, *Scolopax*, and Alcidae, extending well past the midshaft region (Figs. 15, S8, S28), whereas it is restricted to the proximal part of the ulnar shaft in Procellariidae, *Pluvialis*, *Larus*, and *Catharacta* (Figs. 9, S4, S10, S32). The ventral margin of the insertion is marked by a distinct linea intermuscularis on the proximal ulna, ventral to which the m. flexor digitorum profundus typically lies (except in *Spheniscus*, where that muscle does not directly attach to the ulna).

As expected, the m. entepicondylo-ulnaris was absent in all taxa examined; the muscle is apparently unique to some members of Palaeognathae and Galloanseres (e.g., Vanden Berge and Zweers 1993).

The m. ulnometacarpalis dorsalis is a fan-shaped muscle on the dorsocaudal aspect of the wrist joint (Fig. 3). The muscle seems to arise tendinously from the dorsal aspect of the distal ulna, adjacent to the protruding tubercle for the incisura tendinosa (Figs. 9, 15), but the attachment site is not clearly marked on the bone (except in *Spheniscus*, where it is marked by a tubercle; Figs. 22, 23). The insertion is fleshy, typically lying along the caudal margin of the os metacarpale minus around the symphysis metacarpalis proximalis of the carpometacarpus (Figs. 9, 17) (rather extended distally in *Spheniscus*). The attachment site is sometimes (e.g., in Procellariidae and *Catharacta*) incised distally by the origin of the m. flexor digiti minoris (Figs. 9, S32).

The m. ulnometacarpalis ventralis is a muscle on the wrist joint with a complicated passage (Figs. 2, 3). The muscle arises fleshily from the cranioventral aspect of the distal ulnar shaft, distal to the attachments of both of the m. flexor digitorum profundus and m. brachialis (Figs. 9, 15). In Procellariidae, the head is slightly bifurcated, and the origin also extends caudoproximally on the ventral aspect of the distal ulna from its distal end (Fig. S32). The belly turns into a distinct tendon before entering the wrist joint from the ventral side. In the wrist joint, the tendon turns around the joint on the cranial side, along a distinct, diagonal sulcus on the cranial aspect of the radiale; here, the tendon of this muscle lies deep to those of the m. extensor carpi radialis and m. extensor longus alulae. The tendon then inserts on a distinct depression on the proximocranial part of the dorsal rim of the trochlea carpalis of the carpometacarpus (Figs. 9, 17). This muscle is absent in *Spheniscus* (see also Schreiweis 1982).

The m. interosseus dorsalis is a small muscle on the dorsal aspect of the manus (Fig. 2). The muscle arises fleshily along the dorsal margin of the interosseal sides of the ossa metacarpalia majus et minus, including the symphysis metacarpalis proximalis (Figs. 9, 17). The tendon passes a retinaculum formed on the caudal aspect of the distal end of the carpometacarpal shaft (at the symphysis metacarpalis distalis), and then runs along a distinct sulcus on the dorsal aspect of the proximal phalanx of the major digit. It then ends on the dorsal apex of the proximal end of the second phalanx of the major digit (Figs. 9, 17); in most charadriiform taxa, the tendon also extends distally to attach on the dorsal margin of the phalangeal shaft. In *Spheniscus*, the tendon of this muscle runs along the caudodorsal margin of the bone, and is also attached to the proximal phalanx of the major digit (Fig. 23).

The m. interosseus ventralis is another muscle on the interosseal space of the manus (Figs. 2, 3). It arises fleshily from the interosseal space of the carpometacarpus, just ventral to the origin of the previous muscle (Figs. 9, 17). The origin of the m. interosseus ventralis seems to extend further distally than that of the previous muscle, extending nearly to the symphysis metacarpalis distalis. The tendon of this muscle passes another retinaculum on the dorsal aspect of the symphysis metacarpalis distalis of the carpometacarpus. The tendon runs along the caudal margin of the proximal phalanx of the major digit, and then parallel to, but separately from, the caudal margin of the second phalanx of the major digit. The tendon then attaches to a distinct eminence on the caudal aspect of the distal end of the second phalanx (Figs. 9, 17), except in *Spheniscus*, where the eminence is absent and the tendon attaches along the entire caudal margin of the bone (Fig. 23). Before the final insertion, the tendon may be attached to the distal end of the proximal phalanx (in *Larus* and *Cepphus*; Figs. S10, S14) or the proximal end of the distal phalanx (in *Catharacta, Cerorhinca*, *Fratercula*, and *Synthliboramphus*; Figs. 9, S12, S18).

The m. extensor brevis alulae is a small, fan-shaped muscle on the dorsal aspect of the alula (Fig. 2). The muscle arises fleshily from a depression on the dorsal aspect of the os metacarpale alulare of the carpometacarpus, particularly around the base of the processus extensorius (Figs. 9, 17). Although the margins of the origin are not clearly marked on the bone, the origin appears to be proximodistally broad in those taxa with a proximodistally elongated os metacarpale alulare (*Gavia*, *Uria*, *Alca*, and, to some extent, *Synthliboramphus*; Figs. 17, S18, S22, S28). The muscle inserts tendinously on the craniodorsal margin of the proximal end of the alular phalanx, cranial to the insertion of the m. extensor digitorum communis (Figs. 9, 17).

The m. abductor alulae is a small muscle on the cranioventral aspect of the alula (Fig. 2). It arises from the ventrocaudal aspect of the tendon of the m. extensor carpi radialis near its distal end, hence its origin does not have any osteological correlates. Its belly lies along the ventral aspect of the processus extensorius of the carpometacarpus, and its insertion lies on the cranial margin of the proximal end of the alular phalanx, distal to the attachment of the lig. obliquum alulae (Figs. 9, 17). In the taxa examined, with the exception of Alcidae, the insertion extends slightly distally along the cranial margin of the phalanx.

The m. flexor alulae is a small, fan-shaped muscle on the ventral aspect of the alula (Fig. 3). The muscle arises fleshily from the caudal part of a depression on the ventral aspect of the os metacarpale alulare of the carpometacarpus (Figs. 9, 17). In *Gavia*, where the os metacarpale alulare is proximodistally elongated, the origin is restricted to the distal half of the metacarpal body (Fig. S28). The muscle ends tendinously on the dorsocaudal aspect of the proximal end of the alular phalanx (Figs. 9, 17).

The m. adductor alulae is a small muscle on the ventral aspect of the alula (Figs. 2, 3). The muscle arises fleshily from the area distal to the facies articularis alularis of the carpometacarpus, along the major metacarpal shaft just distal to the ventral margin of the articular surface (Figs. 9, 17). The muscle inserts on much of the caudal aspect of the body of the alular phalanx (Figs. 9, 17). Neither the origin nor insertion can be clearly discerned on the bones.

*Spheniscus* lacks all of the intrinsic muscles associated with the alula, including the mm. extensor brevis alulae, abductor alulae, flexor alulae, et adductor alulae (see also Schreiweis 1982).

The m. abductor digiti majoris is a small muscle on the ventrocranial aspect of the manus (Fig. 3). The muscle arises fleshily from the ventrocranial aspect of the shaft of the os metacarpale majus, typically along an elongated area on the metacarpal shaft (Figs. 9, 17). In most taxa examined, the proximal margin of the origin usually reaches the area just cranial to the processus pisiformis. However, it ends on the area just distal to the processus in *Scolopax* (Fig. S8), whereas it does not extend as far proximally in *Cerorhinca* and *Fratercula* (Fig. S12). In *Larus* and *Cepphus*, the origin is separated into two parts lying on the proximal and distal ends of the typical origin (Figs. S10, S14). Its insertion is tendinous, and lies typically on the cranioventral aspect of the proximal end of the proximal phalanx of the major digit (Figs. 9, 17). In *Scolopax*, the tendon also extends to the distal end of the phalanx (Fig. S8). In Procellariidae, the insertion lies on the second phalanx, rather than the proximal phalanx, of the major digit (Fig. S32).

The m. flexor digiti minoris is a small muscle on the caudal aspect of the manus (Fig. 3). Its origin is fleshy, and occupies a large part of the caudal aspect of the os metacarpale minus that is not occupied by the m. ulnometacarpalis dorsalis (Figs. 9, 17). Its insertion is tendinous, and lies on a prominence on the caudal margin of the phalanx of the minor digit (Figs. 9, 17), which is developed into a proximally protruding process in *Spheniscus* (Figs. 22, 23).

### Reconstructed musculature in extinct auks

From osteological correlates observable on fossil and subfossil bones it was possible to reliably infer (Level I inference; Witmer 1995) the presence of most of the wings muscles and ligaments described above in *Pinguinus* and *Mancalla* (Figs. 24–35). The positions of the attachment sites of most of these muscles and ligaments could also be determined from osteological correlates, although it was not feasible to delineate the margins of some fleshy attachment sites (e.g., m. deltoideus pars major, m. humerotriceps). For some muscles lacking clear osteological correlates in extant taxa, it was necessary to rely on Level I’ inference (Witmer 1995) to infer their presence in the extinct taxa (e.g., the m. flexor digitorum superficialis which arises from the lig. humerocarpale). Given the lack of associated complete skeletons in *Pinguinus* and *Mancalla*, the extent and positions of the muscles attached to vertebrae, ribs, carpal elements and phalanges were unclear (e.g., mm. rhomboidei, latissimus dorsi, serratia, digital flexors and extensors). In addition, inferences regarding the presence of some structures in *Mancalla* were equivocal from character optimization alone: examples include the caput humerale of the m. biceps brachii (which is present in the two successive outgroups, *Larus* and *Catharacta*, but absent in crown-group Alcidae), and the caput ventrale of the m. deltoideus pars minor (developed in crown-group Alcidae, but not in the other charadriiform taxa examined). For such muscles, it is tentatively considered here (Level II or II’ inference) that the conditions of these characters in *Mancalla* were similar to those of crown-group Alcidae, as *Mancalla* was probably a wing-propelled diver (which itself is an inference; see below), and shares numerous osteological features with crown-group Alcidae. On this basis, the overall wing musculature of *Pinguinus* and *Mancalla* was reconstructed as illustrated in Figures 36–39.

**Figure 24.**
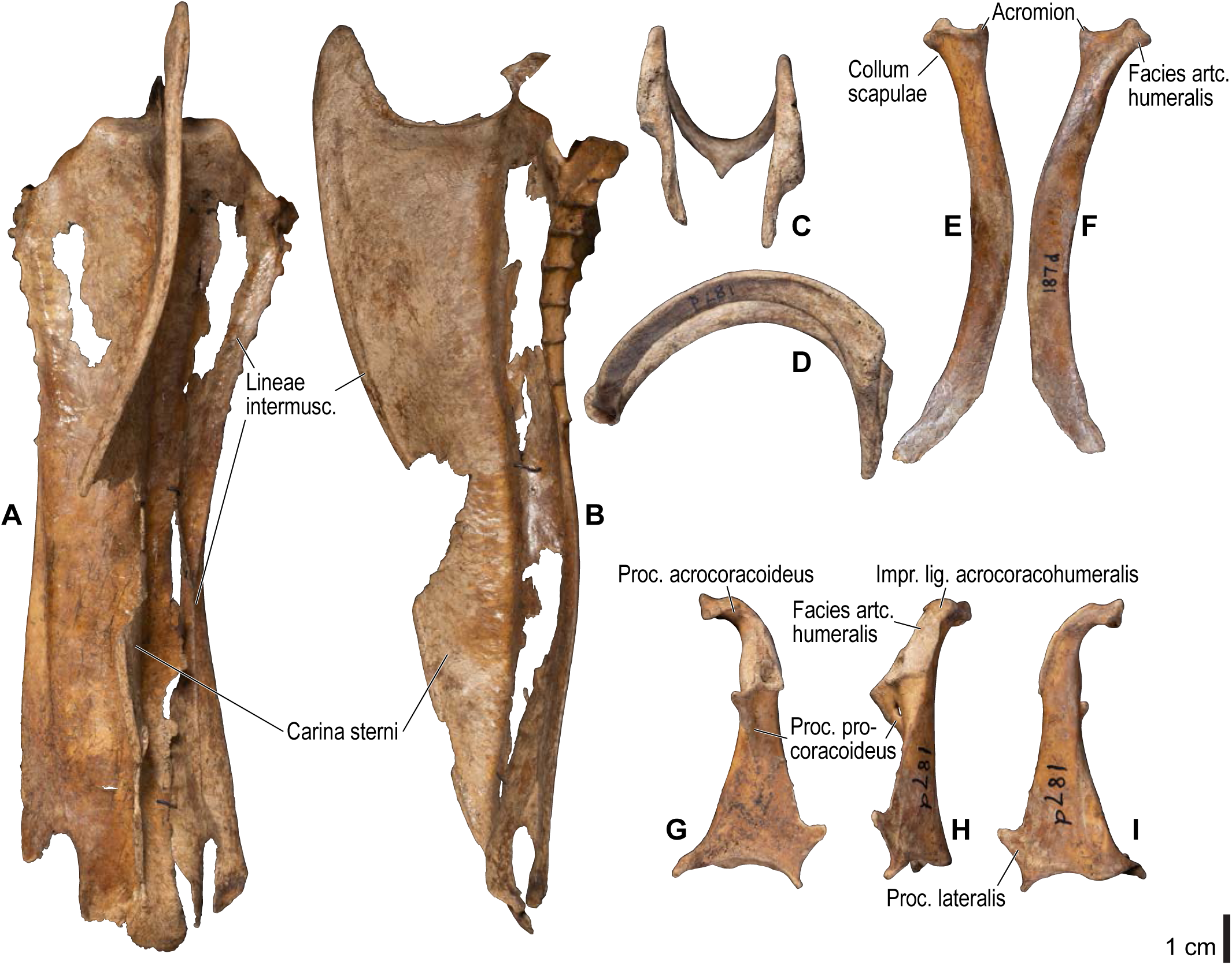
Osteology of *Pinguinus impennis*, pectoral girdle elements. Drawn on UMZC 187.G (sternum) and 187.d (other elements). Sternum in ventral (**A**) and left lateral (**B**) views; furcula in dorsal (**C**) and left lateral (**D**) views; right scapula in medial (**E**) and lateral (**F**) views; right coracoid in dorsal (**G**), lateral (**H**), and ventral (**I**) views. See Figure 4 for abbreviations.

**Figure 25.**
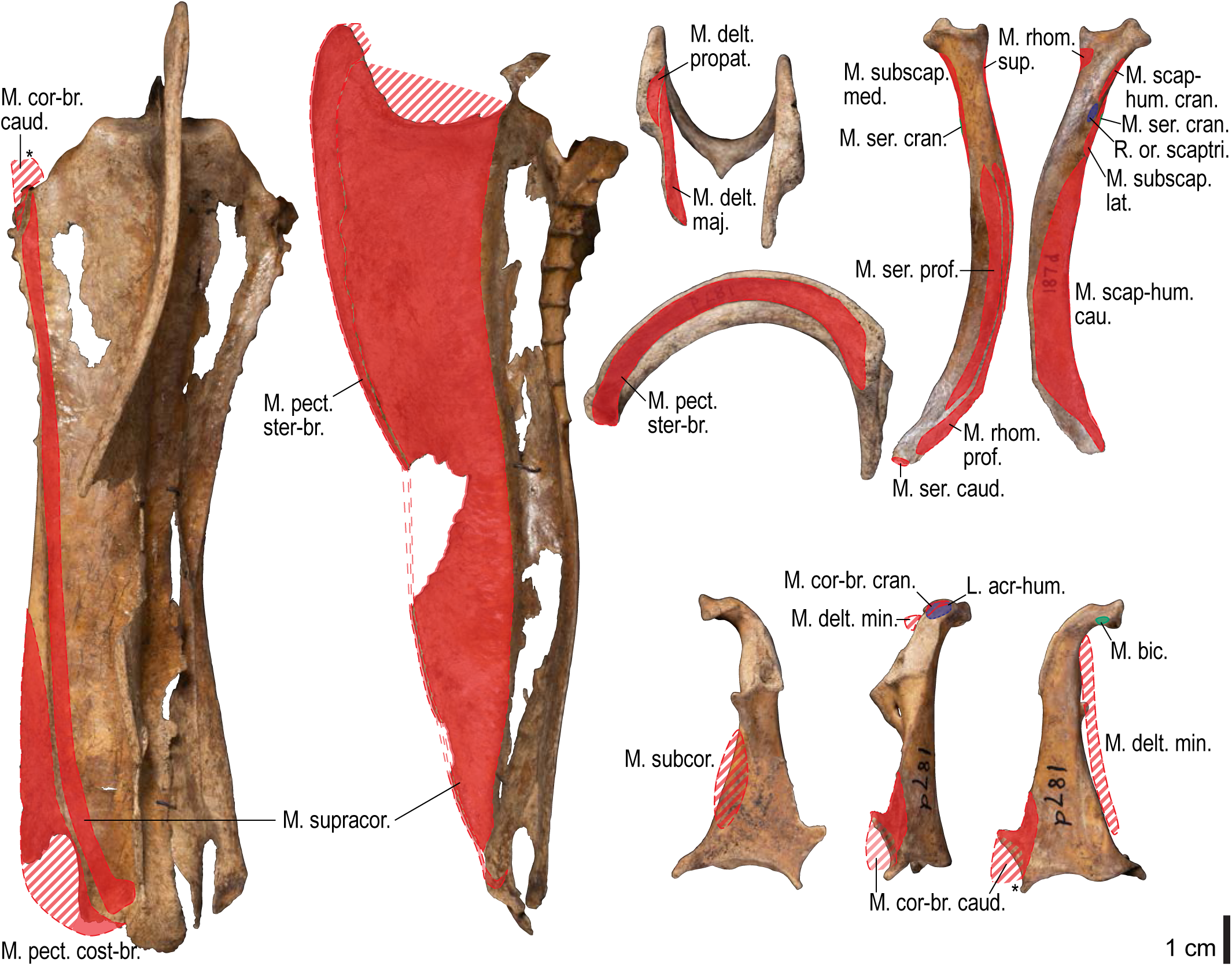
Osteological correlates of major wing muscles and ligaments in *Pinguinus impennis*, pectoral girdle elements. Drawn on UMZC 187.G (sternum) and 187.d (other elements). See Figure 5 for legends.

**Figure 26.**
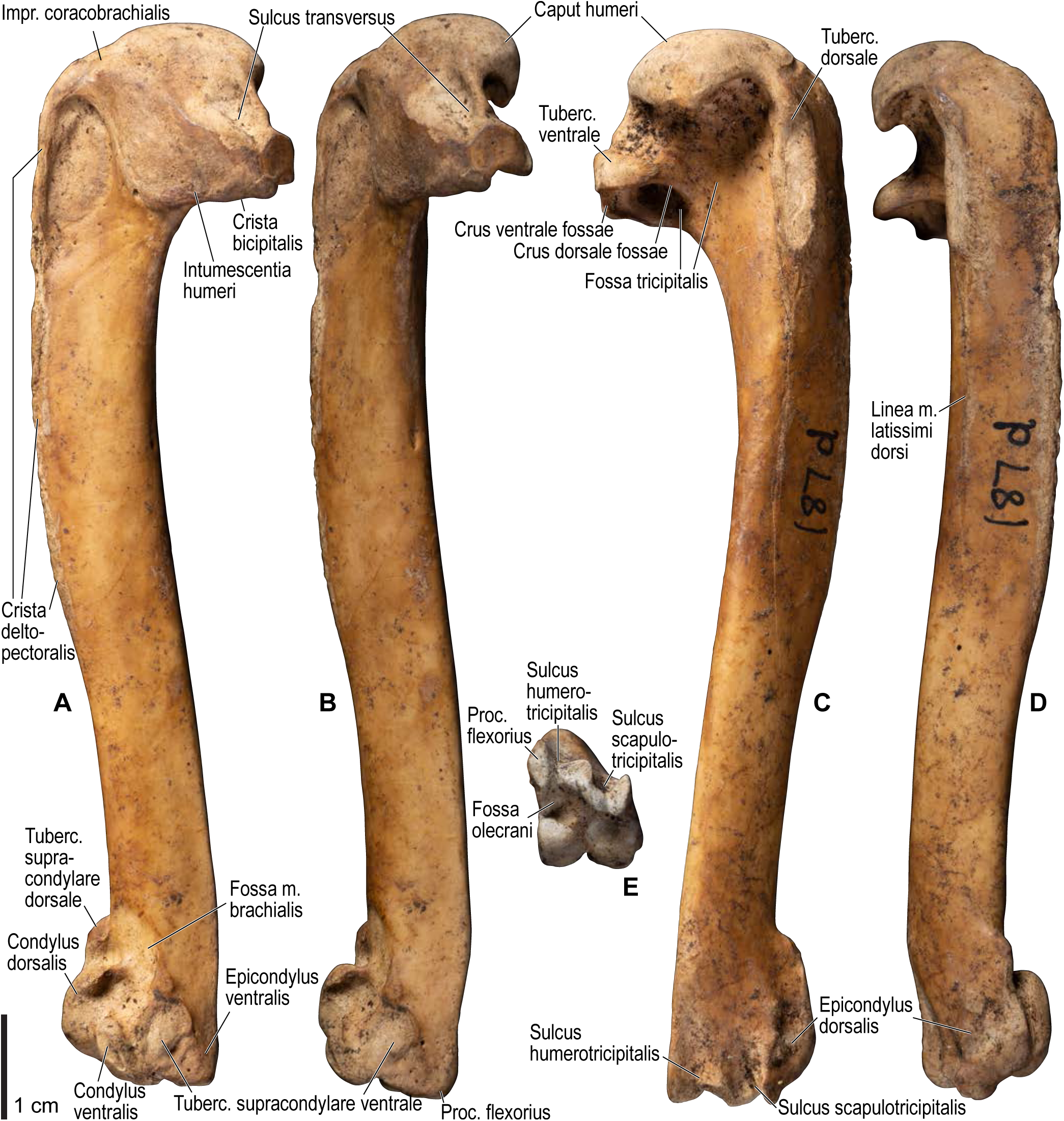
Osteology of *Pinguinus impennis*, humerus. Drawn on UMZC 187.d. Right humerus in cranial (**A**), ventral (**B**), caudal (**C**), dorsal (**D**), and distal (**E**) views. See Figure 4 for abbreviations.

**Figure 27.**
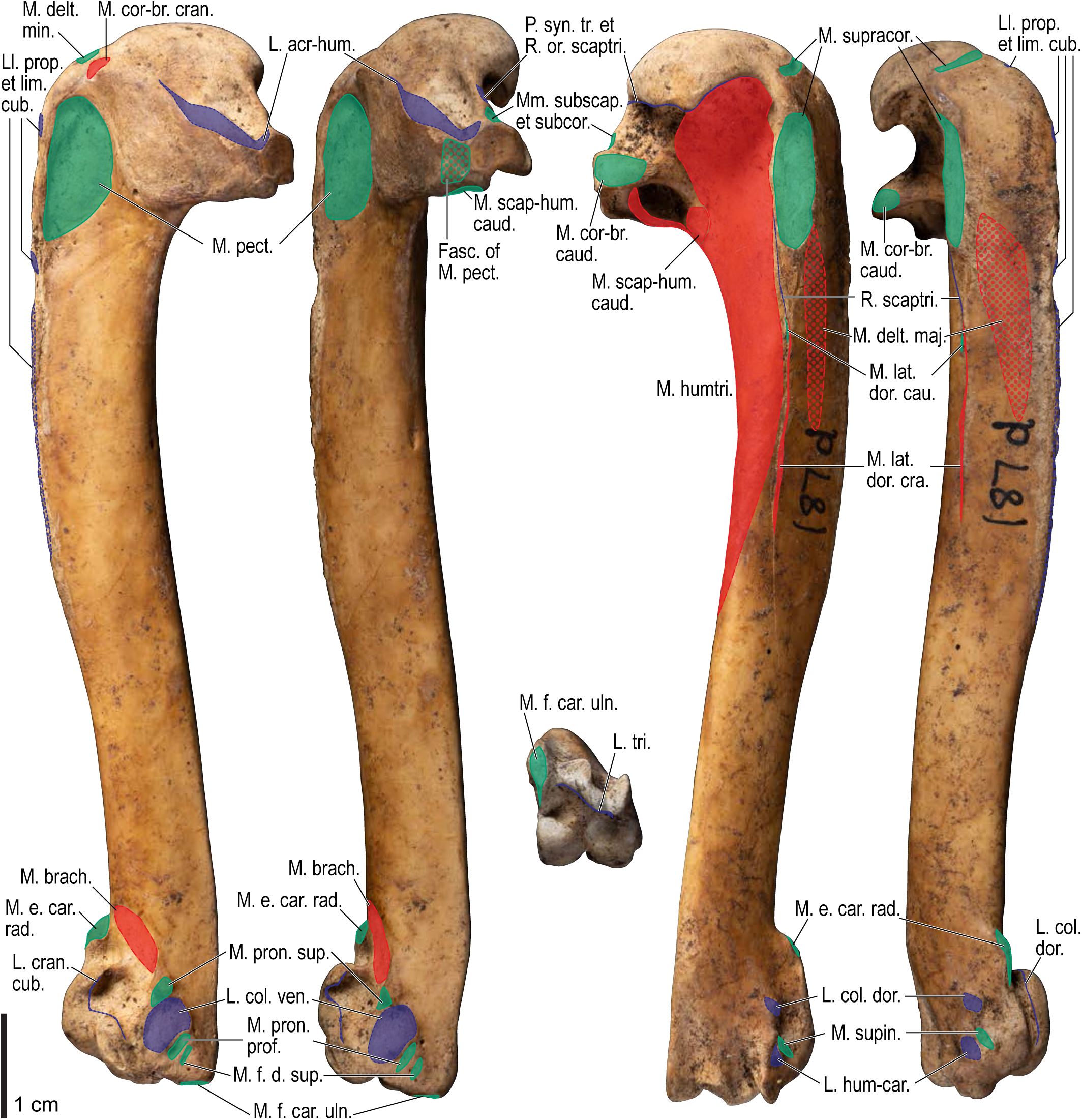
Osteological correlates of major wing muscles and ligaments in *Pinguinus impennis*, humerus. Drawn on UMZC 187.d. Dotted fill represents uncertainty in the extent of attachment sites. See Figure 5 for other legends.

**Figure 28.**
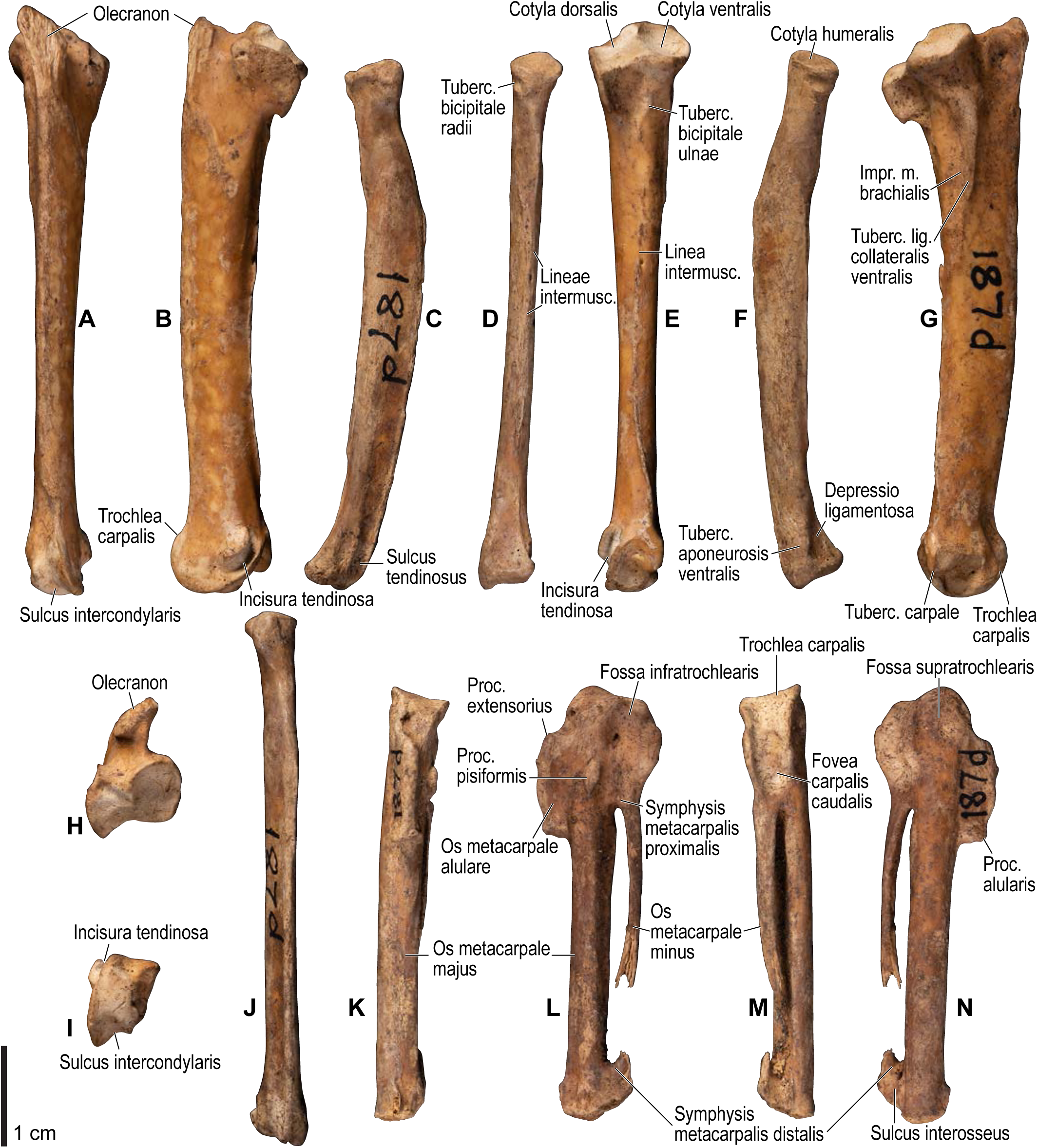
Osteology of *Pinguinus impennis*, distal wing elements. Drawn on UMZC 187.d. Right ulna in caudal (**A**), dorsal (**B**), cranial (**E**), ventral (**G**), proximal (**H**), and distal (**I**) views; right radius in dorsal (**C**), caudal (**D**), ventral (**F**), and cranial (**J**) views; right carpometacarpus in cranial (**K**), ventral (**L**), caudal (**M**), and dorsal (**N**) views. See Figure 4 for abbreviations.

**Figure 29.**
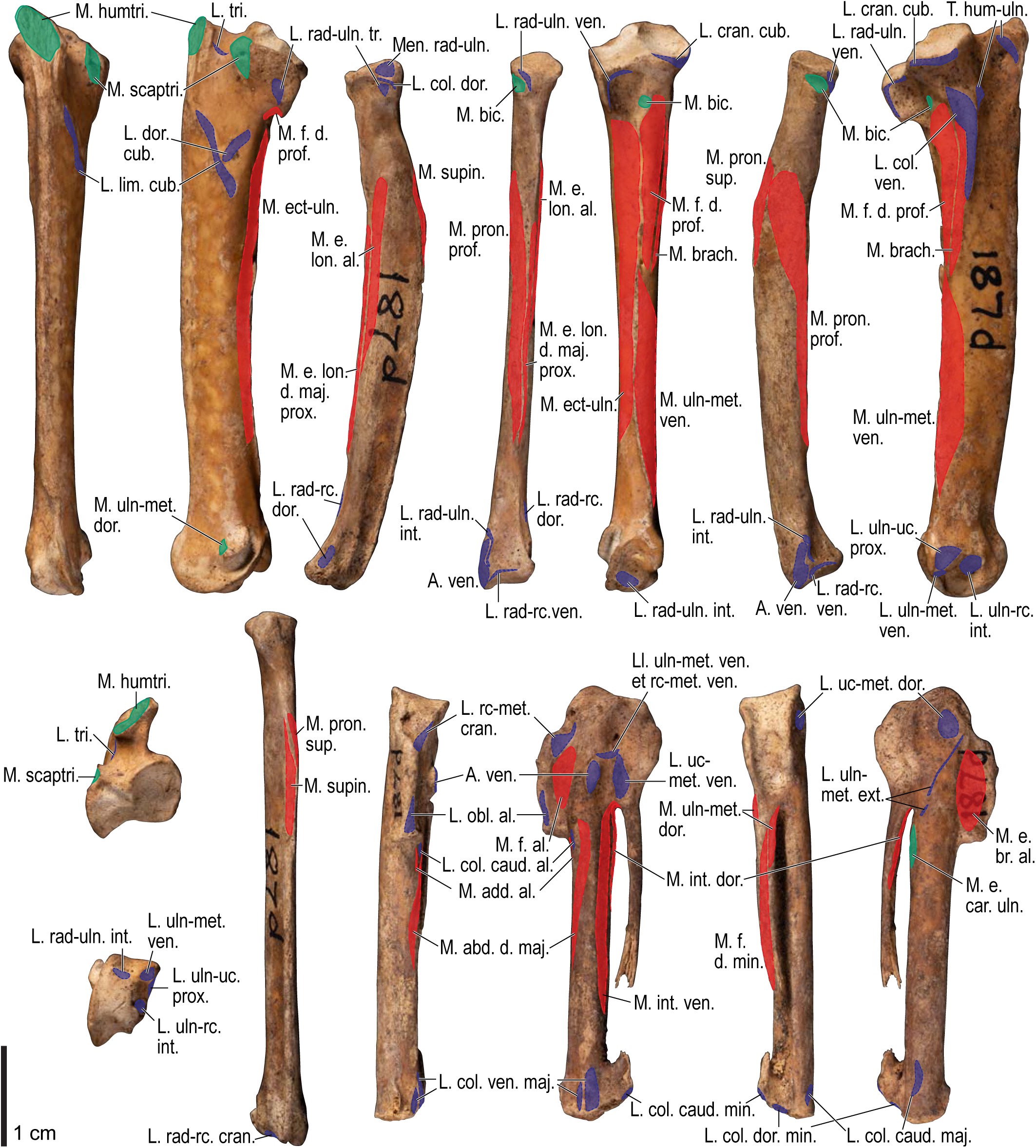
Osteological correlates of major wing muscles and ligaments in *Pinguinus impennis*, distal wing elements. Drawn on UMZC 187.d. See Figure 5 for legends.

**Figure 30.**
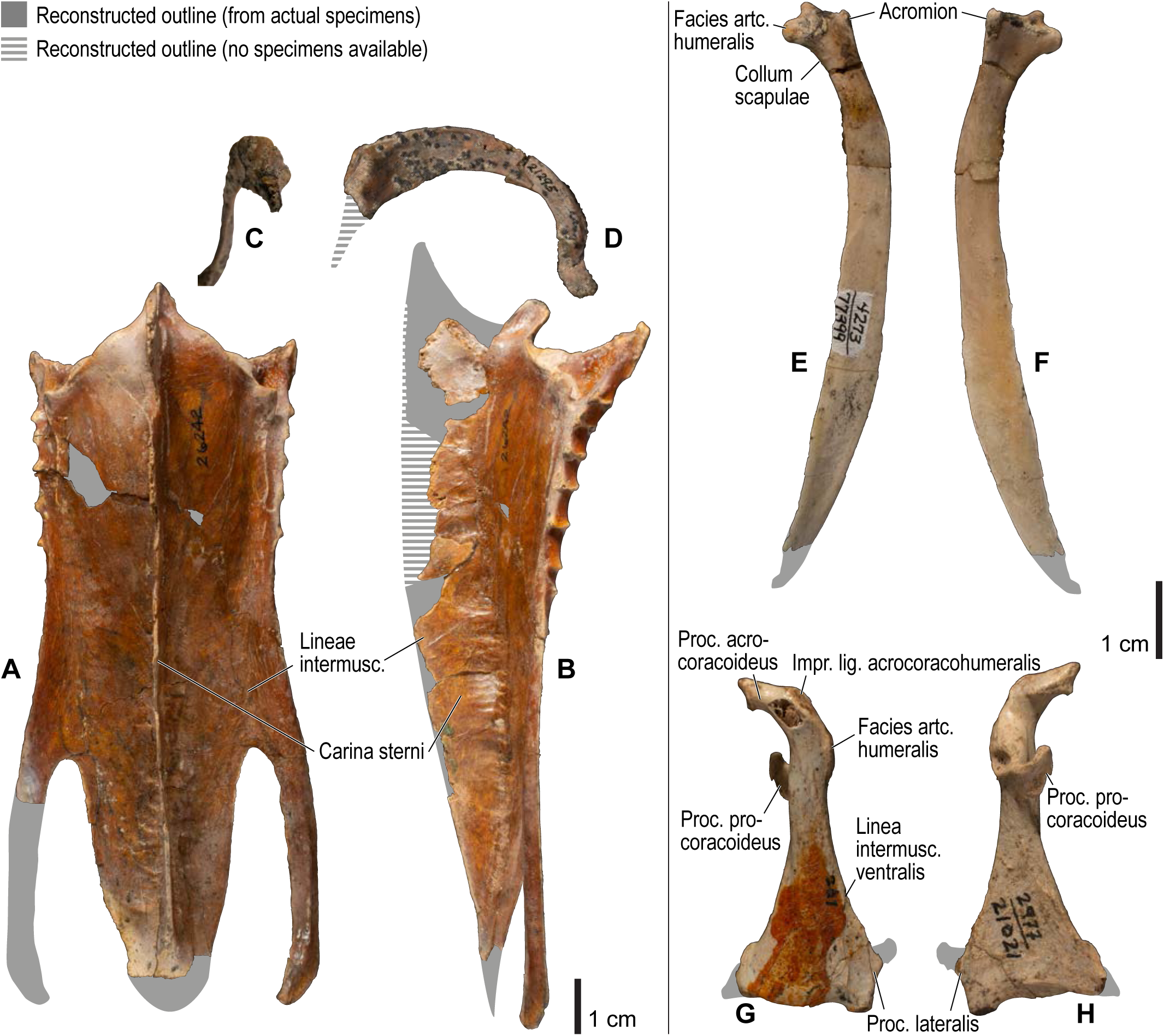
Osteology of *Mancalla*, pectoral girdle elements. Drawn on SDSNH 26242 (sternum), 21295 (furcula), 77399 (scapula), and 21021 (coracoid). Sternum in ventral (**A**) and left lateral (**B**) views; furcula in dorsal (**C**) and right lateral (**D**) views; left scapula in lateral (**E**) and medial (**F**) views; left coracoid in ventral (**G**) and dorsal (**H**) views. Gray shade represents reconstructed outlines. See Figure 4 for abbreviations.

**Figure 31.**
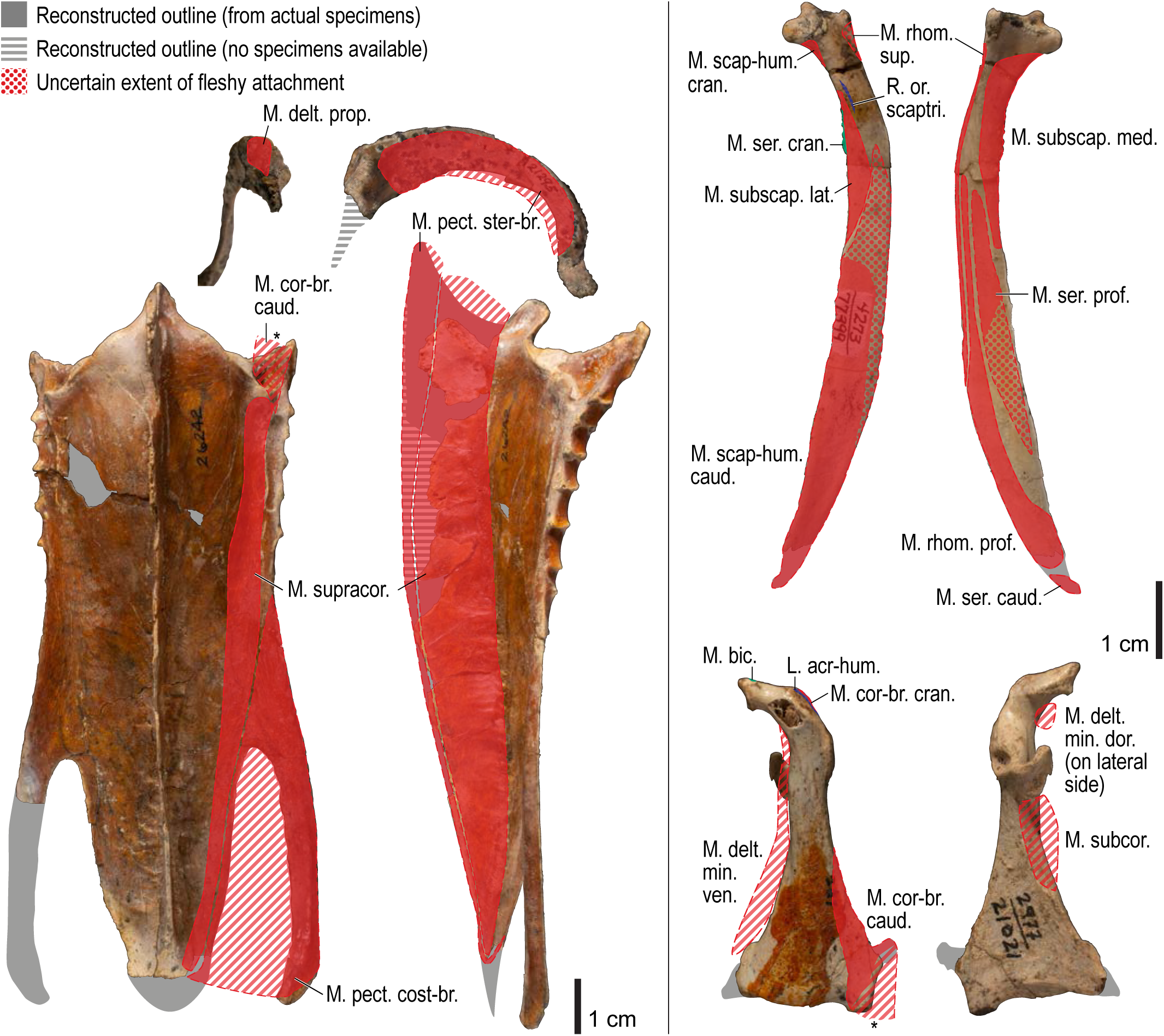
Osteological correlates of major wing muscles and ligaments in *Mancalla*, pectoral girdle elements. Drawn on SDSNH 26242 (sternum), 21295 (furcula), 77399 (scapula), and 21021 (coracoid). See Figures 5 and 30 for legends.

**Figure 32.**
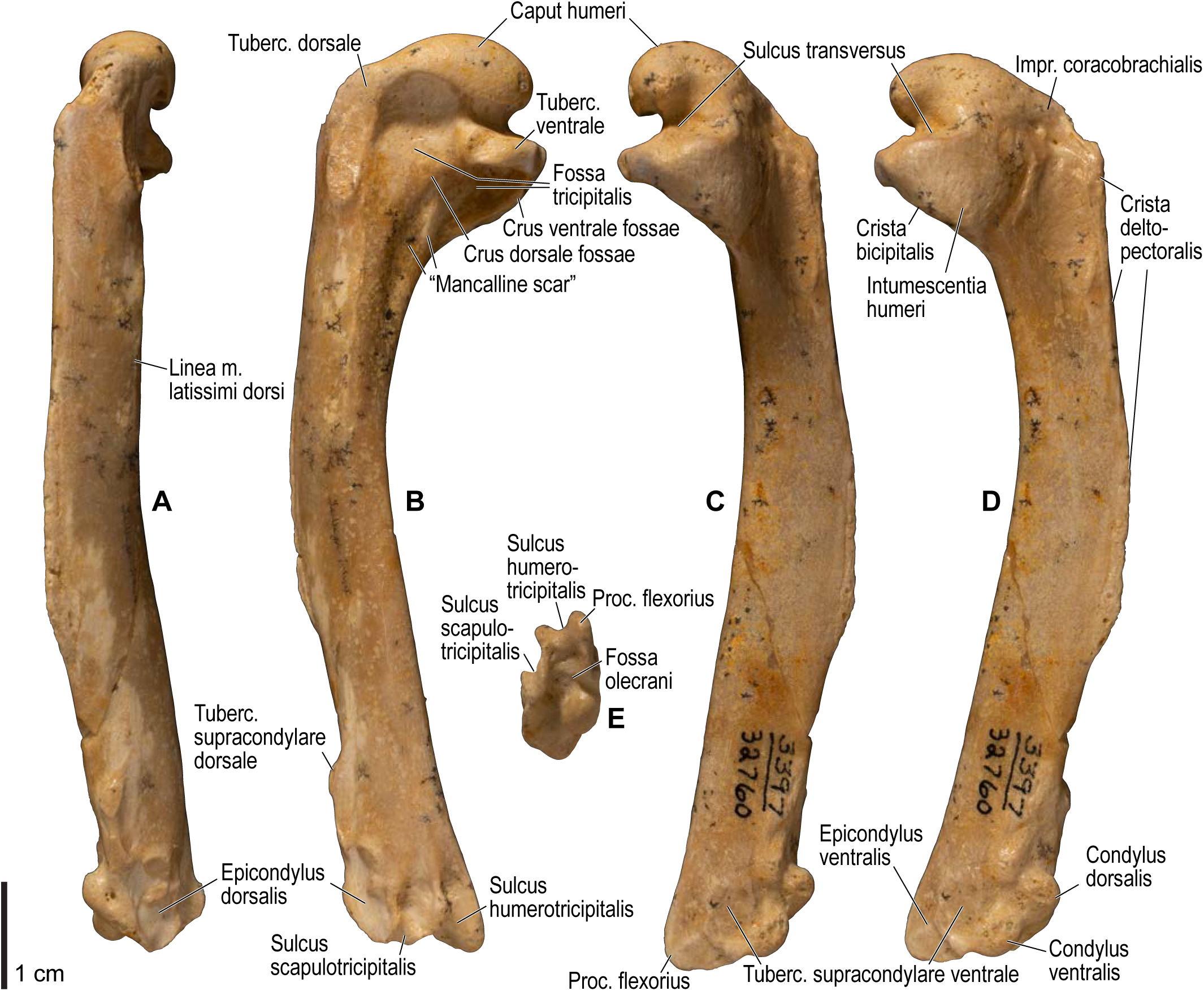
Osteology of *Mancalla*, humerus. Drawn on SDSNH 32760. Left humerus in dorsal (**A**), caudal (**B**), ventral (**C**), cranial (**D**), and distal (**E**) views. See Figure 4 for abbreviations.

**Figure 33.**
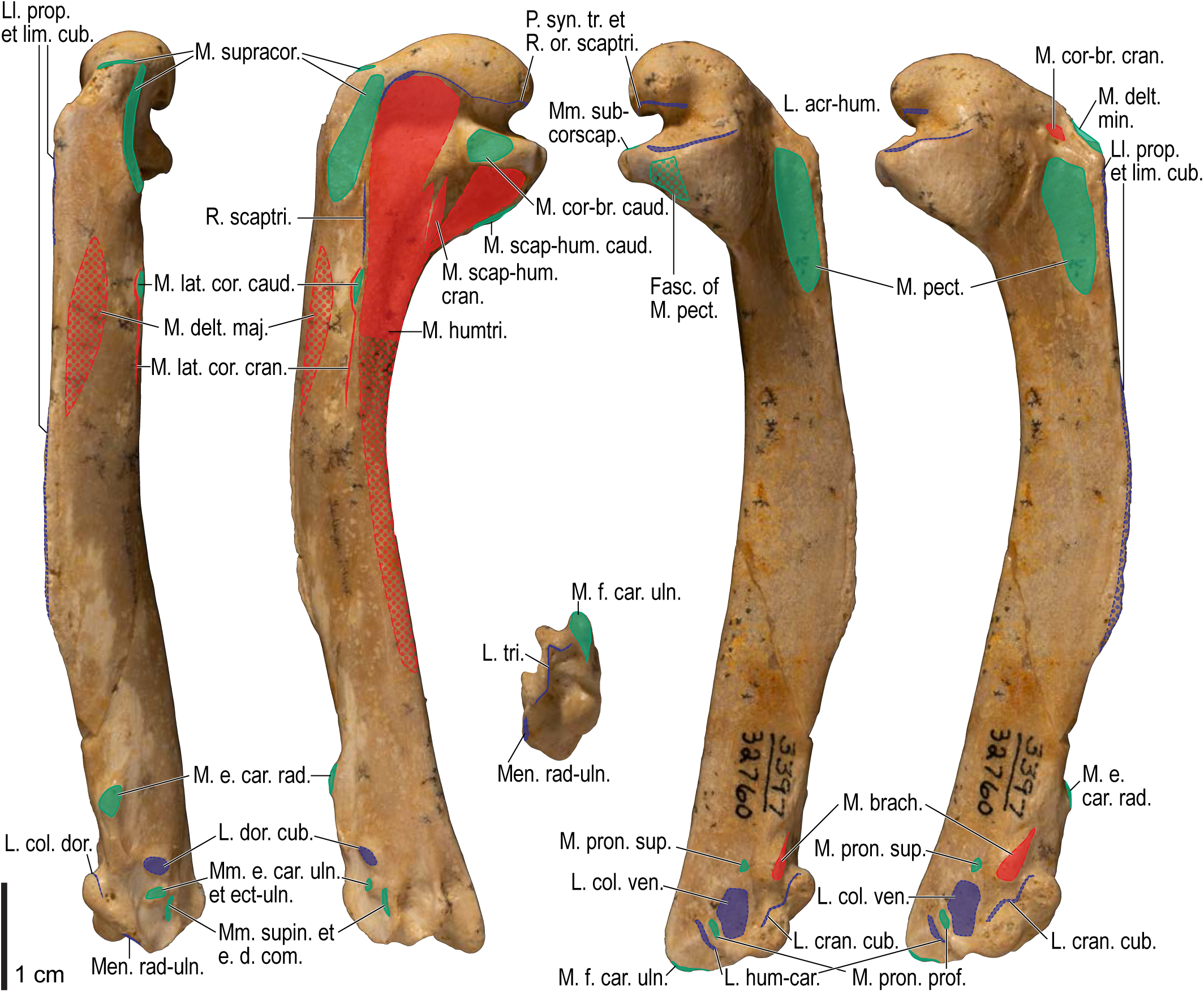
Osteological correlates of major wing muscles and ligaments in *Mancalla*, humerus. Drawn on SDSNH 32760. See Figures 5 and 27 for legends.

**Figure 34.**
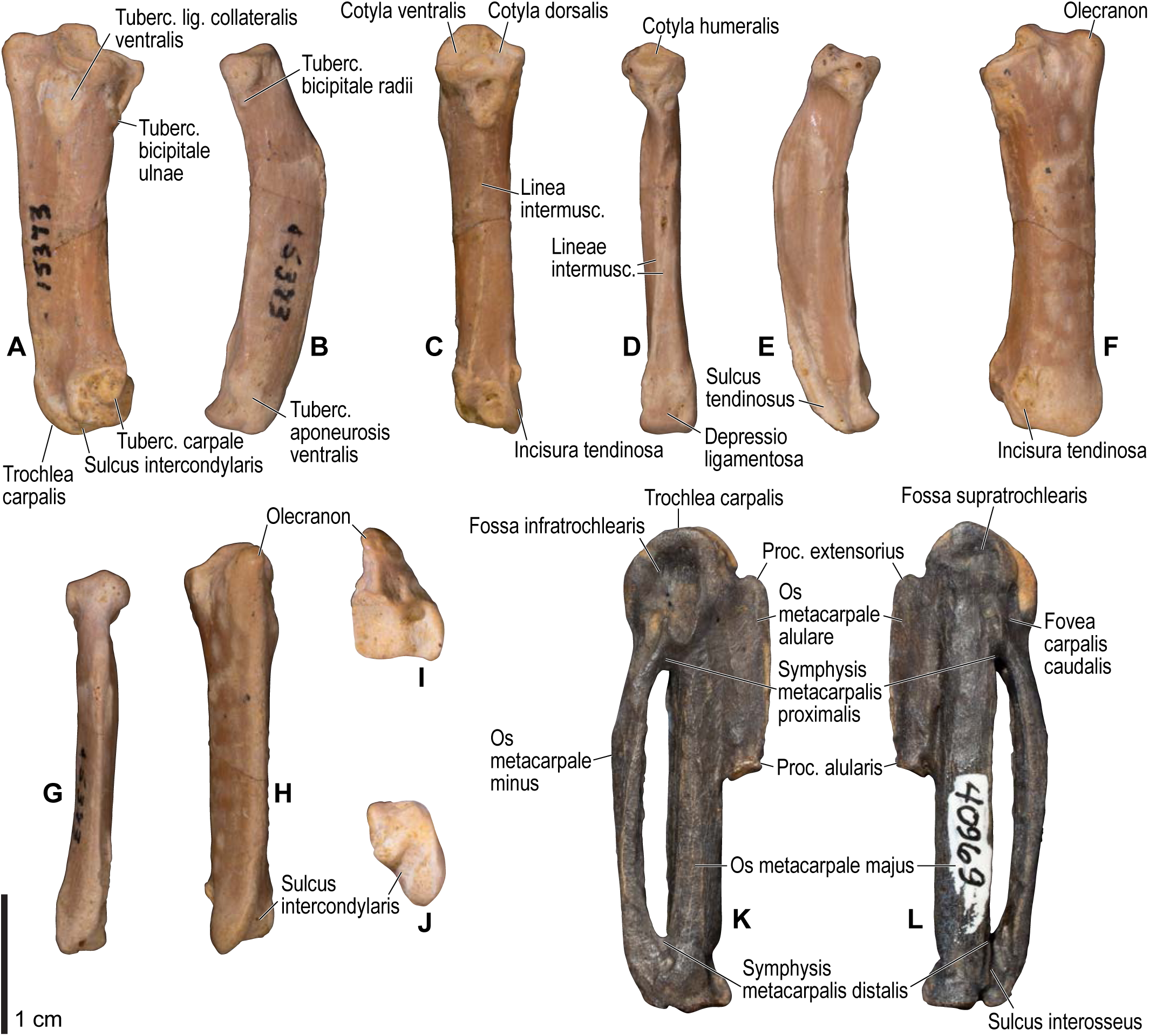
Osteology of *Mancalla*, distal wing elements. Drawn on LACM 15373 (radius and ulna) and SDSNH 40969 (carpometacarpus). Left ulna in ventral (**A**), cranial (**C**), dorsal (**F**), caudal (**H**), proximal (**I**), and distal (**J**) views; left radius in ventral (**B**), caudal (**D**), dorsal (**E**), and cranial (**G**) views; left carpometacarpus in ventral (**K**) and dorsal (**L**) views. Note that the carpometacarpus figured is from a distinct size class (probably a different species) from that of the forearm elements. See Figure 4 for abbreviations.

**Figure 35.**
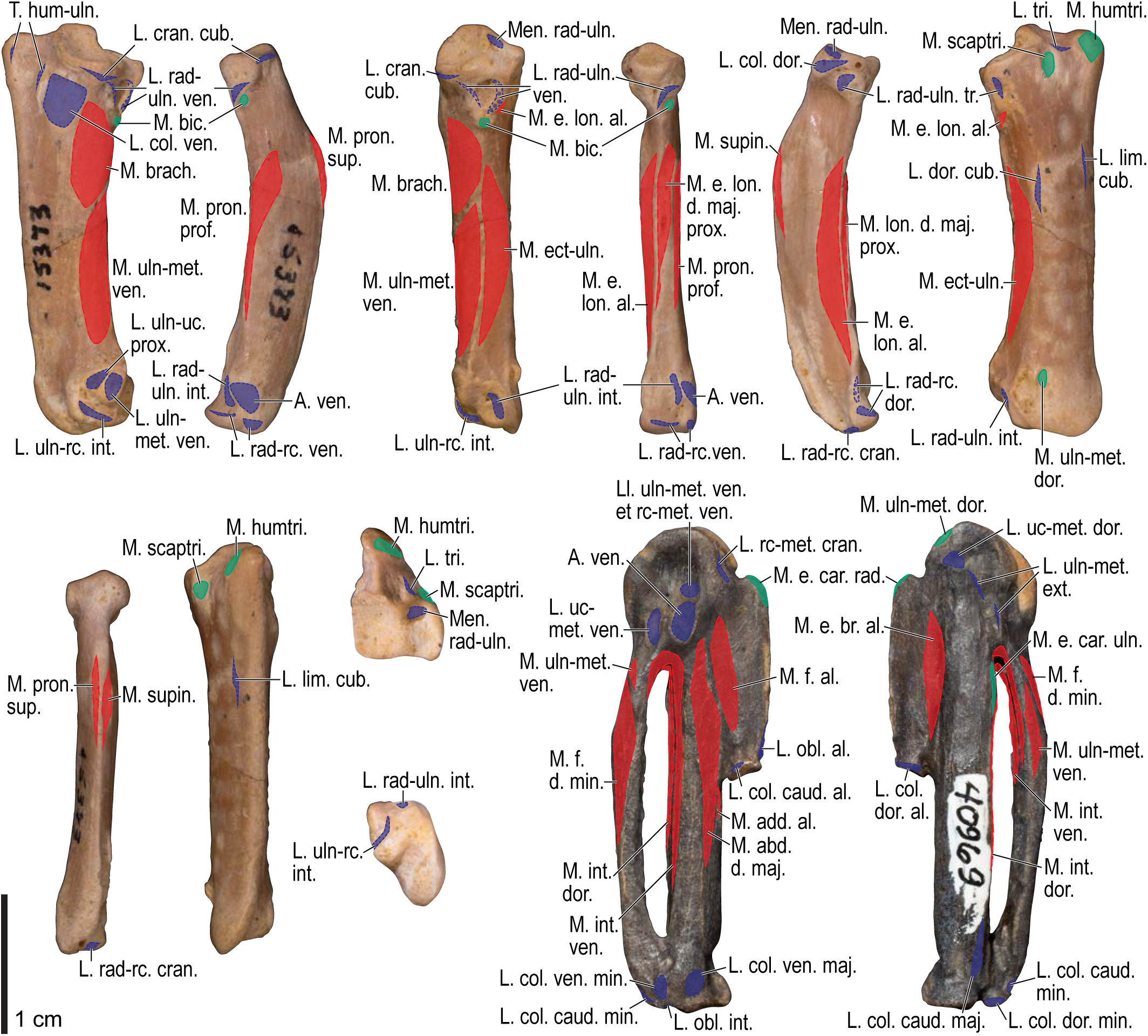
Osteological correlates of major wing muscles and ligaments in *Mancalla*, distal wing elements. Drawn on LACM 15373 (radius and ulna) and SDSNH 40969 (carpometacarpus). See Figure 5 for legends.

**Figure 36.**
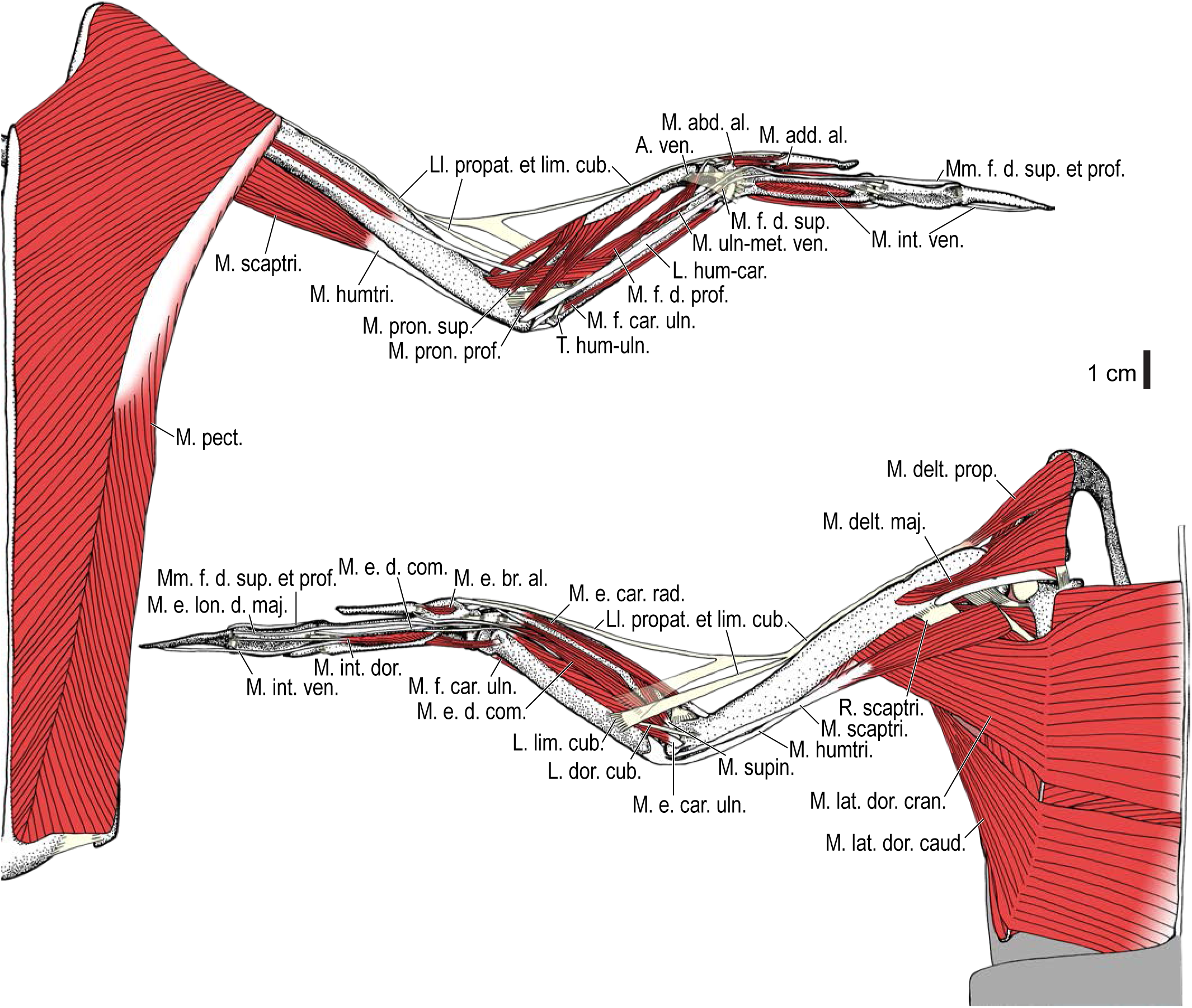
Reconstructed wing musculature in *Pinguinus impennis*, ventral (top) and dorsal (bottom) views, superficial layer. The reconstruction is based on a composite of elements, thus some proportions may be inaccurate. This illustration is partly schematic, and is not an accurate representation of muscle volume, pennation, or other architectural properties. See Table 3 for abbreviations.

**Figure 37.**
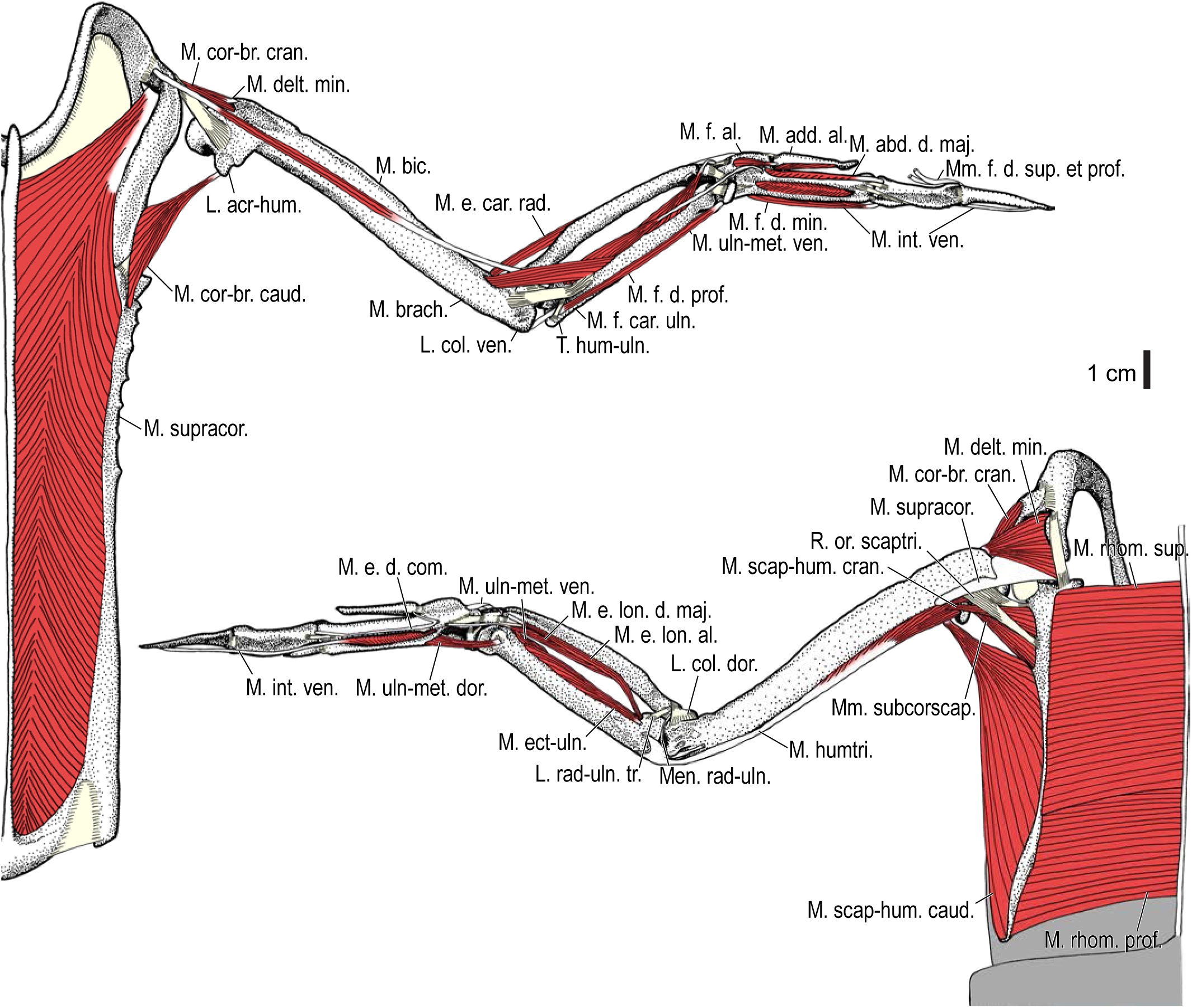
Reconstructed wing musculature in *Pinguinus impennis*, ventral (top) and dorsal (bottom) views, deep layer. See Table 3 for abbreviations and Figure 36 for further information.

**Figure 38.**
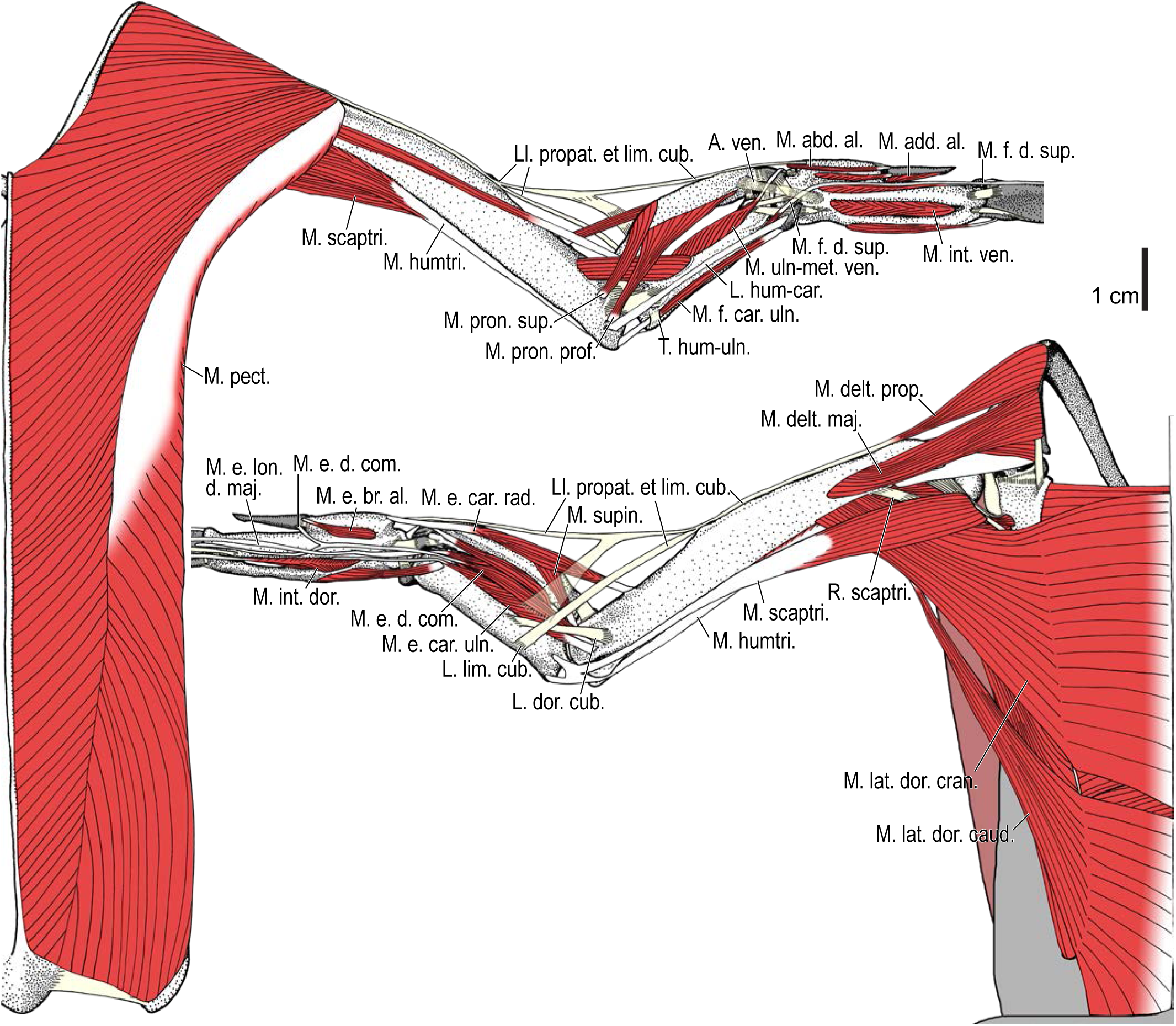
Reconstructed wing musculature in *Mancalla*, ventral (top) and dorsal (bottom) views, superficial layer. The reconstruction is based on a composite of elements, with proportions roughly scaled to the holotype of *Mancalla cedrosensis* (LACM 15373). The omal end of the furcula, free carpal bones and phalanges are not known for this taxon (silhouettes shown in gray). This illustration is partly schematic, and is not an accurate representation of muscle volume, pennation, or other architectural properties. See Table 3 for abbreviations.

**Figure 39.**
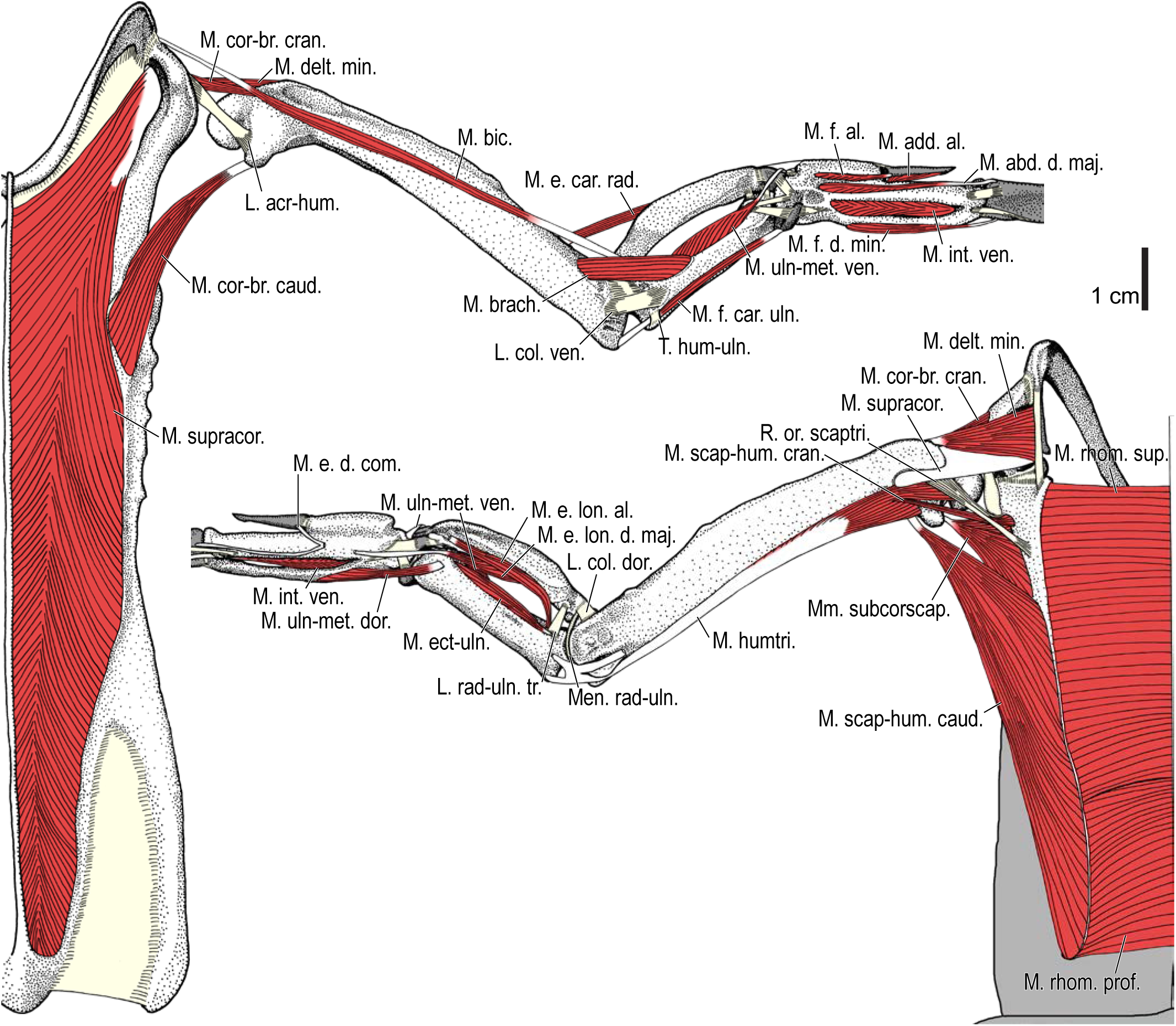
Reconstructed wing musculature in *Mancalla*, ventral (top) and dorsal (bottom) views, deep layer. See Table 3 for abbreviations and Figure 38 for further information.

There is some uncertainty regarding the nature of the attachment sites for the m. brachialis in *Mancalla*. In the extant taxa examined (except *Spheniscus*), the origin and insertion of this muscle, both being fleshy, are marked by fossae (the fossa m. brachialis of the humerus and the impressio m. brachialis of the ulna). However, in *Mancalla*, the corresponding areas are marked by a broad tubercle on the humerus and a raised, smooth platform on the ulna (Figs. 32–35). It is therefore tempting to speculate that the muscle did not retain fleshy attachments in *Mancalla*, because fleshy attachments typically do not exhibit these sorts of osteological correlates in the taxa examined. Nevertheless, it will be impossible to definitively evaluate this idea without further analyses (e.g., histological assessment).

For *Pinguinus*, the reconstructed musculature was subsequently compared with a dried skeletal specimen in which remnants of the elbow and forearm musculature are preserved (NHMUK 1972.1.156). The relative positions of the muscles and ligaments reconstructed from osteological correlates were consistent with those preserved in this specimen (Fig. 40), partially confirming the validity of the present reconstruction.

**Figure 40.**
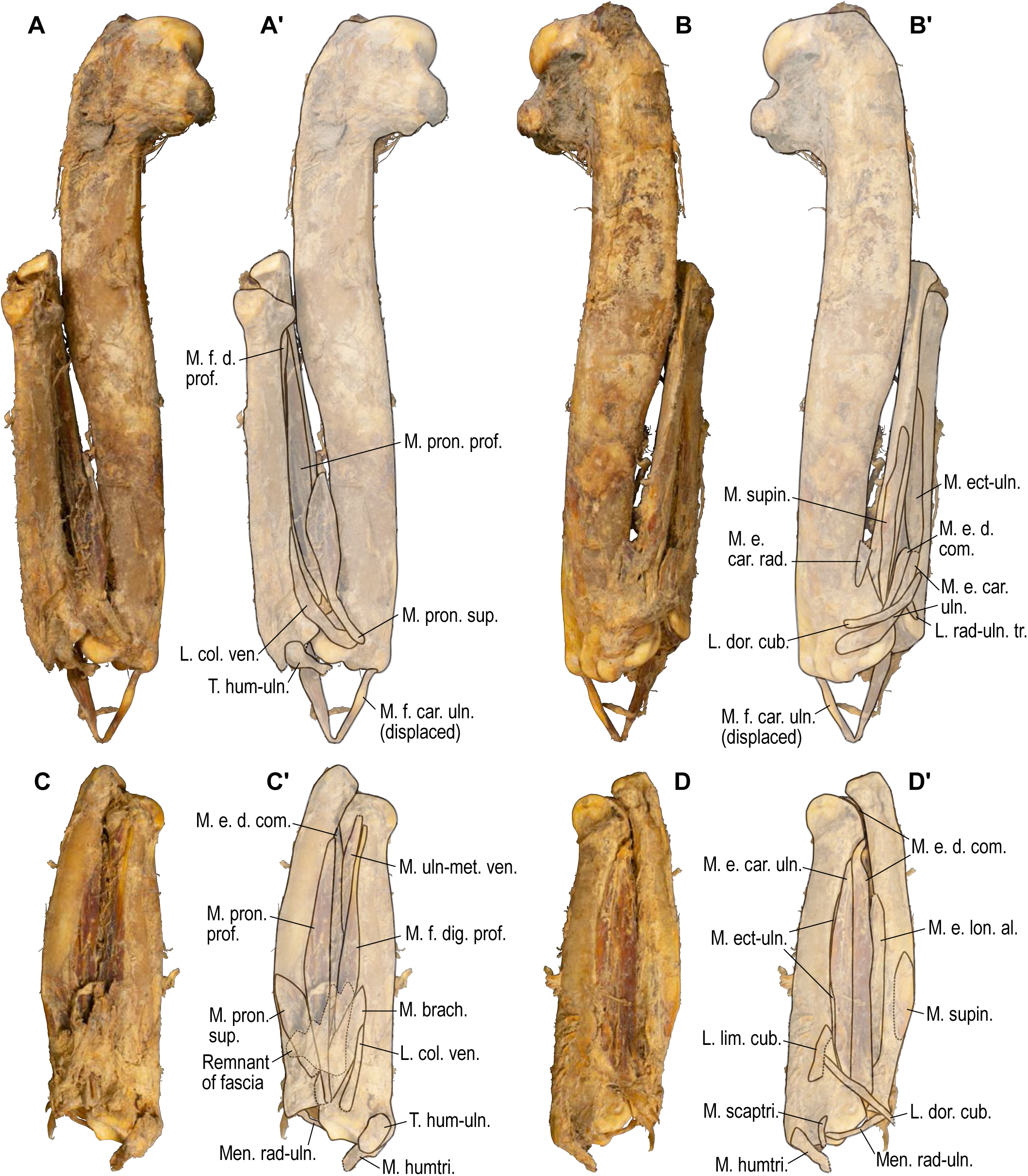
Dried skeletal specimen of *Pinguinus impennis* with remnants of soft tissues (NHMUK S/1972.1.156), photographs (**A**–**D**) and interpretative drawings (**A’**–**D’**). **A**, **B**, right humerus and forearm in original articulation, ventral and dorsal views, respectively. **C**, **D**, left forearm in original articulation, ventral and dorsal views, respectively. Dotted outlines represent damaged parts. See Table 3 for abbreviations.

## Discussion

The extant charadriiform taxa used as the basis for reconstructing the musculature of the extinct flightless auks showed little variation in the positions of osteological correlates of major wing muscles and ligaments. This relative stability largely validates the reconstruction of musculature based on the Extant Phylogenetic Bracket, as ancestral state reconstruction based on parsimony tends to be more accurate for characters with low evolutionary lability (unless transition rates are biased toward derived states; Frumhoff and Reeve 1994; Schultz et al. 1996; Cunningham 1999). Of course, considerable uncertainty accompanies any inferences concerning structures lacking osteological correlates (Witmer 1995) and the identification of the extent of fleshy attachment sites from bones alone (Bryant and Seymour 1990). Given the unavailability of the radiale, ulnare, and phalanges of Mancallinae, there is simply no way to examine any potential specializations of these elements and their associated soft parts. Furthermore, little definitive information can be gleaned regarding the range and limits of joint motion in the extinct taxa examined; this is something to be explored in the future based on the present results. These limitations should be kept in mind when interpreting the results of the present reconstructions.

Some clarification regarding the habits of the extinct taxa investigated here is required as a basis for further discussion. Most previous authors have assumed that Mancallinae were flightless wing-propelled divers, possibly convergent with penguins (e.g., Storer 1960; Livezey 1988). However, both flightlessness and wing-propelled diving in Mancallinae are hypotheses to be evaluated from external evidence, if we are to critically assess the hypothesis of convergent evolution between flightless wing-propelled divers. Because Mancallinae lies outside crown-group Alcidae (Smith 2011) and no extant charadriiforms other than Alcidae are wing-propelled divers, the phylogenetic placement of Mancallinae alone does not provide decisive evidence for wing-propelled diving, let alone flightlessness. Nonetheless, the inference of flightlessness in Mancallinae is relatively straightforward; the relative size of mancalline wing skeletal elements is extremely small—smaller even than those of *Pinguinus*, whose estimated wing loading was clearly too large to allow for aerial flight (Livezey 1988). Hence, it is almost certain that mancalline auks were incapable of aerial flight. Evidence for wing-propelled diving in Mancallinae is less obvious, but can be inferred from the dorsoventrally flattened and craniocaudally widened shafts of wing bones, especially the humerus (Livezey 1988; Smith and Clarke 2014). Torsion of the shaft, rather than longitudinal bending, is a critical factor in the mechanical design of the humerus for aerial flapping flight in birds (Biewener and Dial 1995), and, for a given amount of material, resistance to torsion is optimized by a tube with a circular cross-section (Vogel 1992; Daegling 2002; Ennos 2012). As such, deviation of the wing bone shafts from a circular cross-section is most likely indicative of specialization for a function other than aerial flapping flight. Dorsoventrally flattened limb bones are seen in the wings of various wing-propelled diving birds (penguins, volant auks, diving petrels and some shearwaters; Kuroda 1954; Storer 1960; Livezey 1988, 1989), and are prevalent in the specialized limbs of many different groups of secondarily aquatic tetrapods (e.g., Thewissen and Taylor 2007; Kelley and Pyenson 2015). When limbs are employed as flapping hydrofoils, dorsoventrally flattened limb bones increase hydrofoil rigidity, enabling the production of increased thrust and improving propulsive efficiency (although hydrofoil flexibility is another crucial factor underlying propulsive efficiency; DeBlois and Motani 2019). Therefore, flattened wing bones in Mancallinae strongly indicate that they were structurally suited to aquatic flight in a manner presumably homologous with that in crown-group Alcidae, and analogous to that in other extant wing-propelled diving birds (as well as other aquatic tetrapods).

### Functional anatomy of flightless auks

With the anatomical data presented here, it is possible to evaluate numerous remarkable features of the wings of extinct flightless auks. Arguably, the most distinctive features of the humerus in Mancallinae are muscle scars within the ventral portion of the fossa tricipitalis (the so-called “mancalline scar”; Fig. 32; Miller and Howard 1949; Smith 2011). This structure comprises an area of raised relief near the base of the crus dorsale fossae and a proximodistally elongated depression lying dorsodistal to this area. It should be noted that this interpretation slightly differs from that of Smith (2011), who stated that some mancalline taxa are characterized by either the depression or the area of raised relief. Here, the present observations confirm the presence of both the depression and the area of raised relief in all specimens examined. Previous authors were inconclusive regarding the identities of the muscle(s) corresponding to these scars, although a few candidates had been proposed (Miller and Howard 1949, Smith 2011). One point that has apparently evaded previous authors’ observations is that a similar area of depression and raised relief is actually present in some extant alcids (e.g., *Synthliboramphus*, *Uria*, and *Alca*; Fig. 41), although these structures are much less distinct than in Mancallinae and there is apparently some intraspecific variation in the development of these structures. To be more specific, the area of raised relief was found in almost all extant taxa comprising the extant phylogenetic bracket for *Mancalla* (i.e., *Larus*, *Catharacta*, and most individuals of extant alcids; Fig. 41B), and in these extant taxa hosts the insertion of the m. scapulohumeralis cranialis. Dorsodistally adjacent to this insertion is part of the origin of the m. humerotriceps; this area is marked by a depression in some alcids (*Synthliboramphus*, *Uria*, and *Alca*; Fig. 41A), whereas the corresponding area falls within the much deeper and broader fossa tricipitalis in other extant alcids (*Cerorhinca*, *Fratercula*, and *Cepphus*) as well as the closest outgroups to Pan-Alcidae (*Larus* and *Catharacta*). The attachment sites of these two muscles are always adjacent to one another in the taxa mentioned. Therefore, it can be safely inferred (Level I inference) that the “mancalline scar” hosts the same combination of muscles, namely the m. scapulohumeralis cranialis on the area of raised relief and the m. humerotriceps in the depression, although the presence of the depression itself might not necessarily be homologous between *Mancalla* and crown-group alcids, as the presence of the depression optimizes equivocally in the most recent common ancestor of Mancallinae and crown-group Alcidae. This conclusion is consistent with Miller and Howard’s (1949) speculation that the scar hosts the “supraspinatus muscle” (= m. scapulohumeralis cranialis in, e.g., Shufeldt’s [1890] terminology).

**Figure 41.**
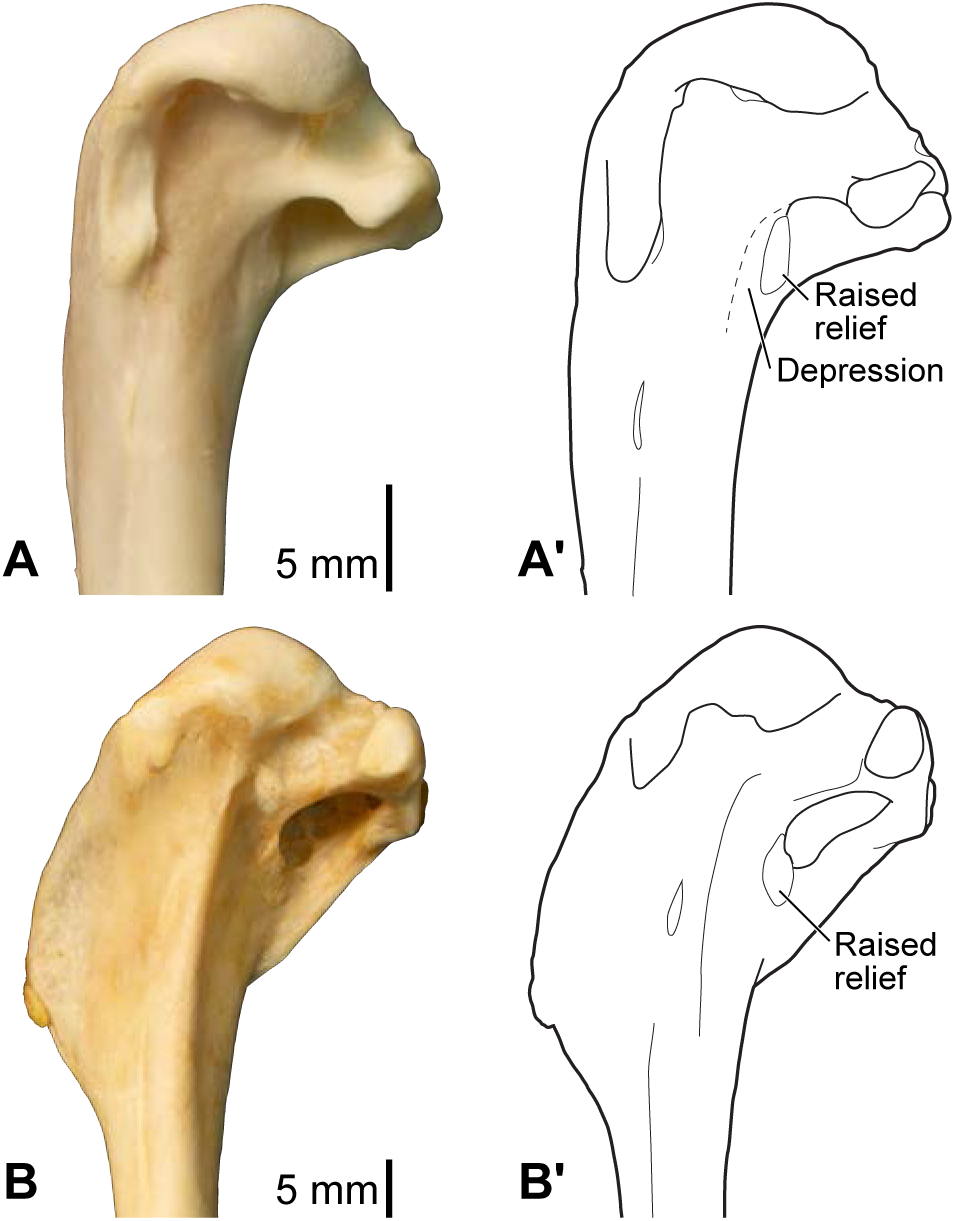
Equivalent of “mancalline muscle scar” in extant charadriiform taxa. **A**, *Alca torda*, NHMUK S/1977.65.7. **B**, *Catharacta antarctica*, NHMUK 1998.63.1. Proximal end of left humeri in caudal view, photographs (**A**, **B**) and interpretative drawings (**A’**, **B’**).

The mechanical advantage of many extrinsic wing muscles appears to have been comparatively greater in *Pinguinus* and *Mancalla* than in extant volant auks. This is partly the product of a reduction in the length of the wing skeleton (especially of distal elements; Livezey 1988; Smith 2011), which would reduce the length of the out-lever in movements of the entire wing. Additionally, distal shifts in the relative positions of the insertions of some extrinsic muscles (and sometimes the elongations of these insertions) would have increased the length of these muscles’ in-levers. Notably, such shifts are most evident in elevators and retractors of the wings (muscle functions following Raikow 1985): the m. supracoracoideus on the distally elongated tuberculum dorsale, the m. scapulohumeralis cranialis on an enlarged scar in the fossa tricipitalis (especially in *Mancalla*; above), and the m. scapulohumeralis caudalis on the distally extended crista bicipitalis (Figs. 26, 27, 32, 33). The increased mechanical advantage of these wing elevators and retractors would have increased the torque generated by these muscles, at the expense of terminal speed of the movement per unit length of muscle fiber contraction under no counteracting force (e.g., Ennos 2012). This modification would have enabled a powerful upstroke of the wings against resistance and drag in the water, facilitating the production of forward thrust through the upstroke phase typical of aquatic flight (see Introduction), and thereby enabling more efficient aquatic locomotion than that achievable by extant volant auks. Nevertheless, in the absence of quantitative data on muscle moment arms and joint range of motion, definitive conclusions regarding the mechanical performance of the wing of these extinct auks might be premature at present. The present results provide a basis for the future testing of such hypotheses through rigorous mechanical investigations of the musculoskeletal system during aquatic flight.

One of the most prominent features of the humerus observed in both *Pinguinus* and Mancallinae is the greatly elongated crista deltopectoralis, associated with the lack of sigmoid curvature of the humeral shaft (e.g., Lucas 1901; Smith 2011). There are several major soft parts associated with the crista deltopectoralis: the m. pectoralis inserts on its ventral surface, the m. deltoideus pars major inserts on its dorsal surface, the m. deltoideus pars minor inserts on its proximal margin, and the ligg. propatagiale et limitans cubiti (often as a common ligament) are anchored on its cranial margin. Of these, the attachment sites of the former three muscles are always restricted in the extant species examined (Figs. 7, 12), and are not particularly enlarged in *Pinguinus* and *Mancalla* (although the attachment site of the m. deltoideus pars major is challenging to delineate). By contrast, the attachment site of the ligg. propatagiale et limitans cubiti is elongated along with the crista deltopectoralis, as clearly indicated by the rugosity of the entire cranial margin of the crest (Figs. 26, 32). Thus, if any muscles or ligaments are causally implicated in the elongation of the crest, the common ligament of the ligg. propatagiale et limitans cubiti would seem to be the most likely candidate (Figs. 27, 33).

The propatagium provides the wings with additional surface area (e.g., Raikow 1985), and the ligg. propatagiale et limitans cubiti support its leading edge and central part, respectively. The distal shift of the anchors of these ligaments in flightless auks indicates that the overall area of the propatagium was substantially reduced in these taxa (Figs. 36, 38). Such a modification is understandable given that the wings of flightless auks would only have been used in aquatic flight. Unlike in aerial flight, additional wing area provided by a large propatagium would not be beneficial, and could even be detrimental, as water poses substantially more drag and resistance to moving wings than air, and thrust production in water is far more important than lift production (Rayner 1988). This expectation is supported by behavioral observations in extant volant auks, where the wings are seen to be kept in a partly folded position during aquatic flight, yielding smaller effective wing area than in the fully extended position (Stettenheim 1959; Spring 1971; Rayner 1995; Gaston and Jones 1998; Kikuchi et al. 2015). As such, both the elongated crista deltopectoralis and attachment site for the ligg. propatagiale et limitans cubiti appear to represent modifications to increase the rigidity of the leading edge of the wings, which would increase thrust production in aquatic flight (DeBlois and Motani 2019). It is evident that these modifications were acquired independently in *Pinguinus* and Mancallinae (Smith 2011).

In *Mancalla*, it is probable that function was lost in the m. flexor digitorum profundus—one of the intrinsic muscles of the wing. The origin of this muscle typically lies on the proximal ulna between the impressio m. brachialis and the linea intermuscularis and, in non-alcid charadriiforms, on the narrow space ventrocaudal to the impressio m. brachialis (see above). In *Mancalla*, however, no substantial spaces are evident in those areas, partly as a result of the enlargement of the impressio m. brachialis (see above; Figs. 34, 35), indicating the virtual loss, or substantial reduction, of the origin of this muscle. In addition, the processus pisiformis of the carpometacarpus, which acts as a pulley for this muscle in the wrist joint (above), is lacking in *Mancalla* (Fig. 34; see also Smith 2011), providing additional evidence for the virtual, if not complete, absence of function in this muscle. These observations may suggest that the function of this muscle—flexion of the wrist and extension and depression of the major digit (Raikow 1985)—was unimportant in *Mancalla*. Correspondingly, the pulley may have been unimportant if the wrist joint was fixed in a straight position. In any case, this probable loss of function would have been associated with reduced mobility of the wrist and digital joints typical of flightless wing-propelled divers (see below), as postulated by Smith (2011). Unfortunately, no wing phalanges are yet known from *Mancalla*, thus conclusions about the insertion of this muscle cannot yet be drawn. Problematically, however, the insertion of this muscle is often indistinguishable from that of the m. flexor digitorum superficialis (see above). In light of this uncertainty concerning its development, the m. flexor digitorum profundus is not illustrated in Figures 38–39. Intriguingly, the processus pisiformis is present in *Miomancalla*, a geologically older, and presumably less specialized member of Mancallinae (Howard 1966, 1976; Smith 2011). The wing elements of *Miomancalla* are apparently less well represented in the fossil record than those of *Mancalla*, thus further fossil discoveries and subsequent anatomical investigation will be needed to shed further light on the reduction of the m. flexor digitorum profundus in Mancallinae.

Lastly, the wings of *Pinguinus* and *Mancalla* are characterized by prominent, well-developed ligaments bracing the elbow joint. Typically, the avian elbow joint is supported by several ligaments: the lig. collaterale ventrale on the ventral aspect of the humero-ulnar articulation, the lig. craniale cubiti on the cranial aspect of the humero-radio-ulnar articulations, and the lig. collaterale dorsale on the dorsal aspect of the humero-radial articulation. In Charadriiformes, but apparently not in other birds, the lig. dorsale cubiti is present on the dorsal aspect of the humero-ulnar articulation, supposedly providing additional support for the elbow joint (see also Stettenheim 1959; below). Among these ligaments, the insertions of the lig. collaterale ventrale and lig. dorsale cubiti are positioned rather distally on the ulna in the flightless auks compared with the volant auks and other charadriiforms examined (Fig. 42). When these ligaments are tensed during extension or dorsoventral bending of the elbow joint, this conformation in flightless auks would confer a greater mechanical advantage of forces exerted by the ligaments, and thereby increase efficiency of the wings under water, at the expense of mobility at the elbow joint. Notably, the conformations of these ligaments in *Pinguinus* and *Mancalla* are not identical. Although the insertions of both of these ligaments are positioned further distally in both *Pinguinus* and *Mancalla* with respect to volant auks, *Pinguinus* is characterized by an extremely elongated insertion of the lig. collaterale ventrale (Figs. 29, 42B), whereas *Mancalla* is distinguished by a rather distally positioned insertion of the lig. dorsale cubiti (Figs. 35, 42C). Although it is conceivable that this conformational difference would have resulted in different responses to dorsal and ventral bending of the elbow joint, and hence might imply differences in wing use between these two taxa, more rigorous mechanical analyses would be required to determine exact functional implications.

**Figure 42.**
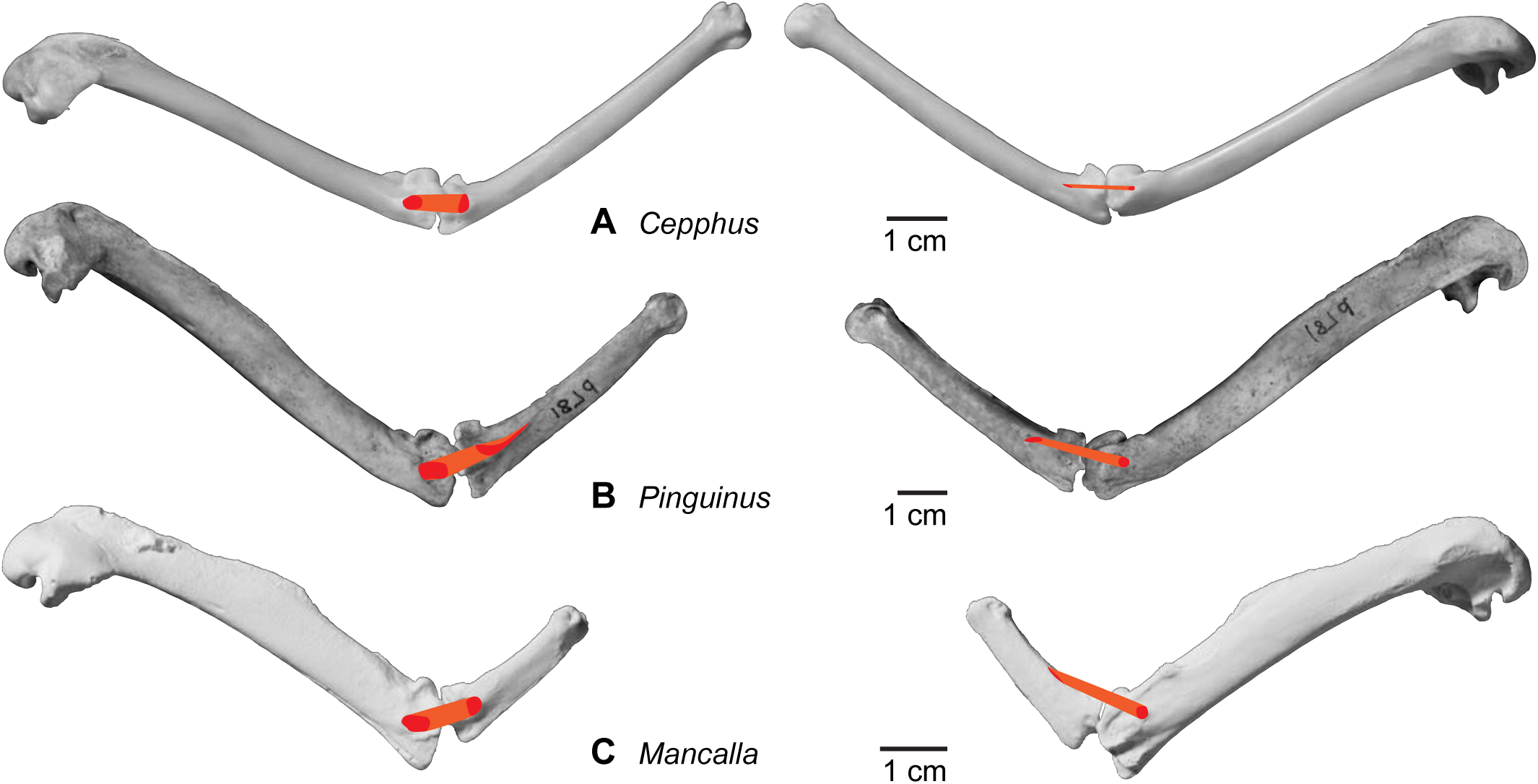
Comparison of elbow joint ligaments in volant and flightless auks. **A**, *Cepphus carbo*, KUGM RAJ 13062101. **B**, *Pinguinus impennis*, UMZC 187.d (mirrored for comparison). **C**, *Mancalla*, cast of LACM 15373. Humeri and ulnae, ventral (left) and dorsal (right) views. Red, attachment sites of the lig. collaterale ventrale and lig. dorsale cubiti; orange, inferred pathways of the ligaments.

To summarize, the wings of *Pinguinus* and Mancallinae are characterized by the following four functionally important modifications: 1) increased mechanical advantages of wing elevators/retractors; 2) reduction of the propatagium; 3) probable loss of function of an intrinsic muscle, the m. flexor digitorum profundus (specifically in *Mancalla*); and 4) increased ligamental bracing of the elbow joint. The first of these features would have assisted with the demanding upstroke of the wings during aquatic flight. The latter three features likely contributed to increasing the rigidity of the wing, transforming the overall wing into a flipper-like apparatus dedicated to aquatic flight.

### Comparison with penguins

The modification of the wing into a flipper-like apparatus in *Pinguinus* and *Mancalla*, as outlined in the previous section, bears numerous similarities to penguins (Sphenisciformes), another major group of flightless wing-propelled divers. It is well understood that extant penguins are characterized by highly modified, flipper-like wings exhibiting small overall size, reduced joint mobility, and reduced intrinsic musculature (Schreiweis 1982; Raikow et al. 1988; Livezey 1989; Louw 1992; Bannasch 1994). At a detailed level, however, flightless auks and penguins exhibit important differences, as well as obvious similarities, in the underlying anatomical architecture of their wings.

Similarities and differences are evident in the conformations of wing elevator/retractor muscles in flightless auks and penguins. In extant Spheniscidae, the m. supracoracoideus, the largest wing elevator, has a single insertion on an oblique ridge on the proximal humerus, which extends in a cranioproximal–caudodistal direction (Figs. 20, 21; Bannasch 1986b). By contrast, in *Pinguinus* and *Mancalla*, this muscle has a dual insertion, on the dorsal margin of the caput humeri and on the elongated tuberculum dorsale, with the latter extending along the shaft more or less in parallel with it (Figs. 26, 27, 32, 33). These conditions may seem radically different at first glance, but are similar insofar as the insertion has extended cranially from the typical position of insertion in volant birds—the tuberculum dorsale on the caudodorsal aspect of the proximal humerus. The specialized conformation in the wing-propelled divers may have some mechanical consequences, especially with regard to dorsal rotation (or “supination”) and retraction/adduction of the humerus, which are induced by the action of this muscle in typical flying birds (Poore et al. 1997; Tobalske and Biewiner 2008). However, the exact consequences of this rearrangement will remain unclear in the absence of rigorous mechanical analyses. Notably, the conformation of this muscle in flightless auks is only slightly modified from the typical condition in Charadriiformes, in which the tendon of the muscle is partly bifurcated and its insertion extends cranially from the tuberculum dorsale (see above). The ancestral condition for Sphenisciformes was probably different, because the insertion is single in *Spheniscus* and in Procellariidae (representing Procellariiformes, the extant sister taxon to Sphenisciformes; e.g., Hackett et al. 2008; Jarvis et al. 2014; Prum et al. 2015; Kimball et al. 2019).

Among other wing elevator/retractor muscles, the m. scapulohumeralis cranialis, which is associated with a prominent scar on the humerus in *Mancalla* (above), is absent in extant Spheniscidae (Schreiweis 1982; see above). By contrast, the m. scapulohumeralis caudalis is prominent and well-developed in extant Spheniscidae, associated with the broadened scapular blade characteristic of penguins (Figs. 18, 19; Schreiweis 1982; Bannasch 1986b, 1994). This differs substantially from the rather unspecialized scapula in the flightless auks (Figs. 24, 30). Notably, the scapula is not broadened in one of the stemward-most penguins known, *Muriwaimanu tuatahi*, whereas it is said to be broadened in other stem-group penguins including *Kupoupou stilwelli*, *Sequiwaimanu rosieae* and *Kumimanu biceae* (although not to the same extent as in extant Spheniscidae, and it is not well preserved in the former two taxa; Slack et al. 2006; Mayr et al. 2017, 2018; Blockland et al. 2019).

The m. latissimus dorsi (partes cranialis et caudalis) is another specialized muscle in Spheniscidae. The two parts of the muscle pass through a retinaculum on the shoulder joint, and share an insertion on a distinct tubercle on the caudal margin of the humerus; both the presence of this retinaculum and the shared insertion are unique to penguins (Schreiweis 1982). There is no evidence of similar muscle specializations in *Pinguinus* and *Mancalla*; neither osteological evidence nor the extant phylogenetic bracket support such an arrangement.

The elongation of the crista deltopectoralis, which was observed in both *Pinguinus* and Mancallinae, is also observed in extant Spheniscidae and many stem-group sphenisciforms (e.g., Ksepka and Clarke 2010). In extant Spheniscidae, this elongated crest hosts a correspondingly elongated attachment site for the lig. propatagiale, and forms a rigid leading edge of the wing (see above; Fig. 21). Although generally similar, it is evident that the elongation of the crista deltopectoralis and the attachment site for the propatagial ligament has been acquired independently in Sphenisciformes, Mancallinae, and *Pinguinus*.

A distinctive feature in *Mancalla*, the lack of the attachment site and pulley for the m. flexor digitorum profundus, is also observed in extant Spheniscidae. In Spheniscidae, the muscle itself is present but is totally tendinous and largely fused with the membrana interossea antebrachii (Schreiweis 1982; see above), thereby lacking contractile function and presumably conferring only passive resilience. In addition, the processus pisiformis of the carpometacarpus—a major osseous pulley for this muscle—is lacking in extant Spheniscidae (Fig. 22). These conditions are associated with the distinctively reduced mobility of forelimb joints in Spheniscidae (Raikow et al. 1988). The processus pisiformis is present in some of the most stemward members of Sphenisciformes (e.g., *Muriwaimanu tuatahi*, *Sequiwaimanu rosieae*, and *Perudyptes devriesi*), whereas it has been lost/reduced in more crownward stem penguins (e.g., *Icadyptes salasi*, and *Kairuku* spp.), potentially reflecting a progressive loss of function in this muscle (Clarke et al. 2007; Ksepka et al. 2008, 2012a; Ksepka and Clarke 2010; Mayr et al. 2018). Although it is tempting to reconstruct the detailed conformation of this muscle in *Mancalla* based on analogy with Spheniscidae, it would be logically circular to regard any such reconstruction as evidence for convergence between these two groups. Nevertheless, based on the osteological features described above, it does seem reasonable to infer a loss or reduction of function of the m. flexor digitorum profundus in *Mancalla*, and therefore a reduction in the mobility of the wrist and digital joints.

Although ligaments of the elbow joints are rather well developed in the flightless auks and *Spheniscus*, their conformation is radically different between the two groups. In *Spheniscus*, the lig. collaterale ventrale has a broad attachment site along the proximal articular surface of the ulna (Fig. 23), unlike the condition in the flightless auks where the attachment site is offset from the articular surface (Figs. 29, 35, 42). The lig. craniale cubiti and lig. collaterale dorsale, which are relatively unspecialized in the flightless auks, are distinctly well developed in *Spheniscus*, with marked attachment scars on the humerus and radius (Figs. 20–23). Most notably, the lig. dorsale cubiti, which is distinctly well developed in *Pinguinus* and *Mancalla* (see above), is absent in *Spheniscus*. In the present study, this ligament was observed in all charadriiform taxa examined, but not in the closest extant relatives of penguins (*Gavia*, Procellariidae, and *Spheniscus*; Fig. 43). Indeed, reference to this ligament is virtually absent in the literature except in studies of Charadriiformes (Stettenheim 1959), suggesting that this ligament is unique to the group. *Pinguinus* and *Mancalla* exhibit a modification to the ancestral lig. dorsale cubiti to enhance bracing of the dorsal side of the elbow joint at the humero-ulnar articulation. By contrast, penguins, lacking this ligament, exhibit a modified lig. collaterale dorsale to enhance bracing at the humero-radial articulation. The exact mechanical consequences of these conformational differences between flightless auks and penguins remain unclear at present, but the overall function (bracing of the elbow joint) appears similar, given the limited range of motion at the elbow joint in extant penguins (Raikow et al. 1988) and flightless auks.

**Figure 43.**
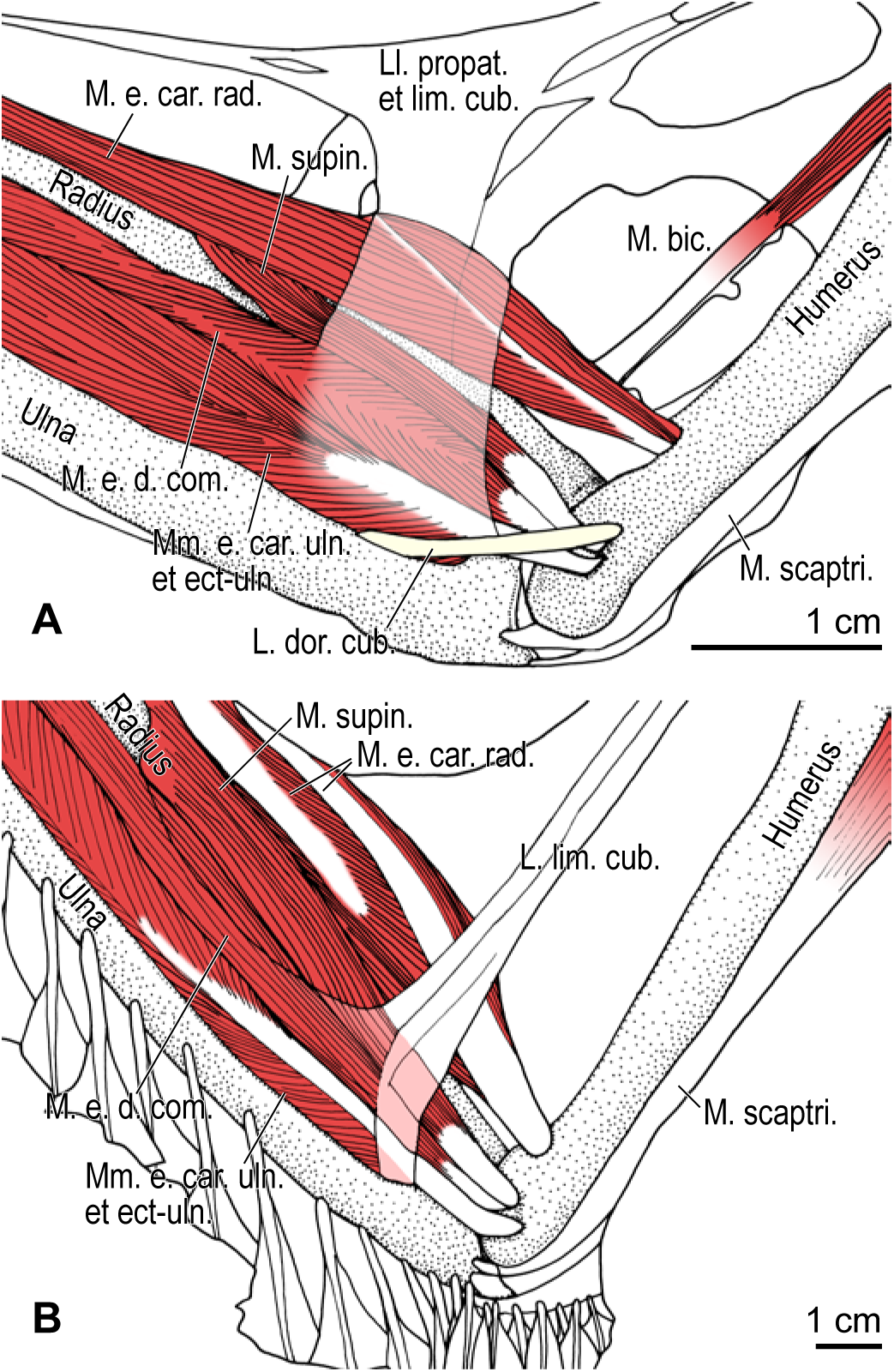
Comparison of the elbow musculature in *Cerorhinca monocerata* (**A**) and *Gavia adamsii* (**B**), dorsal view. Note the presence and absence of the lig. dorsale cubiti in the former and latter species, respectively. See Table 3 for abbreviations.

### Incomplete convergence of anatomical structures

Taken together, the wings of the different lineages of flightless wing-propelled divers—the auks *Pinguinus* and *Mancalla* in Charadriiformes, and penguins Sphenisciformes—appear to be modified to accommodate similar functional demands: powerful wing elevation and increased wing rigidity. In this sense, the wings of these wing-propelled divers are functionally convergent as apparatuses facilitating aquatic flight. However, at finer levels of anatomical detail, the mechanisms by which these modifications have been achieved are clearly distinct. As such, the degree of anatomical convergence between flightless auks and penguins is incomplete (here, the term incomplete convergence is used in a broad, general sense as in Herrel et al. 2004; not in the restrictive sense of Collar et al. 2014). Importantly, several key anatomical differences between these groups appear to reflect differences in the ancestral conditions for these lineages. For instance, flightless auks exhibit modified ancestral anatomical structures unseen in penguins, such as the bifurcated insertion of the m. supracoracoideus and the development of the lig. dorsale cubiti. These flightless wing-propelled divers illustrate that, even in the face of similar functional demands, the nature of convergent evolutionary change of anatomical structures is dictated to an important degree by ancestral starting points.

The recruitment of the ancestrally present lig. dorsale cubiti for a role in limiting elbow joint mobility during aquatic flight in flightless auks can be regarded as an exaptation (Gould and Vrba 1982). The origin of this structure evidently predates the transition to wing-propelled diving, as it is present in all charadriiform taxa examined. It was Bock (1959) who articulated the concepts of exaptation (preadaptation in his terminology) and the ancestral-state dependency of anatomical evolution. That is, lineages with ancestrally differing anatomical structures often respond to novel, similar selection forces in dissimilar ways, and can acquire disparate structures specialized—or “exapted”—for the same function through multiple evolutionary pathways (i.e., non-convergent evolutionary trajectories). The resultant specialized structures, despite differing anatomically, can exhibit qualitatively similar performance for that function (Bock 1959, 1977; Bock and Miller 1959), illustrating redundancy in form–function relationships. More recently, this redundancy has come to be investigated in more quantitative ways in mechanically tractable systems (e.g., Wainwright et al. 2005; Wainwright 2007; Muñoz 2019). These systems have illustrated that various anatomical outcomes may result from selection for a particular function, depending on ancestral starting points (Alfaro et al. 2004; Thompson et al. 2017).

Recent studies have shown that superficially convergent phenotypes do arise via diverse evolutionary pathways and/or developmental mechanisms in different clades (Ng and Smith 2016; Morinaga and Bergmann 2017; Bergmann and Morinaga 2019; Arbour and Zanno 2020). The prevalence of incomplete convergence is now widely appreciated in many different systems, although the detailed underpinnings of incompleteness largely remain to be explored (e.g., Losos 2010, 2011; Kaeuffer et al. 2012; Collar et al. 2014; Moen et al. 2016; Dobler et al. 2019). Also unexplored is the extent to which the retention and/or modification of ancestral conditions have influenced evolutionary trajectories in these examples. However, for convergence to truly exemplify the predictability of anatomical evolution, it is clear that ancestral conditions must be taken into account.

It could be argued that anatomical differences between the flightless auks and extant penguins may be the result of a lesser degree of specialization for flightless wing-propelled diving in the auks, possibly reflecting a shorter amount of elapsed time since these lineages transitioned to this lifestyle. The oldest known fossil penguins, e.g., *Waimanu manneringi* and *Kupoupou stilwelli*, are Paleocene in age (>60 Ma), and are considered to have already been flightless wing-propelled divers (Slack et al. 2006; Blockland et al. 2019). The transition to flightlessness in penguins would therefore have taken place even earlier, in the Paleocene or perhaps the latest Cretaceous (Prum et al. 2015). On the other hand, the oldest known species of Mancallinae, *Miomancalla wetmorei*, is middle Miocene in age (<10 Ma), and that of *Pinguinus*, *Pinguinus alfrednewtoni*, is Pliocene (∼4.4 Ma), although the dates of divergence from their respective volant sister groups are inferred to be substantially older (>28 Ma and >11 Ma, respectively; Smith and Clarke 2015). As such, extant penguins are the product of a much longer flightless evolutionary history than are either of the extinct flightless auk lineages. Nonetheless, there is little evidence that early stem-group penguins passed through a phase where their wings more closely resembled those of flightless auks; instead, these lineages seem to reflect non-parallel trajectories toward broadly similar morphological solutions (but see also Mayr et al. 2020b).

### Concluding remarks

Reconstruction of the wing musculature in extinct, flightless auks (*Pinguinus* and *Mancalla*) has facilitated an investigation into their functional anatomy with respect to wing-propelled diving, as well as a critical evaluation of the hypothesis of their evolutionary convergence with penguins at a previously unmatched level of detail. Although most major wing muscles and ligaments in the flightless auks could be reliably reconstructed from osteological correlates on bones, some uncertainty remains, partly due to a lack of suitable fossil material for some anatomical regions (i.e., free carpals and phalanges in Mancallinae). Recovery of further fossils and the detailed examination of previously collected specimens in museums will be required to complement the present results.

At a broad scale, the wings of the flightless auks were functionally convergent with those of penguins. This holds with respect to the powerful action of wing elevator/retractor muscles and reduced joint mobility in the wings, both of which are associated with clear functional advantages in aquatic flight. Nevertheless, important differences emerged at finer anatomical levels, which are most likely the product of differences in the ancestral conditions from which penguins and flightless auks arose.

It has previously been suggested that differing skeletal proportions in the wings of mancalline auks and penguins may have had mechanical consequences (Smith 2011; but see above for potential caveats). It has also been proposed that variability of limb skeletal proportions in birds is concentrated along the direction of clade-specific ontogeny, potentially biasing the directionality of evolutionary change (Watanabe 2018a). As such, it is possible that developmental bias precludes some wing-propelled divers from obtaining skeletal proportions that would be functionally optimal. Rigorous mechanical analyses will be necessary to fully evaluate qualitative speculations about the function of the anatomical structures discussed in the present study. Such investigations could evaluate whether the anatomical differences identified between flightless auks and penguins would have been associated with notable mechanical consequences. The avian musculoskeletal system leaves much to be explored, but synthesis of information from varied approaches, including soft tissue anatomy, the identification of osteological correlates, and mechanical analyses, will pave the way toward a more comprehensive understanding of avian morphological evolution.

## Acknowledgements

The authors would like to thank Yutaka Watanuki, Makoto Hasebe, Shin Matsui, Tatsuo Sato, Kensuke Yasui, Kentaro Kazama, Dale M. Kikuchi, Nobuhiko Sato, Sakiko Matsumoto, Kenji Hoshina, and John R. Hutchinson for help with modern anatomical specimens, and Judith White, Joanne H. Cooper, Samuel A. McLeod, Vanessa R. Rhue, Kesler A. Randall, Matt Lowe, and Michael Brooke for access to museum collections. The authors declare no conflict of interest. The work was partly supported by the Newton International Fellowship by the Royal Society [NIF\R1\180520 to JW]; and UKRI Future Leaders Fellowship [MR/S032177/1 to DJF].

## Supplementary Figures S1–S32

**Figure S1.**
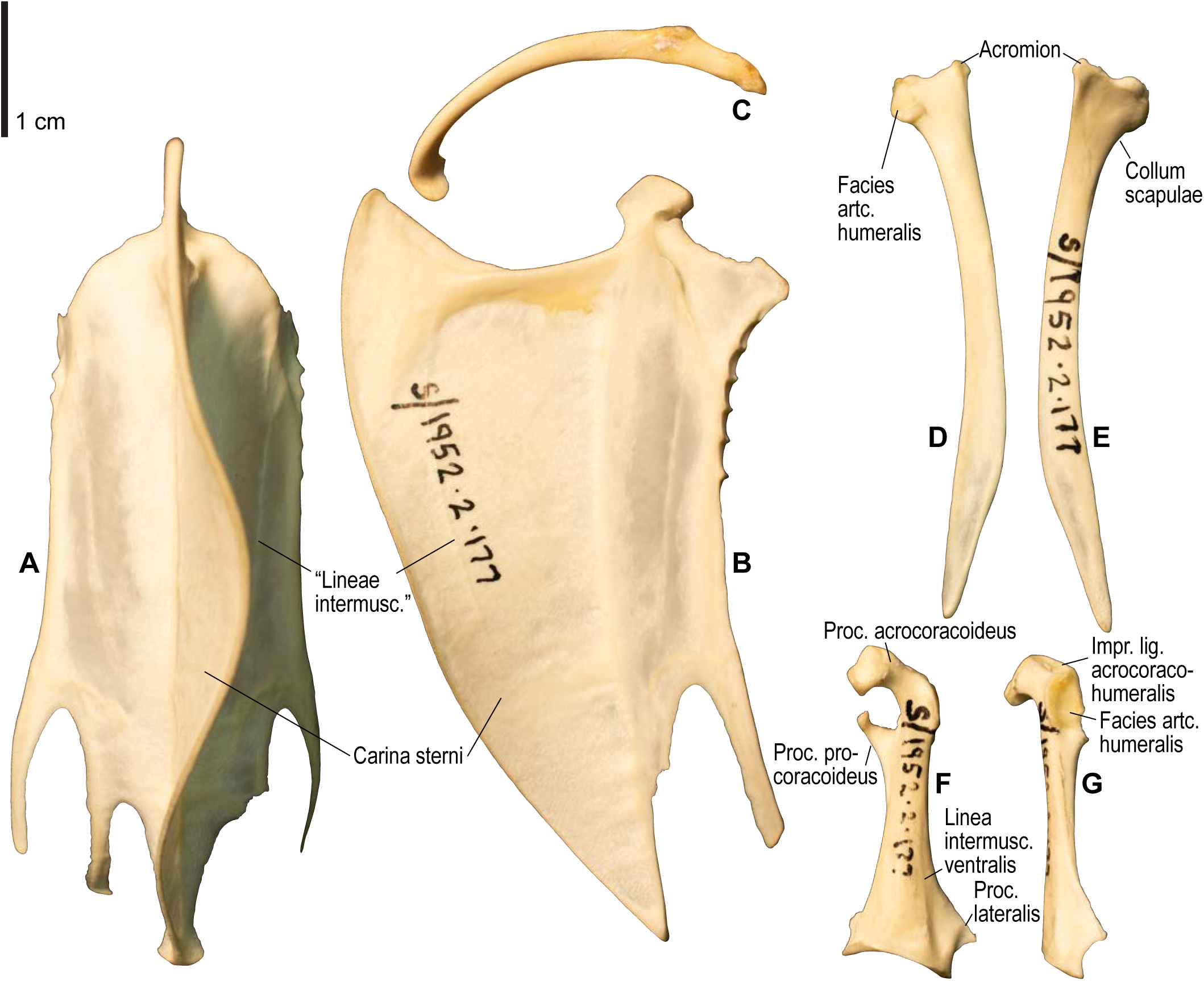
Osteology of *Pluvialis apricaria*, pectoral girdle elements. Drawn on NHMUK S/1952.2.178 (furcula) and S/1952.2.177 (other elements). Sternum in ventral (**A**) and left lateral (**B**) views; furcula in left lateral view (**C**); left scapula in lateral (**D**) and medial (**E**) views; left coracoid in ventral (**F**) and lateral (**G**) views. Major osteological landmarks mentioned in text are designated. Abbreviations: artc., articularis; impr., impressio; intermusc., intermuscularis/intermusculares; lig., ligamenti; m., musculi; proc., processus; tuberc., tuberculum.

**Figure S2.**
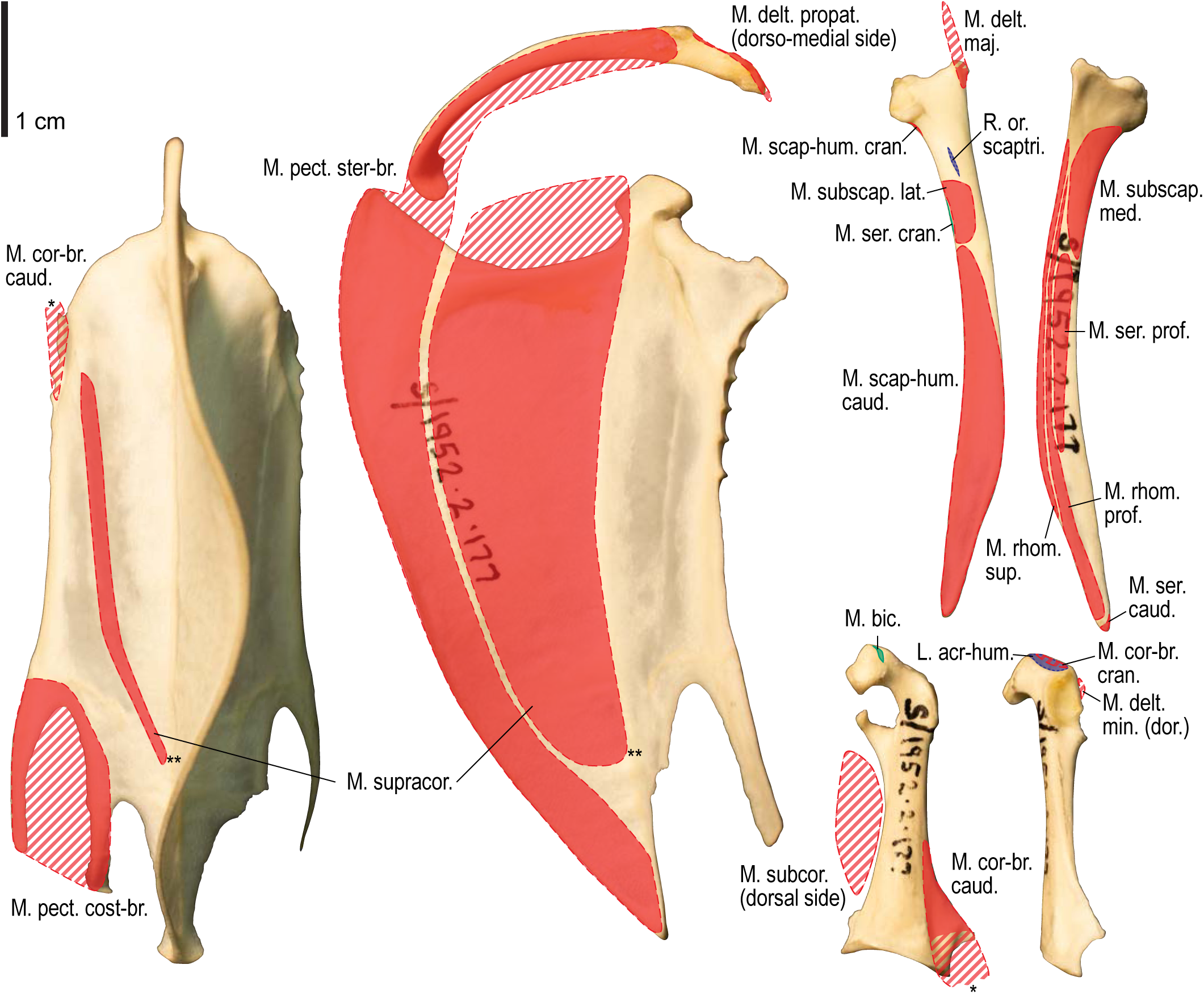
Osteological correlates of major wing muscles and ligaments in *Pluvialis apricaria*, pectoral girdle elements. Drawn on NHMUK S/1952.2.178 (furcula) and S/1952.2.177 (other elements). Note that only reliably identified attachment sites are shown, and the gaps between some adjacent attachment sites are exaggerated for distinction. Asterisks denote continuous attachment sites across panels. Red fill with broken outline, fleshy (direct) attachment of muscles; green fill with solid outline, tendinous/aponeurotic (indirect) attachment of muscles; blue fill with dotted outline, attachment of ligaments; stroked fill, attachment on ligaments/membranes.

**Figure S3.**
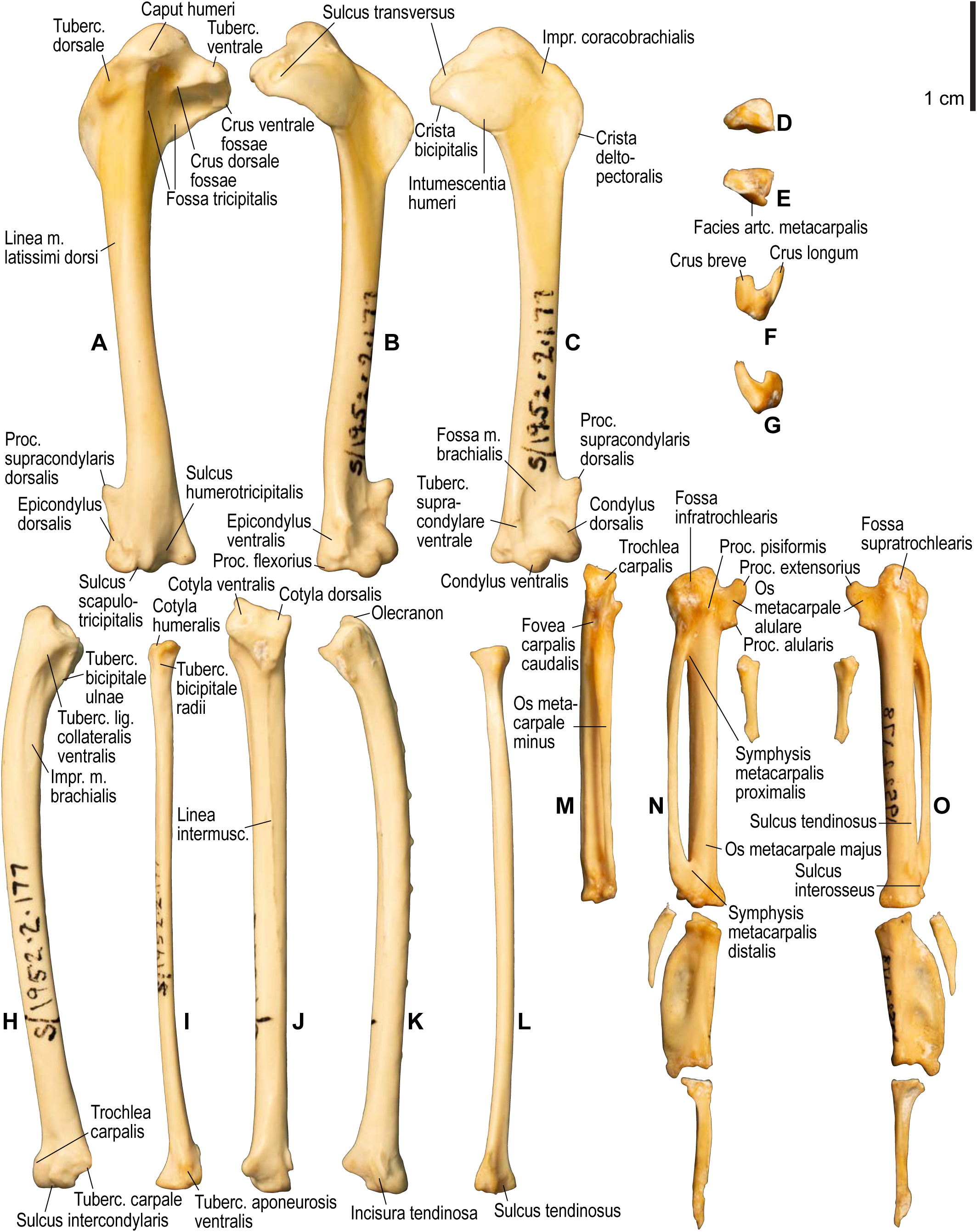
Osteology of *Pluvialis apricaria*, wing elements. Drawn on NHMUK S/1952.2.178 (manual elements) and S/1952.2.177 (other elements). Left humerus in caudal (**A**), ventral (**B**), and cranial (**C**) views; left radiale in cranial (**D**) and caudal (**E**) views; left ulnare in proximal (**F**) and distal (**G**) views; left ulna in ventral (**H**), cranial (**J**), and dorsal (**K**) views; left radius in ventral (**I**) and cranial (**L**) views; right carpometacarpus and phalanges (mirrored for comparison) in caudal (**M**; phalanges not shown), ventral (**N**), and dorsal (**O**) views. See Figure S1 for abbreviations.

**Figure S4.**
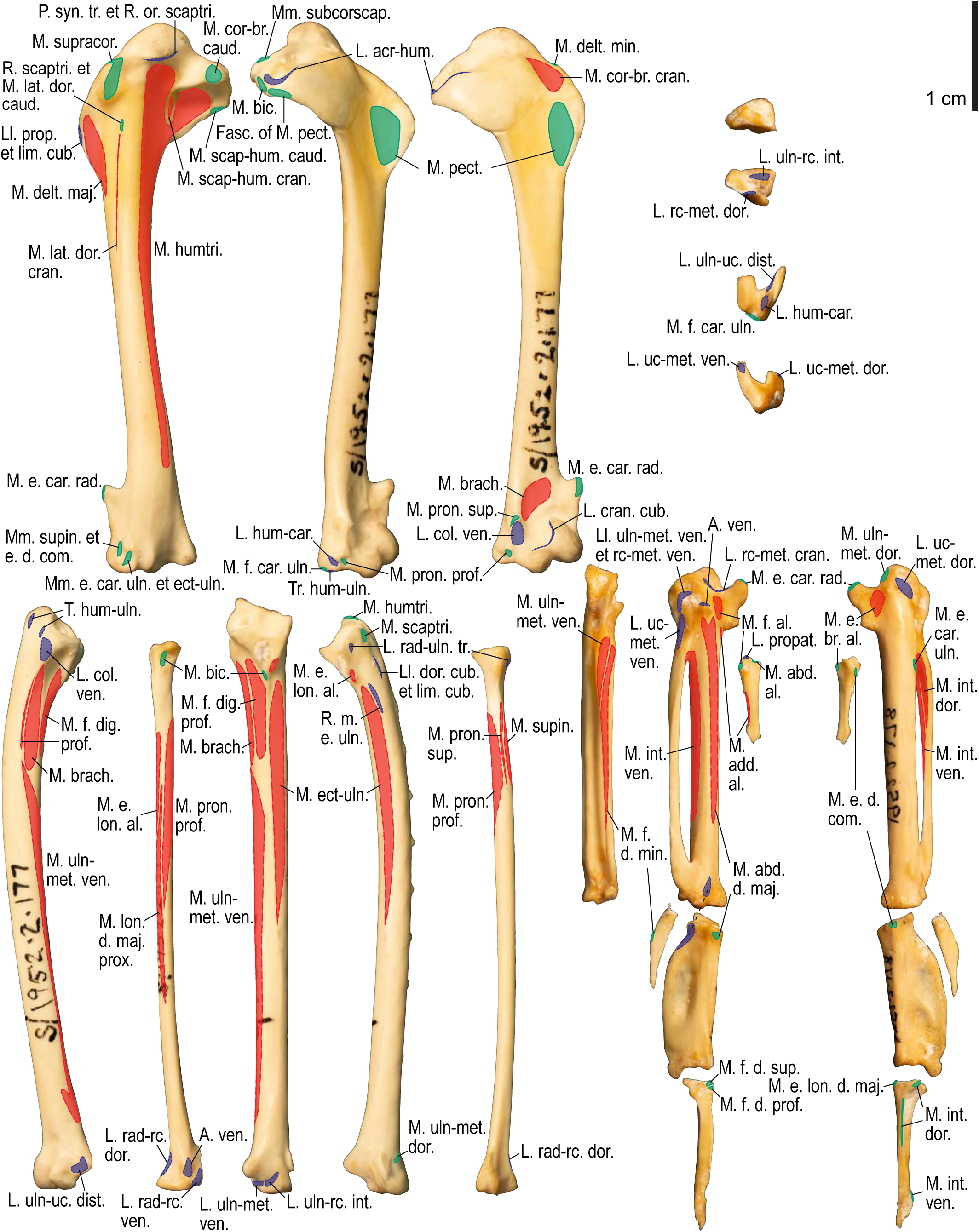
Osteological correlates of major wing muscles and ligaments in *Pluvialis apricaria*, wing elements. Drawn on NHMUK S/1952.2.178 (manual elements) and S/1952.2.177 (other elements). Manual elements are mirrored for comparison. Due to space restrictions, the labels for some distal wing ligaments are not shown; broken lines show correspondence of attachment sites for these ligaments. See Figure S2 for legends.

**Figure S5.**
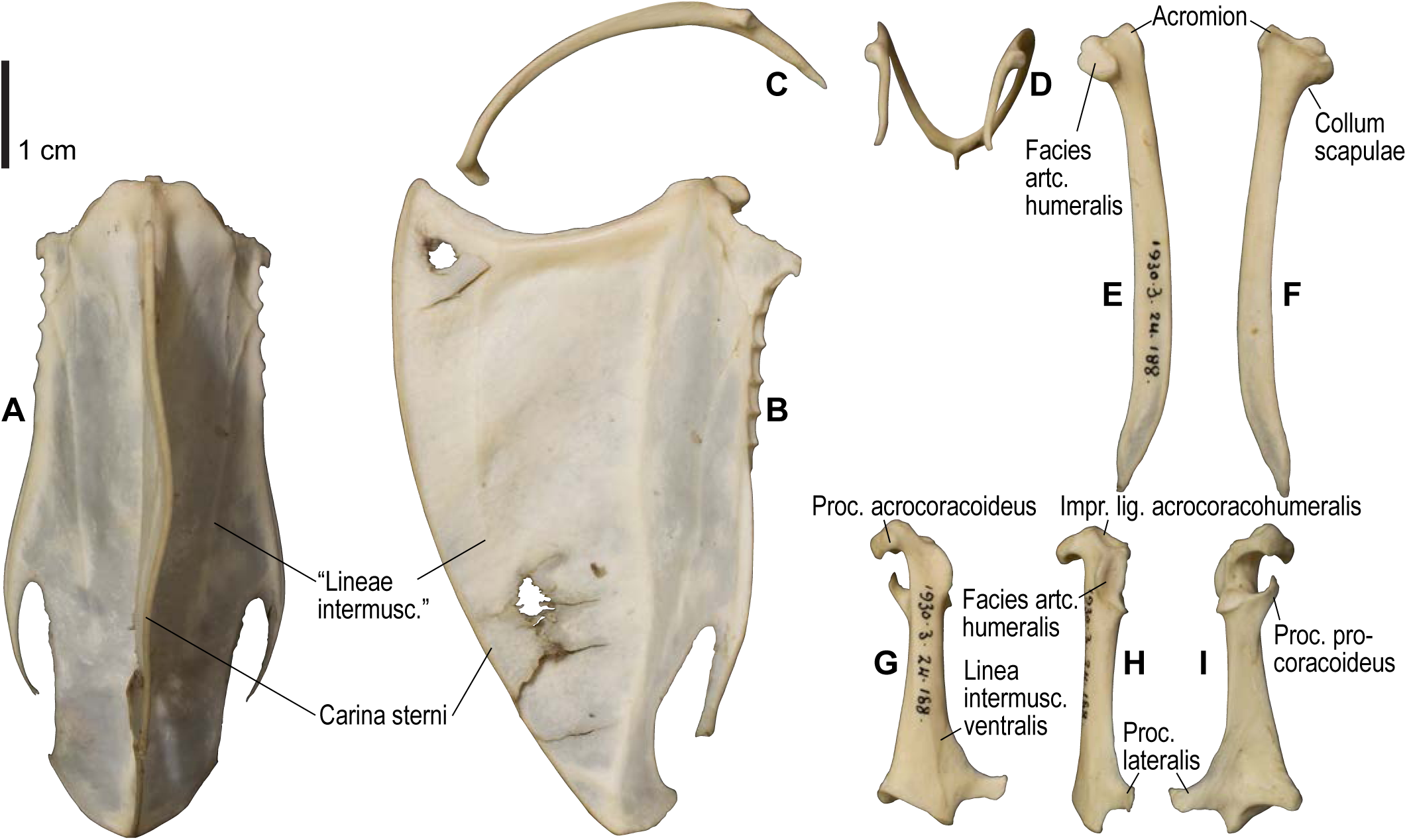
Osteology of *Scolopax rusticola*, pectoral girdle elements. Drawn on NHMUK 1930.3-24.188. Sternum in ventral (**A**) and left lateral (**B**) views; furcula in left lateral (**C**) and dorsal (**D**) views; left scapula in lateral (**E**) and medial (**F**) views; left coracoid in ventral (**G**), lateral (**H**), and dorsal (**I**) views. See Figure S1 for abbreviations.

**Figure S6.**
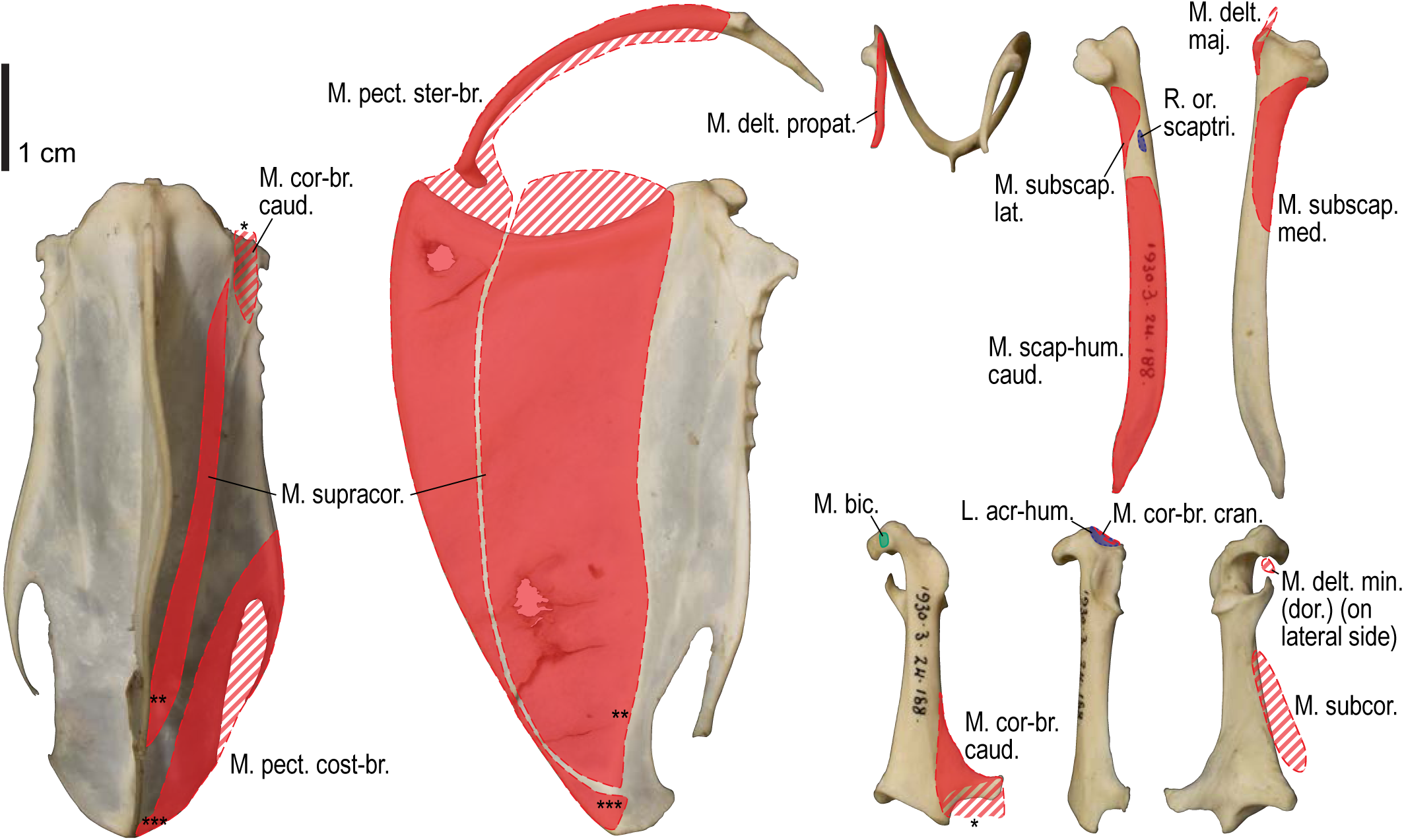
Osteological correlates of major wing muscles and ligaments in *Scolopax rusticola*, pectoral girdle elements. Drawn on NHMUK 1930.3-24.188. See Figure S2 for legends.

**Figure S7.**
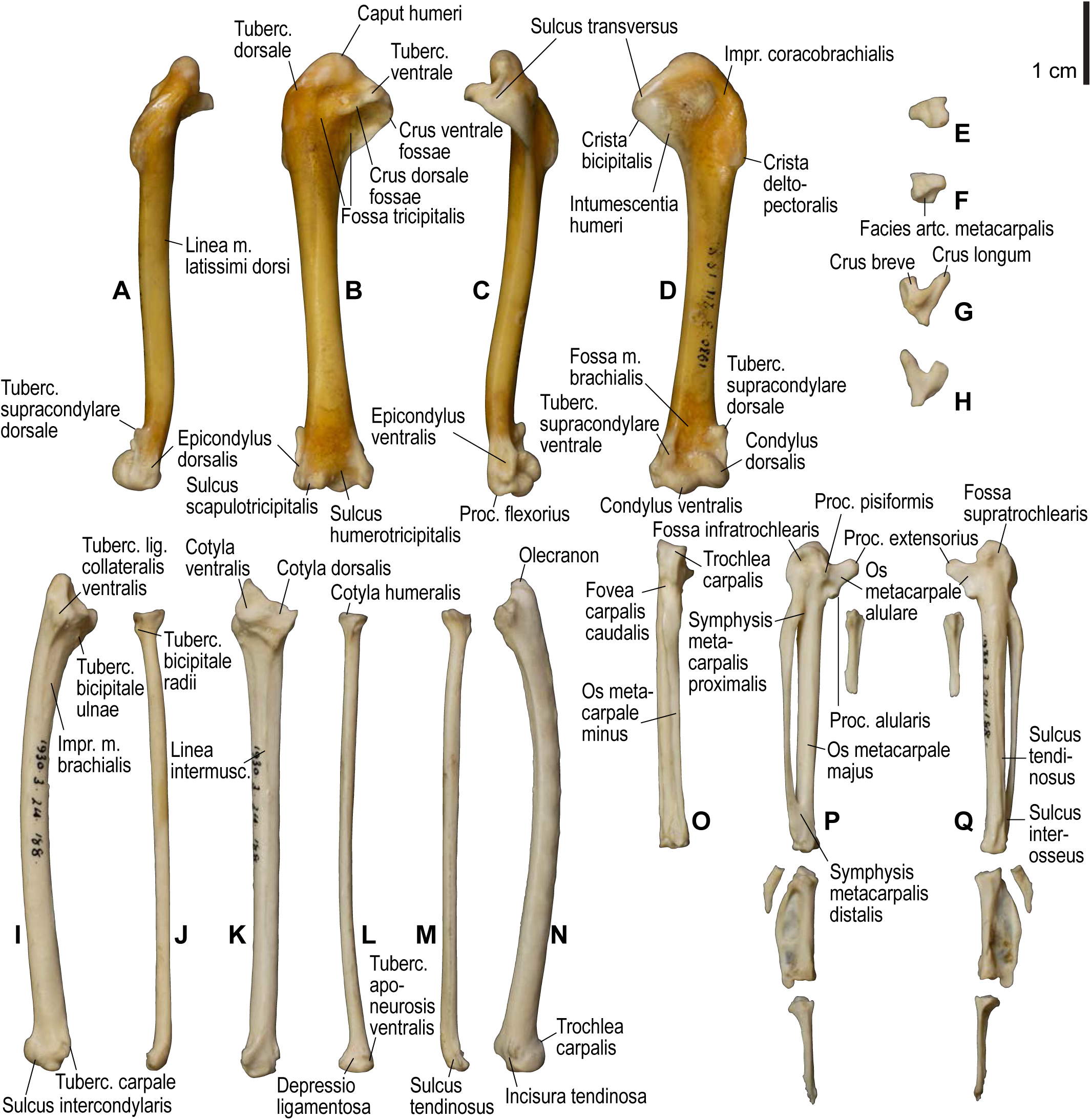
Osteology of *Scolopax rusticola*, wing elements. Drawn on NHMUK 1930.3-24.188. Left humerus in dorsal (**A**), caudal (**B**), ventral (**C**), and cranial (**D**) views; left radiale in cranial (**E**) and caudal (**F**) views; left ulnare in proximal (**G**) and distal (**H**) views; left ulna in ventral (**I**), cranial (**K**), and dorsal (**N**) views; left radius in ventral (**J**), caudal (**L**), and dorsal (**L**) views; left carpometacarpus and phalanges in caudal (**O**; phalanges not shown), ventral (**P**), and dorsal (**Q**) views. See Figure S1 for abbreviations.

**Figure S8.**
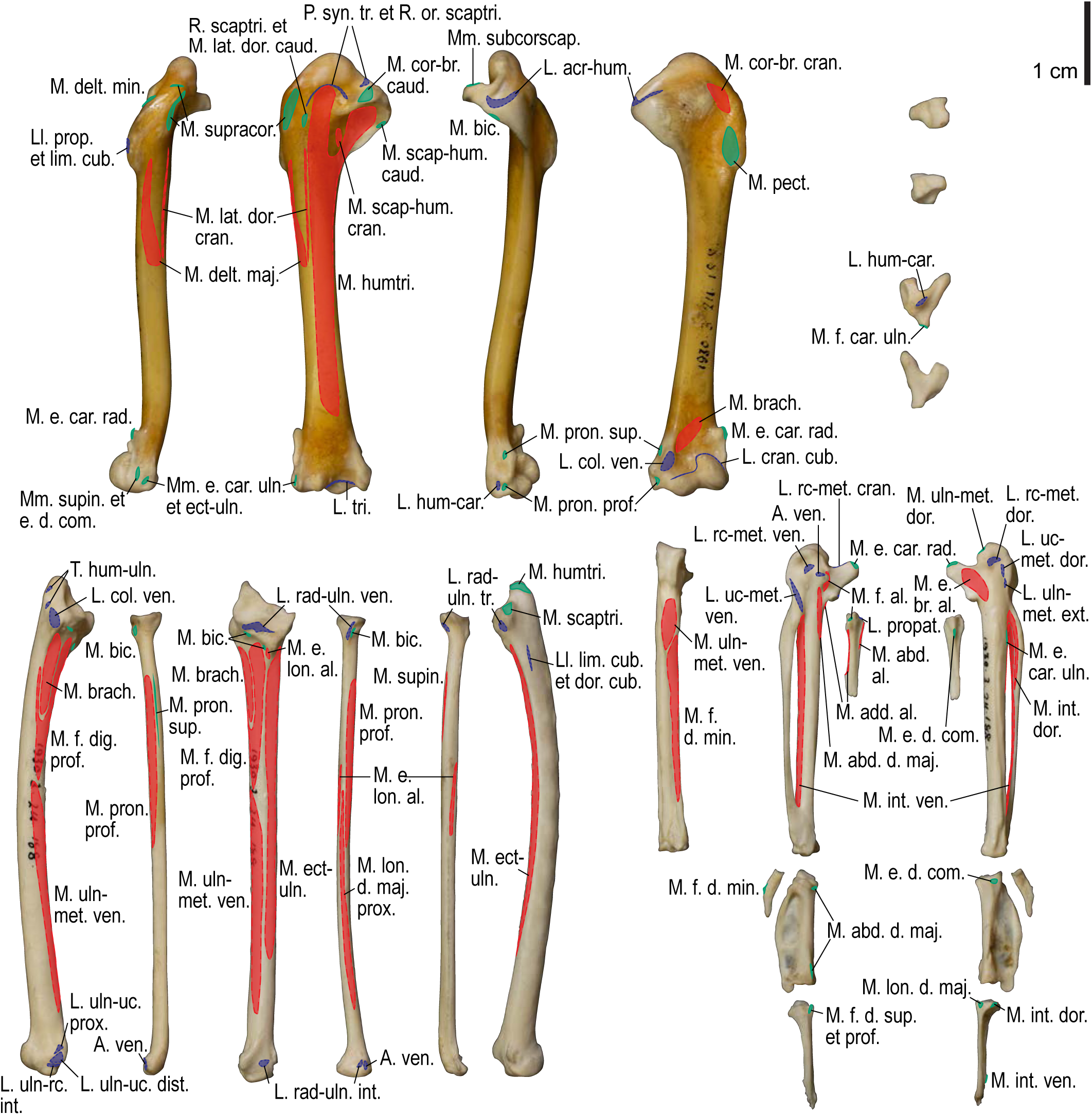
Osteological correlates of major wing muscles and ligaments in *Scolopax rusticola*, wing elements. Drawn on NHMUK 1930.3-24.188. See Figure S2 for legends.

**Figure S9.**
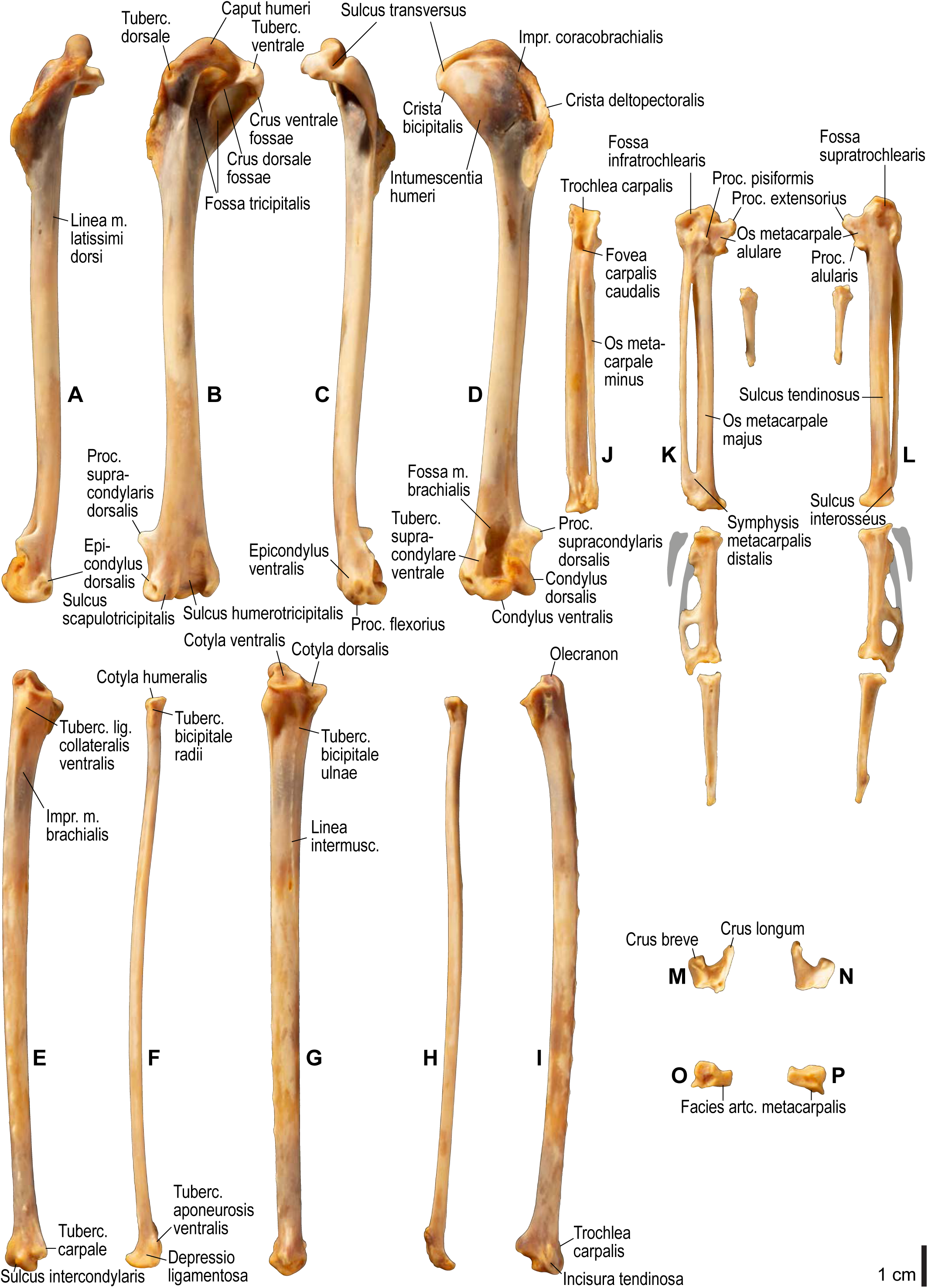
Osteology of *Larus schistisagus*, wing elements. Drawn on KUGM RAJ AO13071502. Left humerus in dorsal (**A**), caudal (**B**), ventral (**C**), and cranial (**D**) views; left ulna in ventral (**E**), cranial (**G**), and dorsal (**I**) views; left radius in ventral (**F**) and dorsal (**H**) views; left carpometacarpus and phalanges in caudal (**J**; phalanges not shown), ventral (**K**), and dorsal (**L**) views; left ulnare in proximal (**M**) and distal (**N**) views; left radiale in cranial (**O**) and caudal (**P**) views. Approximate outlines of missing/broken phalanges are shown with gray shading. See Figure S1 for abbreviations.

**Figure S10.**
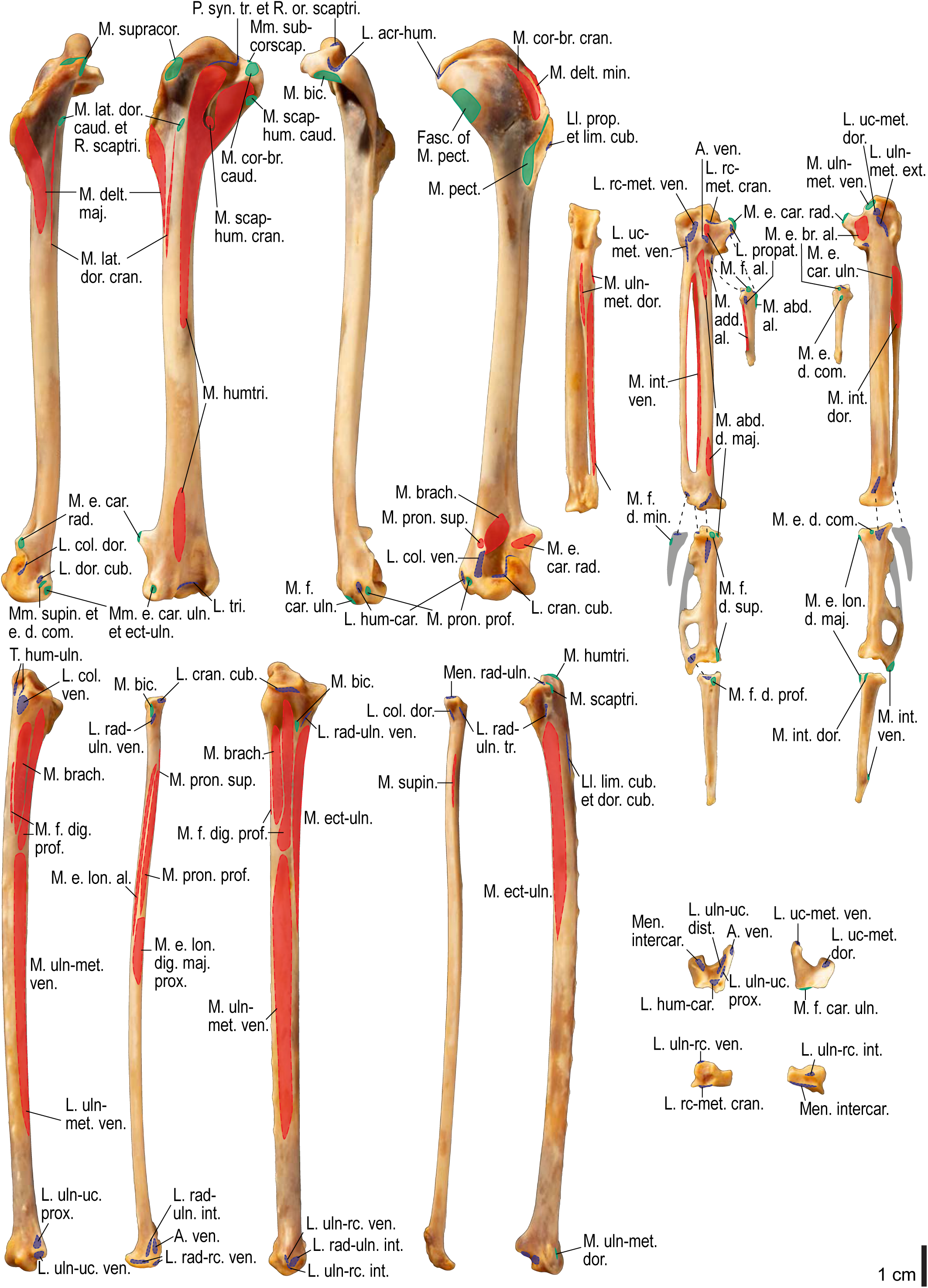
Osteological correlates of major wing muscles and ligaments in *Larus schistisagus*, wing elements. Drawn on KUGM RAJ AO13071502. See Figures S2, S4, and S9 for legends.

**Figure S11.**
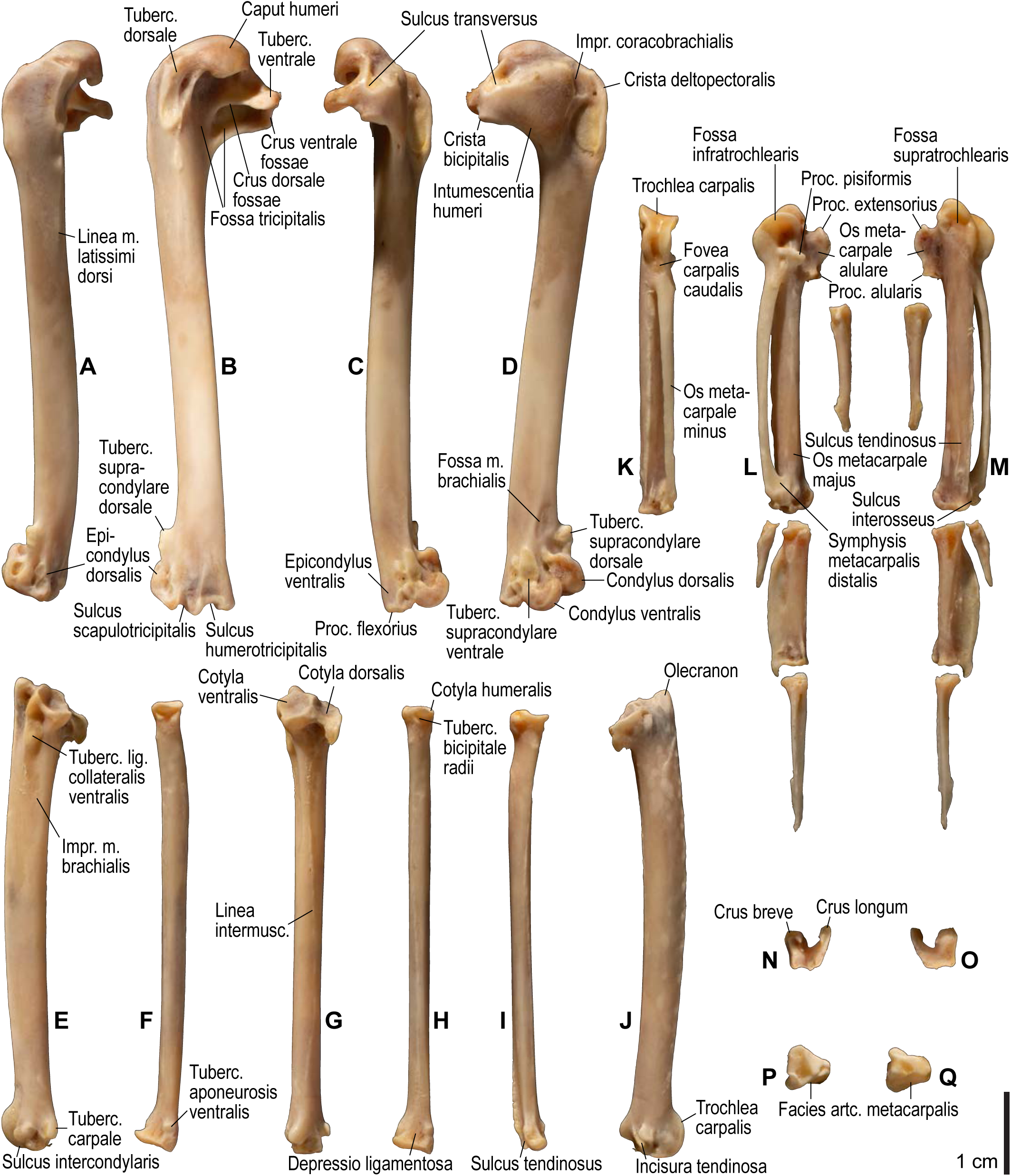
Osteology of *Cerorhinca monocerata*, wing elements. Drawn on KUGM RAJ AO13070802. Left humerus in dorsal (**A**), caudal (**B**), ventral (**C**), and cranial (**D**) views; left ulna in ventral (**E**), cranial (**G**), and dorsal (**J**) views; left radius in ventral (**F**), caudal (**H**), and dorsocaudal (**I**) views; left carpometacarpus and phalanges in caudal (**K**; phalanges not shown), ventral (**L**), and dorsal (**M**) views; left ulnare in proximal (**N**) and distal (**O**) views; left radiale in cranial (**P**) and caudal (**Q**) views. See Figure S1 for abbreviations.

**Figure S12.**
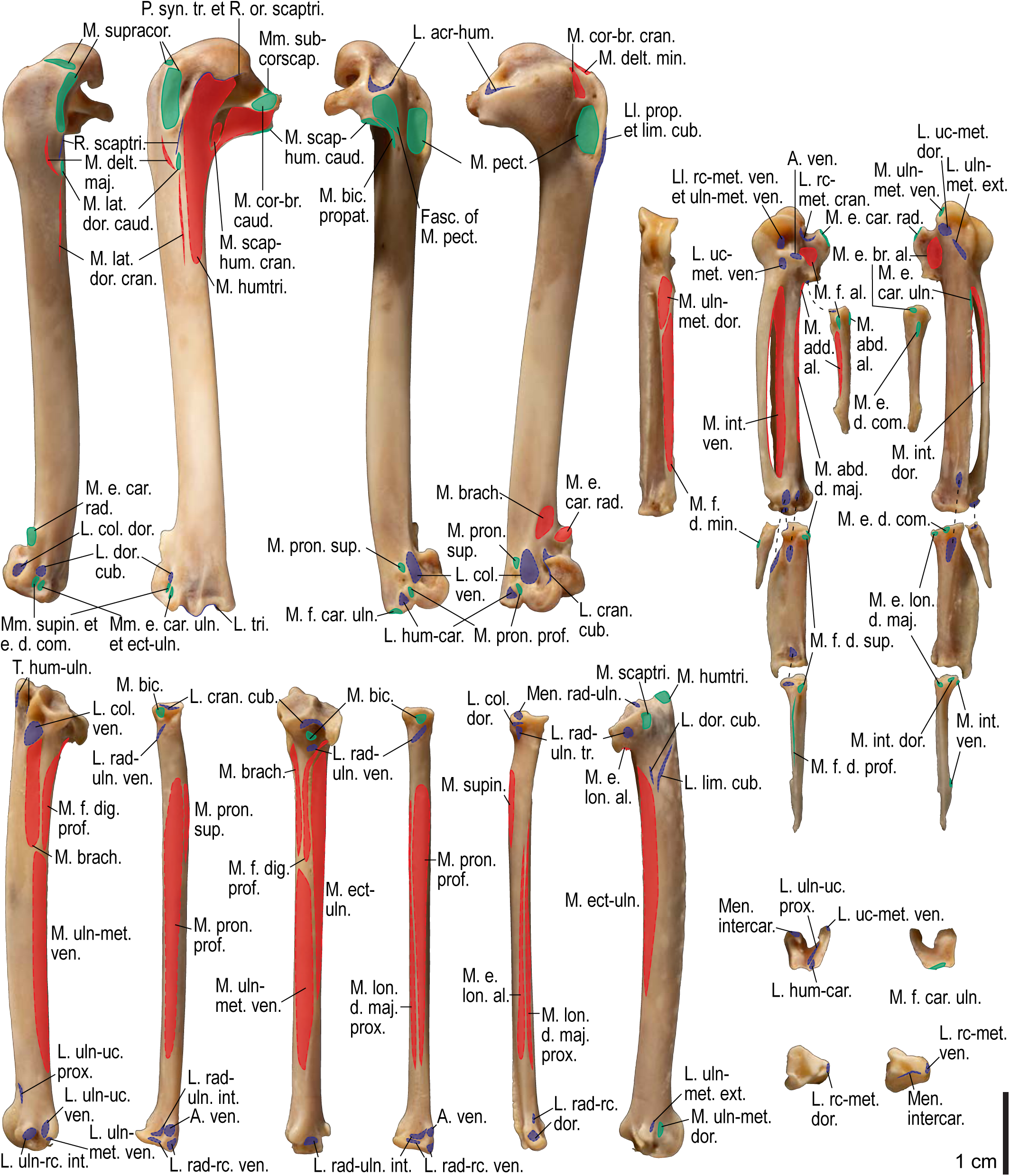
Osteological correlates of major wing muscles and ligaments in *Cerorhinca monocerata*, wing elements. Drawn on KUGM RAJ AO13070802. See Figures S1 and S2 for legends.

**Figure S13.**
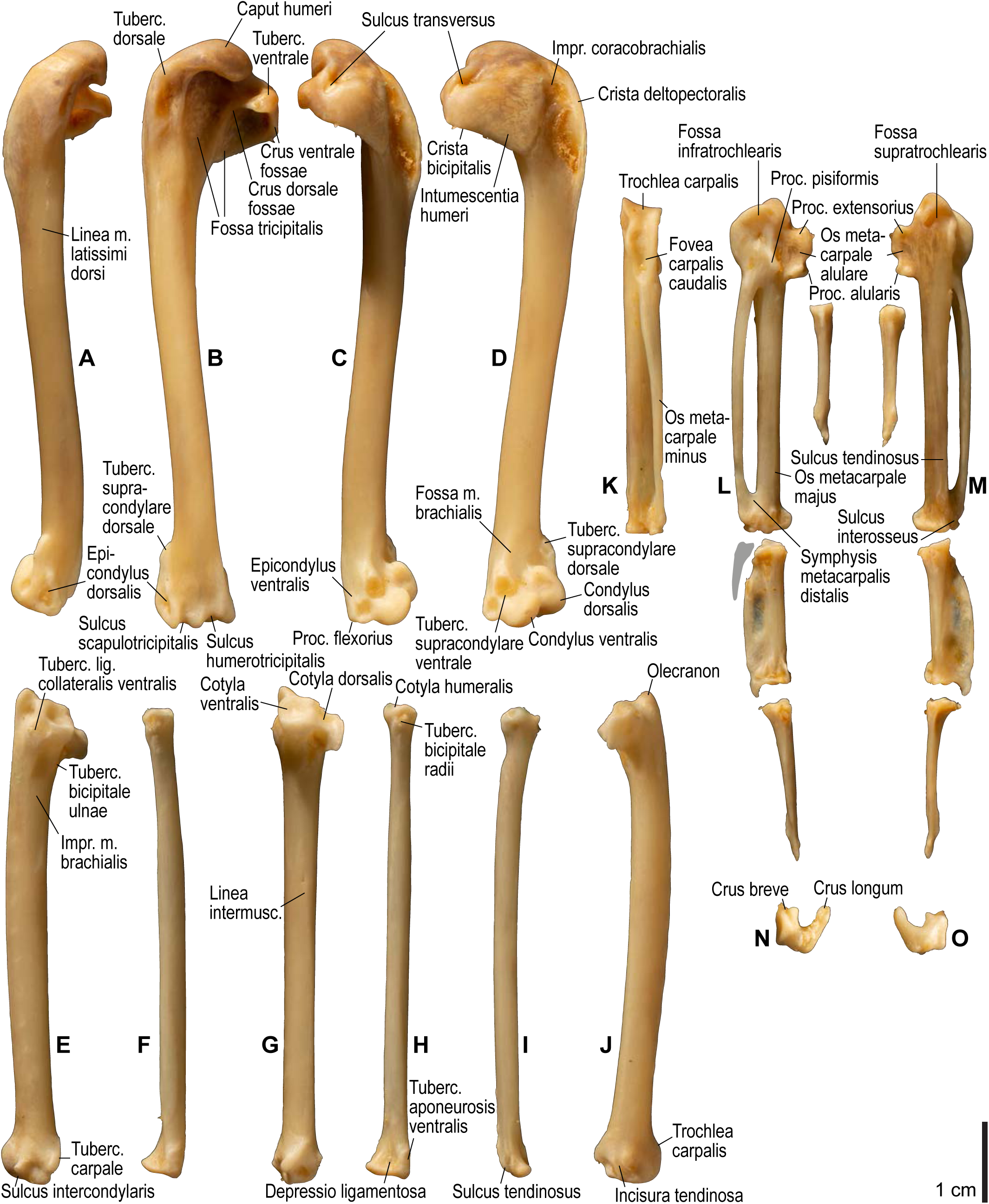
Osteology of *Cepphus carbo*, wing elements. Drawn on KUGM RAJ AO13062101. Left humerus in dorsal (**A**), caudal (**B**), ventral (**C**), and cranial (**D**) views; left ulna in ventral (**E**), cranial (**G**), and dorsal (**J**) views; left radius in ventral (**F**), caudal (**H**), and dorsocaudal (**I**) views; left carpometacarpus and phalanges in caudal (**K**; phalanges not shown), ventral (**L**), and dorsal (**M**) views; left ulnare in proximal (**N**) and distal (**O**) views. Approximate outline of the missing phalanx of the minor digit is shown with gray shading. See Figure S1 for abbreviations.

**Figure S14.**
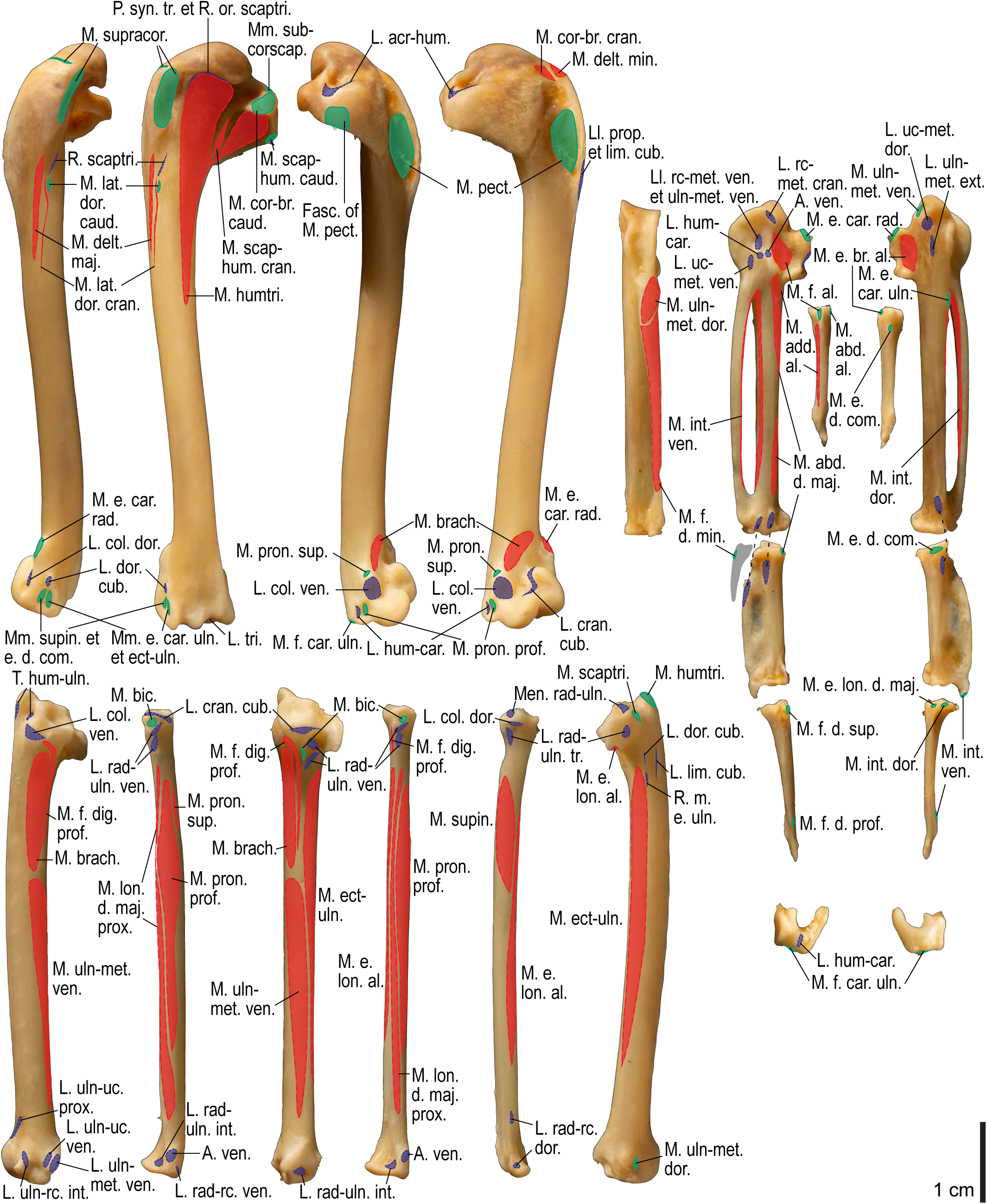
Osteological correlates of major wing muscles and ligaments in *Cepphus carbo*, wing elements. Drawn on KUGM RAJ AO13062101. See Figures S2, S4, and S13 for legends.

**Figure S15.**
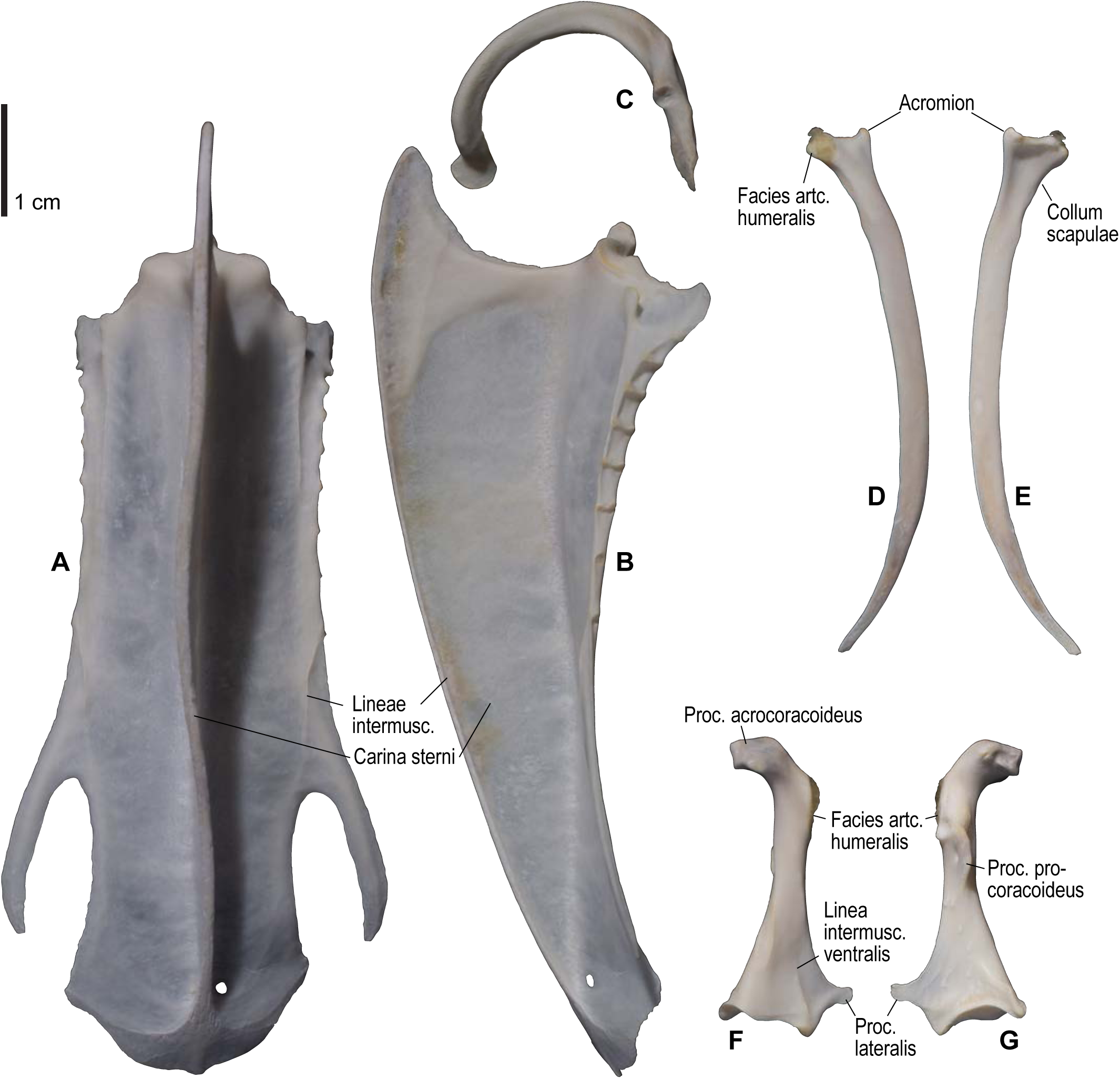
Osteology of *Synthliboramphus antiquus*, pectoral girdle elements. Drawn on KUGM RA 1311. Sternum in ventral (**A**) and left lateral (**B**) views; furcula in left lateral view (**C**); left scapula in lateral (**D**) and medial (**E**) views; left coracoid in ventral (**F**) and dorsal (**G**) views. See Figure S1 for abbreviations.

**Figure S16.**
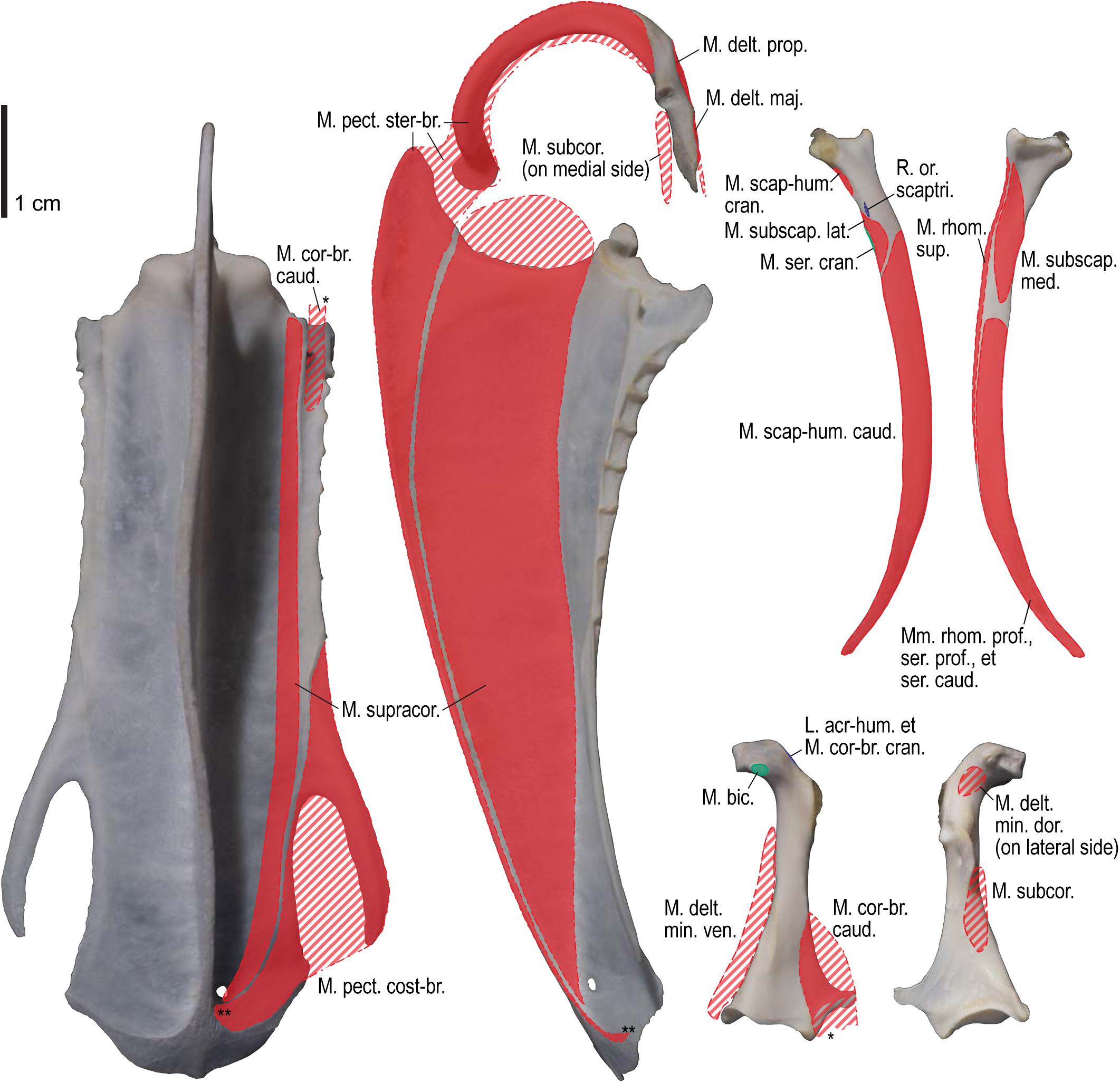
Osteological correlates of major wing muscles and ligaments in *Synthliboramphus antiquus*, pectoral girdle elements. Drawn on KUGM RA 1311. See Figure S2 for legends.

**Figure S17.**
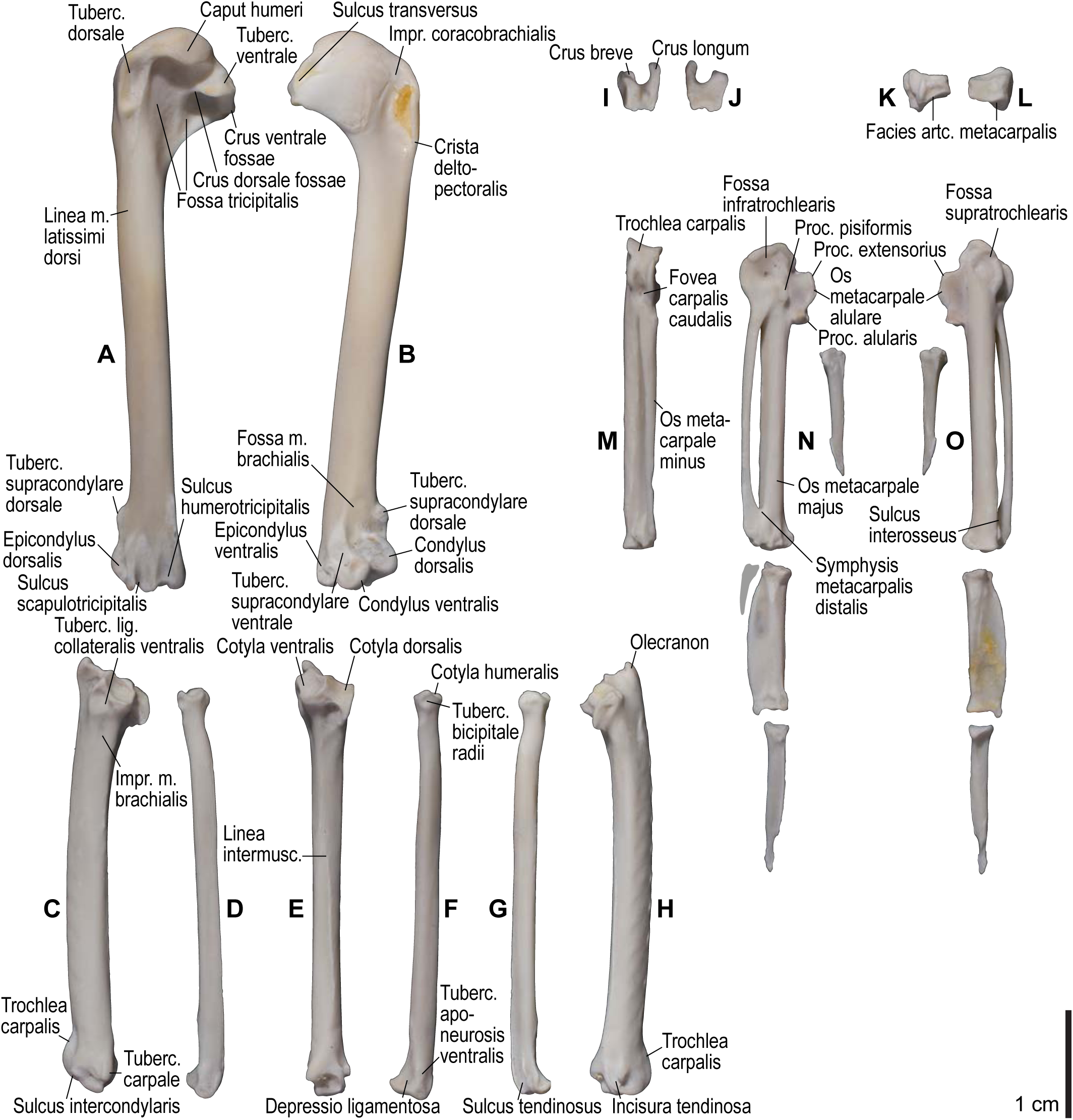
Osteology of *Synthliboramphus antiquus*, wing elements. Drawn on KUGM RA 1311. Left humerus in caudal (**A**) and cranial (**B**) views; left ulna in ventral (**C**), cranial (**E**), and dorsal (**H**) views; left radius in ventral (**D**), caudoventral (**F**), and dorsal (**G**) views; left ulnare in proximal (**I**) and distal (**J**) views; left radiale in cranial (**K**) and caudal (**L**) views; left carpometacarpus and phalanges in caudal (**M**; phalanges not shown), ventral (**N**), and dorsal (**O**) views. Approximate outline of the missing phalanx of the minor digit is shown with gray shading. See Figure S1 for abbreviations.

**Figure S18.**
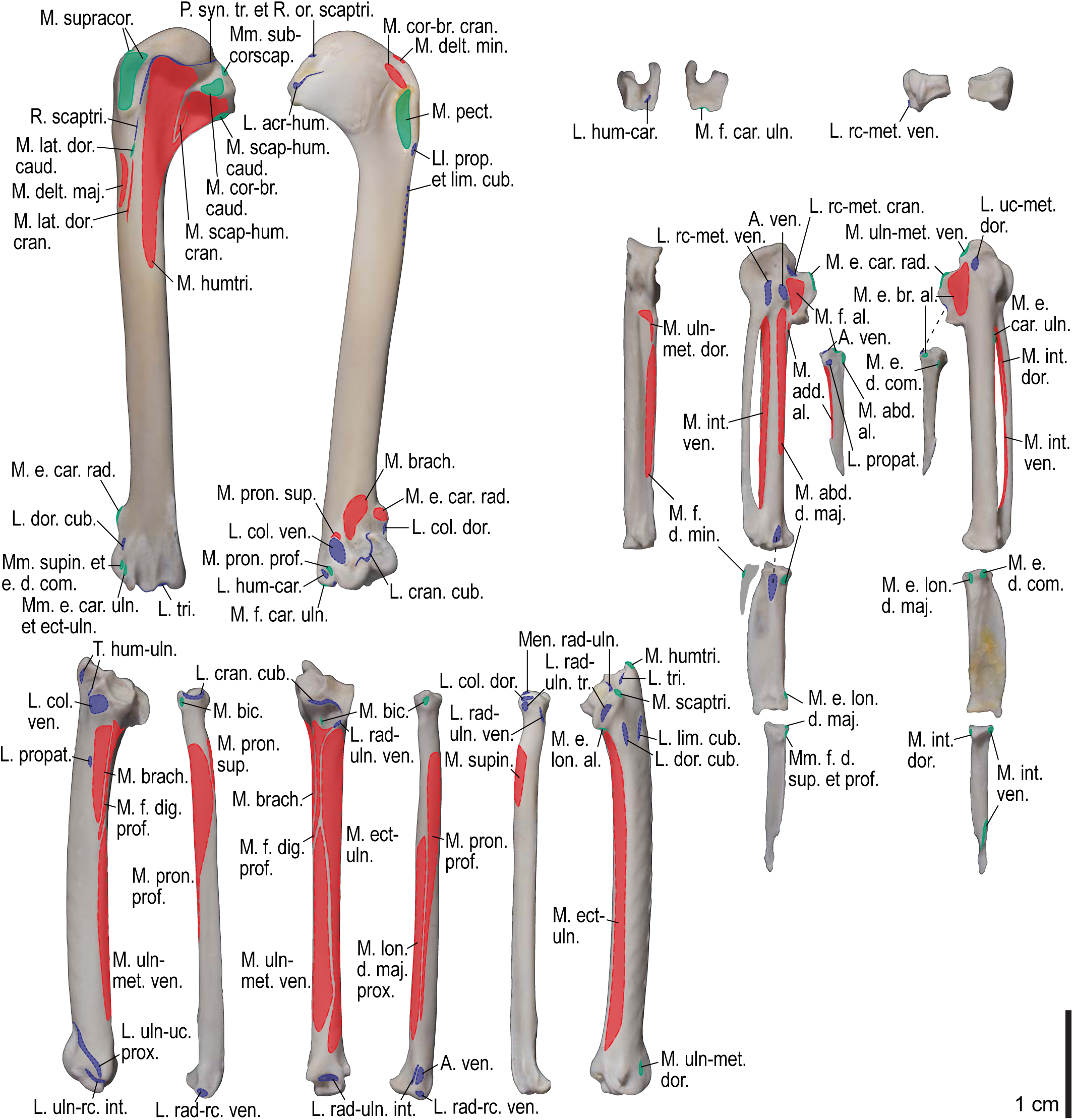
Osteological correlates of major wing muscles and ligaments in *Synthliboramphus antiquus*, wing elements. Drawn on KUGM RA 1311. See Figures S2, S4, and S17 for legends.

**Figure S19.**
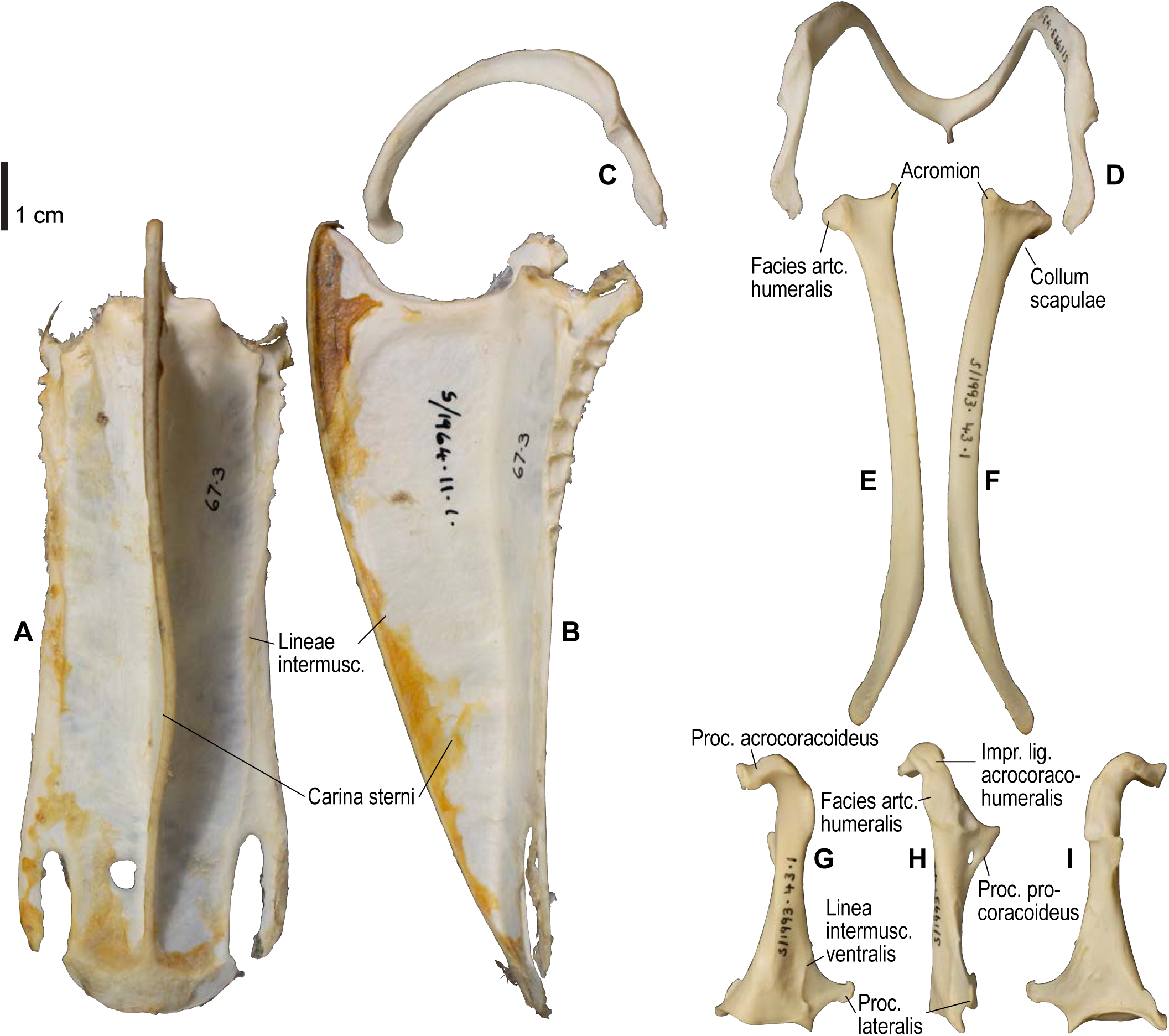
Osteology of *Uria lomvia*, pectoral girdle elements. Drawn on NHMUK S/1964.11.1 (sternum) and S/1993.43.1 (other elements). Sternum in ventral (**A**) and left lateral (**B**) views; furcula in left lateral (**C**) and dorsal (**D**) views; left scapula in lateral (**E**) and medial (**F**) views; left coracoid in ventral (**G**), lateral (**H**) and dorsal (**I**) views. See Figure S1 for abbreviations.

**Figure S20.**
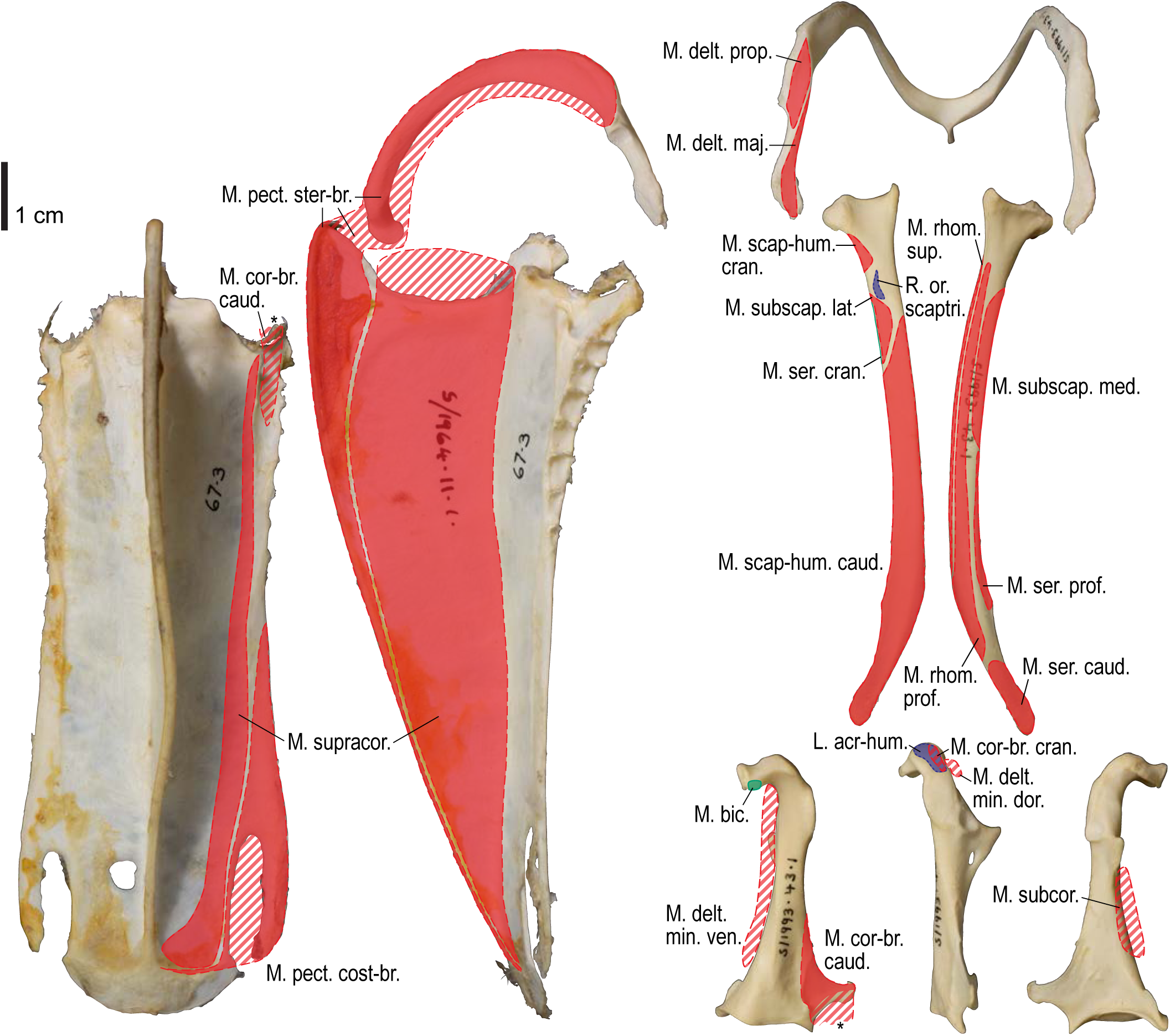
Osteological correlates of major wing muscles and ligaments in *Uria lomvia*, pectoral girdle elements. Drawn on NHMUK S/1964.11.1 (sternum) and S/1993.43.1 (other elements). See Figure S2 for legends.

**Figure S21.**
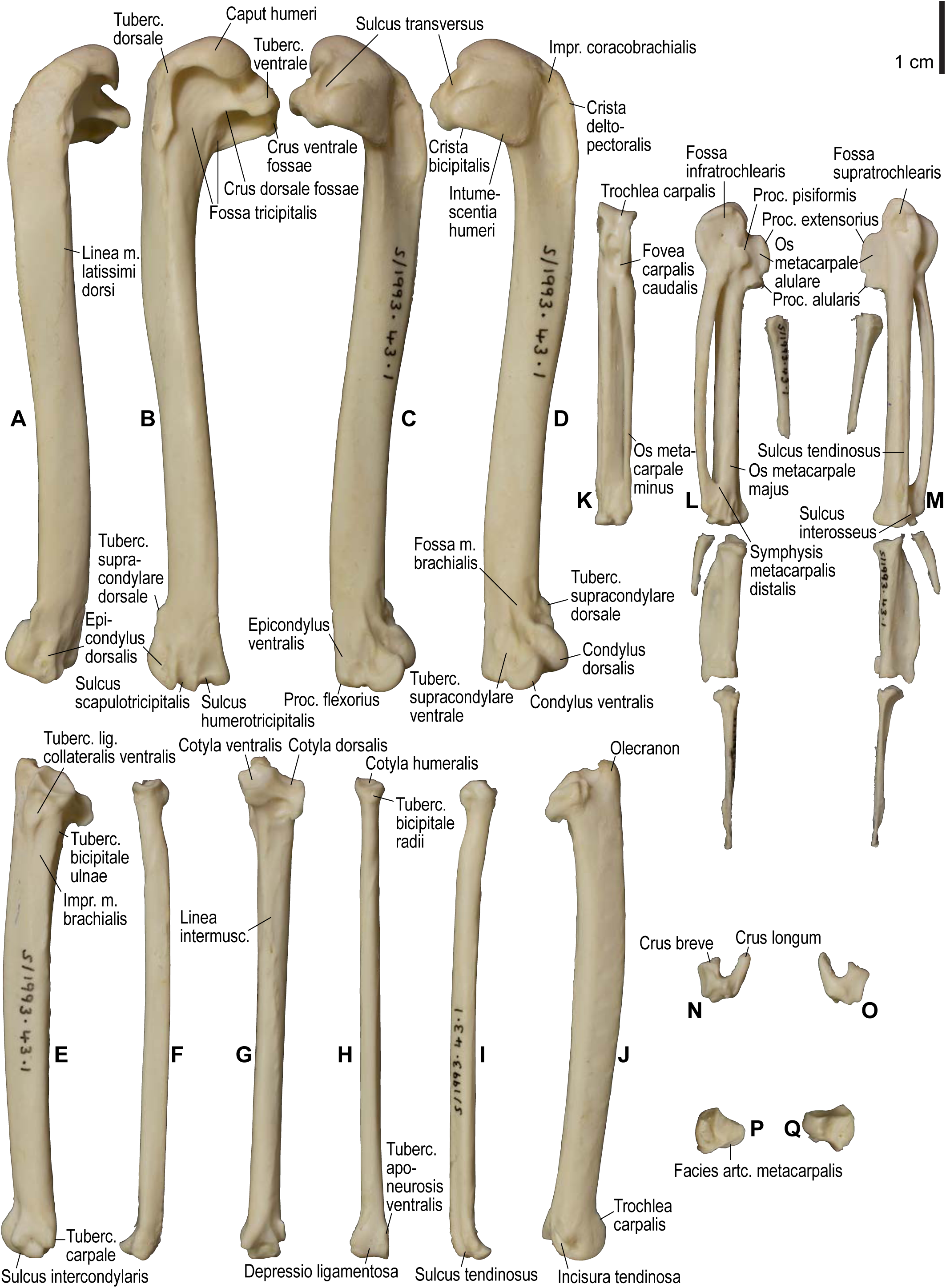
Osteology of *Uria lomvia*, wing elements. Drawn on NHMUK S/1993.43.1. Left humerus in dorsal (**A**), caudal (**B**), ventral (**C**), and cranial (**D**) views; left ulna in ventral (**E**), cranial (**G**), and dorsal (**J**) views; left radius in ventral (**F**), caudal (**H**), and dorsal (**I**) views; left carpometacarpus and phalanges in caudal (**K**; phalanges not shown), ventral (**L**), and dorsal (**M**) views; left ulnare in proximal (**N**) and distal (**O**) views; right radiale (mirrored for comparison) in cranial (**P**) and caudal (**Q**) views. See Figure S1 for abbreviations.

**Figure S22.**
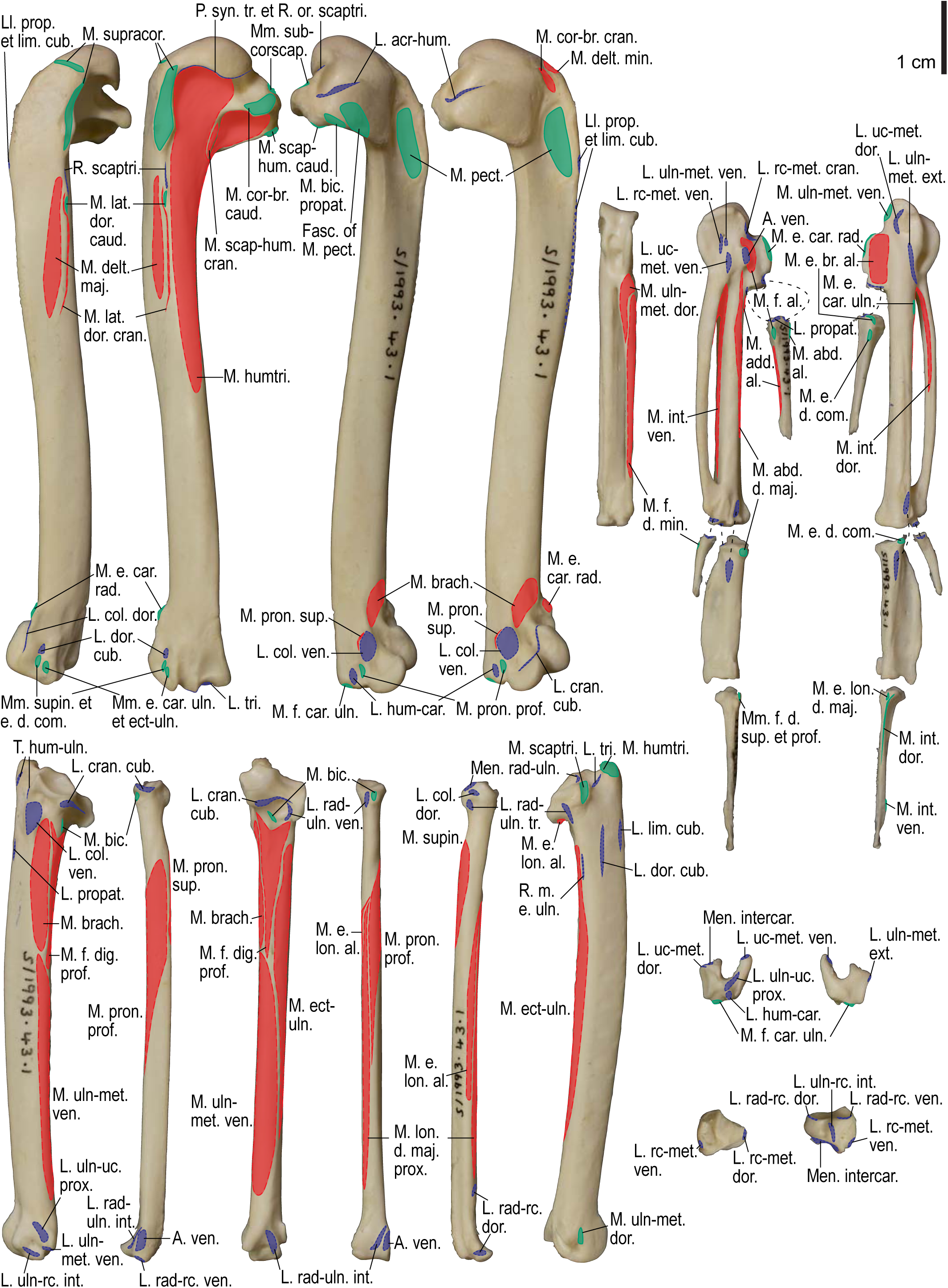
Osteological correlates of major wing muscles and ligaments in *Uria lomvia*, wing elements. Drawn on NHMUK S/1993.43.1. See Figures S2 and S4 for legends.

**Figure S23.**
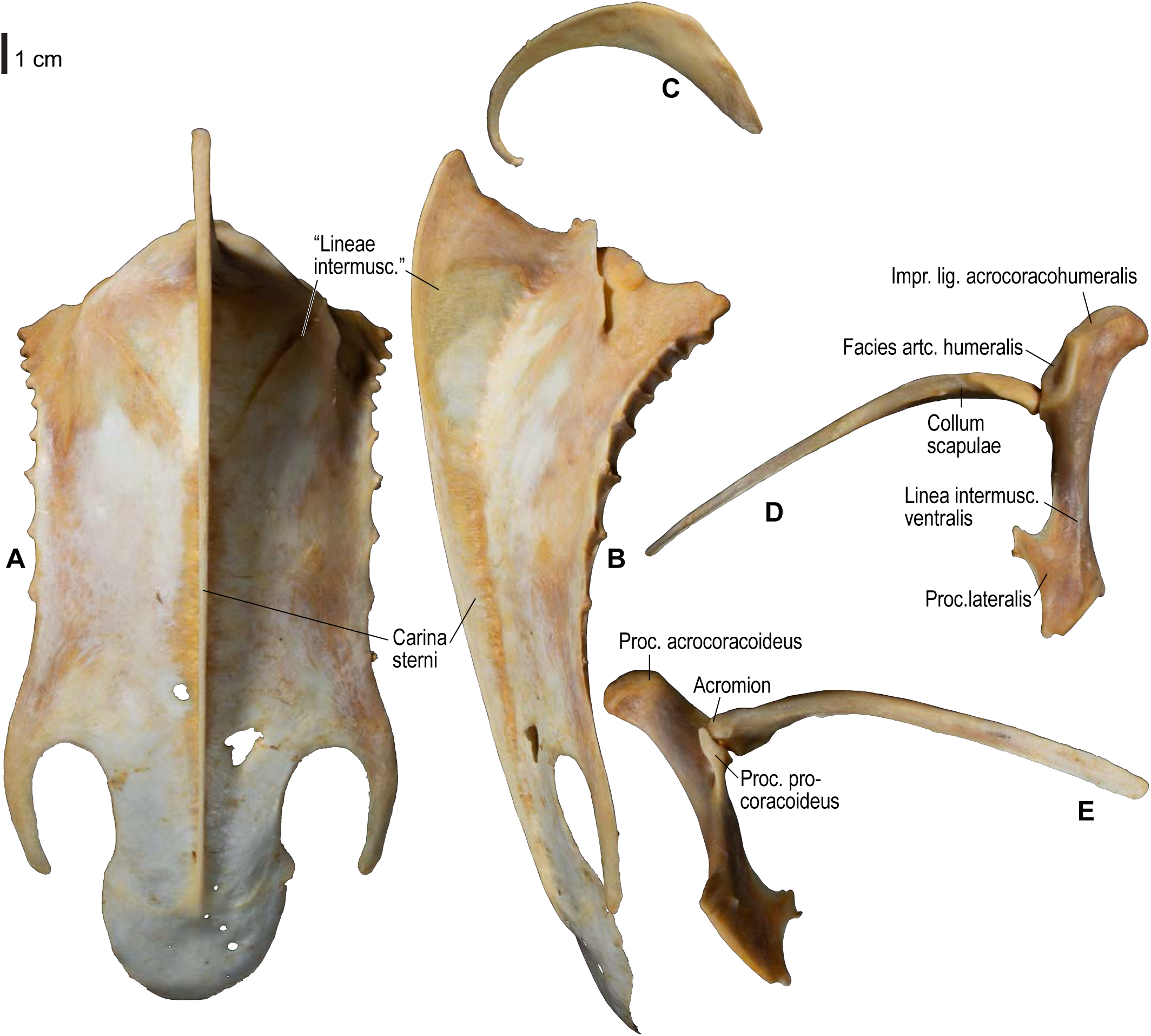
Osteology of *Gavia adamsii*, pectoral girdle elements. Drawn on KUGM RAJ AO14052401. Sternum in ventral (**A**) and left lateral (**B**) views; furcula in left lateral view (**C**); right scapula and coracoid in lateral (**D**) and medial (**E**) views. See Figure S1 for abbreviations.

**Figure S24.**
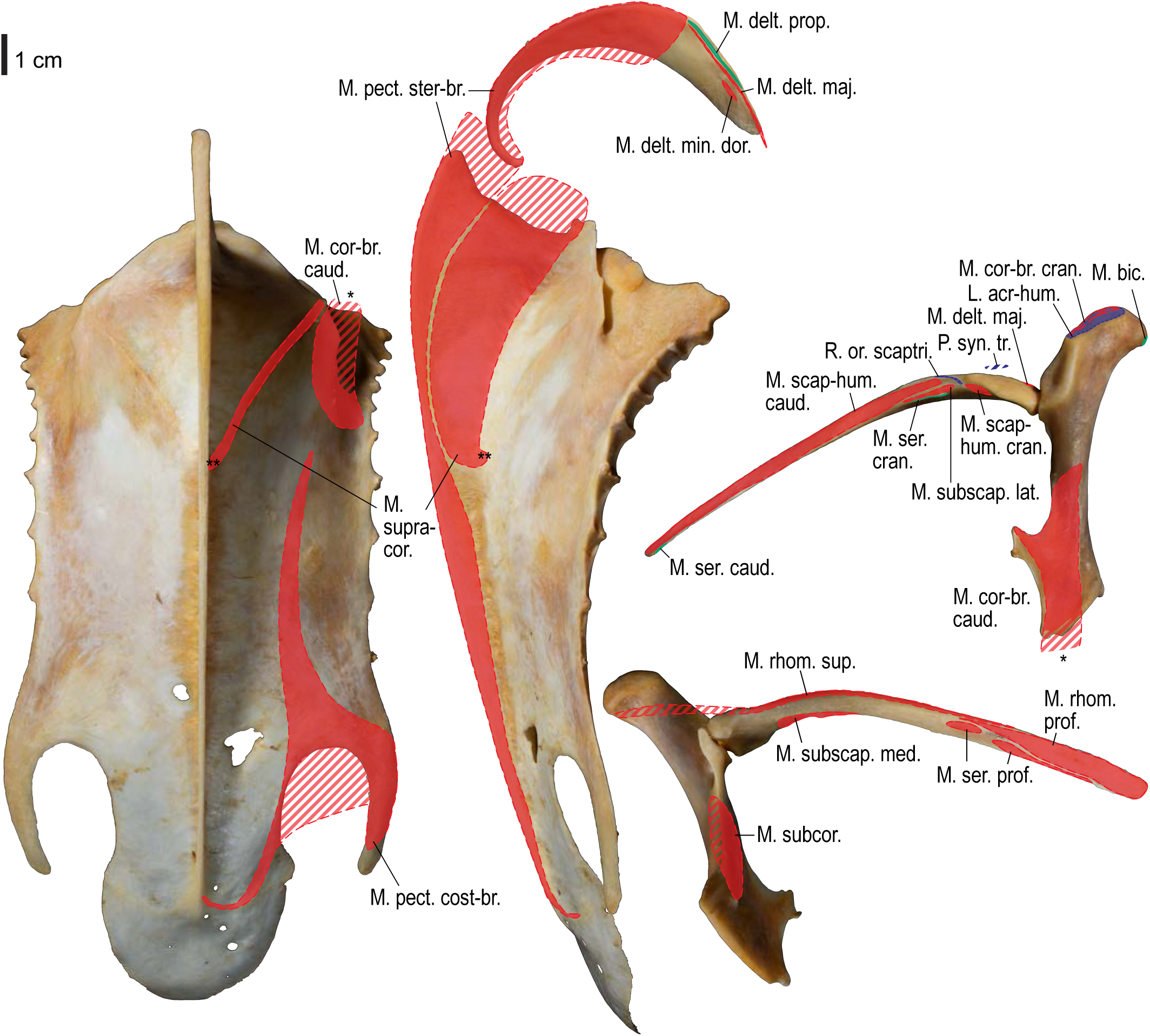
Osteological correlates of major wing muscles and ligaments in *Gavia adamsii*, pectoral girdle elements. Drawn on KUGM RAJ AO14052401. See Figure S2 for legends.

**Figure S25.**
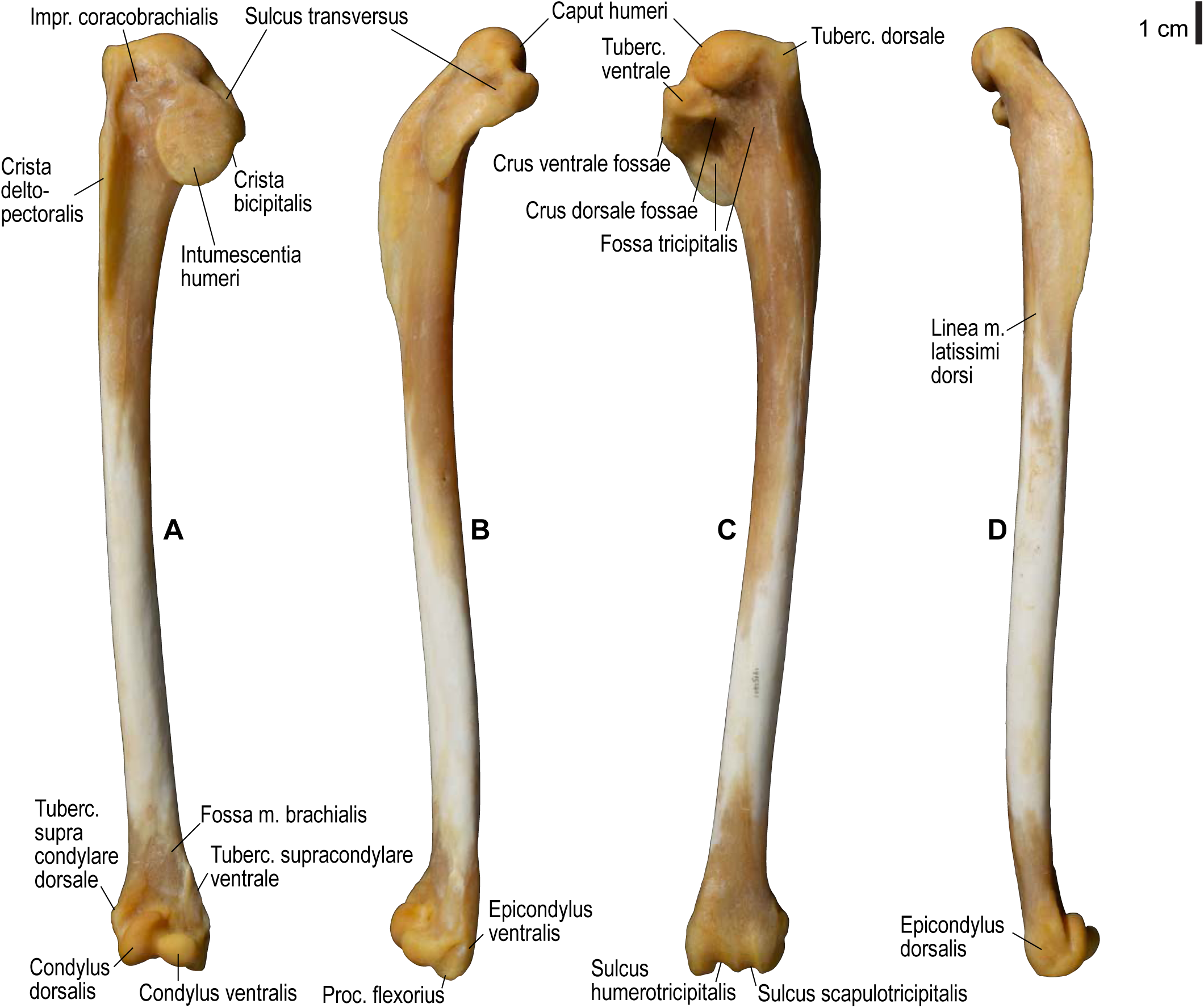
Osteology of *Gavia adamsii*, humerus. Drawn on KUGM RAJ AO14052401. Right humerus in cranial (**A**), ventral (**B**), caudal (**C**), and dorsal (**D**) views. See Figure S1 for abbreviations.

**Figure S26.**
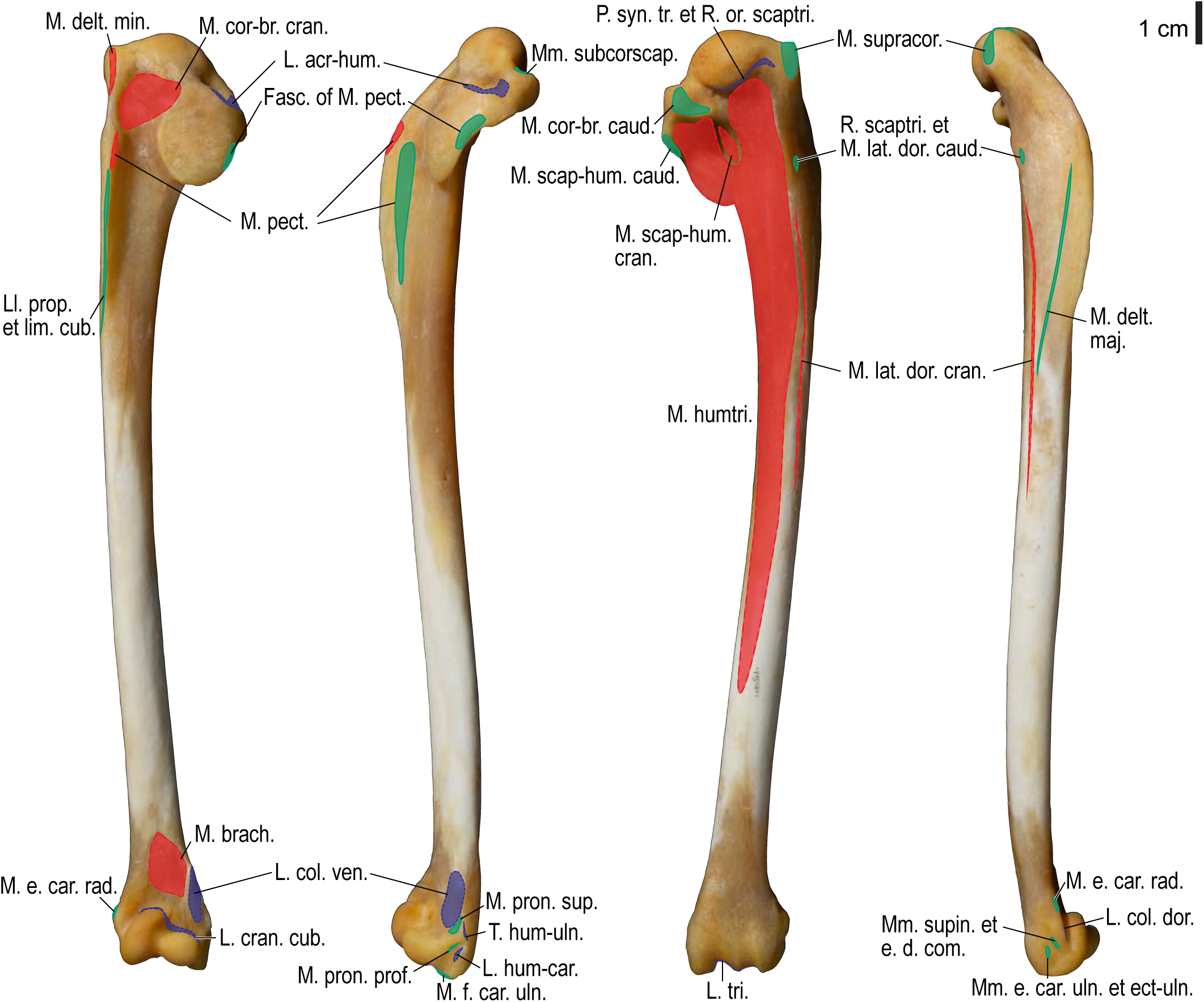
Osteological correlates of major wing muscles and ligaments in *Gavia adamsii*, humerus. Drawn on KUGM RAJ AO14052401. See Figure S2 for legends.

**Figure S27.**
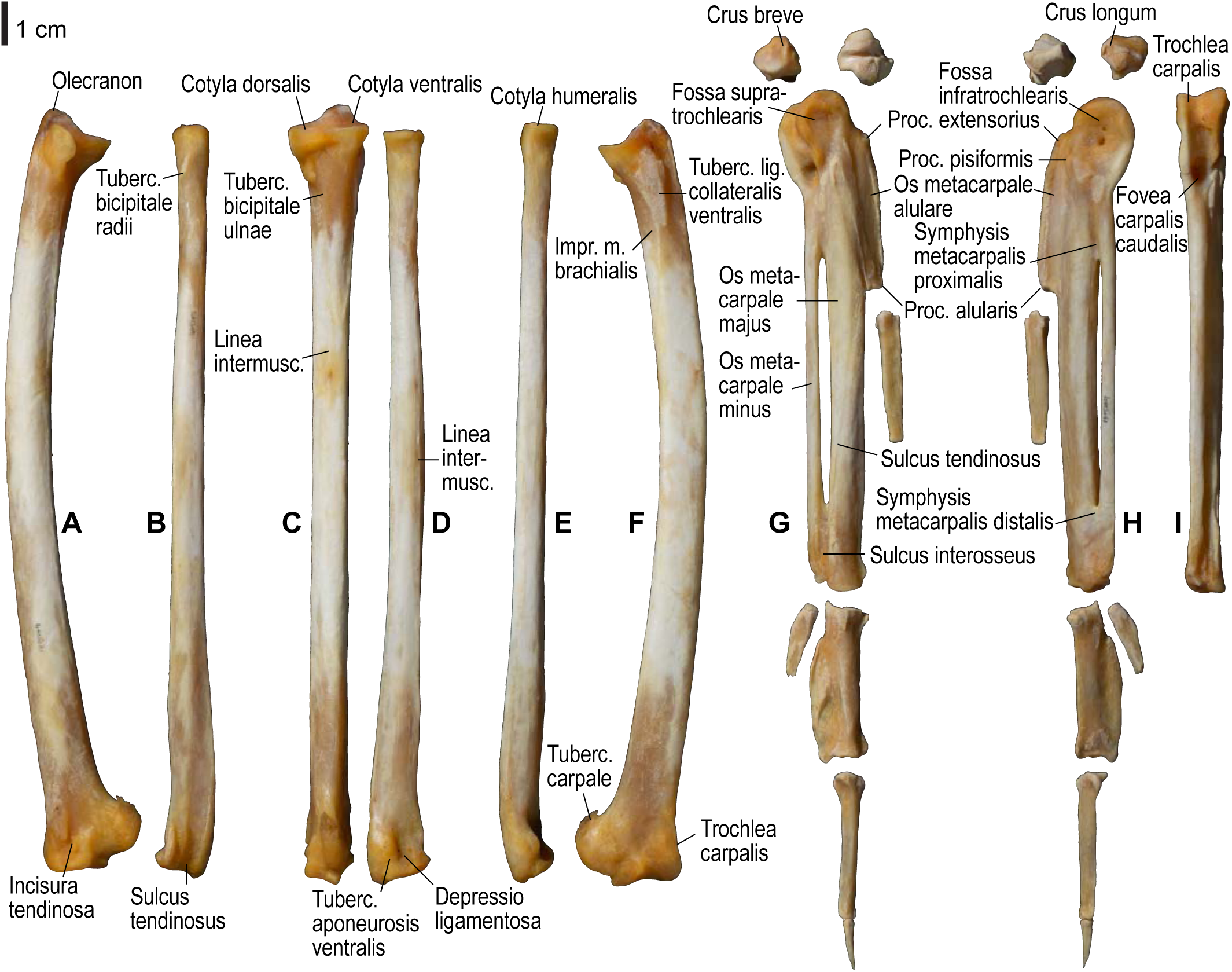
Osteology of *Gavia adamsii*, distal wing elements. Drawn on KUGM RAJ AO14052401. Right ulna in dorsal (**A**), cranial (**C**), and ventral (**F**) views; right radius in dorsal (**B**), caudal (**D**), and ventral (**E**) views; right ulnare, radiale, carpometacarpus, and phalanges in dorsal (**G**), ventral (**H**), and caudal (**I**; carpometacarpus only) views. See Figure S1 for abbreviations.

**Figure S28.**
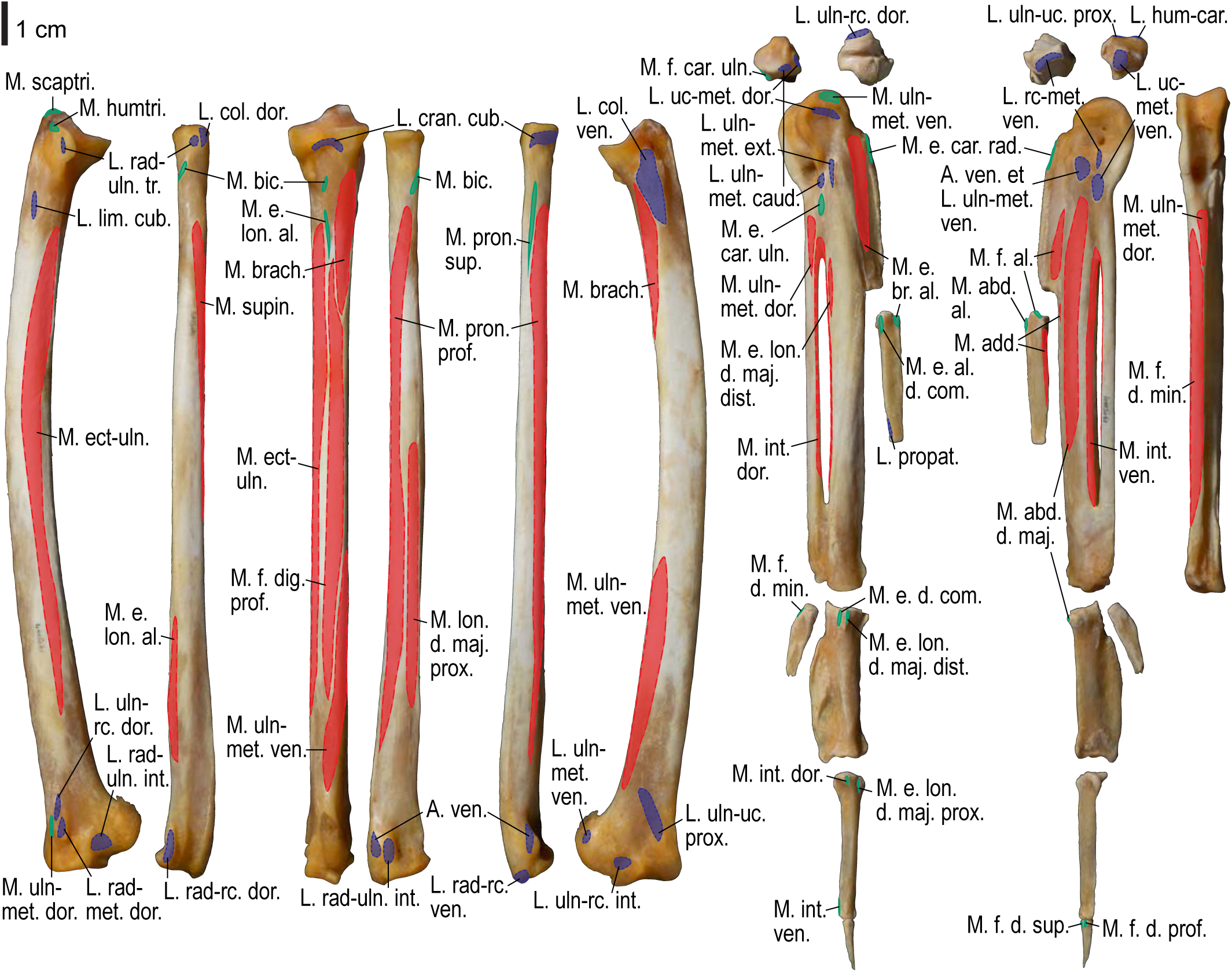
Osteological correlates of major wing muscles and ligaments in *Gavia adamsii*, distal wing elements. Drawn on KUGM RAJ AO14052401. See Figure S2 for legends.

**Figure S29.**
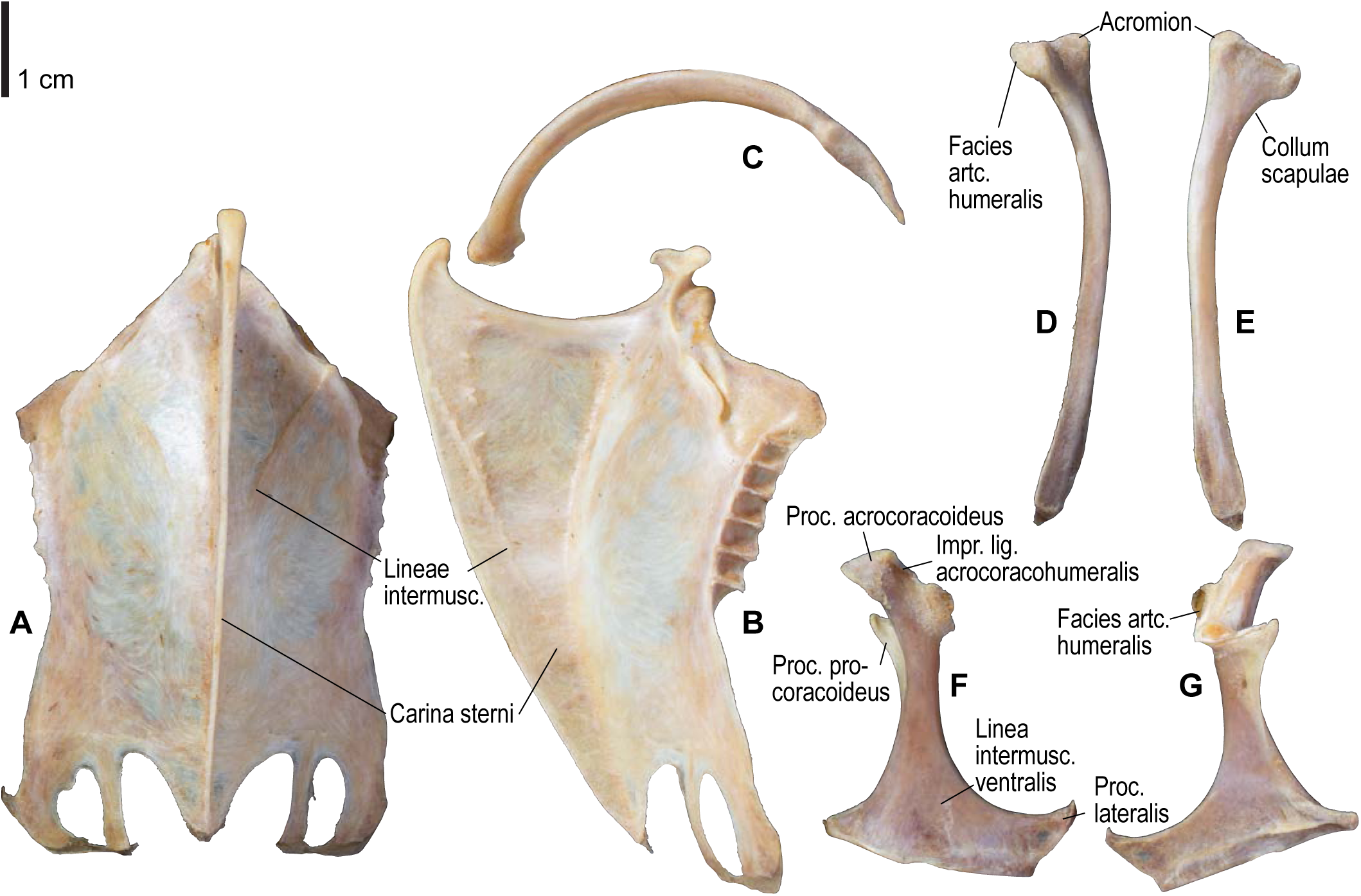
Osteology of *Ardenna tenuirostris*, pectoral girdle elements. Drawn on KUGM RAJ AO09110484. Sternum in ventral (**A**) and left lateral (**B**) views; furcula in left lateral view (**C**); left scapula in lateral (**D**) and medial (**E**) views; left coracoid in ventral (**F**) and dorsal (**G**) views. See Figure S1 for abbreviations.

**Figure S30.**
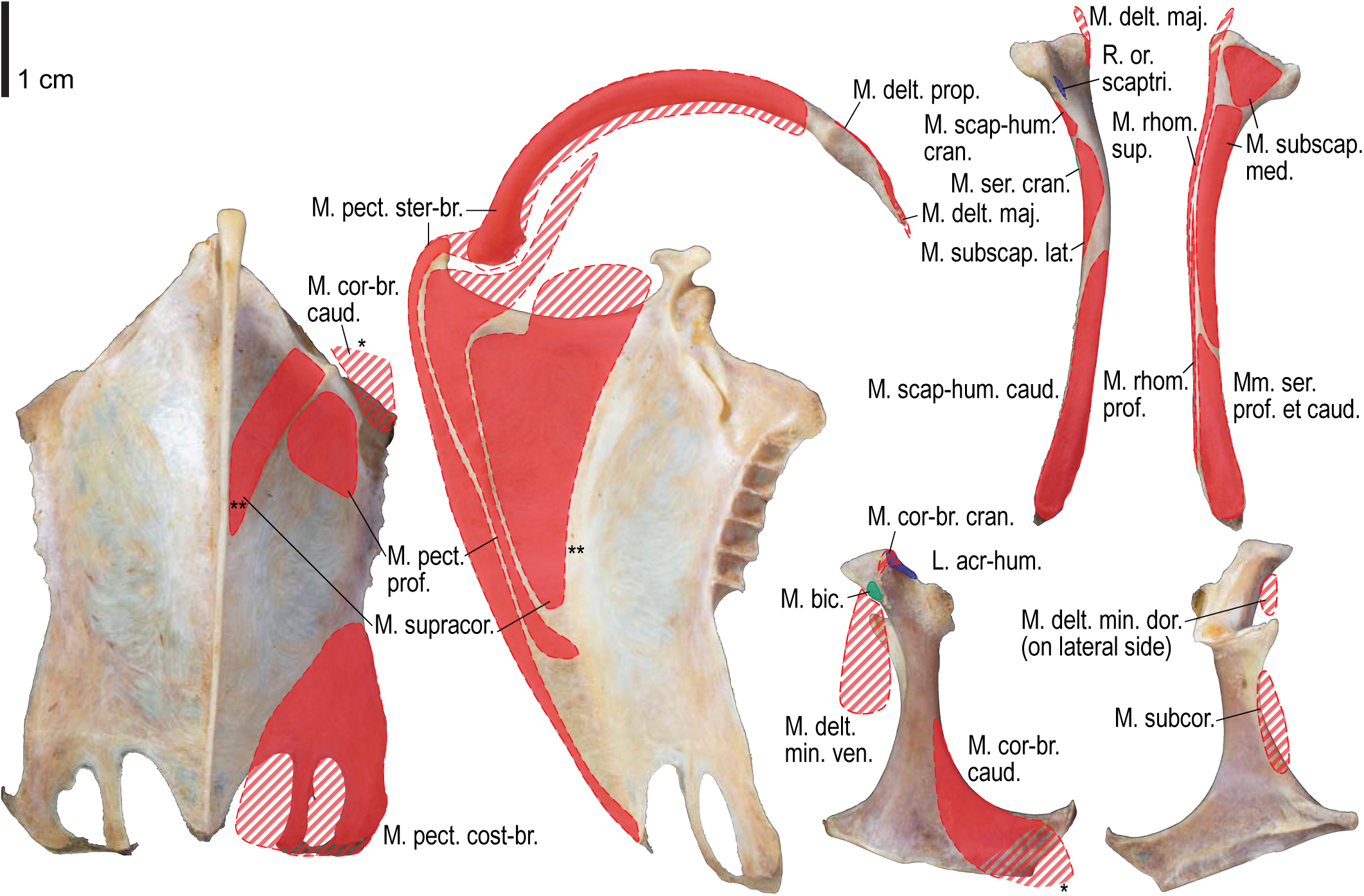
Osteological correlates of major wing muscles and ligaments in *Ardenna tenuirostris*, pectoral girdle elements. Drawn on KUGM RAJ AO09110484. See Figure S2 for legends.

**Figure S31.**
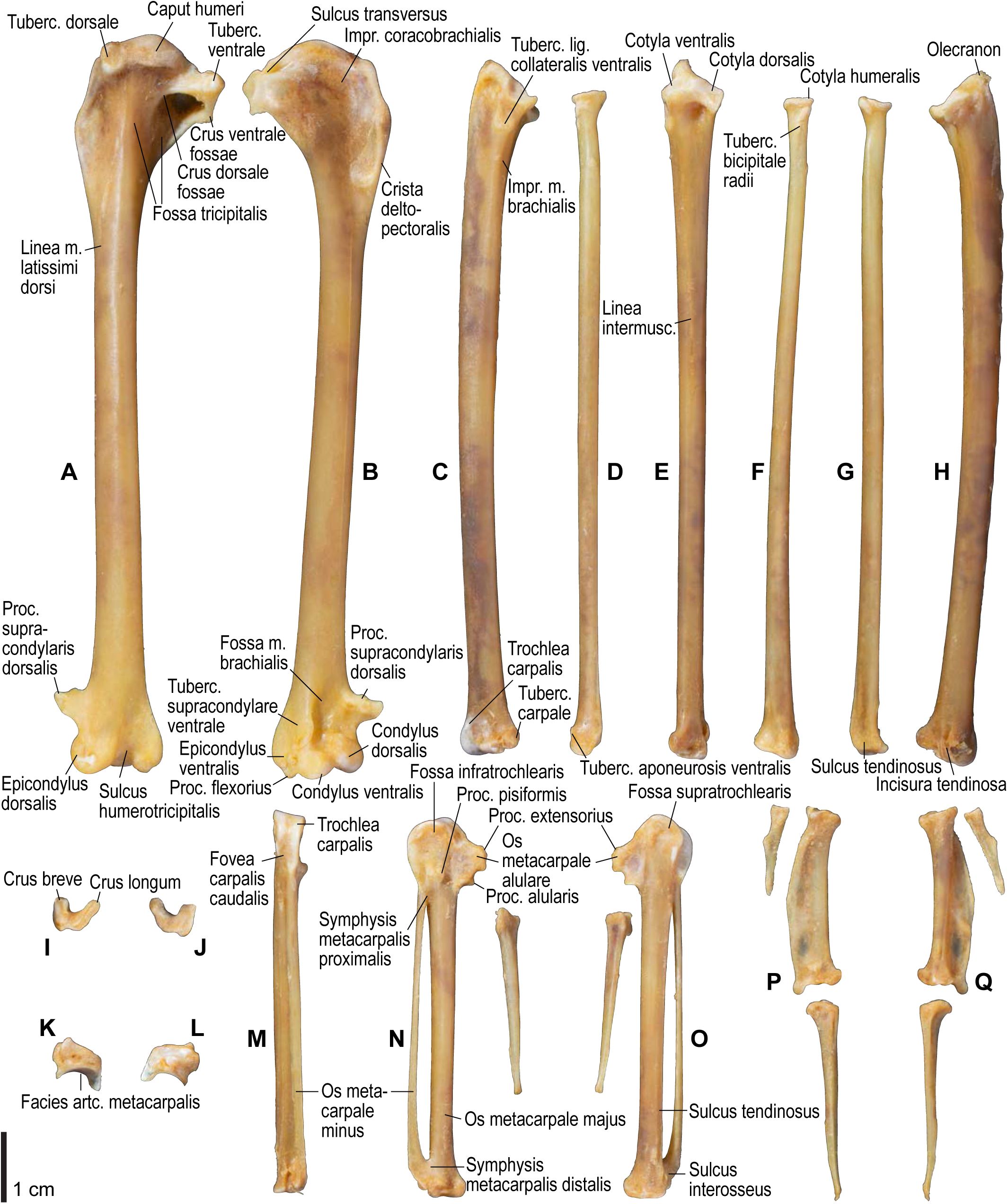
Osteology of *Ardenna tenuirostris*, wing elements. Drawn on KUGM RAJ AO09110484. Left humerus in caudal (**A**) and cranial (**B**) views; left ulna in ventral (**C**), cranial (**E**), and dorsal (**H**) views; left radius in ventral (**D**), caudal (**F**), and dorsal (**G**) views; left ulnare in proximal (**I**) and distal (**J**) views; left radiale in cranial (**K**) and caudal (**L**) views; left carpometacarpus and phalanges in caudal (**M**; phalanges not shown), ventral (**N**, **P**), and dorsal (**O**, **Q**) views. See Figure S1 for abbreviations.

**Figure S32.**
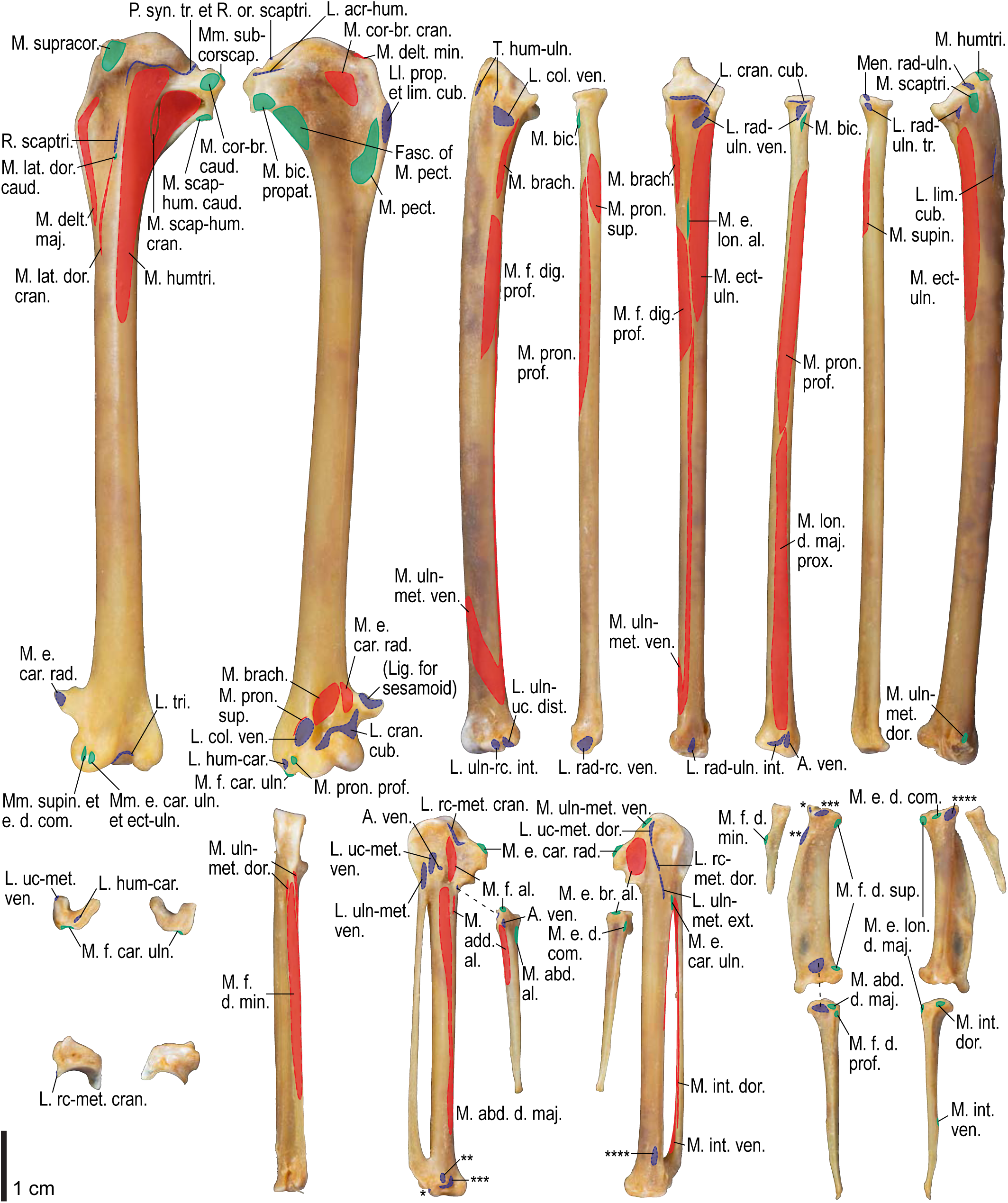
Osteological correlates of major wing muscles and ligaments in *Ardenna tenuirostris*, wing elements. Drawn on KUGM RAJ AO09110484. See Figures S2 and S4 for legends. Correspondence of attachment sites of some ligaments across panels are shown with asterisks.

